## Supplemental Tables for "Identification of a novel modulator of the actin cytoskeleton, mitochondria, nutrient metabolism and lifespan in yeast"

**Supplementary Table 1**

**Downregulated** (significant)

| ID | log2FoldChange | *p* value | padj | WT | *ykl075c*Δ |
|---|---|---|---|---|---|
| *ARO9* | -1.132967918 | 0.0198432 | 0.99979869 | 5654.08208 | 2578.12671 |
| *ILV3* | -0.988520585 | 0.02040871 | 0.99979869 | 147701.676 | 74440.7177 |
| *MAE1* | -0.964816095 | 0.02162205 | 0.99979869 | 90300.4513 | 46264.8169 |
| *ILV5* | -0.956956141 | 0.02181281 | 0.99979869 | 163099.197 | 84019.2529 |
| *CPA2* | -0.926243179 | 0.02414165 | 0.99979869 | 36644.2269 | 19283.1576 |
| *ISU2* | -0.934657685 | 0.02490066 | 0.99979869 | 16864.6948 | 8823.0364 |
| *PHO81* | -0.917420978 | 0.02567054 | 0.99979869 | 18273.3377 | 9674.89457 |
| *ADH5* | -0.91371471 | 0.02695653 | 0.99979869 | 12147.1066 | 6447.87951 |
| *ARN1* | -0.901899438 | 0.02700869 | 0.99979869 | 15219.9773 | 8145.44525 |
| *PHO8* | -0.869847459 | 0.0271153 | 0.99979869 | 49257.6295 | 26953.9816 |
| *BAP2* | -0.859375381 | 0.02732046 | 0.99979869 | 104377.126 | 57531.6919 |
| *snR8* | -1.172352598 | 0.02807933 | 0.99979869 | 1781.28949 | 790.352163 |
| *PDR15* | -0.852804039 | 0.0292684 | 0.99979869 | 22391.96 | 12398.5855 |
| *YKL075C* | -11.33667083 | 0.03020634 | 0.99979869 | 7956.29631 | 3.07530025 |
| *LEU4* | -0.80361025 | 0.0315355 | 0.99979869 | 63202.6093 | 36209.6102 |
| *SIT1* | -0.777667489 | 0.03472474 | 0.99979869 | 26958.3434 | 15725.0353 |
| *SCR1* | -0.730694306 | 0.04153027 | 0.99979869 | 12528.5328 | 7549.8621 |
| *RDT1* | -0.879633238 | 0.04160457 | 0.99979869 | 2549.99492 | 1385.93531 |
| *DRE2* | -0.697804801 | 0.0428953 | 0.99979869 | 25325.3321 | 15613.2993 |
| | -0.730380301 | 0.04479517 | 0.99979869 | 7224.66043 | 4354.62515 |
| *YGL117W* | -0.78387062 | 0.04511035 | 0.99979869 | 3849.38024 | 2235.74328 |
| *PDR5* | -0.656679405 | 0.04563112 | 0.99979869 | 448547.423 | 284529.854 |
| *PHO86* | -0.662221282 | 0.04656403 | 0.99979869 | 36305.7234 | 22941.7398 |
| *HIS4* | -0.646604422 | 0.04757081 | 0.99979869 | 91155.9775 | 58228.7599 |
| *ENB1* | -0.649001298 | 0.04769307 | 0.99979869 | 56569.1107 | 36075.3221 |
| *snR37* | -0.789126897 | 0.04774091 | 0.99979869 | 3059.21349 | 1770.34784 |
| *ILV2* | -0.641585459 | 0.04818287 | 0.99979869 | 103079.691 | 66074.876 |
| *ARG7* | -0.650009152 | 0.04934722 | 0.99979869 | 22474.8787 | 14322.6983 |
| *ARG4* | -0.641630383 | 0.04990071 | 0.99979869 | 28613.7915 | 18341.0907 |
| *YBL029C-A* | -0.685684794 | 0.05039391 | 0.99979869 | 7185.63985 | 4467.38616 |
| *RPS9A* | -0.627858014 | 0.05143214 | 0.99979869 | 35796.5048 | 23165.2116 |
| *RPL41A* | -0.611552767 | 0.05228042 | 0.99979869 | 215663.82 | 141150.131 |
| *snR30* | -0.64649778 | 0.05357696 | 0.99979869 | 9942.44385 | 6351.52011 |
| *TAT1* | -0.607411136 | 0.0538619 | 0.99979869 | 58926.9293 | 38678.0512 |
| *RSB1* | -0.640075459 | 0.05422371 | 0.99979869 | 10470.1972 | 6718.50594 |
| *snR34* | -1.001977535 | 0.05480759 | 0.99979869 | 903.326433 | 451.044036 |
| *GGC1* | -0.61634977 | 0.05537588 | 0.99979869 | 17334.8928 | 11307.879 |
| *ATX1* | -0.636496277 | 0.05559253 | 0.99979869 | 9276.16744 | 5967.10758 |
| *VTC2* | -0.592120934 | 0.05686444 | 0.99979869 | 43931.3203 | 29142.5702 |
| *DAN1* | -1.008196019 | 0.05769053 | 0.99979869 | 816.505642 | 405.939632 |
| *HOM2* | -0.580398717 | 0.05794906 | 0.99979869 | 102150.026 | 68315.7448 |
| *RPL9B* | -0.578552121 | 0.05846595 | 0.99979869 | 83318.694 | 55793.1222 |

| | | | | | |
|---|---|---|---|---|---|
| *FUI1* | -0.586779695 | 0.06140963 | 0.99979869 | 14513.7048 | 9663.61847 |
| *YNL217W* | -0.569496016 | 0.06233414 | 0.99979869 | 25990.633 | 17513.8349 |
| *ARG3* | -0.898692519 | 0.06308403 | 0.99979869 | 982.343108 | 526.901442 |
| *FSF1* | -0.552696325 | 0.06436445 | 0.99979869 | 45707.7322 | 31160.9923 |
| *FET4* | -0.553728631 | 0.06472389 | 0.99979869 | 34901.958 | 23777.1964 |
| *CIT2* | -0.55709209 | 0.0658536 | 0.99979869 | 19700.5155 | 13389.8573 |
| *snR13* | -0.761161468 | 0.06632988 | 0.99979869 | 1554.97012 | 917.464573 |
| *HIS1* | -0.538944665 | 0.06646391 | 0.99979869 | 81031.1124 | 55771.595 |
| *FTR1* | -0.540502682 | 0.06676627 | 0.99979869 | 50851.6202 | 34962.0634 |
| *snR35* | -0.756470089 | 0.07153246 | 0.99979869 | 1355.96516 | 802.653364 |
| *SAM2* | -0.518377746 | 0.0724467 | 0.99979869 | 38626.4724 | 26967.3079 |
| *CPA1* | -0.526170685 | 0.07245569 | 0.99979869 | 21417.421 | 14872.152 |
| *RCL1* | -0.542447265 | 0.07473695 | 0.99979869 | 8525.02127 | 5853.32147 |
| *snR10* | -0.661996124 | 0.07521134 | 0.99979869 | 2043.70289 | 1291.6261 |
| *ASN1* | -0.498395007 | 0.07571696 | 0.99979869 | 150597.978 | 106607.333 |
| *YIR042C* | -0.636288911 | 0.07599752 | 0.99979869 | 2390.01054 | 1537.65012 |
| *ODC2* | -0.530722194 | 0.07608421 | 0.99979869 | 9990.24406 | 6915.32515 |
| *IDH1* | -0.502403122 | 0.07633061 | 0.99979869 | 42139.3001 | 29747.3793 |
| *YBR056W-A* | -0.508147985 | 0.07867005 | 0.99979869 | 15551.6523 | 10934.7426 |
| *ATR1* | -0.531617999 | 0.07889186 | 0.99979869 | 7437.32259 | 5144.97731 |
| *MET22* | -0.503528424 | 0.07981024 | 0.99979869 | 15860.8904 | 11187.9423 |
| *AAC3* | -0.553920991 | 0.08077673 | 0.99979869 | 4459.07681 | 3037.37154 |
| *YMC2* | -0.514351554 | 0.08116162 | 0.99979869 | 9189.34665 | 6433.52811 |
| *snR32* | -0.804822317 | 0.08149737 | 0.99979869 | 872.109968 | 499.22374 |
| *snR42* | -0.733383249 | 0.08181027 | 0.99979869 | 1152.08263 | 692.967655 |
| *HIS5* | -0.486788169 | 0.08309192 | 0.99979869 | 20829.1857 | 14863.9512 |
| *YGR035C* | -0.561212 | 0.08341377 | 0.99979869 | 3563.55449 | 2415.13579 |
| *COT1* | -0.491241644 | 0.08479077 | 0.99979869 | 12838.7464 | 9133.64173 |
| *ECL1* | -0.485779205 | 0.08874641 | 0.99979869 | 9854.64754 | 7037.31206 |
| *BDH2* | -0.461225945 | 0.0901993 | 0.99979869 | 25136.0823 | 18258.0576 |
| *SFG1* | -0.462421447 | 0.09152972 | 0.99979869 | 17902.6422 | 12993.1435 |
| *YPR145C-A* | -0.524299802 | 0.09179596 | 0.99979869 | 3834.74752 | 2666.28531 |
| *FCY2* | -0.437585451 | 0.09573666 | 0.99979869 | 62140.274 | 45882.4546 |
| *FET3* | -0.433586133 | 0.09658551 | 0.99979869 | 92665.0984 | 68610.9736 |
| *snR57* | -0.670578431 | 0.09720055 | 0.99979869 | 1096.4783 | 688.867255 |
| *snR128* | -0.678918337 | 0.09745934 | 0.99979869 | 1048.67809 | 655.038952 |
| *PCL5* | -0.446980472 | 0.09811279 | 0.99979869 | 14879.5228 | 10915.2657 |
| *RPL41B* | -0.427255228 | 0.09862812 | 0.99979869 | 122395.854 | 91022.7367 |
| *TPI1* | -0.423152984 | 0.09937501 | 0.99979869 | 788863.463 | 588328.514 |
| *YPL080C* | -0.803314046 | 0.09981292 | 0.99979869 | 624.329284 | 357.759929 |
| *DLD3* | -0.428624241 | 0.10038424 | 0.99979869 | 36863.7177 | 27388.624 |
| *RPR1* | -0.528499771 | 0.10050292 | 0.99979869 | 2629.01159 | 1822.62795 |
| *CMK2* | -0.419518523 | 0.10159778 | 0.99979869 | 131283.767 | 98157.4332 |
| *snR17a* | -0.610923167 | 0.10227829 | 0.99979869 | 1351.08759 | 884.661371 |
| *HOM3* | -0.424130839 | 0.10313743 | 0.99979869 | 28346.5005 | 21126.2876 |

| | | | | | |
|---|---|---|---|---|---|
| *PRM4* | -0.450733846 | 0.10369562 | 0.99979869 | 7653.88681 | 5600.12175 |
| *FAL1* | -0.48178056 | 0.10433375 | 0.99979869 | 4023.99734 | 2881.55633 |
| *FIT1* | -0.551684204 | 0.10509698 | 0.99979869 | 1882.743 | 1284.4504 |
| *YOR302W* | -0.590271172 | 0.10604185 | 0.99979869 | 1410.59398 | 936.941475 |
| *ARO10* | -0.461962077 | 0.10699737 | 0.99979869 | 4941.95649 | 3587.85029 |
| *INO2* | -0.476706921 | 0.10886836 | 0.99979869 | 3613.30573 | 2596.57851 |
| *RPL7B* | -0.405739116 | 0.10910979 | 0.99979869 | 46690.0753 | 35243.9659 |
| *snR61* | -0.603569983 | 0.11308811 | 0.99979869 | 1135.49889 | 747.29796 |
| *FRE6* | -0.409009233 | 0.11332301 | 0.99979869 | 15120.4748 | 11387.8368 |
| *snR190* | -1.003920656 | 0.11345588 | 0.99979869 | 306.311555 | 152.739912 |
| *GPP1* | -0.390377153 | 0.11368389 | 0.99979869 | 302124.647 | 230499.904 |
| *YOR192C-B* | -0.438879611 | 0.11527319 | 0.99979869 | 5159.49622 | 3806.1966 |
| *YOR385W* | -0.393176914 | 0.11592892 | 0.99979869 | 33475.7558 | 25490.1386 |
| *snR82* | -0.517898871 | 0.11602677 | 0.99979869 | 1893.47366 | 1322.37911 |
| *YHI9* | -0.41612226 | 0.11690305 | 0.99979869 | 8227.48934 | 6165.97699 |
| *ATG5* | -0.410294925 | 0.1170388 | 0.99979869 | 9912.2029 | 7458.6282 |
| *REG2* | -0.571489308 | 0.11734386 | 0.99979869 | 1261.34026 | 848.782868 |
| *PDR3* | -0.400730572 | 0.11797525 | 0.99979869 | 13705.0033 | 10381.1885 |
| *NSR1* | -0.382086057 | 0.11849646 | 0.99979869 | 108138.71 | 82977.7512 |
| *MPC2* | -0.384949655 | 0.11858086 | 0.99979869 | 49711.2437 | 38069.1417 |
| *ERG4* | -0.380466534 | 0.11971534 | 0.99979869 | 81869.0794 | 62890.9151 |
| *SRG1* | -0.413206321 | 0.11999619 | 0.99979869 | 7335.86909 | 5508.88784 |
| *ISU1* | -0.382309942 | 0.12356117 | 0.99979869 | 21320.845 | 16357.522 |
| *NRG2* | -0.396442858 | 0.12383846 | 0.99979869 | 9729.78168 | 7391.99669 |
| *IDH2* | -0.373649101 | 0.12392324 | 0.99979869 | 57812.8917 | 44621.5815 |
| *snR5* | -0.588655478 | 0.12693659 | 0.99979869 | 969.661419 | 644.787951 |
| *FAF1* | -0.387433381 | 0.12737942 | 0.99979869 | 10517.9974 | 8040.88504 |
| *MET16* | -0.440192449 | 0.12839943 | 0.99979869 | 3100.1851 | 2284.94808 |
| *INM1* | -0.3805267 | 0.12882359 | 0.99979869 | 12681.6886 | 9741.52608 |
| *FRE1* | -0.363398642 | 0.12897613 | 0.99979869 | 71408.6374 | 55508.1443 |
| *HGH1* | -0.373298776 | 0.12999193 | 0.99979869 | 16912.495 | 13056.6997 |
| *SDT1* | -0.394510933 | 0.13055912 | 0.99979869 | 6749.58487 | 5134.72631 |
| *OAZ1* | -0.433035077 | 0.13066228 | 0.99979869 | 3183.10383 | 2357.73019 |
| *GTO3* | -0.494440907 | 0.13105873 | 0.99979869 | 1656.42363 | 1175.78979 |
| *TMT1* | -0.40561327 | 0.13134391 | 0.99979869 | 4946.83406 | 3734.4396 |
| *RPS23B* | -0.355363401 | 0.1320871 | 0.99979869 | 230759.907 | 180378.661 |
| *YLR126C* | -0.394957096 | 0.13238813 | 0.99979869 | 6023.80208 | 4581.17227 |
| *MET14* | -0.375292083 | 0.13486636 | 0.99979869 | 9640.03435 | 7431.97559 |
| *JJJ3* | -0.480793544 | 0.13486737 | 0.99979869 | 1726.66068 | 1237.2958 |
| *HIS7* | -0.359254623 | 0.1350893 | 0.99979869 | 25251.193 | 19684.9969 |
| *TCA17* | -0.478956765 | 0.13542494 | 0.99979869 | 1734.46479 | 1244.4715 |
| *RRP45* | -0.365456501 | 0.13659725 | 0.99979869 | 13158.7152 | 10214.0972 |
| *YGR018C* | -0.440267687 | 0.13955238 | 0.99979869 | 2315.87144 | 1706.79164 |
| *RPS12* | -0.341534745 | 0.13995368 | 0.99979869 | 446773.938 | 352594.449 |
| *ATF2* | -0.369490733 | 0.1400837 | 0.99979869 | 8525.99678 | 6599.59433 |

| | | | | | |
|---|---|---|---|---|---|
| *VIP1* | -0.348703204 | 0.14010177 | 0.99979869 | 31477.9021 | 24719.2634 |
| *BUD23* | -0.353276969 | 0.14069341 | 0.99979869 | 18453.8079 | 14445.7104 |
| *ARO4* | -0.342003781 | 0.14151246 | 0.99979869 | 72340.2537 | 57072.4471 |
| *GCD10* | -0.357238253 | 0.14166804 | 0.99979869 | 12976.294 | 10130.039 |
| *YOL014W* | -0.348583746 | 0.14205036 | 0.99979869 | 22774.3617 | 17885.9462 |
| *AFT1* | -0.360695436 | 0.14258601 | 0.99979869 | 10351.1844 | 8061.38704 |
| *YDR261W-B* | -0.385960964 | 0.14380109 | 0.99979869 | 4620.0367 | 3535.57018 |
| *RLP7* | -0.34323571 | 0.14383757 | 0.99979869 | 31062.3329 | 24485.5406 |
| *PGK1* | -0.335026294 | 0.14390407 | 0.99979869 | 1772238.66 | 1404976.54 |
| *MTM1* | -0.354796597 | 0.14415419 | 0.99979869 | 12098.3309 | 9460.64865 |
| *SMF3* | -0.346242983 | 0.1459663 | 0.99979869 | 16994.4382 | 13368.3302 |
| *RCR1* | -0.34038753 | 0.1463186 | 0.99979869 | 27335.8675 | 21590.6579 |
| *snR18* | -0.4834756 | 0.14632502 | 0.99979869 | 1371.5734 | 981.020778 |
| *CIA2* | -0.363286628 | 0.14648537 | 0.99979869 | 7355.37938 | 5718.00826 |
| *EFG1* | -0.358538953 | 0.14695235 | 0.99979869 | 8439.17599 | 6582.16763 |
| *LSO1* | -0.917497405 | 0.14727886 | 0.99979869 | 255.584801 | 135.313211 |
| *snR49* | -0.400309691 | 0.14757004 | 0.99979869 | 3175.29972 | 2405.90989 |
| *snR62* | -0.621567242 | 0.14834763 | 0.99979869 | 621.40274 | 403.889432 |
| *QDR3* | -0.34442216 | 0.14900695 | 0.99979869 | 13853.2815 | 10911.1653 |
| *ORT1* | -0.364675871 | 0.1490262 | 0.99979869 | 6206.22329 | 4820.02058 |
| *CCC2* | -0.356318571 | 0.1498209 | 0.99979869 | 7724.12386 | 6033.73908 |
| *snR4* | -0.371403787 | 0.15300846 | 0.99979869 | 4516.63216 | 3491.49088 |
| *IRC10* | -0.381647342 | 0.15420442 | 0.99979869 | 3564.53001 | 2735.99212 |
| *HTA1* | -0.321710891 | 0.15530518 | 0.99979869 | 74035.6979 | 59237.4584 |
| *LOC1* | -0.332164432 | 0.15633727 | 0.99979869 | 15283.3858 | 12140.2603 |
| *BCP1* | -0.328716619 | 0.1575168 | 0.99979869 | 17386.595 | 13843.9766 |
| *snR86* | -0.3289192 | 0.15751841 | 0.99979869 | 17117.353 | 13627.6805 |
| *MRX14* | -0.348892337 | 0.15768577 | 0.99979869 | 6633.49864 | 5208.53352 |
| *CTR2* | -0.342066911 | 0.15834099 | 0.99979869 | 8136.7665 | 6419.17671 |
| *SUP17* | -0.995237127 | 0.15871508 | 0.99979869 | 194.127387 | 97.3845078 |
| *YNL034W* | -0.586853131 | 0.15882699 | 0.99979869 | 638.962002 | 425.416534 |
| *DBP2* | -0.315027857 | 0.15986878 | 0.99979869 | 86770.0643 | 69748.8347 |
| *TRM1* | -0.319180421 | 0.16010246 | 0.99979869 | 32220.2686 | 25825.3464 |
| *TMA23* | -0.344059154 | 0.16040085 | 0.99979869 | 6805.18919 | 5361.27343 |
| *SRY1* | -0.339565098 | 0.16123053 | 0.99979869 | 7605.11109 | 6010.16178 |
| *RRG8* | -0.38220908 | 0.16167458 | 0.99979869 | 2899.22911 | 2224.46718 |
| *APT2* | -0.352641839 | 0.16211798 | 0.99979869 | 4997.56081 | 3913.83211 |
| *CMS1* | -0.332114554 | 0.16263761 | 0.99979869 | 9433.22527 | 7493.4816 |
| *ERG29* | -0.338096546 | 0.16366377 | 0.99979869 | 7095.89252 | 5613.44805 |
| *UGA3* | -0.348855566 | 0.16463003 | 0.99979869 | 5009.26699 | 3933.30901 |
| *RPL30* | -0.306591626 | 0.16495731 | 0.99979869 | 246792.488 | 199543.932 |
| *snR189* | -0.645535412 | 0.16533619 | 0.99979869 | 468.246963 | 299.329224 |
| *snR33* | -0.344760908 | 0.16622091 | 0.99979869 | 5242.41496 | 4128.07803 |
| *YER138W-A* | -0.353766358 | 0.16639635 | 0.99979869 | 4222.02678 | 3303.89756 |
| *HUG1* | -0.47494077 | 0.16730578 | 0.99979869 | 1064.28633 | 765.749761 |

| | | | | | |
|---|---|---|---|---|---|
| *HES1* | -0.506164243 | 0.16772709 | 0.99979869 | 869.183425 | 611.984749 |
| *DOM34* | -0.326968067 | 0.16775337 | 0.99979869 | 8714.27108 | 6947.10325 |
| *CIC1* | -0.309778871 | 0.16963677 | 0.99979869 | 22008.5828 | 17755.7585 |
| *ADH3* | -0.302064261 | 0.16989122 | 0.99979869 | 87142.7108 | 70680.6506 |
| *NOC2* | -0.306043348 | 0.16995641 | 0.99979869 | 33285.5305 | 26923.2285 |
| *PAN5* | -0.31251279 | 0.17046293 | 0.99979869 | 15856.9883 | 12768.6466 |
| *REI1* | -0.314328763 | 0.17106694 | 0.99979869 | 13289.4341 | 10687.6935 |
| *QDR2* | -0.306248606 | 0.17159214 | 0.99979869 | 24758.5582 | 20023.2799 |
| *SER33* | -0.307334551 | 0.17175907 | 0.99979869 | 21541.3113 | 17408.2496 |
| *TDH2* | -0.296771907 | 0.1723718 | 0.99979869 | 937559.188 | 763240.316 |
| *LYS20* | -0.300024266 | 0.17238484 | 0.99979869 | 62294.4053 | 50597.9149 |
| *snR67* | -0.463347102 | 0.17249986 | 0.99979869 | 1076.96801 | 781.126262 |
| *MTD1* | -0.30778708 | 0.17296068 | 0.99979869 | 18004.0957 | 14545.1451 |
| *VMR1* | -0.310147836 | 0.17381603 | 0.99979869 | 13928.3961 | 11234.0718 |
| *RSA3* | -0.310182426 | 0.17419957 | 0.99979869 | 13469.9043 | 10864.0107 |
| *SSF1* | -0.310397235 | 0.17464835 | 0.99979869 | 12829.9668 | 10346.3351 |
| *OCA2* | -0.317743061 | 0.17482398 | 0.99979869 | 8716.22211 | 6993.23276 |
| *HTA2* | -0.297154591 | 0.17520265 | 0.99979869 | 54435.6605 | 44302.7753 |
| *BMT6* | -0.357918334 | 0.17567804 | 0.99979869 | 3042.62974 | 2374.13179 |
| *snR84* | -0.381875784 | 0.1758316 | 0.99979869 | 2155.88706 | 1654.51153 |
| *snR38* | -0.585479803 | 0.1766248 | 0.99979869 | 530.679891 | 353.659528 |
| *PEX13* | -0.32915757 | 0.17755041 | 0.99979869 | 5269.72936 | 4194.70953 |
| *ADH1* | -0.289662918 | 0.17833891 | 0.99979869 | 2504230.61 | 2048693.27 |
| *LYS21* | -0.295048623 | 0.17927693 | 0.99979869 | 32346.11 | 26363.5239 |
| *CTF19* | -0.354704915 | 0.17938582 | 0.99979869 | 2944.10278 | 2302.37478 |
| *YPL088W* | -0.338037966 | 0.18088627 | 0.99979869 | 3852.30678 | 3047.62254 |
| *MIG3* | -0.320712718 | 0.18119013 | 0.99979869 | 5880.40144 | 4708.28468 |
| *YDL241W* | -0.332182104 | 0.18126436 | 0.99979869 | 4332.25992 | 3441.26097 |
| *FYV7* | -0.316117081 | 0.18140222 | 0.99979869 | 6769.09516 | 5437.13083 |
| *YPR014C* | -0.419024048 | 0.1827772 | 0.99979869 | 1319.87113 | 987.171379 |
| *DFR1* | -0.316196527 | 0.18302979 | 0.99979869 | 6312.55437 | 5070.145 |
| *snR63* | -0.335677954 | 0.1836125 | 0.99979869 | 3764.51048 | 2983.04124 |
| *RUF20* | -0.370692332 | 0.18565438 | 0.99979869 | 2072.96833 | 1603.25653 |
| *ACS2* | -0.284326438 | 0.18581447 | 0.99979869 | 68537.6982 | 56277.9945 |
| *NOP15* | -0.290904901 | 0.1860479 | 0.99979869 | 20805.7734 | 17006.4104 |
| *GCV1* | -0.292507289 | 0.18728211 | 0.99979869 | 15904.7885 | 12985.9678 |
| *YHM2* | -0.287377992 | 0.18740159 | 0.99979869 | 26318.4059 | 21565.0304 |
| *PNO1* | -0.286534101 | 0.18783284 | 0.99979869 | 27664.6159 | 22681.3644 |
| *FRE3* | -0.322369579 | 0.1882352 | 0.99979869 | 4450.29718 | 3559.14748 |
| *UTP9* | -0.291981212 | 0.18864196 | 0.99979869 | 14746.8528 | 12044.926 |
| *YLR157C-B* | -0.559930511 | 0.18939321 | 0.99979869 | 522.875775 | 354.684628 |
| *OAC1* | -2.875904137 | 0.18982934 | 0.99979869 | 32839.7203 | 4473.53676 |
| *YPR011C* | -0.296232576 | 0.18994669 | 0.99979869 | 10357.0375 | 8434.52347 |
| *RRP1* | -0.290569212 | 0.19018518 | 0.99979869 | 14511.7538 | 11864.5083 |
| *NCB2* | -0.292880225 | 0.19022866 | 0.99979869 | 12354.8912 | 10084.9346 |

| | | | | | |
|---|---|---|---|---|---|
| *RTP1* | -0.30648993 | 0.19023551 | 0.99979869 | 6580.82086 | 5321.29452 |
| *ARO1* | -0.281126015 | 0.19033428 | 0.99979869 | 44138.1293 | 36323.3963 |
| *PRO1* | -0.286028363 | 0.19038403 | 0.99979869 | 21104.2808 | 17308.8149 |
| *RNH202* | -0.30920631 | 0.19107556 | 0.99979869 | 5824.79712 | 4701.10898 |
| *AAT2* | -0.277924265 | 0.191343 | 0.99979869 | 82949.9495 | 68415.1795 |
| *UTP18* | -0.287417331 | 0.19280787 | 0.99979869 | 14854.1594 | 12171.0133 |
| *NOP16* | -0.282827224 | 0.19299158 | 0.99979869 | 22262.2165 | 18299.0616 |
| *YLR363W-A* | -0.310440432 | 0.19364095 | 0.99979869 | 5133.15733 | 4139.35413 |
| *ARX1* | -0.280258166 | 0.19402274 | 0.99979869 | 26751.5343 | 22028.3757 |
| *NOP7* | -0.274277587 | 0.19485788 | 0.99979869 | 86167.1963 | 71248.5561 |
| *LEU9* | -0.277276859 | 0.19562834 | 0.99979869 | 32594.8662 | 26895.5508 |
| *IPI1* | -0.302447298 | 0.19569482 | 0.99979869 | 6104.76978 | 4950.20829 |
| *SSF2* | -0.292511948 | 0.19607832 | 0.99979869 | 8884.01061 | 7253.60818 |
| *NOC4* | -0.285059534 | 0.19623816 | 0.99979869 | 13519.6555 | 11095.6833 |
| *snR54* | -0.487580349 | 0.19632444 | 0.99979869 | 701.39493 | 500.24884 |
| *UTP23* | -0.282928126 | 0.19804188 | 0.99979869 | 13716.7095 | 11274.0507 |
| *RPF1* | -0.283554723 | 0.19857238 | 0.99979869 | 12644.619 | 10388.3642 |
| *NAT2* | -0.280536256 | 0.20011641 | 0.99979869 | 13979.1229 | 11508.7986 |
| *OTU2* | -0.29248439 | 0.20146961 | 0.99979869 | 6964.19806 | 5686.23015 |
| *ARO3* | -0.269570454 | 0.20219414 | 0.99979869 | 39940.4904 | 33133.2848 |
| *snR17b* | -0.586363454 | 0.20229531 | 0.99979869 | 415.56918 | 276.777022 |
| *OPI3* | -0.279658245 | 0.20239592 | 0.99979869 | 12632.9129 | 10406.816 |
| *RPS30B* | -0.264965955 | 0.20456399 | 0.99979869 | 76925.1719 | 64018.5252 |
| *PRS4* | -0.274828868 | 0.20666017 | 0.99979869 | 13158.7152 | 10876.3119 |
| *NOP13* | -0.266259229 | 0.20668321 | 0.99979869 | 33011.4109 | 27448.0798 |
| *LEU1* | -2.628672633 | 0.2067837 | 0.99979869 | 171751.035 | 27769.9612 |
| *PEX21* | -0.305045806 | 0.20709091 | 0.99979869 | 3995.70742 | 3234.19076 |
| *DDP1* | -0.274715124 | 0.20810666 | 0.99979869 | 12037.849 | 9950.64649 |
| *CBF1* | -0.271243514 | 0.20938493 | 0.99979869 | 14110.8173 | 11692.2915 |
| *RPS24A* | -0.258386944 | 0.20989079 | 0.99979869 | 218965.937 | 183060.322 |
| *RPS23A* | -0.258059577 | 0.2102504 | 0.99979869 | 218810.83 | 182972.164 |
| *GDE1* | -0.276116567 | 0.21038429 | 0.99979869 | 9703.44279 | 8013.20734 |
| *DBP7* | -0.276574575 | 0.21092544 | 0.99979869 | 9182.51805 | 7580.6151 |
| *NIP7* | -0.267954775 | 0.21140988 | 0.99979869 | 15767.241 | 13094.6284 |
| *RPP2B* | -0.255626388 | 0.21266162 | 0.99979869 | 309856.575 | 259543.04 |
| *HCA4* | -0.2615771 | 0.21320443 | 0.99979869 | 26477.4147 | 22086.8064 |
| *GRE2* | -0.262340147 | 0.21387078 | 0.99979869 | 21952.0029 | 18302.1369 |
| *snR41* | -0.382263487 | 0.21407697 | 0.99979869 | 1214.51556 | 931.815974 |
| *NME1* | -0.390397558 | 0.214215 | 0.99979869 | 1128.67028 | 861.084069 |
| *RKI1* | -0.262917284 | 0.21433923 | 0.99979869 | 19589.3068 | 16325.7439 |
| *EFM3* | -0.289662679 | 0.21497818 | 0.99979869 | 4814.16409 | 3938.43451 |
| *RPS7A* | -0.252832055 | 0.21594899 | 0.99979869 | 239304.439 | 200835.558 |
| *SNL1* | -0.268835477 | 0.21609328 | 0.99979869 | 10755.0474 | 8926.57151 |
| *snR83* | -0.296457745 | 0.21616643 | 0.99979869 | 3904.00905 | 3178.83535 |
| *ASC1* | -0.249940036 | 0.2187841 | 0.99979869 | 544438.548 | 457835.324 |

| | | | | | |
|---|---|---|---|---|---|
| *SPO24* | -0.251944341 | 0.21993776 | 0.99979869 | 55099.9858 | 46270.9675 |
| *AKR1* | -0.252784877 | 0.22071692 | 0.99979869 | 37770.9462 | 31700.1949 |
| *PHO89* | -2.420646933 | 0.22083825 | 0.99979869 | 3951.80926 | 738.072059 |
| *POC4* | -0.295527572 | 0.22173895 | 0.99979869 | 3500.14605 | 2851.82843 |
| *FRD1* | -0.254776125 | 0.22175519 | 0.99979869 | 23503.071 | 19698.3232 |
| *ACO1* | -0.248993711 | 0.222015 | 0.99979869 | 85382.8827 | 71848.2396 |
| *DML1* | -0.282753588 | 0.22271486 | 0.99979869 | 4717.58815 | 3877.95361 |
| *LYS12* | -0.250358378 | 0.22273916 | 0.99979869 | 44151.7865 | 37117.8489 |
| *ICT1* | -0.260872222 | 0.22337013 | 0.99979869 | 11732.513 | 9791.75598 |
| *DUR1%2C2* | -0.25649051 | 0.22464151 | 0.99979869 | 15047.3113 | 12596.4298 |
| *YIL002W-A* | -0.268347331 | 0.22495036 | 0.99979869 | 7278.31373 | 6042.96498 |
| *snR36* | -0.446652691 | 0.22570271 | 0.99979869 | 669.202951 | 491.022939 |
| *snR79* | -0.773493977 | 0.22581639 | 0.99979869 | 189.249814 | 110.710809 |
| *tL(UAG)L1* | -0.678748969 | 0.22610145 | 0.99979869 | 247.780685 | 154.790112 |
| *DUS1* | -0.278755869 | 0.22612075 | 0.99979869 | 4823.91923 | 3976.36322 |
| *DIN7* | -0.318416576 | 0.22799883 | 0.99979869 | 2075.89487 | 1664.76253 |
| *MRT4* | -0.246845957 | 0.22940803 | 0.99979869 | 29083.9895 | 24510.143 |
| *LTV1* | -0.248145413 | 0.22953632 | 0.99979869 | 23564.5284 | 19840.8121 |
| *ADH2* | -0.292014514 | 0.23110037 | 0.99979869 | 3096.28304 | 2528.9219 |
| *snR11* | -0.301433084 | 0.23121315 | 0.99979869 | 2582.1869 | 2095.30457 |
| *RPL43B* | -0.240481686 | 0.23147914 | 0.99979869 | 128564.032 | 108824.625 |
| *RRP9* | -0.251491068 | 0.23166277 | 0.99979869 | 13849.3794 | 11633.8608 |
| *PHO12* | -2.254943076 | 0.23179591 | 0.99979869 | 50163.8824 | 10509.326 |
| *YML045W* | -0.534493618 | 0.23183549 | 0.99979869 | 403.863006 | 278.827222 |
| *YGR146C-A* | -0.542726282 | 0.23228709 | 0.99979869 | 388.254773 | 266.526021 |
| *NOP12* | -0.245654386 | 0.23265267 | 0.99979869 | 23324.5518 | 19672.6957 |
| *LYS2* | -0.243860251 | 0.23312084 | 0.99979869 | 28833.2823 | 24349.2022 |
| *TRP4* | -0.251600991 | 0.23425107 | 0.99979869 | 11706.1741 | 9832.75998 |
| *SGD1* | -0.249957996 | 0.23477537 | 0.99979869 | 12785.0931 | 10751.2497 |
| *SPL2* | -2.208866882 | 0.23477866 | 0.99979869 | 11331.5765 | 2451.0143 |
| *HIS3* | -0.247370366 | 0.23478671 | 0.99979869 | 15902.8375 | 13397.033 |
| *BIO5* | -0.316224583 | 0.23504276 | 0.99979869 | 1924.69012 | 1545.85092 |
| *snR45* | -0.463764819 | 0.23509102 | 0.99979869 | 557.018783 | 403.889432 |
| *YLR152C* | -0.252441174 | 0.23614274 | 0.99979869 | 9996.09714 | 8391.46927 |
| *MOD5* | -0.259788246 | 0.23617258 | 0.99979869 | 6879.3283 | 5745.68596 |
| *PUF6* | -0.241149637 | 0.23685062 | 0.99979869 | 27720.2202 | 23453.2648 |
| *OYE2* | -0.235517909 | 0.23724233 | 0.99979869 | 195524.323 | 166074.414 |
| *SVF1* | -0.238137091 | 0.23752641 | 0.99979869 | 46318.4043 | 39270.559 |
| *RCN1* | -0.266665182 | 0.237867 | 0.99979869 | 5046.33654 | 4194.70953 |
| *ALG3* | -0.244823863 | 0.23788108 | 0.99979869 | 15937.956 | 13450.3382 |
| *YPL162C* | -0.265141863 | 0.23791128 | 0.99979869 | 5300.94583 | 4411.00565 |
| *EFM4* | -0.248775469 | 0.23798921 | 0.99979869 | 11484.7323 | 9665.66867 |
| *SUF7* | -0.763837905 | 0.23848231 | 0.99979869 | 177.54364 | 104.560208 |
| *RRP15* | -0.244251719 | 0.23873324 | 0.99979869 | 15765.2899 | 13309.8995 |
| *YOL103W-B* | -0.314051147 | 0.23890396 | 0.99979869 | 1875.9144 | 1508.94732 |

| | | | | | |
|---|---|---|---|---|---|
| *SFM1* | -0.240199867 | 0.23899314 | 0.99979869 | 24683.4435 | 20897.6903 |
| *PGA3* | -0.23778733 | 0.23922162 | 0.99979869 | 36583.745 | 31024.654 |
| *snR43* | -0.354772296 | 0.23923266 | 0.99979869 | 1170.61741 | 915.414373 |
| *PHO11* | -2.137787348 | 0.2393391 | 0.99979869 | 11490.5854 | 2610.92991 |
| *SHQ1* | -0.265082455 | 0.23941295 | 0.99979869 | 5081.45506 | 4228.53784 |
| *RPL28* | -0.232525973 | 0.24055361 | 0.99979869 | 387032.454 | 329420.012 |
| *AFT2* | -0.257334288 | 0.24060681 | 0.99979869 | 6508.63278 | 5445.33163 |
| *snR81* | -0.404031433 | 0.24092589 | 0.99979869 | 767.729916 | 580.206646 |
| *HIT1* | -0.299739494 | 0.24114134 | 0.99979869 | 2248.56094 | 1826.72835 |
| *RKM3* | -0.253706638 | 0.24125839 | 0.99979869 | 7452.93083 | 6251.0603 |
| *RPA34* | -0.239127072 | 0.2414745 | 0.99979869 | 21859.3291 | 18520.4832 |
| *PHO84* | -2.102264653 | 0.24160698 | 0.99979869 | 158659.63 | 36949.7324 |
| *ENO1* | -0.231023958 | 0.24241927 | 0.99979869 | 479846.806 | 408843.741 |
| *PPH3* | -0.270920032 | 0.24288784 | 0.99979869 | 3919.61729 | 3248.54216 |
| *MNT4* | -0.257628591 | 0.24373049 | 0.99979869 | 5805.28683 | 4855.89909 |
| *TRM44* | -0.247888396 | 0.24395926 | 0.99979869 | 8964.0028 | 7548.837 |
| *ERG24* | -0.236945776 | 0.24430543 | 0.99979869 | 21815.4309 | 18511.2573 |
| *UTP21* | -0.234906642 | 0.24501099 | 0.99979869 | 27175.8831 | 23092.4295 |
| *GFD2* | -0.252349985 | 0.2451605 | 0.99979869 | 6854.94043 | 5754.91186 |
| *snR80* | -0.615052584 | 0.24562791 | 0.99979869 | 265.339946 | 173.241914 |
| *YJR011C* | -0.27446208 | 0.245711 | 0.99979869 | 3376.25571 | 2791.34752 |
| *RPC10* | -0.242538496 | 0.24599222 | 0.99979869 | 11339.3806 | 9584.68576 |
| *PEX15* | -0.273274084 | 0.24613243 | 0.99979869 | 3441.61518 | 2847.72803 |
| *RPS6A* | -0.228326937 | 0.2464517 | 0.99979869 | 254109.823 | 216914.253 |
| *YML108W* | -0.263149758 | 0.24652538 | 0.99979869 | 4462.00335 | 3718.038 |
| *ARG80* | -0.255005559 | 0.24683971 | 0.99979869 | 5813.09094 | 4871.27559 |
| *HMT1* | -0.234036067 | 0.24717026 | 0.99979869 | 24097.1593 | 20488.6753 |
| *NOP9* | -0.237604595 | 0.2477336 | 0.99979869 | 15168.2751 | 12865.006 |
| *GND1* | -0.2276622 | 0.24776108 | 0.99979869 | 174922.433 | 149386.81 |
| *RAD34* | -0.266907313 | 0.24820143 | 0.99979869 | 3858.15987 | 3206.51306 |
| *PHO5* | -1.996302098 | 0.24821413 | 0.99979869 | 31910.055 | 7997.83084 |
| *YBR072C-A* | -0.839200848 | 0.24851021 | 0.99979869 | 137.547545 | 76.8825061 |
| *YOR008C-A* | -0.472543316 | 0.24857537 | 0.99979869 | 475.075564 | 342.383427 |
| *PLC1* | -0.250082179 | 0.24868642 | 0.99979869 | 6678.37231 | 5615.49825 |
| *GPD2* | -0.229036855 | 0.24888884 | 0.99979869 | 52726.5591 | 44986.5171 |
| *MER1* | -0.36002253 | 0.2494395 | 0.99979869 | 999.902369 | 779.076062 |
| *LRP1* | -0.237996472 | 0.24996667 | 0.99979869 | 12655.3497 | 10730.7477 |
| *CGR1* | -0.239564134 | 0.25016864 | 0.99979869 | 11129.645 | 9426.82035 |
| *ALD5* | -1.964265294 | 0.25018198 | 0.99979869 | 68930.8305 | 17664.5246 |
| *TFB5* | -0.268295427 | 0.25024799 | 0.99979869 | 3560.62795 | 2956.38864 |
| *DIA1* | -0.228842175 | 0.25182167 | 0.99979869 | 32417.3226 | 27662.3257 |
| *YGR038C-B* | -0.460252137 | 0.25193447 | 0.99979869 | 493.61034 | 358.785029 |
| *TIR4* | -0.273381531 | 0.25199528 | 0.99979869 | 3055.31143 | 2527.8968 |
| *YPR078C* | -0.502862803 | 0.25207605 | 0.99979869 | 398.009918 | 280.877422 |
| *GLN1* | -0.223861413 | 0.25323224 | 0.99979869 | 149539.545 | 128046.276 |

| | | | | | |
|---|---|---|---|---|---|
| *ARF1* | -0.224391575 | 0.25334839 | 0.99979869 | 96063.791 | 82226.3529 |
| *PIN2* | -0.256934026 | 0.2533848 | 0.99979869 | 4532.2404 | 3792.8703 |
| *FOL2* | -0.229515208 | 0.25353318 | 0.99979869 | 23417.2257 | 19973.05 |
| *CIN2* | -0.272990278 | 0.25402064 | 0.99979869 | 2966.53961 | 2455.1147 |
| *MET6* | -0.225386519 | 0.25411358 | 0.99979869 | 50440.9286 | 43145.4373 |
| *YHB1* | -0.223511361 | 0.2544913 | 0.99979869 | 99956.0939 | 85610.2082 |
| *TOM6* | -0.232537598 | 0.25459748 | 0.99979869 | 15035.6051 | 12797.3494 |
| *HOM6* | -0.22326356 | 0.25572792 | 0.99979869 | 72214.4123 | 61860.6895 |
| *ANB1* | -0.255473688 | 0.25596989 | 0.99979869 | 4444.44409 | 3723.1635 |
| *SSU1* | -0.235077461 | 0.25639891 | 0.99979869 | 10964.783 | 9316.10954 |
| *GPM3* | -0.237937227 | 0.25662336 | 0.99979869 | 9024.4847 | 7652.37211 |
| *RSA4* | -0.237390578 | 0.25702882 | 0.99979869 | 9155.20364 | 7766.15822 |
| *MRS4* | -0.242090967 | 0.25719203 | 0.99979869 | 7094.917 | 5998.88568 |
| *snR191* | -0.449915741 | 0.25778429 | 0.99979869 | 498.487913 | 364.935629 |
| *CIA1* | -0.235813637 | 0.25783533 | 0.99979869 | 9721.00205 | 8255.13096 |
| *snR31* | -0.270182253 | 0.25818545 | 0.99979869 | 2918.7394 | 2420.26129 |
| *TCB2* | -0.222591746 | 0.25884086 | 0.99979869 | 43033.8469 | 36881.0507 |
| *BAT1* | -1.810682643 | 0.2593446 | 0.99979869 | 125928.192 | 35895.9296 |
| *TOM7* | -0.236974264 | 0.25951844 | 0.99979869 | 8426.4943 | 7150.07307 |
| *VBA2* | -0.24945571 | 0.26011336 | 0.99979869 | 4851.23364 | 4080.92343 |
| *PHM6* | -1.790709128 | 0.2603808 | 0.99979869 | 2677.78732 | 773.950562 |
| *DRN1* | -0.23811329 | 0.26212043 | 0.99979869 | 7186.61537 | 6093.19489 |
| *IMT1* | -1.928458219 | 0.26230694 | 0.99979869 | 27.3144062 | 7.17570057 |
| *DIC1* | -1.754189183 | 0.26259254 | 0.99979869 | 37437.3202 | 11097.7335 |
| *YLR407W* | -0.246649792 | 0.26283399 | 0.99979869 | 4968.29538 | 4187.53383 |
| *MAM33* | -0.232562774 | 0.26288206 | 0.99979869 | 9449.80902 | 8042.93524 |
| *VTC3* | -1.747565328 | 0.26297529 | 0.99979869 | 82583.156 | 24593.1761 |
| *DEG1* | -0.235177788 | 0.26323739 | 0.99979869 | 8002.14549 | 6798.46374 |
| *TIM8* | -0.237992023 | 0.26324437 | 0.99979869 | 6946.6388 | 5890.22507 |
| *SUA5* | -0.230891773 | 0.26336407 | 0.99979869 | 10301.4332 | 8777.932 |
| *FKH1* | -0.232213107 | 0.26370753 | 0.99979869 | 9305.43287 | 7921.97343 |
| *snR39B* | -0.424757569 | 0.26454044 | 0.99979869 | 546.288123 | 406.964732 |
| *snR51* | -0.359388453 | 0.26526087 | 0.99979869 | 861.379309 | 671.440554 |
| *ARO2* | -0.215261828 | 0.26644316 | 0.99979869 | 101622.273 | 87536.3713 |
| *QRI5* | -0.233508919 | 0.26673325 | 0.99979869 | 7642.18064 | 6500.15962 |
| *CHO1* | -0.221180779 | 0.26680403 | 0.99979869 | 20277.0445 | 17394.9233 |
| *ALR1* | -0.220117238 | 0.26702877 | 0.99979869 | 23029.9465 | 19771.1053 |
| *ERV25* | -0.216271773 | 0.26709229 | 0.99979869 | 52613.3994 | 45288.9216 |
| *MTO1* | -0.242290715 | 0.26724891 | 0.99979869 | 5149.74108 | 4353.60005 |
| *TIF2* | -0.214281374 | 0.26751402 | 0.99979869 | 122813.374 | 105862.085 |
| *RPS3* | -0.213286714 | 0.26761415 | 0.99979869 | 440831.103 | 380247.549 |
| *KTR2* | -0.241776611 | 0.26781178 | 0.99979869 | 5167.30034 | 4370.00165 |
| *snR66* | -0.369172275 | 0.26824613 | 0.99979869 | 774.558518 | 599.683548 |
| *MET8* | -0.258313811 | 0.26825174 | 0.99979869 | 3164.56906 | 2645.78331 |
| *GID8* | -0.221986549 | 0.26909248 | 0.99979869 | 15425.8109 | 13225.8413 |

| | | | | | |
|---|---|---|---|---|---|
| *YLR179C* | -0.214181607 | 0.27070504 | 0.99979869 | 47650.9571 | 41076.7854 |
| *snR70* | -0.320334391 | 0.27190068 | 0.99979869 | 1164.76432 | 932.841074 |
| *CIS1* | -1.579094386 | 0.27204068 | 0.99979869 | 2235.87925 | 748.32306 |
| *RPC40* | -0.213321164 | 0.27211145 | 0.99979869 | 46746.6551 | 40321.2866 |
| *SOL3* | -0.214735151 | 0.27258919 | 0.99979869 | 30645.7882 | 26407.6032 |
| *TIM9* | -0.224898927 | 0.27323604 | 0.99979869 | 9669.29978 | 8273.58276 |
| *TRM11* | -0.219109627 | 0.27343445 | 0.99979869 | 15328.2594 | 13168.4357 |
| *RIX7* | -0.217539631 | 0.27344351 | 0.99979869 | 18240.1702 | 15687.1066 |
| *NOC3* | -0.218109247 | 0.27386893 | 0.99979869 | 16529.1178 | 14209.9373 |
| *ERG11* | -0.210098192 | 0.27389027 | 0.99979869 | 124919.51 | 107990.193 |
| *MNN1* | -0.216520975 | 0.2739623 | 0.99979869 | 19861.4753 | 17093.5439 |
| *TAH18* | -0.229696444 | 0.27418585 | 0.99979869 | 7087.11289 | 6043.99008 |
| *snR56* | -0.409809639 | 0.27517645 | 0.99979869 | 550.190181 | 414.140433 |
| *TOD6* | -0.226497033 | 0.27532233 | 0.99979869 | 8044.09262 | 6875.34625 |
| *YEL1* | -0.240305878 | 0.27539387 | 0.99979869 | 4481.51364 | 3793.8954 |
| *COG7* | -0.220400865 | 0.2756 | 0.99979869 | 11944.1996 | 10252.0259 |
| *SAS2* | -0.254614584 | 0.27572152 | 0.99979869 | 3008.48674 | 2521.7462 |
| *PSY4* | -0.223414776 | 0.27578293 | 0.99979869 | 9507.36438 | 8143.39505 |
| *tK(UUU)K* | -0.646230306 | 0.27615503 | 0.99979869 | 194.127387 | 124.03711 |
| *LAC1* | -0.214203157 | 0.27744569 | 0.99979869 | 19947.3206 | 17195.0288 |
| *RRP36* | -0.233665111 | 0.27768964 | 0.99979869 | 5351.67258 | 4551.44436 |
| *YGR161W-C* | -0.241437652 | 0.27845332 | 0.99979869 | 4031.80145 | 3410.50797 |
| *RPS21A* | -0.207171896 | 0.27863764 | 0.99979869 | 116328.154 | 100767.338 |
| *PMP2* | -0.208122059 | 0.27864766 | 0.99979869 | 67224.6556 | 58193.9065 |
| *SYO1* | -0.21606444 | 0.27935811 | 0.99979869 | 14055.213 | 12100.2814 |
| *PEP3* | -0.21832468 | 0.27987215 | 0.99979869 | 11238.9026 | 9660.54317 |
| *YDR261C-D* | -0.240402025 | 0.28025335 | 0.99979869 | 3998.63396 | 3384.88047 |
| *MPP10* | -0.209410483 | 0.28029678 | 0.99979869 | 32979.2189 | 28523.4098 |
| *LST7* | -0.232154243 | 0.28067246 | 0.99979869 | 5231.6843 | 4454.05986 |
| *snR9* | -0.279066585 | 0.28074641 | 0.99979869 | 1772.50986 | 1460.76762 |
| *RIB3* | -0.209311226 | 0.2810146 | 0.99979869 | 30585.3063 | 26454.7578 |
| *ATX2* | -0.231740714 | 0.28170074 | 0.99979869 | 5171.2024 | 4403.82995 |
| *YML133C* | -0.316286032 | 0.28233438 | 0.99979869 | 1092.57625 | 877.48567 |
| *GDH1* | -1.377721547 | 0.28243216 | 0.99979869 | 100477.019 | 38665.75 |
| *YGL039W* | -0.214335371 | 0.28245589 | 0.99979869 | 13681.5909 | 11792.7513 |
| *IZH2* | -0.214971435 | 0.28247744 | 0.99979869 | 12890.4487 | 11105.9343 |
| *ELF1* | -0.207926444 | 0.28450352 | 0.99979869 | 26218.9034 | 22699.8162 |
| *snR74* | -1.002467496 | 0.28457251 | 0.99979869 | 78.0411605 | 38.9538031 |
| *SHE9* | -0.226962866 | 0.28461821 | 0.99979869 | 5777.97242 | 4936.88199 |
| *HAS1* | -0.206058435 | 0.28474594 | 0.99979869 | 37709.4887 | 32690.4416 |
| *SEN15* | -0.272164418 | 0.2848162 | 0.99979869 | 1872.98785 | 1550.97642 |
| *RRN3* | -0.21963602 | 0.28499557 | 0.99979869 | 8204.077 | 7045.51286 |
| *PXP2* | -0.240698926 | 0.28516086 | 0.99979869 | 3579.16272 | 3029.17074 |
| *SPS4* | -0.411108754 | 0.28533695 | 0.99979869 | 504.341 | 379.28703 |
| *TTI1* | -0.224259283 | 0.28553372 | 0.99979869 | 6335.96672 | 5423.80453 |

| | | | | | |
|---|---|---|---|---|---|
| *RSM27* | -0.224219339 | 0.28564297 | 0.99979869 | 6326.21157 | 5415.60373 |
| *FIT2* | -1.298273271 | 0.28611766 | 0.99979869 | 17554.3835 | 7137.77187 |
| *YMR102C* | -0.205242264 | 0.2864159 | 0.99979869 | 35759.4353 | 31017.4783 |
| *DBP3* | -0.204649163 | 0.28645282 | 0.99979869 | 41495.4605 | 36007.6655 |
| *FIT3* | -1.276879907 | 0.28711704 | 0.99979869 | 51008.678 | 21050.4302 |
| *NOG2* | -0.206572035 | 0.28775314 | 0.99979869 | 23435.7605 | 20309.2828 |
| *DBP8* | -0.216428843 | 0.28791143 | 0.99979869 | 8884.98612 | 7647.24661 |
| *TIS11* | -1.25368386 | 0.28800035 | 0.99979869 | 4702.95543 | 1972.29256 |
| *REX4* | -0.221215011 | 0.28847717 | 0.99979869 | 6671.54371 | 5723.13376 |
| *YIL102C-A* | -0.265655458 | 0.28909232 | 0.99979869 | 1970.5393 | 1639.13503 |
| *TDH1* | -0.200305363 | 0.2892301 | 0.99979869 | 163770.351 | 142540.166 |
| *ATC1* | -0.222574588 | 0.29008409 | 0.99979869 | 5961.36915 | 5109.09881 |
| *ARG1* | -1.208768851 | 0.29008958 | 0.99979869 | 17981.6589 | 7779.48452 |
| *SES1* | -0.199947553 | 0.29043383 | 0.99979869 | 117058.814 | 101909.3 |
| *ENP1* | -0.20652863 | 0.29057535 | 0.99979869 | 18380.6443 | 15929.0302 |
| *TCD2* | -0.211927575 | 0.29067551 | 0.99979869 | 10756.9985 | 9287.40674 |
| *VTC4* | -1.180652651 | 0.29132726 | 0.99979869 | 78792.3067 | 34759.0936 |
| *TSR2* | -0.208938227 | 0.2913314 | 0.99979869 | 13400.6428 | 11593.8819 |
| *YRB1* | -0.199521584 | 0.2917559 | 0.99979869 | 91010.6258 | 79255.6128 |
| *TMA19* | -0.197735377 | 0.2926652 | 0.99979869 | 405319.449 | 353405.303 |
| *MOG1* | -0.219038773 | 0.29271799 | 0.99979869 | 6481.31838 | 5568.34364 |
| *snR59* | -0.609400563 | 0.29272523 | 0.99979869 | 197.05393 | 129.16261 |
| *DFG5* | -0.204256332 | 0.29297741 | 0.99979869 | 20508.2415 | 17800.8629 |
| *RPC17* | -0.222176863 | 0.29317105 | 0.99979869 | 5555.55511 | 4762.61498 |
| *VTC1* | -1.113495074 | 0.29409656 | 0.99979869 | 76308.6467 | 35267.5432 |
| *ARN2* | -1.093444935 | 0.29485473 | 0.99979869 | 19120.0843 | 8960.39981 |
| *PHO4* | -0.204297014 | 0.29501249 | 0.99979869 | 17327.0887 | 15039.2433 |
| *YJL028W* | -1.25039295 | 0.29510695 | 0.99979869 | 48.7757253 | 20.5020016 |
| *YMR084W* | -0.269507008 | 0.29554221 | 0.99979869 | 1705.19936 | 1414.63811 |
| *HMF1* | -0.202683185 | 0.29600691 | 0.99979869 | 19792.2138 | 17198.1041 |
| *MAS1* | -0.205438036 | 0.29656751 | 0.99979869 | 13783.0445 | 11953.6921 |
| *MTC2* | -0.210166534 | 0.29686154 | 0.99979869 | 9244.95097 | 7991.68024 |
| *EHT1* | -0.202021142 | 0.29716294 | 0.99979869 | 19717.0992 | 17140.6985 |
| *snR87* | -0.344144502 | 0.29726782 | 0.99979869 | 736.513452 | 580.206646 |
| *RIB5* | -0.202219764 | 0.29748558 | 0.99979869 | 18693.7845 | 16248.8614 |
| *TRP2* | -0.198262401 | 0.2981369 | 0.99979869 | 35403.3725 | 30857.5627 |
| *SRO77* | -0.216363863 | 0.29833104 | 0.99979869 | 6201.34572 | 5337.69613 |
| *TLC1* | -0.197554536 | 0.29868778 | 0.99979869 | 39163.0054 | 34151.2092 |
| *MRD1* | -0.199474264 | 0.29901046 | 0.99979869 | 24755.6316 | 21558.8798 |
| *LDB7* | -0.199430078 | 0.29922959 | 0.99979869 | 24394.6913 | 21245.1992 |
| *STR2* | -0.213984197 | 0.29964527 | 0.99979869 | 6688.12745 | 5766.18796 |
| *HTZ1* | -0.204168219 | 0.29993664 | 0.99979869 | 12814.3586 | 11123.361 |
| *YDR514C* | -0.219490132 | 0.30001231 | 0.99979869 | 5169.25137 | 4439.70845 |
| *YDR034C-A* | -0.762458274 | 0.30008745 | 0.99979869 | 119.988284 | 70.7319056 |
| *ALG14* | -0.232632134 | 0.30018459 | 0.99979869 | 3346.01476 | 2847.72803 |

| | | | | | |
|---|---|---|---|---|---|
| *POP8* | -0.215872594 | 0.30057052 | 0.99979869 | 5940.88334 | 5115.24941 |
| *GDT1* | -0.197388083 | 0.30061616 | 0.99979869 | 31016.4837 | 27050.341 |
| *ERG3* | -0.193618264 | 0.30132774 | 0.99979869 | 116606.175 | 101961.58 |
| *HIP1* | -0.195917414 | 0.30174717 | 0.99979869 | 37818.7464 | 33016.4234 |
| *TRF5* | -0.218623184 | 0.30203636 | 0.99979869 | 5081.45506 | 4366.92635 |
| *SPR6* | -0.217589507 | 0.30239986 | 0.99979869 | 5248.26804 | 4513.51566 |
| *ISA1* | -0.204865997 | 0.30286741 | 0.99979869 | 10412.6418 | 9034.20702 |
| *HHT1* | -0.193048624 | 0.30341384 | 0.99979869 | 77699.7304 | 67968.2358 |
| *RIX1* | -0.202289719 | 0.30351466 | 0.99979869 | 12521.7042 | 10883.4876 |
| *CIR1* | -0.201059977 | 0.30371078 | 0.99979869 | 13949.8574 | 12135.1348 |
| *MET5* | -0.202514908 | 0.30383438 | 0.99979869 | 12061.2614 | 10481.6483 |
| *YBL028C* | -0.201131917 | 0.30396608 | 0.99979869 | 13640.6193 | 11865.5334 |
| *YCR007C* | -0.251075377 | 0.30430626 | 0.99979869 | 2078.82141 | 1746.77054 |
| *GSC2* | -0.198544843 | 0.30491373 | 0.99979869 | 17113.451 | 14913.156 |
| *MRPL23* | -0.209735469 | 0.30492978 | 0.99979869 | 7038.33716 | 6086.01919 |
| *MAK10* | -0.204486825 | 0.30510936 | 0.99979869 | 9691.73662 | 8410.94617 |
| *FTH1* | -0.199151167 | 0.30541192 | 0.99979869 | 15390.6924 | 13406.2589 |
| *RMT2* | -0.196018772 | 0.30542891 | 0.99979869 | 23668.9085 | 20661.9172 |
| *CEP3* | -0.210090629 | 0.30548721 | 0.99979869 | 6782.75236 | 5863.57247 |
| *DIB1* | -0.224360401 | 0.30564212 | 0.99979869 | 3829.86995 | 3278.27006 |
| *OSW1* | -1.867056823 | 0.30568074 | 0.99979869 | 22.4368336 | 6.15060049 |
| *YOR192C-C* | -0.35890446 | 0.30575224 | 0.99979869 | 612.62311 | 477.696638 |
| *PEX5* | -0.205661873 | 0.30644281 | 0.99979869 | 8471.36797 | 7345.86719 |
| *YGK3* | -0.24108818 | 0.30675038 | 0.99979869 | 2448.54141 | 2071.72727 |
| *RPC37* | -0.199361712 | 0.30676323 | 0.99979869 | 13829.8692 | 12044.926 |
| *RPF2* | -0.194913482 | 0.30703563 | 0.99979869 | 24469.8059 | 21377.4371 |
| *PUS2* | -0.209284511 | 0.30724626 | 0.99979869 | 6690.07848 | 5786.68996 |
| *RPL3* | -0.188956869 | 0.30775526 | 0.99979869 | 964112.693 | 845757.797 |
| *YDL009C* | -0.27582653 | 0.30785663 | 0.99979869 | 1354.01413 | 1118.38419 |
| *ALG1* | -0.199221314 | 0.30815514 | 0.99979869 | 12947.0285 | 11277.126 |
| *NOP4* | -0.192305949 | 0.30876599 | 0.99979869 | 34431.76 | 30134.8671 |
| *YJL047C-A* | -0.315871899 | 0.3087681 | 0.99979869 | 857.477251 | 688.867255 |
| *RPP1B* | -0.189101895 | 0.30884946 | 0.99979869 | 157319.273 | 137992.822 |
| *MRP2* | -0.217384633 | 0.30915941 | 0.99979869 | 4488.34224 | 3860.52691 |
| *MRP7* | -0.197628479 | 0.30956424 | 0.99979869 | 14002.5352 | 12209.9671 |
| *PRP9* | -0.209599413 | 0.30976229 | 0.99979869 | 6104.76978 | 5279.26542 |
| *RPC31* | -0.195877732 | 0.31042574 | 0.99979869 | 16115.4996 | 14069.4986 |
| *URK1* | -0.201142262 | 0.3104939 | 0.99979869 | 9816.60247 | 8539.08368 |
| *YHR020W* | -0.188938745 | 0.31057765 | 0.99979869 | 83517.6989 | 73265.953 |
| *MRPL31* | -0.204698654 | 0.31100832 | 0.99979869 | 7608.03763 | 6601.64453 |
| *DLD2* | -0.194411819 | 0.31123981 | 0.99979869 | 18317.2359 | 16007.9629 |
| *TMA16* | -0.203084342 | 0.31142385 | 0.99979869 | 8273.33853 | 7186.97667 |
| *YPL108W* | -0.236592437 | 0.31158779 | 0.99979869 | 2510.97434 | 2131.18307 |
| *NAR1* | -0.195856055 | 0.31166532 | 0.99979869 | 14900.0086 | 13008.52 |
| *PET123* | -0.202720181 | 0.31199554 | 0.99979869 | 8288.94676 | 7202.35317 |

| | | | | | |
|---|---|---|---|---|---|
| *TMA7* | -0.190422493 | 0.3124147 | 0.99979869 | 33282.6039 | 29167.1726 |
| *RPB10* | -0.19497601 | 0.31252364 | 0.99979869 | 15586.7708 | 13616.4044 |
| *snR58* | -0.531810272 | 0.31298644 | 0.99979869 | 231.196938 | 159.915613 |
| *RPL31B* | -0.187394135 | 0.31309345 | 0.99979869 | 93407.465 | 82029.5336 |
| *YML018C* | -0.19631392 | 0.31311689 | 0.99979869 | 12838.7464 | 11205.369 |
| *TEF2* | -0.185927673 | 0.31341368 | 0.99979869 | 732391.903 | 643833.583 |
| *PCL7* | -0.201089349 | 0.31379012 | 0.99979869 | 8662.56881 | 7535.5107 |
| *CTR1* | -0.1871522 | 0.31416675 | 0.99979869 | 76003.3107 | 66756.5675 |
| *SAP185* | -0.195223762 | 0.31478831 | 0.99979869 | 13337.2343 | 11649.2373 |
| *TIM17* | -0.189588293 | 0.31507864 | 0.99979869 | 29944.3933 | 26256.9135 |
| *RPS19B* | -0.184666041 | 0.31633806 | 0.99979869 | 259228.347 | 228082.718 |
| *POA1* | -0.206503382 | 0.3166829 | 0.99979869 | 5810.1644 | 5035.2916 |
| *SUF10* | -1.397948262 | 0.31722863 | 0.99979869 | 35.1185222 | 13.3263011 |
| *PMU1* | -0.190512719 | 0.31725044 | 0.99979869 | 19857.5733 | 17401.0739 |
| *YGR283C* | -0.204621347 | 0.31778196 | 0.99979869 | 6165.25168 | 5349.99733 |
| *SER1* | -0.186157057 | 0.3179415 | 0.99979869 | 48283.0905 | 42438.1183 |
| *NOB1* | -0.189408409 | 0.3188761 | 0.99979869 | 20529.7028 | 18003.8327 |
| *UTP4* | -0.188128363 | 0.31973932 | 0.99979869 | 23482.5852 | 20611.6873 |
| *NMD3* | -0.185493219 | 0.3198998 | 0.99979869 | 42520.7263 | 37390.5255 |
| *ESF1* | -0.184615167 | 0.32015134 | 0.99979869 | 55594.5717 | 48916.7508 |
| *RPL14A* | -0.182737956 | 0.32034796 | 0.99979869 | 204608.315 | 180265.9 |
| *MDM35* | -0.200887352 | 0.32040396 | 0.99979869 | 6922.25094 | 6022.46298 |
| *SEC53* | -0.183494855 | 0.32076985 | 0.99979869 | 82633.8828 | 72764.6791 |
| *SAM4* | -0.185118504 | 0.32124701 | 0.99979869 | 38393.3244 | 33769.872 |
| *STB6* | -0.205260152 | 0.32161997 | 0.99979869 | 5386.7911 | 4672.40617 |
| *RPL18B* | -0.183310372 | 0.32165464 | 0.99979869 | 70560.9153 | 62141.567 |
| *AAH1* | -0.18640363 | 0.32180041 | 0.99979869 | 26384.7408 | 23186.7388 |
| *HEM2* | -0.186574552 | 0.32190355 | 0.99979869 | 25249.242 | 22186.2411 |
| *ADH4* | -0.193481157 | 0.32240637 | 0.99979869 | 10545.3118 | 9221.80034 |
| *PCC1* | -0.207730905 | 0.32258241 | 0.99979869 | 4744.90256 | 4108.60113 |
| *THI80* | -0.210690443 | 0.32265282 | 0.99979869 | 4238.61053 | 3662.68259 |
| *MCH5* | -0.19350786 | 0.32265439 | 0.99979869 | 10404.8377 | 9098.78833 |
| *YDR090C* | -0.200831074 | 0.32294911 | 0.99979869 | 6418.88545 | 5584.74525 |
| *ERG26* | -0.187485093 | 0.3239801 | 0.99979869 | 18181.6394 | 15965.9338 |
| *EGO2* | -0.229094494 | 0.32470577 | 0.99979869 | 2442.68832 | 2084.02847 |
| *RHO3* | -0.189028435 | 0.32524747 | 0.99979869 | 13856.208 | 12154.6117 |
| *CAF16* | -0.194824963 | 0.32550749 | 0.99979869 | 8402.10644 | 7340.74169 |
| *FBA1* | -0.17941559 | 0.32559021 | 0.99979869 | 2611696.21 | 2306285.54 |
| *tG(GCC)D2* | -0.332602618 | 0.32589813 | 0.99979869 | 642.86406 | 510.499841 |
| *BUR2* | -0.216727616 | 0.32634478 | 0.99979869 | 3253.34088 | 2799.54832 |
| *GIS4* | -0.195829859 | 0.32666692 | 0.99979869 | 7560.23742 | 6600.61943 |
| *NSL1* | -0.203767198 | 0.32677009 | 0.99979869 | 5048.28757 | 4383.32795 |
| *RPL35A* | -0.179473658 | 0.32694697 | 0.99979869 | 166863.707 | 147344.81 |
| *PPM2* | -0.22324114 | 0.32698521 | 0.99979869 | 2708.02827 | 2319.80149 |
| *AQR1* | -0.200330822 | 0.32725017 | 0.99979869 | 5821.87057 | 5067.0697 |

| | | | | | |
|---|---|---|---|---|---|
| *CUL3* | -0.199074469 | 0.32729019 | 0.99979869 | 6188.66403 | 5391.00133 |
| *FLR1* | -0.206996766 | 0.3273261 | 0.99979869 | 4383.96219 | 3797.9958 |
| *IPI3* | -0.18862277 | 0.32773793 | 0.99979869 | 12587.0637 | 11044.4283 |
| *CUP9* | -0.186944849 | 0.32847078 | 0.99979869 | 14422.0065 | 12669.2119 |
| *RKM2* | -0.205310391 | 0.32856185 | 0.99979869 | 4547.84863 | 3944.58511 |
| *MATALPHA2* | -0.451147723 | 0.32905362 | 0.99979869 | 298.507439 | 218.346317 |
| *RPL4B* | -0.177770476 | 0.33002581 | 0.99979869 | 202890.434 | 179368.937 |
| *PAP2* | -0.188369592 | 0.33002601 | 0.99979869 | 11479.8547 | 10074.6836 |
| *RAI1* | -0.19029537 | 0.33056595 | 0.99979869 | 9539.55636 | 8360.71627 |
| *COQ2* | -0.191835995 | 0.33066875 | 0.99979869 | 8494.78032 | 7437.10109 |
| *NHP6A* | -0.181365115 | 0.33114879 | 0.99979869 | 27192.4669 | 23980.1662 |
| *NPP2* | -0.192856454 | 0.3315074 | 0.99979869 | 7707.54011 | 6743.10834 |
| *PEX4* | -0.211229472 | 0.33151812 | 0.99979869 | 3491.36642 | 3015.84444 |
| *ARE1* | -0.180895842 | 0.33180749 | 0.99979869 | 27968.0009 | 24672.1088 |
| *CBF5* | -0.177001675 | 0.33251731 | 0.99979869 | 116385.709 | 102947.726 |
| *YSA1* | -0.181837616 | 0.33271285 | 0.99979869 | 21416.4455 | 18880.2933 |
| *YOR019W* | -0.208439559 | 0.33418429 | 0.99979869 | 3627.93845 | 3139.88155 |
| *GPM1* | -0.175121715 | 0.33420508 | 0.99979869 | 1505790.53 | 1333667.51 |
| *NHP2* | -0.177937605 | 0.33448556 | 0.99979869 | 42806.552 | 37839.5193 |
| *FLO9* | -0.211389887 | 0.33450483 | 0.99979869 | 3292.36146 | 2843.62763 |
| *SET5* | -0.190620333 | 0.33473363 | 0.99979869 | 7955.3208 | 6970.68056 |
| *AIM20* | -0.187003219 | 0.33481084 | 0.99979869 | 10363.8661 | 9103.91383 |
| *YHL009W-B* | -0.370591141 | 0.33505585 | 0.99979869 | 454.58976 | 351.609328 |
| *snR50* | -0.352678963 | 0.3354579 | 0.99979869 | 513.12063 | 401.839232 |
| *BSC5* | -0.219113674 | 0.33590636 | 0.99979869 | 2614.37888 | 2245.99428 |
| *YJL114W* | -0.492630498 | 0.33630546 | 0.99979869 | 235.098996 | 167.091313 |
| *TRZ1* | -0.186847533 | 0.33661259 | 0.99979869 | 9715.14897 | 8534.98328 |
| *NCS2* | -0.193314347 | 0.33664147 | 0.99979869 | 6380.84038 | 5580.64485 |
| *PEX12* | -0.192705922 | 0.33680897 | 0.99979869 | 6563.2616 | 5742.61066 |
| *RPC82* | -0.182809287 | 0.33697088 | 0.99979869 | 13942.0533 | 12282.7492 |
| *ARP9* | -0.182121861 | 0.33770796 | 0.99979869 | 14442.4923 | 12729.6928 |
| *MIC12* | -0.194699721 | 0.33852085 | 0.99979869 | 5644.32693 | 4931.75649 |
| *snR73* | -0.531398189 | 0.33889459 | 0.99979869 | 197.05393 | 136.338311 |
| *SAS10* | -0.176081258 | 0.33921396 | 0.99979869 | 36686.174 | 32471.0702 |
| *RPI1* | -0.190334948 | 0.33922847 | 0.99979869 | 6975.90423 | 6113.69689 |
| *snR19* | -0.288257634 | 0.33965134 | 0.99979869 | 866.256881 | 709.369257 |
| *SIW14* | -0.186247518 | 0.33971898 | 0.99979869 | 8978.63551 | 7891.22043 |
| *ADE12* | -0.174595719 | 0.33987405 | 0.99979869 | 55075.598 | 48797.8392 |
| *TUB4* | -0.190010362 | 0.33988246 | 0.99979869 | 6966.14909 | 6106.52119 |
| *DOG1* | -0.233369993 | 0.34005899 | 0.99979869 | 1828.11418 | 1555.07682 |
| *HNM1* | -0.177919848 | 0.34009627 | 0.99979869 | 22086.6239 | 19524.0562 |
| *FRE4* | -0.208671863 | 0.34046378 | 0.99979869 | 3216.27133 | 2783.14672 |
| *snR48* | -0.413065148 | 0.34051432 | 0.99979869 | 338.503534 | 254.22482 |
| *RPL8A* | -0.1726129 | 0.34067951 | 0.99979869 | 173983.012 | 154363.671 |
| *RPS6B* | -0.172216391 | 0.34124865 | 0.99979869 | 208129.922 | 184710.734 |

| | | | | | |
|---|---|---|---|---|---|
| *IRS4* | -0.191141824 | 0.34163948 | 0.99979869 | 6206.22329 | 5436.10573 |
| *PAU6* | -0.340026775 | 0.3417248 | 0.99979869 | 538.484007 | 425.416534 |
| *FOL1* | -0.180170866 | 0.34198799 | 0.99979869 | 14075.6988 | 12423.1879 |
| *NFU1* | -0.182088895 | 0.34225791 | 0.99979869 | 11382.3033 | 10032.6545 |
| *YFR035C* | -0.202164947 | 0.34245668 | 0.99979869 | 3816.21275 | 3317.22386 |
| *UTP25* | -0.180400524 | 0.34270455 | 0.99979869 | 13181.152 | 11631.8106 |
| *CNS1* | -0.180486457 | 0.34271144 | 0.99979869 | 13055.3106 | 11520.0747 |
| *SSK22* | -0.184380028 | 0.34274614 | 0.99979869 | 9182.51805 | 8080.86394 |
| *EMG1* | -0.1770087 | 0.34310528 | 0.99979869 | 20015.6066 | 17704.5035 |
| *SEN34* | -0.206911892 | 0.34387107 | 0.99979869 | 3195.78552 | 2768.79532 |
| *SKP2* | -0.194057358 | 0.34390932 | 0.99979869 | 5076.57749 | 4437.65825 |
| *NTR2* | -0.189222001 | 0.34414509 | 0.99979869 | 6407.17928 | 5619.59865 |
| *RLP24* | -0.176062028 | 0.34416487 | 0.99979869 | 21483.756 | 19015.6065 |
| *GPN2* | -0.184560179 | 0.34458001 | 0.99979869 | 8450.88217 | 7436.07599 |
| *TEF1* | -0.170080525 | 0.34487057 | 0.99979869 | 594718.517 | 528581.581 |
| *FET5* | -0.177436916 | 0.34496004 | 0.99979869 | 16426.6888 | 14525.6682 |
| *SPP381* | -0.209334994 | 0.34507808 | 0.99979869 | 2922.64146 | 2527.8968 |
| *FMP23* | -0.239318757 | 0.34538119 | 0.99979869 | 1550.09255 | 1313.1532 |
| *RCM1* | -0.181719803 | 0.34578622 | 0.99979869 | 10034.1422 | 8846.61371 |
| *DTR1* | -0.268320041 | 0.34581669 | 0.99979869 | 1027.21677 | 852.883268 |
| *RRP12* | -0.172699122 | 0.34587857 | 0.99979869 | 38097.7435 | 33799.5999 |
| *PAM16* | -0.178337113 | 0.34711078 | 0.99979869 | 12980.196 | 11470.8699 |
| *INA22* | -0.193952428 | 0.34722901 | 0.99979869 | 4718.56367 | 4125.00273 |
| *YNL019C* | -0.234433146 | 0.34753422 | 0.99979869 | 1643.74194 | 1397.21141 |
| *DYS1* | -0.17051401 | 0.34785456 | 0.99979869 | 61169.6371 | 54350.8063 |
| *TRM732* | -0.177739269 | 0.34840006 | 0.99979869 | 12922.6407 | 11424.7404 |
| *ELP3* | -0.174417411 | 0.34915577 | 0.99979869 | 18864.4995 | 16716.307 |
| *NOP2* | -0.170856969 | 0.34923514 | 0.99979869 | 41037.9442 | 36454.6091 |
| *CSL4* | -0.177276436 | 0.34964194 | 0.99979869 | 12716.8071 | 11246.373 |
| *ARF3* | -0.188843317 | 0.34977066 | 0.99979869 | 5600.42878 | 4913.30469 |
| *RPL2B* | -0.167434204 | 0.35056309 | 0.99979869 | 512647.506 | 456473.991 |
| *ENO2* | -0.167186087 | 0.35064742 | 0.99979869 | 3482928.22 | 3101818.56 |
| *URA7* | -0.169230889 | 0.35106626 | 0.99979869 | 54811.2336 | 48744.534 |
| *FRS2* | -0.168664643 | 0.3510809 | 0.99979869 | 72784.1128 | 64753.522 |
| *NPL6* | -0.173066821 | 0.35145209 | 0.99979869 | 19598.0864 | 17382.6221 |
| *ECM1* | -0.178087016 | 0.35150492 | 0.99979869 | 10732.6106 | 9486.27616 |
| *PDH1* | -0.237308202 | 0.35154602 | 0.99979869 | 1498.39028 | 1271.1241 |
| *SMI1* | -0.170005054 | 0.35164746 | 0.99979869 | 37411.9568 | 33253.2216 |
| *YDL157C* | -0.177755633 | 0.3521236 | 0.99979869 | 10757.974 | 9510.87856 |
| *MST27* | -0.225037331 | 0.35247431 | 0.99979869 | 1842.7469 | 1576.60393 |
| *ITR1* | -0.168206305 | 0.35273217 | 0.99979869 | 61430.0995 | 54669.6125 |
| *RMD6* | -0.237923871 | 0.35302309 | 0.99979869 | 1460.34522 | 1238.3209 |
| *YNR040W* | -0.192472527 | 0.35355045 | 0.99979869 | 4363.47639 | 3818.4978 |
| *YBL101W-B* | -0.206640579 | 0.35367587 | 0.99979869 | 2748.02436 | 2381.30749 |
| *IGO2* | -0.182355042 | 0.35371085 | 0.99979869 | 7175.88471 | 6323.8424 |

| | | | | | |
|---|---|---|---|---|---|
| *snR78* | -0.524927246 | 0.35427994 | 0.99979869 | 184.372242 | 128.13751 |
| *HTS1* | -0.167411056 | 0.35442746 | 0.99979869 | 61598.8635 | 54850.0301 |
| *HTB1* | -0.166499555 | 0.35535886 | 0.99979869 | 81095.4964 | 72256.2295 |
| *TEL2* | -0.187050931 | 0.35587541 | 0.99979869 | 5237.53738 | 4600.64917 |
| *CBF2* | -0.18489983 | 0.35702123 | 0.99979869 | 5655.05759 | 4974.8107 |
| *TPC1* | -0.202997871 | 0.35717189 | 0.99979869 | 2876.79228 | 2499.194 |
| *IES6* | -0.172730546 | 0.35731931 | 0.99979869 | 13942.0533 | 12368.8576 |
| *PNT1* | -0.197232049 | 0.35763779 | 0.99979869 | 3397.71702 | 2963.56434 |
| *ZDS2* | -0.187719913 | 0.35802302 | 0.99979869 | 4827.82129 | 4238.78884 |
| *YNL208W* | -0.164772911 | 0.35867204 | 0.99979869 | 93953.7531 | 83813.2078 |
| *SOF1* | -0.174717515 | 0.35875258 | 0.99979869 | 10568.7242 | 9363.26415 |
| *TSR1* | -0.167768758 | 0.3588678 | 0.99979869 | 27674.371 | 24636.2303 |
| *snR46* | -0.276854645 | 0.35896634 | 0.99979869 | 832.113874 | 686.817055 |
| *MSI1* | -0.187325523 | 0.35926714 | 0.99979869 | 4775.14351 | 4193.68443 |
| *CWH41* | -0.167028251 | 0.36007 | 0.99979869 | 28905.4703 | 25745.3886 |
| *YER186C* | -0.177357996 | 0.36092072 | 0.99979869 | 7880.20618 | 6968.63036 |
| *IZH1* | -0.174724756 | 0.36152942 | 0.99979869 | 9405.91087 | 8333.03856 |
| *TOA2* | -0.170323112 | 0.36184924 | 0.99979869 | 14391.7655 | 12789.1486 |
| *RPT2* | -0.164364272 | 0.36212403 | 0.99979869 | 49447.8548 | 44123.3828 |
| *BMT5* | -0.181526802 | 0.36225048 | 0.99979869 | 5875.52387 | 5180.85581 |
| *ERG1* | -0.163585469 | 0.36273676 | 0.99979869 | 62293.4298 | 55615.7798 |
| *DAT1* | -0.177335431 | 0.36280974 | 0.99979869 | 7412.93473 | 6555.51502 |
| *NAT5* | -0.171884459 | 0.36293859 | 0.99979869 | 11441.8096 | 10156.6916 |
| *SFI1* | -0.177508959 | 0.3629964 | 0.99979869 | 7285.14233 | 6441.72891 |
| *CFD1* | -0.182167738 | 0.36351893 | 0.99979869 | 5498.97527 | 4846.67319 |
| *YBL071C* | -0.225401745 | 0.36392157 | 0.99979869 | 1604.72136 | 1372.60901 |
| *PEX28* | -0.182307 | 0.36396847 | 0.99979869 | 5399.47279 | 4758.51458 |
| *snR77* | -0.305358834 | 0.36401774 | 0.99979869 | 606.770023 | 491.022939 |
| *BUD27* | -0.169723256 | 0.36415769 | 0.99979869 | 13582.0885 | 12074.6539 |
| *RPL29* | -0.161859042 | 0.36416273 | 0.99979869 | 142154.9 | 127068.331 |
| *HAT1* | -0.189751628 | 0.36489579 | 0.99979869 | 3851.33127 | 3376.67967 |
| *RPC25* | -0.17700904 | 0.36505907 | 0.99979869 | 7051.99436 | 6237.734 |
| *RPL42A* | -0.161357536 | 0.36525521 | 0.99979869 | 144727.332 | 129412.735 |
| *RIT1* | -0.18673082 | 0.36527378 | 0.99979869 | 4297.1414 | 3775.4436 |
| *MNN11* | -0.166223542 | 0.36574924 | 0.99979869 | 20064.3824 | 17880.8207 |
| *PUG1* | -0.304420053 | 0.36578786 | 0.99979869 | 603.843479 | 488.972739 |
| *RPL24A* | -0.160770047 | 0.36608141 | 0.99979869 | 197204.159 | 176408.448 |
| *RTG3* | -0.175004867 | 0.3664628 | 0.99979869 | 7700.71151 | 6821.01594 |
| *RPC11* | -0.176310489 | 0.36647263 | 0.99979869 | 7057.84745 | 6245.9348 |
| *tR(ACG)L* | -1.58053598 | 0.36657687 | 0.99979869 | 21.4613191 | 7.17570057 |
| *RNT1* | -0.174214079 | 0.36694923 | 0.99979869 | 8009.94961 | 7098.81807 |
| *MAK16* | -0.164437916 | 0.36736655 | 0.99979869 | 24477.61 | 21840.7823 |
| *NMT1* | -0.166059065 | 0.36740975 | 0.99979869 | 18201.1497 | 16222.2088 |
| *RRP14* | -0.162911963 | 0.36750433 | 0.99979869 | 35225.8288 | 31464.4219 |
| *COQ8* | -0.175790148 | 0.36806296 | 0.99979869 | 6947.61431 | 6150.60049 |

| | | | | | |
|---|---|---|---|---|---|
| *DLT1* | -0.190621502 | 0.36830364 | 0.99979869 | 3505.02362 | 3071.19985 |
| *UBP16* | -0.191587677 | 0.36837465 | 0.99979869 | 3387.96188 | 2966.63964 |
| *TIM22* | -0.176682796 | 0.36856594 | 0.99979869 | 6480.34286 | 5733.38476 |
| *NOP1* | -0.159818377 | 0.36856607 | 0.99979869 | 159421.507 | 142704.182 |
| *RRP43* | -0.167637546 | 0.3686148 | 0.99979869 | 13614.2804 | 12120.7834 |
| *TIM54* | -0.165532119 | 0.36895406 | 0.99979869 | 17710.4659 | 15790.6417 |
| *YEH2* | -0.166167009 | 0.36903194 | 0.99979869 | 16055.0177 | 14308.3469 |
| *ARP1* | -0.182769316 | 0.36903286 | 0.99979869 | 4690.27375 | 4132.17843 |
| *RNH70* | -0.16750584 | 0.3692583 | 0.99979869 | 13352.8426 | 11889.1107 |
| *URM1* | -0.169047848 | 0.36927696 | 0.99979869 | 11292.5559 | 10043.9306 |
| *MHF2* | -0.189977893 | 0.36960208 | 0.99979869 | 3501.12156 | 3069.14964 |
| *FOL3* | -0.173534975 | 0.36976561 | 0.99979869 | 7643.15616 | 6776.93664 |
| *GUA1* | -0.159111523 | 0.37060613 | 0.99979869 | 127498.77 | 114184.873 |
| *HPT1* | -0.160875372 | 0.37075549 | 0.99979869 | 42807.5276 | 38290.5634 |
| *IME1* | -0.209829215 | 0.37084595 | 0.99979869 | 2009.55988 | 1737.54464 |
| *YMR122C* | -0.255528374 | 0.37095362 | 0.99979869 | 961.857303 | 805.728664 |
| *SPS2* | -0.291347663 | 0.37112133 | 0.99979869 | 653.594719 | 534.077143 |
| *YGR161W-B* | -0.636702325 | 0.37123824 | 0.99979869 | 113.159683 | 72.7821058 |
| *ATG41* | -0.167628253 | 0.37163686 | 0.99979869 | 11679.8352 | 10398.6152 |
| *YAH1* | -0.167691919 | 0.37178222 | 0.99979869 | 11523.7529 | 10259.2016 |
| *YJL113W* | -0.211087283 | 0.37216684 | 0.99979869 | 1924.69012 | 1662.71233 |
| *NDJ1* | -0.185025634 | 0.37231732 | 0.99979869 | 3987.9033 | 3507.89248 |
| *FAP7* | -0.169049774 | 0.37241114 | 0.99979869 | 9850.74548 | 8761.5304 |
| *SPO7* | -0.178227622 | 0.37268543 | 0.99979869 | 5338.99089 | 4718.53568 |
| *SMP1* | -0.197546387 | 0.37340626 | 0.99979869 | 2614.37888 | 2279.82258 |
| *RPL9A* | -0.1572356 | 0.3737978 | 0.99979869 | 271229.127 | 243222.421 |
| *MAF1* | -0.170170907 | 0.37383773 | 0.99979869 | 8499.65789 | 7553.9625 |
| *YHR022C-A* | -0.385154815 | 0.37421664 | 0.99979869 | 319.968758 | 244.99892 |
| *GSP1* | -0.157465116 | 0.37443913 | 0.99979869 | 125160.462 | 112218.731 |
| *LAP3* | -0.161133806 | 0.37461264 | 0.99979869 | 24843.4279 | 22218.0192 |
| *YNL050C* | -0.17500572 | 0.3752157 | 0.99979869 | 5936.98128 | 5258.76342 |
| *AAD3* | -0.190506533 | 0.37567446 | 0.99979869 | 3101.16061 | 2717.54032 |
| *AIM45* | -0.162176535 | 0.3760358 | 0.99979869 | 18211.8803 | 16275.514 |
| *SPC24* | -0.205323454 | 0.37655427 | 0.99979869 | 2067.11524 | 1792.90004 |
| *YIL046W-A* | -0.517681014 | 0.37726877 | 0.99979869 | 165.837466 | 115.836309 |
| *RRP4* | -0.166061715 | 0.37790153 | 0.99979869 | 10285.825 | 9167.47003 |
| *GPM2* | -0.188107865 | 0.37791219 | 0.99979869 | 3227.9775 | 2833.37663 |
| *NBP2* | -0.16896938 | 0.3780049 | 0.99979869 | 8043.1171 | 7154.17347 |
| *YDR169C-A* | -0.366461162 | 0.37805309 | 0.99979869 | 350.209708 | 271.651522 |
| *snR71* | -0.39323245 | 0.37828265 | 0.99979869 | 297.531924 | 226.547118 |
| *UTP5* | -0.159501016 | 0.37839838 | 0.99979869 | 24701.9783 | 22116.5343 |
| *SAS5* | -0.176718071 | 0.37904798 | 0.99979869 | 4942.932 | 4373.07695 |
| *YGR121W-A* | -0.347073858 | 0.37924265 | 0.99979869 | 395.083375 | 310.605325 |
| *MRPL36* | -0.166122378 | 0.37945585 | 0.99979869 | 9596.1362 | 8552.40998 |
| *RSR1* | -0.17135939 | 0.37984619 | 0.99979869 | 6465.71015 | 5741.58556 |

| | | | | | |
|---|---|---|---|---|---|
| *IMP3* | -0.170882642 | 0.37987476 | 0.99979869 | 6653.98445 | 5910.72707 |
| *MRPL32* | -0.171791885 | 0.3801036 | 0.99979869 | 6254.0235 | 5551.94204 |
| *CTR86* | -0.178508698 | 0.38041797 | 0.99979869 | 4420.05623 | 3905.63131 |
| *URA6* | -0.159961001 | 0.38084242 | 0.99979869 | 18517.2164 | 16573.8181 |
| *TSR4* | -0.160033601 | 0.38086468 | 0.99979869 | 18268.4602 | 16350.3463 |
| *UTP13* | -0.16017358 | 0.38089879 | 0.99979869 | 17817.7725 | 15945.4318 |
| *YGR027W-B* | -0.164483354 | 0.38118736 | 0.99979869 | 10381.4254 | 9262.80434 |
| *SLP1* | -0.169260771 | 0.38141229 | 0.99979869 | 7056.87194 | 6275.6627 |
| *MAK5* | -0.158483723 | 0.38203655 | 0.99979869 | 22002.7297 | 19713.6997 |
| *NUC1* | -0.167441518 | 0.38231126 | 0.99979869 | 7783.63024 | 6930.70165 |
| *MRP21* | -0.166869452 | 0.38280352 | 0.99979869 | 7986.53726 | 7114.19457 |
| *SAM3* | -0.159307348 | 0.38301568 | 0.99979869 | 17661.6901 | 15815.2441 |
| *PMT6* | -0.160976007 | 0.38314275 | 0.99979869 | 13848.4039 | 12386.2843 |
| *YML007C-A* | -0.388494535 | 0.38319047 | 0.99979869 | 296.55641 | 226.547118 |
| *snR39* | -0.258859689 | 0.38320408 | 0.99979869 | 834.064903 | 697.068056 |
| *DBP9* | -0.156573151 | 0.38359854 | 0.99979869 | 29056.6751 | 26068.2951 |
| *EFM6* | -0.202875615 | 0.3840976 | 0.99979869 | 1988.09856 | 1727.29364 |
| *MSW1* | -0.172872547 | 0.3843153 | 0.99979869 | 5283.38657 | 4686.75757 |
| *GRX6* | -0.173717778 | 0.38470222 | 0.99979869 | 5017.0711 | 4447.90925 |
| *RPS22B* | -0.154565134 | 0.38486044 | 0.99979869 | 46813.9656 | 42057.8062 |
| *SFH5* | -0.158616396 | 0.38568774 | 0.99979869 | 16440.346 | 14728.638 |
| *UIP5* | -0.165958038 | 0.38578582 | 0.99979869 | 7731.92798 | 6891.74785 |
| *YNL010W* | -0.154693844 | 0.38585789 | 0.99979869 | 38131.8865 | 34254.7443 |
| *RPL11B* | -0.152766114 | 0.38591589 | 0.99979869 | 106414.975 | 95722.8205 |
| *DSE1* | -0.154837789 | 0.38619818 | 0.99979869 | 34743.9246 | 31208.1469 |
| *LHP1* | -0.154287992 | 0.38635439 | 0.99979869 | 40861.3761 | 36717.0347 |
| *LYS1* | -0.163441786 | 0.38663279 | 0.99979869 | 9158.13018 | 8177.22335 |
| *LSR1* | -0.191840522 | 0.38691215 | 0.99979869 | 2513.90088 | 2200.88988 |
| *RBG1* | -0.154752626 | 0.38694795 | 0.99979869 | 32424.1512 | 29126.1686 |
| *TOM5* | -0.159963062 | 0.38751291 | 0.99979869 | 12436.8344 | 11131.5618 |
| *RPL39* | -0.15175195 | 0.38770019 | 0.99979869 | 143260.158 | 128956.565 |
| *IMP4* | -0.15949173 | 0.38821817 | 0.99979869 | 12677.7865 | 11350.9332 |
| *ADE13* | -0.151912969 | 0.38834573 | 0.99979869 | 92934.3405 | 83646.1165 |
| *snR40* | -0.30811345 | 0.38851379 | 0.99979869 | 497.512398 | 401.839232 |
| *NUG1* | -0.153137134 | 0.38855765 | 0.99979869 | 44585.8905 | 40095.7646 |
| *AMS1* | -0.172266484 | 0.38910374 | 0.99979869 | 4868.7929 | 4320.79684 |
| *SPG3* | -0.222835102 | 0.38924517 | 0.99979869 | 1289.63018 | 1105.05789 |
| *YNL018C* | -0.257867534 | 0.38933984 | 0.99979869 | 802.848439 | 671.440554 |
| *CSS1* | -0.151763808 | 0.38966596 | 0.99979869 | 70038.0395 | 63044.6801 |
| *COX17* | -0.163515531 | 0.38973335 | 0.99979869 | 8136.7665 | 7264.88428 |
| *GCV2* | -0.155671713 | 0.39010073 | 0.99979869 | 19542.4821 | 17543.5628 |
| *ERG6* | -0.151392761 | 0.39013511 | 0.99979869 | 78771.8209 | 70924.6245 |
| *ADY4* | -0.178658553 | 0.39031663 | 0.99979869 | 3621.10985 | 3199.33736 |
| *MRPL24* | -0.16744799 | 0.39077809 | 0.99979869 | 6045.26339 | 5382.80053 |
| *GFD1* | -0.16067065 | 0.39087041 | 0.99979869 | 9934.63973 | 8887.61771 |

| | | | | | |
|---|---|---|---|---|---|
| *VPS71* | -0.177246387 | 0.39159807 | 0.99979869 | 3732.3185 | 3300.82226 |
| *CPR8* | -0.160605532 | 0.39248247 | 0.99979869 | 9372.74337 | 8385.31867 |
| *NCE103* | -0.168691032 | 0.39274247 | 0.99979869 | 5361.42773 | 4769.79068 |
| *GCN3* | -0.15566229 | 0.39332773 | 0.99979869 | 15713.5877 | 14106.4022 |
| *CDC60* | -0.149524813 | 0.39390976 | 0.99979869 | 102171.487 | 92112.418 |
| *TIM10* | -0.152357535 | 0.39427428 | 0.99979869 | 27159.2994 | 24437.3608 |
| *SPT15* | -0.152068933 | 0.39508728 | 0.99979869 | 26845.1837 | 24159.5587 |
| *NOP53* | -0.153804653 | 0.39540363 | 0.99979869 | 18356.2565 | 16500.0109 |
| *MRM1* | -0.15891383 | 0.39567281 | 0.99979869 | 9641.00986 | 8635.44309 |
| *RPS25A* | -0.148122648 | 0.39605659 | 0.99979869 | 200901.359 | 181298.175 |
| *SPC2* | -0.164971662 | 0.396684 | 0.99979869 | 5980.87944 | 5334.62083 |
| *RPL11A* | -0.147751992 | 0.39721589 | 0.99979869 | 171515.936 | 154819.84 |
| *snR53* | -0.785103093 | 0.3973552 | 0.99979869 | 65.3594719 | 37.928703 |
| *JLP2* | -0.18748193 | 0.39737721 | 0.99979869 | 2445.61487 | 2147.58467 |
| *UBP10* | -0.151862782 | 0.39746977 | 0.99979869 | 22522.6789 | 20272.3792 |
| *ADE8* | -0.150975438 | 0.3976804 | 0.99979869 | 27035.409 | 24349.2022 |
| *RPA43* | -0.151908129 | 0.39776125 | 0.99979869 | 21770.5572 | 19594.7881 |
| *YEL073C* | -0.256406446 | 0.39788058 | 0.99979869 | 762.852344 | 638.637351 |
| *RPS31* | -0.146824318 | 0.39828999 | 0.99979869 | 753813.226 | 680871.474 |
| *SHU2* | -0.19473129 | 0.3984132 | 0.99979869 | 2008.58437 | 1754.97134 |
| *YSR3* | -0.167620634 | 0.39843241 | 0.99979869 | 4955.61369 | 4412.03075 |
| *PRM5* | -0.152438204 | 0.39970491 | 0.99979869 | 17130.0347 | 15412.3797 |
| *RPL20A* | -0.14654367 | 0.40011082 | 0.99979869 | 187089.05 | 169018.501 |
| *HRI1* | -0.148743878 | 0.40023886 | 0.99979869 | 38497.7045 | 34726.2904 |
| *EMP65* | -0.162841318 | 0.40042798 | 0.99979869 | 6154.52102 | 5497.61174 |
| *ENT4* | -0.163458812 | 0.40090859 | 0.99979869 | 5854.06255 | 5226.98532 |
| *YCR016W* | -0.15305565 | 0.40137946 | 0.99979869 | 14129.3521 | 12707.1406 |
| *YAT2* | -0.164560956 | 0.40199708 | 0.99979869 | 5350.69707 | 4773.89108 |
| *URA5* | -0.14700313 | 0.40247368 | 0.99979869 | 55049.2591 | 49716.3289 |
| *MTF1* | -0.172245734 | 0.40263641 | 0.99979869 | 3702.07755 | 3285.44576 |
| *RPP2A* | -0.14524044 | 0.40313131 | 0.99979869 | 230225.325 | 208176.3 |
| *OAF3* | -0.159921173 | 0.40344275 | 0.99979869 | 6831.52809 | 6114.72199 |
| *HXT1* | -0.145395927 | 0.40436141 | 0.99979869 | 97814.8395 | 88437.4343 |
| *RPS29B* | -0.145395438 | 0.40439911 | 0.99979869 | 96564.2299 | 87306.7489 |
| *YOR283W* | -0.153518874 | 0.40485432 | 0.99979869 | 11225.2454 | 10092.1103 |
| *ATP11* | -0.154031973 | 0.40492895 | 0.99979869 | 10595.0631 | 9522.15466 |
| *snR60* | -0.448336724 | 0.40521343 | 0.99979869 | 190.225329 | 139.413611 |
| *ELP2* | -0.147753586 | 0.4056993 | 0.99979869 | 26904.6901 | 24285.646 |
| *GPI17* | -0.150410608 | 0.40616638 | 0.99979869 | 15564.3339 | 14023.3691 |
| *NSA1* | -0.148765974 | 0.40618906 | 0.99979869 | 20569.6989 | 18554.3115 |
| *YPS3* | -0.148407515 | 0.40648939 | 0.99979869 | 21573.5033 | 19464.6004 |
| *tE(UUC)E1* | -1.201481632 | 0.40650964 | 0.99979869 | 28.2899207 | 12.301201 |
| *EMP47* | -0.148345476 | 0.40652539 | 0.99979869 | 21786.1655 | 19657.3192 |
| *MEH1* | -0.154361577 | 0.40653932 | 0.99979869 | 9598.08723 | 8624.16699 |
| *GNP1* | -0.145332967 | 0.4069295 | 0.99979869 | 52583.1584 | 47544.1418 |

| | | | | | |
|---|---|---|---|---|---|
| *RPS14B* | -0.14595727 | 0.40710539 | 0.99979869 | 39191.2953 | 35420.2831 |
| *SRP72* | -0.148220471 | 0.40711996 | 0.99979869 | 21286.702 | 19208.3253 |
| *GAT1* | -0.155346276 | 0.40730059 | 0.99979869 | 8540.6295 | 7668.77371 |
| *MNN9* | -0.147021132 | 0.40743224 | 0.99979869 | 27196.3689 | 24561.398 |
| *YPQ2* | -0.154458602 | 0.40756446 | 0.99979869 | 9143.49747 | 8215.15206 |
| *MRF1* | -0.156906149 | 0.40759087 | 0.99979869 | 7467.56354 | 6698.00393 |
| *STN1* | -0.1708603 | 0.40844268 | 0.99979869 | 3519.65634 | 3126.55525 |
| *YBL055C* | -0.151402549 | 0.40903829 | 0.99979869 | 11732.513 | 10563.6563 |
| *PET117* | -0.171597075 | 0.40910049 | 0.99979869 | 3384.05982 | 3004.56834 |
| *PWP1* | -0.144694313 | 0.40942956 | 0.99979869 | 45003.4107 | 40708.7744 |
| *MRP1* | -0.149560522 | 0.40958751 | 0.99979869 | 14404.4472 | 12985.9678 |
| *KRS1* | -0.142926438 | 0.40963671 | 0.99979869 | 156452.041 | 141695.484 |
| *ALT2* | -0.154907279 | 0.40997287 | 0.99979869 | 8072.38254 | 7250.53288 |
| *MMT1* | -0.158858041 | 0.41055024 | 0.99979869 | 6001.36524 | 5375.62483 |
| *PWP2* | -0.146145878 | 0.41055396 | 0.99979869 | 24873.6689 | 22477.3695 |
| *YMR320W* | -0.220258322 | 0.41078269 | 0.99979869 | 1097.45382 | 942.066975 |
| *YGR168C* | -0.169696245 | 0.41093584 | 0.99979869 | 3522.58288 | 3131.68075 |
| *YMR187C* | -0.182343055 | 0.41132733 | 0.99979869 | 2327.57761 | 2051.22526 |
| *SUP19* | -0.735822411 | 0.41192277 | 0.99979869 | 68.2860154 | 41.0040033 |
| *SDD2* | -0.148740489 | 0.41210104 | 0.99979869 | 14026.9231 | 12652.8103 |
| *YNR066C* | -0.194638878 | 0.41242172 | 0.99979869 | 1708.1259 | 1492.54572 |
| *snR3* | -0.241312291 | 0.41266409 | 0.99979869 | 810.652555 | 685.791955 |
| *ELO2* | -0.144141052 | 0.41285233 | 0.99979869 | 33736.2182 | 30528.5055 |
| *YJL118W* | -0.175540725 | 0.4131519 | 0.99979869 | 2770.4612 | 2453.0645 |
| *LIA1* | -0.142514318 | 0.41343393 | 0.99979869 | 61382.2993 | 55608.6041 |
| *LSB3* | -0.143465602 | 0.4135593 | 0.99979869 | 38745.4852 | 35077.8997 |
| *RRP3* | -0.147164174 | 0.41358865 | 0.99979869 | 16215.0021 | 14642.5296 |
| *STE20* | -0.144578212 | 0.41377677 | 0.99979869 | 26814.9427 | 24257.9683 |
| *LDB16* | -0.157565073 | 0.41427755 | 0.99979869 | 5893.08313 | 5283.36582 |
| *TEX1* | -0.161642979 | 0.41453406 | 0.99979869 | 4673.69 | 4178.30793 |
| *CLF1* | -0.152793989 | 0.41481612 | 0.99979869 | 8142.61958 | 7324.34008 |
| *PAU5* | -0.323399674 | 0.41486807 | 0.99979869 | 370.695512 | 296.253924 |
| *YJR029W* | -0.233608481 | 0.41527515 | 0.99979869 | 875.036512 | 744.222659 |
| *MSH1* | -0.157596164 | 0.41531743 | 0.99979869 | 5726.27015 | 5133.70121 |
| *ESF2* | -0.149281032 | 0.41670408 | 0.99979869 | 10523.8505 | 9489.35146 |
| *YDR286C* | -0.186853525 | 0.41690004 | 0.99979869 | 1933.46975 | 1698.59084 |
| *PDX3* | -0.143582363 | 0.41692242 | 0.99979869 | 25324.3566 | 22925.3382 |
| *THG1* | -0.150263838 | 0.41705758 | 0.99979869 | 9399.08227 | 8469.37688 |
| *tG(GCC)P2* | -0.964089397 | 0.41709796 | 0.99979869 | 39.9960948 | 20.5020016 |
| *TAD1* | -0.16103786 | 0.41733854 | 0.99979869 | 4534.19142 | 4055.29592 |
| *SPN1* | -0.145089766 | 0.41739851 | 0.99979869 | 17842.1603 | 16135.0753 |
| *EFM2* | -0.152792847 | 0.41766306 | 0.99979869 | 7417.8123 | 6672.37643 |
| *PRP21* | -0.164168374 | 0.41773991 | 0.99979869 | 3896.20494 | 3477.13948 |
| *YCR087C-A* | -0.145042839 | 0.41926084 | 0.99979869 | 15928.2009 | 14404.7063 |
| *TRM82* | -0.143620553 | 0.41964029 | 0.99979869 | 19933.6634 | 18044.8367 |

| | | | | | |
|---|---|---|---|---|---|
| *snR65* | -0.340559884 | 0.42002316 | 0.99979869 | 318.017729 | 251.14952 |
| *LYS4* | -0.141645037 | 0.42052458 | 0.99979869 | 29322.9905 | 26580.8451 |
| *RBG2* | -0.144218496 | 0.4210679 | 0.99979869 | 16237.439 | 14692.7595 |
| *PRS2* | -0.145484377 | 0.4212741 | 0.99979869 | 13320.6506 | 12042.8758 |
| *YGR021W* | -0.148154215 | 0.42246099 | 0.99979869 | 9344.45345 | 8432.47327 |
| *TUB3* | -0.139703112 | 0.42246641 | 0.99979869 | 44785.871 | 40652.3939 |
| *MRX3* | -0.159104355 | 0.42300266 | 0.99979869 | 4423.95829 | 3962.01182 |
| *PAM18* | -0.147547234 | 0.4232624 | 0.99979869 | 9610.76891 | 8676.44709 |
| *MFM1* | -0.16742019 | 0.42342109 | 0.99979869 | 3106.03819 | 2765.72002 |
| *EMW1* | -0.144251012 | 0.42357962 | 0.99979869 | 13945.9554 | 12618.982 |
| *INO4* | -0.158306608 | 0.42387545 | 0.99979869 | 4517.60768 | 4048.12022 |
| *PSH1* | -0.151942431 | 0.42406544 | 0.99979869 | 6519.36344 | 5867.67287 |
| *RRB1* | -0.1385717 | 0.42432176 | 0.99979869 | 55317.5256 | 50251.4311 |
| *RPA49* | -0.139784329 | 0.42457243 | 0.99979869 | 32354.8896 | 29367.0671 |
| *BET2* | -0.147118592 | 0.42461224 | 0.99979869 | 9510.29092 | 8588.28849 |
| *SHR3* | -0.139628204 | 0.42473145 | 0.99979869 | 33380.1554 | 30300.9333 |
| *SAH1* | -0.137052116 | 0.42522764 | 0.99979869 | 152719.722 | 138879.534 |
| *SPC3* | -0.146845039 | 0.42548161 | 0.99979869 | 9444.93145 | 8530.88288 |
| *MRX12* | -0.146988003 | 0.4258376 | 0.99979869 | 9189.34665 | 8299.21026 |
| *NAN1* | -0.139296403 | 0.4266084 | 0.99979869 | 29712.2208 | 26977.5589 |
| *MRPS17* | -0.149952657 | 0.4267286 | 0.99979869 | 6957.36946 | 6270.5372 |
| *YKL033W-A* | -0.139711521 | 0.42716347 | 0.99979869 | 25081.4535 | 22766.4477 |
| *RPL31A* | -0.136066361 | 0.42745155 | 0.99979869 | 203122.606 | 184840.921 |
| *HXK2* | -0.135977167 | 0.42755159 | 0.99979869 | 226731.033 | 206337.27 |
| *CCS1* | -0.137583512 | 0.4280744 | 0.99979869 | 45258.02 | 41141.3667 |
| *FEN2* | -0.145243584 | 0.42811916 | 0.99979869 | 9999.02369 | 9041.38272 |
| *IPA1* | -0.158378335 | 0.42847503 | 0.99979869 | 4099.11195 | 3672.93359 |
| *RSM26* | -0.145340395 | 0.42860689 | 0.99979869 | 9708.32036 | 8777.932 |
| *HOS1* | -0.169526231 | 0.42865996 | 0.99979869 | 2669.00769 | 2373.10669 |
| *RNR4* | -0.136068178 | 0.42958885 | 0.99979869 | 79120.0795 | 71998.9293 |
| *ALR2* | -0.158948363 | 0.42974214 | 0.99979869 | 3898.15597 | 3491.49088 |
| *ENP2* | -0.138302532 | 0.42974716 | 0.99979869 | 28207.0019 | 25628.5271 |
| *ROK1* | -0.140940308 | 0.42996499 | 0.99979869 | 15878.4496 | 14400.6059 |
| *BUD22* | -0.1402182 | 0.43106471 | 0.99979869 | 16707.6369 | 15160.2051 |
| *IKI1* | -0.148749432 | 0.43124302 | 0.99979869 | 6654.95996 | 6002.98608 |
| *TAF9* | -0.140829095 | 0.43131112 | 0.99979869 | 14909.7637 | 13523.1203 |
| *YMC1* | -0.141974562 | 0.43216503 | 0.99979869 | 12154.9107 | 11015.7255 |
| *FMP41* | -0.1422002 | 0.4323975 | 0.99979869 | 11689.5903 | 10592.3591 |
| *RPS28B* | -0.135566114 | 0.43249492 | 0.99979869 | 54896.1033 | 49972.6039 |
| *RPS2* | -0.133799317 | 0.4327371 | 0.99979869 | 647331.916 | 589996.352 |
| *SLM3* | -0.156073101 | 0.43387422 | 0.99979869 | 4103.98953 | 3683.18459 |
| *DDR48* | -0.134758219 | 0.43396974 | 0.99979869 | 64467.8517 | 58718.7578 |
| *BNA7* | -0.151175766 | 0.43436596 | 0.99979869 | 5244.36598 | 4722.63608 |
| *KRR1* | -0.136812108 | 0.43454735 | 0.99979869 | 26065.7476 | 23707.4896 |
| *MNL2* | -0.141108884 | 0.43625817 | 0.99979869 | 11168.6656 | 10127.9888 |

| | | | | | |
|---|---|---|---|---|---|
| *TCD1* | -0.140175358 | 0.43628376 | 0.99979869 | 12529.5083 | 11369.385 |
| *YDL177C* | -0.171049176 | 0.43662474 | 0.99979869 | 2281.72843 | 2026.62286 |
| *YET2* | -0.19404897 | 0.43756813 | 0.99979869 | 1328.65076 | 1161.43839 |
| *YRF1-8* | -0.761357036 | 0.4377892 | 0.99979869 | 55.6043268 | 32.8032026 |
| *ATG40* | -0.16339927 | 0.43781022 | 0.99979869 | 2856.30647 | 2550.449 |
| *YAP7* | -0.150579904 | 0.43808676 | 0.99979869 | 4977.07501 | 4483.78776 |
| *YEL067C* | -0.202731765 | 0.43815618 | 0.99979869 | 1132.57234 | 984.096079 |
| *SSS1* | -0.135219627 | 0.43846547 | 0.99979869 | 27202.222 | 24768.4682 |
| *GLE2* | -0.13819142 | 0.43848435 | 0.99979869 | 14780.9958 | 13430.8613 |
| *MSS2* | -0.161176982 | 0.43853056 | 0.99979869 | 3058.23798 | 2734.96702 |
| *MRPL39* | -0.145775273 | 0.43889936 | 0.99979869 | 6627.64555 | 5990.68488 |
| *GRC3* | -0.145526874 | 0.43931392 | 0.99979869 | 6669.59268 | 6029.63868 |
| *BNA5* | -0.138252931 | 0.43942902 | 0.99979869 | 13890.3511 | 12621.0322 |
| *HHF2* | -0.132262876 | 0.43951358 | 0.99979869 | 95415.0738 | 87056.6244 |
| *DBR1* | -0.155577639 | 0.43997073 | 0.99979869 | 3737.19607 | 3355.15257 |
| *TIF3* | -0.131762739 | 0.44013327 | 0.99979869 | 130658.462 | 119253.993 |
| *DIP2* | -0.13500215 | 0.4403482 | 0.99979869 | 24047.4081 | 21899.213 |
| *MNP1* | -0.135255201 | 0.44048404 | 0.99979869 | 22361.719 | 20360.5378 |
| *YMR265C* | -0.141835929 | 0.44090839 | 0.99979869 | 8600.13589 | 7794.86102 |
| *ARP10* | -0.271071546 | 0.44096715 | 0.99979869 | 479.953137 | 397.738832 |
| *HAP1* | -0.132173189 | 0.44168493 | 0.99979869 | 57346.5958 | 52326.2337 |
| *ITT1* | -0.149377087 | 0.4418172 | 0.99979869 | 4892.20525 | 4411.00565 |
| *YOR314W* | -0.461137021 | 0.44183565 | 0.99979869 | 145.351661 | 105.585308 |
| *UTP8* | -0.135115852 | 0.44224901 | 0.99979869 | 20014.6311 | 18225.2544 |
| *SNZ3* | -0.201383418 | 0.44235694 | 0.99979869 | 1115.01308 | 969.744677 |
| *RGT2* | -0.140801495 | 0.44334214 | 0.99979869 | 8688.90771 | 7880.96943 |
| *GEP3* | -0.149010852 | 0.44348672 | 0.99979869 | 4810.26203 | 4338.22355 |
| *SET4* | -0.187363117 | 0.44360341 | 0.99979869 | 1425.22669 | 1251.6472 |
| *YDR524W-A* | -0.131203874 | 0.44420488 | 0.99979869 | 59975.6074 | 54761.8715 |
| *YDR210W-B* | -0.173963997 | 0.44436413 | 0.99979869 | 1913.95946 | 1696.54064 |
| *YER137C* | -0.190864901 | 0.4444033 | 0.99979869 | 1319.87113 | 1156.31289 |
| *SLO1* | -0.179847383 | 0.44445944 | 0.99979869 | 1659.35017 | 1464.86802 |
| *IRC19* | -0.179288899 | 0.44476875 | 0.99979869 | 1674.95841 | 1479.21942 |
| *SCC4* | -0.144220641 | 0.44484445 | 0.99979869 | 6271.58276 | 5674.95405 |
| *YLL054C* | -0.151031429 | 0.44528425 | 0.99979869 | 4169.349 | 3754.9416 |
| *APM4* | -0.135263889 | 0.44531917 | 0.99979869 | 15793.5799 | 14380.1039 |
| *RPL22B* | -0.13588069 | 0.44542997 | 0.99979869 | 14220.075 | 12941.8885 |
| *LCB2* | -0.130862197 | 0.44580003 | 0.99979869 | 52736.3142 | 48163.3022 |
| *GAR1* | -0.130265911 | 0.44625647 | 0.99979869 | 69143.4927 | 63173.8427 |
| *YDR274C* | -0.167962849 | 0.44641299 | 0.99979869 | 2193.93212 | 1952.81566 |
| *HIF1* | -0.137838132 | 0.44686772 | 0.99979869 | 10290.7025 | 9353.01315 |
| *YHR175W-A* | -0.239877511 | 0.44714089 | 0.99979869 | 633.108914 | 536.127343 |
| *YDR109C* | -0.145074158 | 0.44770857 | 0.99979869 | 5499.95079 | 4973.7856 |
| *YJL027C* | -0.519828931 | 0.44792084 | 0.99979869 | 110.233139 | 76.8825061 |
| *PLP1* | -0.156405991 | 0.44813254 | 0.99979869 | 3137.25465 | 2814.92482 |

| | | | | | |
|---|---|---|---|---|---|
| *AQY2* | -0.216845584 | 0.4495564 | 0.99979869 | 829.18733 | 713.469657 |
| *CAF20* | -0.130162834 | 0.44995704 | 0.99979869 | 37137.8372 | 33933.888 |
| *PRO2* | -0.129411419 | 0.45092214 | 0.99979869 | 44282.5055 | 40483.2524 |
| *PGU1* | -0.16060045 | 0.45097607 | 0.99979869 | 2586.08896 | 2313.65088 |
| *RSN1* | -0.129609549 | 0.45104142 | 0.99979869 | 39834.1593 | 36411.5549 |
| *STB4* | -0.138050059 | 0.45179061 | 0.99979869 | 8347.47763 | 7585.74061 |
| *AIM46* | -0.133712815 | 0.45182073 | 0.99979869 | 13881.5714 | 12652.8103 |
| *BRX1* | -0.130101744 | 0.4518912 | 0.99979869 | 29893.6665 | 27315.8419 |
| *MED6* | -0.146405375 | 0.45238656 | 0.99979869 | 4571.26098 | 4130.12823 |
| *TRP5* | -0.127709384 | 0.45399925 | 0.99979869 | 62861.1793 | 57535.7923 |
| *YLR406C-A* | -0.972858876 | 0.45414146 | 0.99979869 | 32.1919787 | 16.4016013 |
| *GCV3* | -0.128645659 | 0.45419979 | 0.99979869 | 37663.6396 | 34450.5384 |
| *XPT1* | -0.136922663 | 0.45468965 | 0.99979869 | 8389.42475 | 7629.81991 |
| *YLF2* | -0.144939052 | 0.45520301 | 0.99979869 | 4662.95934 | 4217.26174 |
| *DJP1* | -0.130263161 | 0.4558682 | 0.99979869 | 19909.2756 | 18190.401 |
| *EST1* | -0.153608009 | 0.45601027 | 0.99979869 | 3078.72378 | 2767.77022 |
| *RPS17B* | -0.126060811 | 0.45611975 | 0.99979869 | 161386.193 | 147882.988 |
| *POP3* | -0.149691146 | 0.45633839 | 0.99979869 | 3602.57507 | 3247.51706 |
| *SPO74* | -0.301928127 | 0.4566216 | 0.99979869 | 333.625961 | 270.626422 |
| *DSS4* | -0.152724897 | 0.45672119 | 0.99979869 | 3150.91185 | 2834.40173 |
| *SGF29* | -0.135592011 | 0.45673271 | 0.99979869 | 8877.18201 | 8080.86394 |
| *YNL042W-B* | -0.43895336 | 0.45724223 | 0.99979869 | 147.30269 | 108.660609 |
| *YDR316W-B* | -0.266165279 | 0.45761452 | 0.99979869 | 448.736673 | 373.13643 |
| *XRS2* | -0.14339398 | 0.45793015 | 0.99979869 | 4792.70277 | 4339.24865 |
| *AAC1* | -0.158691838 | 0.4580008 | 0.99979869 | 2493.41508 | 2233.69308 |
| *ARL1* | -0.131003289 | 0.45817963 | 0.99979869 | 14873.6697 | 13582.5761 |
| *MAK31* | -0.143169298 | 0.45840214 | 0.99979869 | 4804.40894 | 4350.52475 |
| *HVG1* | -0.206004045 | 0.45893315 | 0.99979869 | 897.473346 | 778.050962 |
| *YOL029C* | -0.147738437 | 0.45901127 | 0.99979869 | 3749.87776 | 3384.88047 |
| *AVT7* | -0.128943679 | 0.45911553 | 0.99979869 | 20730.6588 | 18958.2009 |
| *DHR2* | -0.135773745 | 0.46017056 | 0.99979869 | 7747.53621 | 7051.66346 |
| *EAF5* | -0.138074992 | 0.46032672 | 0.99979869 | 6391.57104 | 5808.21706 |
| *DPH5* | -0.127178699 | 0.46045855 | 0.99979869 | 29340.5498 | 26864.7978 |
| *SHB17* | -0.125512857 | 0.4606586 | 0.99979869 | 59388.3476 | 54439.99 |
| *TRP3* | -0.127500396 | 0.46084381 | 0.99979869 | 25624.815 | 23457.3652 |
| *MRPL13* | -0.132639285 | 0.46100971 | 0.99979869 | 10334.6007 | 9426.82035 |
| *OLE1* | -0.124543173 | 0.46105895 | 0.99979869 | 126487.162 | 116025.953 |
| *SEY1* | -0.128049698 | 0.46141669 | 0.99979869 | 21187.1996 | 19387.7178 |
| *MAK21* | -0.125404575 | 0.46181848 | 0.99979869 | 50229.2419 | 46047.4957 |
| *YIP1* | -0.131068795 | 0.46200926 | 0.99979869 | 12039.8 | 10994.1984 |
| *YDR381C-A* | -0.134797312 | 0.46237004 | 0.99979869 | 7869.47552 | 7167.49977 |
| *PAU2* | -0.257931656 | 0.46289931 | 0.99979869 | 468.246963 | 391.588231 |
| *FMC1* | -0.141669024 | 0.4635065 | 0.99979869 | 4679.54309 | 4241.86414 |
| *RPS16A* | -0.123461833 | 0.46373773 | 0.99979869 | 167984.573 | 154206.83 |
| *SNU13* | -0.12511476 | 0.46383336 | 0.99979869 | 41379.3743 | 37942.0293 |

| | | | | | |
|---|---|---|---|---|---|
| *DYN3* | -0.135024438 | 0.46393397 | 0.99979869 | 7334.89357 | 6679.55213 |
| *UFO1* | -0.130104105 | 0.46394225 | 0.99979869 | 12519.7532 | 11440.1169 |
| *POL3* | -0.132859607 | 0.46414061 | 0.99979869 | 8921.08016 | 8136.21935 |
| *BMT2* | -0.143821326 | 0.46445608 | 0.99979869 | 4087.40578 | 3699.5862 |
| *SUV3* | -0.136060474 | 0.46537084 | 0.99979869 | 6463.75912 | 5882.02427 |
| *SAL1* | -0.139974337 | 0.46537476 | 0.99979869 | 4957.56472 | 4499.16426 |
| *RPS25B* | -0.123217393 | 0.46562792 | 0.99979869 | 98776.6968 | 90690.6042 |
| *NHP10* | -0.134218906 | 0.46587454 | 0.99979869 | 7398.30201 | 6741.05814 |
| *SIR2* | -0.136486767 | 0.46639217 | 0.99979869 | 6088.18603 | 5538.61574 |
| *RRP17* | -0.128428109 | 0.46652382 | 0.99979869 | 14057.164 | 12859.8805 |
| *BNA1* | -0.132231611 | 0.46688362 | 0.99979869 | 8605.98897 | 7852.26663 |
| *DRS1* | -0.124076227 | 0.46722542 | 0.99979869 | 39491.7538 | 36237.2879 |
| *RKM4* | -0.131413472 | 0.46722949 | 0.99979869 | 9247.87752 | 8442.72427 |
| *TPN1* | -0.126317566 | 0.46732985 | 0.99979869 | 19784.4097 | 18125.8196 |
| *SPC97* | -0.144138523 | 0.46803175 | 0.99979869 | 3758.65739 | 3401.28207 |
| *PUP2* | -0.123777834 | 0.46806864 | 0.99979869 | 39756.1182 | 36487.4123 |
| *TRI1* | -0.142225771 | 0.46823945 | 0.99979869 | 4115.6957 | 3729.3141 |
| *SHM2* | -0.122282142 | 0.46839209 | 0.99979869 | 100494.578 | 92327.6891 |
| *TRM2* | -0.128768895 | 0.46869914 | 0.99979869 | 11957.8568 | 10936.7928 |
| *CSN9* | -0.164098425 | 0.46886302 | 0.99979869 | 1851.52653 | 1652.46133 |
| *RPL23B* | -0.12149664 | 0.4690492 | 0.99979869 | 265745.76 | 244282.375 |
| *EEB1* | -0.138520333 | 0.46940501 | 0.99979869 | 4939.02994 | 4486.86306 |
| *CUE3* | -0.130192698 | 0.46989567 | 0.99979869 | 9531.75224 | 8709.25029 |
| *PAM17* | -0.127574162 | 0.47012066 | 0.99979869 | 13227.9767 | 12108.4822 |
| *tF(GAA)H2* | -1.291030175 | 0.4702725 | 0.99979869 | 17.5592611 | 7.17570057 |
| *RTC4* | -0.143449082 | 0.47077336 | 0.99979869 | 3692.32241 | 3342.85137 |
| *YLR361C-A* | -0.139872871 | 0.47134148 | 0.99979869 | 4380.06013 | 3975.33812 |
| *ELM1* | -0.145700934 | 0.4720109 | 0.99979869 | 3266.02257 | 2952.28824 |
| *YDR061W* | -0.129674202 | 0.47238989 | 0.99979869 | 9164.95878 | 8377.11787 |
| *YAR029W* | -0.178035525 | 0.47260349 | 0.99979869 | 1289.63018 | 1139.91129 |
| *MIX23* | -0.135230652 | 0.47305771 | 0.99979869 | 5596.52672 | 5095.77251 |
| *ISN1* | -0.127851048 | 0.47344453 | 0.99979869 | 10860.403 | 9939.37039 |
| *YPS7* | -0.123603554 | 0.47354461 | 0.99979869 | 22886.5458 | 21007.376 |
| *CWC2* | -0.144145172 | 0.47397725 | 0.99979869 | 3379.18225 | 3057.87354 |
| *RDN37-2* | -0.691427996 | 0.47455792 | 0.99979869 | 54.6288123 | 33.8283027 |
| *TMA20* | -0.124216048 | 0.47478995 | 0.99979869 | 18045.0673 | 16556.3914 |
| *VHT1* | -0.135265803 | 0.47486532 | 0.99979869 | 5339.96641 | 4862.04969 |
| *YML096W* | -0.131483133 | 0.47506964 | 0.99979869 | 7045.16576 | 6431.47791 |
| *YBR196C-B* | -0.196498673 | 0.47560966 | 0.99979869 | 903.326433 | 788.301963 |
| *GNT1* | -0.124912691 | 0.47650755 | 0.99979869 | 14261.0466 | 13078.2268 |
| *PGA2* | -0.122309599 | 0.47662235 | 0.99979869 | 24588.8186 | 22590.1305 |
| *TAF13* | -0.131314439 | 0.47670365 | 0.99979869 | 6805.18919 | 6213.1316 |
| *MKC7* | -0.122411695 | 0.47684721 | 0.99979869 | 23428.9319 | 21523.0013 |
| *KTR5* | -0.131487881 | 0.4771959 | 0.99979869 | 6613.98835 | 6037.83948 |
| *RIF2* | -0.158628181 | 0.47755877 | 0.99979869 | 1938.34732 | 1736.51954 |

| | | | | | |
|---|---|---|---|---|---|
| *RPC34* | -0.126282059 | 0.47764218 | 0.99979869 | 11032.0935 | 10107.4868 |
| *YNL040W* | -0.127199531 | 0.47777198 | 0.99979869 | 9801.96976 | 8974.75122 |
| *AHK1* | -0.126243425 | 0.4779067 | 0.99979869 | 10957.9544 | 10039.8302 |
| *BUD17* | -0.126933217 | 0.47794816 | 0.99979869 | 10041.9463 | 9196.17283 |
| *SDA1* | -0.120313746 | 0.4781835 | 0.99979869 | 41875.9112 | 38525.3113 |
| *TFB3* | -0.131626688 | 0.47833121 | 0.99979869 | 6334.9912 | 5782.58956 |
| *ERB1* | -0.120812922 | 0.47833285 | 0.99979869 | 33239.6813 | 30569.5095 |
| *RIO1* | -0.126633161 | 0.47847643 | 0.99979869 | 10185.347 | 9329.43584 |
| *NBP35* | -0.128943701 | 0.4787327 | 0.99979869 | 7880.20618 | 7206.45357 |
| *ECT1* | -0.123750823 | 0.47893458 | 0.99979869 | 15118.5238 | 13875.7547 |
| *MAS2* | -0.122849935 | 0.48023243 | 0.99979869 | 16468.6359 | 15124.3266 |
| *HBN1* | -0.169839104 | 0.48049345 | 0.99979869 | 1418.39809 | 1260.8731 |
| *PTH4* | -0.183896869 | 0.48055605 | 0.99979869 | 1070.13941 | 942.066975 |
| *RTC6* | -0.133899827 | 0.48065615 | 0.99979869 | 5086.33263 | 4635.50257 |
| *SSO2* | -0.120686012 | 0.48086734 | 0.99979869 | 26221.8299 | 24117.5296 |
| *THS1* | -0.117816643 | 0.48101673 | 0.99979869 | 164255.182 | 151374.479 |
| *YCL019W* | -0.821550691 | 0.48115648 | 0.99979869 | 38.0450657 | 21.5271017 |
| *SEC66* | -0.123854172 | 0.48168677 | 0.99979869 | 12785.0931 | 11733.2955 |
| *GTR2* | -0.126391627 | 0.48173906 | 0.99979869 | 9200.07731 | 8428.37287 |
| *MRI1* | -0.12002914 | 0.48206449 | 0.99979869 | 28494.7787 | 26220.0099 |
| *YLL053C* | -0.201048233 | 0.4821124 | 0.99979869 | 798.946381 | 695.017855 |
| *APC2* | -0.130500688 | 0.48221189 | 0.99979869 | 6211.10086 | 5673.92895 |
| *FPR2* | -0.124107341 | 0.48250048 | 0.99979869 | 11838.844 | 10862.9856 |
| *HSH155* | -0.127397485 | 0.48301467 | 0.99979869 | 7922.1533 | 7252.58308 |
| *MSG5* | -0.131409004 | 0.48314853 | 0.99979869 | 5664.81274 | 5171.62991 |
| *RPS24B* | -0.117381347 | 0.48321448 | 0.99979869 | 109646.855 | 101078.968 |
| *MIM2* | -0.132751284 | 0.48342128 | 0.99979869 | 5135.10836 | 4683.68227 |
| *YGR039W* | -0.22881752 | 0.48383967 | 0.99979869 | 550.190181 | 469.495837 |
| *RPS19A* | -0.116996023 | 0.48394089 | 0.99979869 | 135443.361 | 124893.068 |
| *RRP40* | -0.126449123 | 0.48411277 | 0.99979869 | 8383.57167 | 7680.04981 |
| *PRI2* | -0.129427046 | 0.48511352 | 0.99979869 | 6234.51321 | 5699.55645 |
| *PGI1* | -0.115983664 | 0.48574676 | 0.99979869 | 415226.774 | 383151.658 |
| *PEP12* | -0.122144177 | 0.48576741 | 0.99979869 | 13488.4391 | 12393.46 |
| *RDN37-1* | -0.634736342 | 0.48577949 | 0.99979869 | 60.4818994 | 38.9538031 |
| *IRC15* | -0.131217922 | 0.48581218 | 0.99979869 | 5372.15839 | 4905.10389 |
| *FPR1* | -0.116769135 | 0.48586607 | 0.99979869 | 84851.2273 | 78254.09 |
| *RSC58* | -0.122955408 | 0.48678979 | 0.99979869 | 11370.5971 | 10441.6694 |
| *CDD1* | -0.145545178 | 0.48710417 | 0.99979869 | 2597.79513 | 2348.50429 |
| *snR161* | -0.37370703 | 0.48765097 | 0.99979869 | 172.666068 | 133.263011 |
| *NOP8* | -0.124919732 | 0.48847909 | 0.99979869 | 8413.81262 | 7715.92832 |
| *FCF1* | -0.123296544 | 0.48860263 | 0.99979869 | 10054.628 | 9231.02624 |
| *NAF1* | -0.123061706 | 0.48863898 | 0.99979869 | 10336.5517 | 9491.40166 |
| *MAK11* | -0.119818952 | 0.48873356 | 0.99979869 | 17207.1004 | 15835.7461 |
| *YBR242W* | -0.125325502 | 0.48897513 | 0.99979869 | 7938.73705 | 7278.21058 |
| *YDL121C* | -0.119722462 | 0.48944931 | 0.99979869 | 16735.9269 | 15403.1538 |

| | | | | | |
|---|---|---|---|---|---|
| *MTQ2* | -0.131647519 | 0.48952363 | 0.99979869 | 4789.77622 | 4372.05185 |
| *SPB4* | -0.121131056 | 0.48985157 | 0.99979869 | 12782.1666 | 11752.7724 |
| *HMS2* | -0.120248525 | 0.4900645 | 0.99979869 | 14608.3297 | 13440.0872 |
| *YGL101W* | -0.128937907 | 0.49008947 | 0.99979869 | 5669.69031 | 5184.95621 |
| *MRX10* | -0.135696771 | 0.49013717 | 0.99979869 | 3769.38805 | 3431.00997 |
| *RRS1* | -0.115286583 | 0.49031425 | 0.99979869 | 91360.8356 | 84344.2096 |
| *BOL3* | -0.129770058 | 0.49071032 | 0.99979869 | 5265.8273 | 4812.84488 |
| *YPR202W* | -0.193537869 | 0.49122267 | 0.99979869 | 831.138359 | 726.795958 |
| *MAN2* | -0.223348698 | 0.49170049 | 0.99979869 | 554.092239 | 474.621338 |
| *SEC1* | -0.121693274 | 0.49174706 | 0.99979869 | 10786.2639 | 9913.74289 |
| *OSW7* | -0.121025102 | 0.49303496 | 0.99979869 | 11141.3512 | 10244.8502 |
| *ELO3* | -0.115102146 | 0.49305941 | 0.99979869 | 53018.2379 | 48952.6293 |
| *ECM14* | -0.116645441 | 0.49317755 | 0.99979869 | 26676.4197 | 24604.4522 |
| *FCF2* | -0.12155535 | 0.49381392 | 0.99979869 | 10053.6525 | 9241.27724 |
| *CPR7* | -0.121733634 | 0.49401211 | 0.99979869 | 9757.09609 | 8967.57552 |
| *COX18* | -0.130791854 | 0.4947366 | 0.99979869 | 4502.97496 | 4112.70153 |
| *REX3* | -0.1275171 | 0.49483972 | 0.99979869 | 5571.16334 | 5099.87291 |
| *KIN4* | -0.122433078 | 0.49490947 | 0.99979869 | 8700.61388 | 7992.70534 |
| *PBR1* | -0.120255045 | 0.49522459 | 0.99979869 | 11214.5148 | 10317.6323 |
| *PLM2* | -0.155362714 | 0.49525097 | 0.99979869 | 1722.75862 | 1546.87602 |
| *SPP2* | -0.131891298 | 0.49529471 | 0.99979869 | 4176.1776 | 3811.3221 |
| *TRM12* | -0.131780283 | 0.49531569 | 0.99979869 | 4200.56546 | 3833.87431 |
| *EBP2* | -0.114202292 | 0.49566547 | 0.99979869 | 55851.132 | 51600.4628 |
| *MRPL44* | -0.126121555 | 0.49579838 | 0.99979869 | 6034.53273 | 5529.38984 |
| *RPL13B* | -0.112662222 | 0.4966758 | 0.99979869 | 284129.331 | 262785.431 |
| *TOG1* | -0.127591818 | 0.49672524 | 0.99979869 | 5289.23965 | 4841.54769 |
| *PLP2* | -0.118477146 | 0.49673349 | 0.99979869 | 13628.9132 | 12554.4007 |
| *PDR16* | -0.115392631 | 0.49697676 | 0.99979869 | 27068.5765 | 24987.8396 |
| *PEX11* | -0.128077473 | 0.49714659 | 0.99979869 | 5065.84683 | 4635.50257 |
| *AIF1* | -0.126028353 | 0.49715973 | 0.99979869 | 5861.86667 | 5371.52443 |
| *tK(CUU)K* | -1.065966151 | 0.4973018 | 0.99979869 | 21.4613191 | 10.2510008 |
| *DPB4* | -0.122288949 | 0.49752406 | 0.99979869 | 8054.82328 | 7400.19749 |
| *TSR3* | -0.119966106 | 0.49760805 | 0.99979869 | 10492.634 | 9655.41767 |
| *FCY1* | -0.114148926 | 0.4978001 | 0.99979869 | 39702.4649 | 36682.1813 |
| *snR14* | -0.295527116 | 0.4985026 | 0.99979869 | 269.242004 | 219.371418 |
| *ALG5* | -0.119662166 | 0.49912261 | 0.99979869 | 10233.1472 | 9418.61955 |
| *SER2* | -0.115956324 | 0.49919561 | 0.99979869 | 19139.5946 | 17661.4493 |
| *RAD55* | -0.147660183 | 0.49941007 | 0.99979869 | 2048.58046 | 1849.28055 |
| *TPO5* | -0.117095663 | 0.49945063 | 0.99979869 | 14884.4003 | 13724.0399 |
| *CCM1* | -0.12153615 | 0.49953178 | 0.99979869 | 8123.10929 | 7466.829 |
| *CDC73* | -0.118148463 | 0.49969359 | 0.99979869 | 12320.7482 | 11351.9583 |
| *MIH1* | -0.129773922 | 0.50021321 | 0.99979869 | 4262.02288 | 3895.38031 |
| *ALG11* | -0.118454259 | 0.50081408 | 0.99979869 | 11171.5921 | 10290.9797 |
| *YLH47* | -0.118455475 | 0.5012544 | 0.99979869 | 10950.1503 | 10086.9848 |
| *CPR2* | -0.116621365 | 0.50168477 | 0.99979869 | 14249.3404 | 13142.8081 |

| | | | | | |
|---|---|---|---|---|---|
| *JNM1* | -0.129436645 | 0.50168986 | 0.99979869 | 4216.1737 | 3854.37631 |
| *MHR1* | -0.118355643 | 0.50177865 | 0.99979869 | 10844.7948 | 9990.6254 |
| *HSH49* | -0.132227835 | 0.50188546 | 0.99979869 | 3614.28125 | 3297.74696 |
| *TPM1* | -0.111456214 | 0.5020358 | 0.99979869 | 114182.022 | 105692.944 |
| *SAM1* | -0.111353934 | 0.50205134 | 0.99979869 | 131620.319 | 121843.396 |
| *UAF30* | -0.137397825 | 0.50217383 | 0.99979869 | 2843.62479 | 2585.30241 |
| *SYF1* | -0.120210622 | 0.50242998 | 0.99979869 | 8442.10254 | 7767.18332 |
| *VMA16* | -0.112615597 | 0.50254472 | 0.99979869 | 40777.4819 | 37715.4822 |
| *TIF35* | -0.11142677 | 0.50312785 | 0.99979869 | 80618.4698 | 74626.2609 |
| *ATP23* | -0.131144417 | 0.50330126 | 0.99979869 | 3721.58784 | 3398.20677 |
| *HMRA1* | -0.118698544 | 0.50351809 | 0.99979869 | 9641.98538 | 8880.44201 |
| *ENV9* | -0.11420757 | 0.50354961 | 0.99979869 | 20963.8067 | 19368.2409 |
| *YEH1* | -0.123486227 | 0.50355758 | 0.99979869 | 6025.7531 | 5531.44004 |
| *SLC1* | -0.114524524 | 0.50386566 | 0.99979869 | 18930.8345 | 17486.1572 |
| *THI20* | -0.12578391 | 0.50463948 | 0.99979869 | 4960.49126 | 4546.31886 |
| *YCR051W* | -0.112266063 | 0.50503078 | 0.99979869 | 33771.3367 | 31243.0003 |
| *YML131W* | -0.127051698 | 0.50521555 | 0.99979869 | 4513.70562 | 4133.20353 |
| *HO* | -0.121261648 | 0.50556235 | 0.99979869 | 6870.54867 | 6316.6667 |
| *ARO7* | -0.117994098 | 0.50565692 | 0.99979869 | 9651.74052 | 8893.76831 |
| *ECM22* | -0.117079088 | 0.50582829 | 0.99979869 | 10794.068 | 9952.69669 |
| *FAA2* | -0.134112049 | 0.50604611 | 0.99979869 | 3076.77275 | 2803.64872 |
| *ANT1* | -0.118441194 | 0.50648672 | 0.99979869 | 8863.5248 | 8164.92215 |
| *YMR316C-A* | -0.149670786 | 0.5065063 | 0.99979869 | 1770.55883 | 1596.08083 |
| *DBP10* | -0.111498737 | 0.50696655 | 0.99979869 | 36319.3806 | 33618.1572 |
| *YCL001W-A* | -0.264378613 | 0.50719246 | 0.99979869 | 328.748389 | 273.701722 |
| *ERG27* | -0.111700961 | 0.50747168 | 0.99979869 | 31333.5259 | 28999.0562 |
| *ARG8* | -0.123951877 | 0.50757853 | 0.99979869 | 5250.21907 | 4817.97038 |
| *PHB1* | -0.111436565 | 0.50770244 | 0.99979869 | 33849.3778 | 31333.2091 |
| *OSW2* | -0.136275101 | 0.50807218 | 0.99979869 | 2719.73444 | 2474.5916 |
| *RDS1* | -0.126139846 | 0.5085032 | 0.99979869 | 4449.32166 | 4076.82303 |
| *ROX3* | -0.115187321 | 0.50870273 | 0.99979869 | 12455.3692 | 11499.5727 |
| *DST1* | -0.113308778 | 0.50874621 | 0.99979869 | 17619.743 | 16288.8403 |
| *MRPL11* | -0.119355794 | 0.50882739 | 0.99979869 | 7422.68988 | 6833.31715 |
| *ATG12* | -0.137766132 | 0.50945891 | 0.99979869 | 2513.90088 | 2284.94808 |
| *PPR1* | -0.126438658 | 0.50953489 | 0.99979869 | 4275.68008 | 3916.90741 |
| *RRI2* | -0.125363747 | 0.50970731 | 0.99979869 | 4549.79966 | 4171.13223 |
| *YIL102C* | -0.319110333 | 0.51152828 | 0.99979869 | 209.735619 | 168.116413 |
| *RSA1* | -0.120340723 | 0.51172461 | 0.99979869 | 6227.68461 | 5729.28436 |
| *YOR289W* | -0.126399464 | 0.51195944 | 0.99979869 | 4078.62615 | 3736.4898 |
| *THI4* | -0.189986627 | 0.51207367 | 0.99979869 | 741.391025 | 649.913452 |
| *YFL002W-A* | -0.446935812 | 0.51271701 | 0.99979869 | 103.404538 | 75.8574061 |
| *RPP1A* | -0.107779282 | 0.51278087 | 0.99979869 | 275816.971 | 255962.365 |
| *TUM1* | -0.113396046 | 0.51296668 | 0.99979869 | 13403.5693 | 12390.3847 |
| *YKL107W* | -0.229244877 | 0.51300325 | 0.99979869 | 445.810129 | 380.31213 |
| *RPB5* | -0.110716192 | 0.51346162 | 0.99979869 | 23502.0955 | 21765.95 |

| | | | | | |
|---|---|---|---|---|---|
| *GDA1* | -0.109248208 | 0.51349861 | 0.99979869 | 41050.6259 | 38056.8405 |
| *SAR1* | -0.109104294 | 0.51398393 | 0.99979869 | 40990.144 | 38004.5604 |
| *BEM1* | -0.114555044 | 0.51404784 | 0.99979869 | 10640.9122 | 9828.65958 |
| *RXT2* | -0.119896317 | 0.51482351 | 0.99979869 | 5935.03026 | 5461.73324 |
| *LYS9* | -0.109203598 | 0.51517328 | 0.99979869 | 33197.7342 | 30777.6049 |
| *YIL001W* | -0.126003517 | 0.51572119 | 0.99979869 | 3872.79259 | 3548.89648 |
| *RNH201* | -0.123758868 | 0.51576673 | 0.99979869 | 4415.17865 | 4052.22062 |
| *MSB4* | -0.116802495 | 0.51616518 | 0.99979869 | 7540.72713 | 6954.27895 |
| *SWM2* | -0.14459585 | 0.51621883 | 0.99979869 | 1830.06521 | 1655.53663 |
| *BUD21* | -0.11295341 | 0.51651471 | 0.99979869 | 12011.5101 | 11106.9594 |
| *SHE4* | -0.112274901 | 0.51668511 | 0.99979869 | 13308.9444 | 12312.4771 |
| *MND2* | -0.123313431 | 0.51729747 | 0.99979869 | 4393.71734 | 4033.76882 |
| *PHB2* | -0.108895837 | 0.51760966 | 0.99979869 | 28147.4956 | 26101.0983 |
| *DSL1* | -0.111877568 | 0.51767749 | 0.99979869 | 13515.7535 | 12507.2461 |
| *RSM18* | -0.115461136 | 0.5177425 | 0.99979869 | 8233.34243 | 7600.09201 |
| *MRPS16* | -0.115236038 | 0.51789797 | 0.99979869 | 8395.27784 | 7750.78172 |
| *SRD1* | -0.12140337 | 0.51813452 | 0.99979869 | 4886.35216 | 4491.98856 |
| *VAC14* | -0.113917767 | 0.51826262 | 0.99979869 | 9696.61419 | 8960.39981 |
| *RPL1A* | -0.106306285 | 0.51829053 | 0.99979869 | 177250.01 | 164658.751 |
| *RPN13* | -0.110091302 | 0.51878259 | 0.99979869 | 18077.2593 | 16749.1102 |
| *RPS13* | -0.106013338 | 0.51882927 | 0.99979869 | 251215.471 | 233417.339 |
| *YHR214C-E* | -0.802934586 | 0.51894572 | 0.99979869 | 32.1919787 | 18.4518015 |
| *POL5* | -0.108276067 | 0.51950396 | 0.99979869 | 28638.1794 | 26567.5188 |
| *RPL40A* | -0.105836327 | 0.51979804 | 0.99979869 | 187608.023 | 174337.746 |
| *tV(AAC)G2* | -0.312384475 | 0.51983705 | 0.99979869 | 208.760104 | 168.116413 |
| *MCX1* | -0.112744081 | 0.51986187 | 0.99979869 | 10608.7203 | 9811.23288 |
| *YLR122C* | -0.179278178 | 0.51986365 | 0.99979869 | 825.285272 | 728.846158 |
| *PHO13* | -0.108514564 | 0.52047351 | 0.99979869 | 23987.9017 | 22249.7973 |
| *RPL18A* | -0.105437791 | 0.520482 | 0.99979869 | 348140.641 | 323604.619 |
| *YHR173C* | -0.208953342 | 0.52048391 | 0.99979869 | 535.557464 | 463.345237 |
| *YDR132C* | -0.124962139 | 0.52088026 | 0.99979869 | 3715.73475 | 3407.43267 |
| *RRT15* | -0.55851821 | 0.52141496 | 0.99979869 | 63.4084429 | 43.0542034 |
| *CWC27* | -0.11636018 | 0.52212971 | 0.99979869 | 6542.77579 | 6035.78928 |
| *ORC6* | -0.120559673 | 0.52232451 | 0.99979869 | 4702.95543 | 4325.92235 |
| *PAC2* | -0.128899585 | 0.5223895 | 0.99979869 | 2978.24579 | 2723.69092 |
| *MIC19* | -0.116253101 | 0.52256809 | 0.99979869 | 6522.28999 | 6017.33748 |
| *MMS21* | -0.14349746 | 0.52288355 | 0.99979869 | 1747.14648 | 1581.72943 |
| *SNZ1* | -0.132491128 | 0.52292993 | 0.99979869 | 2537.31323 | 2314.67598 |
| *ESA1* | -0.11595536 | 0.52302287 | 0.99979869 | 6612.03732 | 6101.39569 |
| *YRF1-7* | -0.745601967 | 0.52337514 | 0.99979869 | 36.0940367 | 21.5271017 |
| *AMD1* | -0.107897298 | 0.52347617 | 0.99979869 | 22030.0441 | 20442.5458 |
| *LIN1* | -0.129037253 | 0.52363698 | 0.99979869 | 2901.18014 | 2652.95901 |
| *GCN4* | -0.104787456 | 0.52398109 | 0.99979869 | 131952.97 | 122708.58 |
| *SOV1* | -0.121115318 | 0.52436127 | 0.99979869 | 4339.08852 | 3989.68952 |
| *YHR127W* | -0.115785824 | 0.52466107 | 0.99979869 | 6401.32619 | 5907.65177 |

| | | | | | |
|---|---|---|---|---|---|
| *SPT14* | -0.115929948 | 0.52472536 | 0.99979869 | 6308.65231 | 5821.54336 |
| *ZUO1* | -0.104418033 | 0.52486695 | 0.99979869 | 161747.134 | 150453.939 |
| *CRP1* | -0.105251557 | 0.52540369 | 0.99979869 | 53290.4064 | 49541.0368 |
| *MED4* | -0.116155221 | 0.52541031 | 0.99979869 | 6069.65126 | 5600.12175 |
| *EMC3* | -0.10655144 | 0.52590203 | 0.99979869 | 26756.4119 | 24851.5013 |
| *ORC3* | -0.123675749 | 0.5260627 | 0.99979869 | 3615.25676 | 3318.24896 |
| *NOP56* | -0.10431444 | 0.52620512 | 0.99979869 | 101852.494 | 94747.9504 |
| *CEG1* | -0.11036672 | 0.52630828 | 0.99979869 | 11199.882 | 10375.0379 |
| *NIC96* | -0.106684998 | 0.52749964 | 0.99979869 | 22070.0402 | 20496.8761 |
| *CRT10* | -0.111072166 | 0.52765329 | 0.99979869 | 9575.65039 | 8866.09061 |
| *CCZ1* | -0.124060908 | 0.52805526 | 0.99979869 | 3416.2518 | 3134.75605 |
| *YDJ1* | -0.103663652 | 0.52805687 | 0.99979869 | 119124.954 | 110865.599 |
| *DAS2* | -0.12395433 | 0.52858724 | 0.99979869 | 3402.5946 | 3122.45485 |
| *RIA1* | -0.106264549 | 0.52868524 | 0.99979869 | 22506.0952 | 20907.9413 |
| *PTH2* | -0.109848774 | 0.5287355 | 0.99979869 | 10825.2845 | 10031.6294 |
| *STD1* | -0.109339459 | 0.52931848 | 0.99979869 | 11375.4747 | 10545.2045 |
| *IME2* | -0.124858202 | 0.52999237 | 0.99979869 | 3173.34869 | 2910.25913 |
| *THR4* | -0.103205502 | 0.53021618 | 0.99979869 | 94806.3528 | 88261.117 |
| *CDA1* | -0.343506778 | 0.53029185 | 0.99979869 | 159.984379 | 126.08731 |
| *YSH1* | -0.110380058 | 0.53033715 | 0.99979869 | 9403.95984 | 8711.3005 |
| *RPL4A* | -0.10234185 | 0.53099668 | 0.99979869 | 424978.993 | 395876.225 |
| *YGR201C* | -0.120257035 | 0.53109169 | 0.99979869 | 3981.0747 | 3662.68259 |
| *NUP133* | -0.106856771 | 0.53148456 | 0.99979869 | 15784.8002 | 14657.9061 |
| *PHO88* | -0.102707973 | 0.53196994 | 0.99979869 | 93714.7521 | 87274.9708 |
| *GCD14* | -0.109744425 | 0.5321621 | 0.99979869 | 9492.73166 | 8797.4089 |
| *ARL3* | -0.117847606 | 0.53323099 | 0.99979869 | 4424.9338 | 4077.84813 |
| *CWC24* | -0.119310318 | 0.53358239 | 0.99979869 | 4007.41359 | 3689.33519 |
| *snR44* | -0.11337425 | 0.53366068 | 0.99979869 | 6138.91279 | 5674.95405 |
| *APA1* | -0.103076577 | 0.5336954 | 0.99979869 | 45544.8213 | 42404.29 |
| *HPR1* | -0.109404995 | 0.53372717 | 0.99979869 | 9319.09008 | 8638.51839 |
| *MF(ALPHA)2* | -0.164768129 | 0.53382464 | 0.99979869 | 949.175614 | 846.732668 |
| *IRC22* | -0.103632402 | 0.53456248 | 0.99979869 | 30690.6619 | 28563.3887 |
| *AAP1* | -0.103686858 | 0.53508492 | 0.99979869 | 28310.4065 | 26347.1223 |
| *CDC31* | -0.109489259 | 0.53551074 | 0.99979869 | 8611.84206 | 7982.45434 |
| *ATS1* | -0.106169606 | 0.53586455 | 0.99979869 | 13931.3227 | 12942.9136 |
| *UBA4* | -0.10725625 | 0.53613894 | 0.99979869 | 11391.0829 | 10574.9324 |
| *IST3* | -0.124276849 | 0.53649236 | 0.99979869 | 2928.49455 | 2686.78731 |
| *CTF13* | -0.143232152 | 0.53687601 | 0.99979869 | 1502.29234 | 1360.30781 |
| *CPR3* | -0.103915605 | 0.53701859 | 0.99979869 | 21775.4348 | 20262.1282 |
| *BRE2* | -0.110122459 | 0.53716612 | 0.99979869 | 7561.21294 | 7005.53396 |
| *CAC2* | -0.106984521 | 0.53721206 | 0.99979869 | 11298.409 | 10490.8742 |
| *RVB2* | -0.103238592 | 0.53728834 | 0.99979869 | 26377.9122 | 24556.2725 |
| *PTC2* | -0.102736402 | 0.53733322 | 0.99979869 | 31859.3283 | 29669.4717 |
| *snR64* | -0.213773481 | 0.53787885 | 0.99979869 | 445.810129 | 384.412531 |
| *EAF7* | -0.105917042 | 0.53794628 | 0.99979869 | 13014.339 | 12093.1057 |

| | | | | | |
|---|---|---|---|---|---|
| *AGP1* | -0.107999931 | 0.53798025 | 0.99979869 | 9445.90696 | 8764.6057 |
| *SPT10* | -0.109088231 | 0.53813584 | 0.99979869 | 8207.00354 | 7609.31791 |
| *CHS7* | -0.106444312 | 0.53823142 | 0.99979869 | 11725.6844 | 10891.6884 |
| *CCH1* | -0.107984741 | 0.53863307 | 0.99979869 | 9221.53863 | 8556.51038 |
| *TIR1* | -0.102942046 | 0.53892456 | 0.99979869 | 24801.4808 | 23093.4546 |
| *NUP53* | -0.105567048 | 0.53895194 | 0.99979869 | 13131.4008 | 12204.8416 |
| *MAK32* | -0.115446745 | 0.53911561 | 0.99979869 | 4566.3834 | 4215.21154 |
| *SHE2* | -0.110145961 | 0.53959505 | 0.99979869 | 6974.92872 | 6462.23092 |
| *GRX3* | -0.103108709 | 0.54028861 | 0.99979869 | 20860.4022 | 19421.5461 |
| *SPE4* | -0.110007472 | 0.54036152 | 0.99979869 | 6905.66719 | 6398.67471 |
| *YKG9* | -0.110448406 | 0.54038414 | 0.99979869 | 6610.08629 | 6122.92279 |
| *IDP3* | -0.131753801 | 0.54046955 | 0.99979869 | 2048.58046 | 1869.78255 |
| *YIP3* | -0.103805749 | 0.54149197 | 0.99979869 | 16059.8953 | 14944.9341 |
| *SEC65* | -0.104237995 | 0.54155644 | 0.99979869 | 14602.4766 | 13584.6263 |
| *GAL2* | -0.144004147 | 0.54157776 | 0.99979869 | 1398.8878 | 1265.9986 |
| *YDR034C-D* | -0.120923719 | 0.54185122 | 0.99979869 | 3147.98531 | 2894.88263 |
| *UBC11* | -0.21036335 | 0.54200608 | 0.99979869 | 450.687702 | 389.538031 |
| *YCH1* | -0.112637773 | 0.54209419 | 0.99979869 | 5225.83121 | 4833.34689 |
| *KTI11* | -0.119890507 | 0.54248965 | 0.99979869 | 3281.6308 | 3019.94484 |
| *TIF6* | -0.100494831 | 0.54369486 | 0.99979869 | 38654.7623 | 36053.795 |
| *NMA2* | -0.109028582 | 0.54430102 | 0.99979869 | 6741.78075 | 6251.0603 |
| *ARP6* | -0.109388666 | 0.54432547 | 0.99979869 | 6505.70624 | 6030.66378 |
| *CKA2* | -0.101881847 | 0.54445155 | 0.99979869 | 20964.7823 | 19535.3323 |
| *SEC28* | -0.10111852 | 0.54549313 | 0.99979869 | 24183.0046 | 22546.0512 |
| *SRP1* | -0.099830181 | 0.54552003 | 0.99979869 | 41827.1355 | 39030.6856 |
| *GCD6* | -0.100225548 | 0.5456326 | 0.99979869 | 33662.0791 | 31402.9159 |
| *MTR4* | -0.099552918 | 0.54641142 | 0.99979869 | 42414.3952 | 39586.2899 |
| *CCT8* | -0.099018438 | 0.54646584 | 0.99979869 | 60423.3685 | 56415.3579 |
| *KAR3* | -0.113223883 | 0.5469265 | 0.99979869 | 4471.7585 | 4134.22863 |
| *GCR1* | -0.100355481 | 0.54717345 | 0.99979869 | 26672.5176 | 24880.2041 |
| *PAU9* | -0.665432101 | 0.54743182 | 0.99979869 | 39.0205802 | 24.602402 |
| *POL1* | -0.104675166 | 0.54785276 | 0.99979869 | 9910.25187 | 9216.67484 |
| *LSM1* | -0.10101082 | 0.54817874 | 0.99979869 | 19792.2138 | 18453.8517 |
| *snR55* | -0.254239859 | 0.54836788 | 0.99979869 | 275.095091 | 230.647518 |
| *AMD2* | -0.112540389 | 0.54865433 | 0.99979869 | 4511.75459 | 4173.18243 |
| *SIF2* | -0.10738309 | 0.54866546 | 0.99979869 | 6950.54086 | 6451.97991 |
| *IES4* | -0.10922023 | 0.54887895 | 0.99979869 | 5807.23785 | 5383.82563 |
| *VPS20* | -0.117894731 | 0.54908586 | 0.99979869 | 3242.61022 | 2988.16674 |
| *YPR195C* | -0.19048518 | 0.54913507 | 0.99979869 | 552.14121 | 483.847239 |
| *UTP6* | -0.105914193 | 0.54927293 | 0.99979869 | 7999.21895 | 7433.00069 |
| *CHO2* | -0.098927866 | 0.54933253 | 0.99979869 | 37991.4124 | 35473.5883 |
| *PDP3* | -0.108898977 | 0.54943551 | 0.99979869 | 5883.32799 | 5455.58264 |
| *HAM1* | -0.101438544 | 0.54991862 | 0.99979869 | 15739.9266 | 14671.2324 |
| *BLI1* | -0.11707271 | 0.55008593 | 0.99979869 | 3327.47998 | 3068.12454 |
| *DPH2* | -0.099129714 | 0.55010668 | 0.99979869 | 31146.2272 | 29077.9889 |

| | | | | | |
|---|---|---|---|---|---|
| *STP2* | -0.104240032 | 0.55048463 | 0.99979869 | 9440.05388 | 8782.0324 |
| *YVH1* | -0.100997095 | 0.55109062 | 0.99979869 | 16106.72 | 15017.7162 |
| *APN1* | -0.102611912 | 0.55141293 | 0.99979869 | 11573.5041 | 10778.9274 |
| *RAD28* | -0.120311308 | 0.55187306 | 0.99979869 | 2749.97539 | 2529.947 |
| *VMA3* | -0.096817594 | 0.55209325 | 0.99979869 | 124487.357 | 116407.29 |
| *YBL029W* | -0.112733699 | 0.55405427 | 0.99979869 | 3970.34404 | 3671.90849 |
| *CDC33* | -0.096592413 | 0.55406013 | 0.99979869 | 80551.1593 | 75334.605 |
| *snR72* | -0.346970497 | 0.55414114 | 0.99979869 | 135.596516 | 106.610409 |
| *MSS1* | -0.117758171 | 0.55420661 | 0.99979869 | 2990.92748 | 2756.49412 |
| *MSP1* | -0.10472414 | 0.55568404 | 0.99979869 | 7354.40386 | 6839.46775 |
| *MRH4* | -0.112928519 | 0.55569249 | 0.99979869 | 3795.72694 | 3509.94268 |
| *GIS2* | -0.096122337 | 0.55572677 | 0.99979869 | 81359.8608 | 76115.7313 |
| *MES1* | -0.09622331 | 0.55584434 | 0.99979869 | 70738.4589 | 66174.3107 |
| *TAL1* | -0.095758589 | 0.55612843 | 0.99979869 | 113656.22 | 106357.209 |
| *NOG1* | -0.096833916 | 0.55621899 | 0.99979869 | 41553.9914 | 38856.4186 |
| *STO1* | -0.097891261 | 0.55644703 | 0.99979869 | 25127.3026 | 23478.8923 |
| *PRP31* | -0.109546351 | 0.55669724 | 0.99979869 | 4656.13074 | 4315.67134 |
| *YEA6* | -0.103363067 | 0.55711898 | 0.99979869 | 8194.32185 | 7627.76971 |
| *IBD2* | -0.113636085 | 0.55714364 | 0.99979869 | 3539.16663 | 3271.09436 |
| *STE7* | -0.108514236 | 0.55721124 | 0.99979869 | 4972.19744 | 4611.92527 |
| *YBR230W-A* | -0.116794241 | 0.55742635 | 0.99979869 | 2973.36821 | 2742.14272 |
| *SWT1* | -0.105811413 | 0.55758063 | 0.99979869 | 6207.1988 | 5768.23816 |
| *RPC53* | -0.098977589 | 0.55766549 | 0.99979869 | 16618.8651 | 15516.9399 |
| *tV(AAC)J* | -1.015924441 | 0.55777278 | 0.99979869 | 16.5837466 | 8.20080065 |
| *POM152* | -0.097617868 | 0.55799674 | 0.99979869 | 23754.7537 | 22200.5925 |
| *HRB1* | -0.097272081 | 0.55814736 | 0.99979869 | 26459.8555 | 24734.6399 |
| *RRT14* | -0.101145068 | 0.55864427 | 0.99979869 | 10438.0052 | 9731.27508 |
| *RHB1* | -0.111084251 | 0.55938313 | 0.99979869 | 3954.73581 | 3661.65749 |
| *MRPS5* | -0.103393744 | 0.55950747 | 0.99979869 | 7509.51067 | 6990.15746 |
| *BRR1* | -0.104510426 | 0.55994284 | 0.99979869 | 6573.01674 | 6113.69689 |
| *YLR225C* | -0.100384298 | 0.55999914 | 0.99979869 | 11063.31 | 10319.6825 |
| *YJR079W* | -0.450889387 | 0.56004815 | 0.99979869 | 77.065646 | 56.3805045 |
| *SUP45* | -0.094920462 | 0.56024051 | 0.99979869 | 77822.6452 | 72867.1891 |
| *CDC45* | -0.117701693 | 0.5605409 | 0.99979869 | 2706.07724 | 2494.0685 |
| *HUB1* | -0.108645973 | 0.5611115 | 0.99979869 | 4495.17084 | 4169.08203 |
| *ACL4* | -0.097246566 | 0.56144174 | 0.99979869 | 19748.3157 | 18461.0274 |
| *SAM37* | -0.107043643 | 0.56156581 | 0.99979869 | 5019.02213 | 4660.10497 |
| *IWR1* | -0.10129224 | 0.56237821 | 0.99979869 | 8753.29166 | 8159.79665 |
| *HSP10* | -0.095115274 | 0.56244064 | 0.99979869 | 41466.1951 | 38820.5401 |
| *PRP28* | -0.10846897 | 0.56246884 | 0.99979869 | 4416.15417 | 4096.29993 |
| *DIF1* | -0.100801759 | 0.56274533 | 0.99979869 | 9218.61208 | 8596.48929 |
| *SPC19* | -0.109155039 | 0.56293776 | 0.99979869 | 4168.37348 | 3864.62731 |
| *POL12* | -0.108218331 | 0.56304224 | 0.99979869 | 4438.591 | 4117.82703 |
| *MNT2* | -0.09859914 | 0.56311107 | 0.99979869 | 12944.102 | 12089.0053 |
| *PAU15* | -0.51343029 | 0.56343665 | 0.99979869 | 58.5308704 | 41.0040033 |

| | | | | | |
|---|---|---|---|---|---|
| *UPS2* | -0.109298433 | 0.56405262 | 0.99979869 | 4032.77697 | 3738.54 |
| *POM34* | -0.098791763 | 0.56465995 | 0.99979869 | 11537.4101 | 10773.8019 |
| *SCJ1* | -0.09527908 | 0.56595656 | 0.99979869 | 24756.6071 | 23174.4375 |
| *PEX34* | -0.11163058 | 0.56646718 | 0.99979869 | 3338.21064 | 3089.65165 |
| *REV3* | -0.103413331 | 0.56667301 | 0.99979869 | 6005.2673 | 5589.87075 |
| *MNN10* | -0.095071843 | 0.56711255 | 0.99979869 | 23792.7988 | 22275.4248 |
| *VPS51* | -0.103682938 | 0.56723071 | 0.99979869 | 5766.26625 | 5366.39893 |
| *CDC42* | -0.094497216 | 0.56754012 | 0.99979869 | 28258.7042 | 26467.059 |
| *MCM10* | -0.10947385 | 0.56778046 | 0.99979869 | 3698.17549 | 3427.93467 |
| *SGE1* | -0.097407932 | 0.56855141 | 0.99979869 | 12111.9881 | 11321.2053 |
| *TGS1* | -0.10173375 | 0.56913227 | 0.99979869 | 6594.47806 | 6145.47499 |
| *SDS3* | -0.10456507 | 0.56979632 | 0.99979869 | 5000.48736 | 4650.87907 |
| *IBA57* | -0.103745283 | 0.57016258 | 0.99979869 | 5303.87237 | 4935.85689 |
| *PET18* | -0.156390639 | 0.57039052 | 0.99979869 | 807.726011 | 724.745758 |
| *PIF1* | -0.100075744 | 0.57063357 | 0.99979869 | 7541.70265 | 7036.28696 |
| *KRE2* | -0.092360479 | 0.57114567 | 0.99979869 | 53642.5672 | 50316.0124 |
| *AIR1* | -0.100214319 | 0.57129393 | 0.99979869 | 7257.82793 | 6770.78604 |
| *LCL3* | -0.109095625 | 0.57135126 | 0.99979869 | 3531.36251 | 3274.16966 |
| *BOL2* | -0.095695095 | 0.57177805 | 0.99979869 | 14189.834 | 13279.1465 |
| *TDA5* | -0.102323632 | 0.57192744 | 0.99979869 | 5739.92735 | 5346.92203 |
| *ALB1* | -0.09642064 | 0.57204894 | 0.99979869 | 12108.086 | 11325.3057 |
| *NVJ3* | -0.113821057 | 0.5723731 | 0.99979869 | 2694.37107 | 2489.9681 |
| *ISA2* | -0.103101277 | 0.57247457 | 0.99979869 | 5279.48451 | 4915.35489 |
| *YJR124C* | -0.096512515 | 0.5729315 | 0.99979869 | 11392.0584 | 10654.8903 |
| *RAD53* | -0.115433154 | 0.57311557 | 0.99979869 | 2471.95376 | 2281.87278 |
| *ILV1* | -0.091803566 | 0.57345351 | 0.99979869 | 51044.772 | 47897.8013 |
| *DGR1* | -0.355890197 | 0.57373159 | 0.99979869 | 114.135197 | 89.1837071 |
| *GCN5* | -0.097409441 | 0.57389161 | 0.99979869 | 9454.68659 | 8837.38781 |
| *THI3* | -0.097941085 | 0.57463851 | 0.99979869 | 8520.1437 | 7960.92724 |
| *FRE2* | -0.124021004 | 0.57483074 | 0.99979869 | 1731.53825 | 1588.90513 |
| *tK(CUU)D1* | -1.027996478 | 0.57494313 | 0.99979869 | 14.6327176 | 7.17570057 |
| *SRP101* | -0.092921566 | 0.57497953 | 0.99979869 | 23705.978 | 22227.2451 |
| *HUT1* | -0.096172236 | 0.57565599 | 0.99979869 | 10596.0386 | 9912.71779 |
| *RET1* | -0.092013565 | 0.57604738 | 0.99979869 | 30311.1867 | 28438.3265 |
| *SQS1* | -0.097050529 | 0.57629044 | 0.99979869 | 9022.53367 | 8435.54857 |
| *PBI1* | -0.134658703 | 0.57650121 | 0.99979869 | 1237.92791 | 1127.61009 |
| *DIM1* | -0.092841041 | 0.57650647 | 0.99979869 | 21180.371 | 19860.289 |
| *FZO1* | -0.096838118 | 0.57700631 | 0.99979869 | 9033.26433 | 8446.82467 |
| *ACO2* | -0.09204987 | 0.57735891 | 0.99979869 | 25754.5585 | 24162.634 |
| *YUR1* | -0.099277961 | 0.57755825 | 0.99979869 | 6578.86983 | 6141.37459 |
| *TIF5* | -0.090233262 | 0.57831716 | 0.99979869 | 63919.6125 | 60044.2122 |
| *CIN4* | -0.106828557 | 0.5787618 | 0.99979869 | 3514.77877 | 3263.91866 |
| *BAS1* | -0.097152096 | 0.57886326 | 0.99979869 | 8062.62739 | 7537.5609 |
| *MET10* | -0.09483006 | 0.57946966 | 0.99979869 | 11088.6734 | 10383.2387 |
| *YPR109W* | -0.098804059 | 0.57979584 | 0.99979869 | 6455.955 | 6028.61358 |

| | | | | | |
|---|---|---|---|---|---|
| *YGR204C-A* | -0.16344053 | 0.58002568 | 0.99979869 | 655.545748 | 585.332147 |
| *YEL023C* | -0.105529544 | 0.58018295 | 0.99979869 | 3707.93064 | 3446.38647 |
| *YNL162W-A* | -0.142941563 | 0.58033286 | 0.99979869 | 974.538992 | 882.61117 |
| *SPC1* | -0.102318443 | 0.5808624 | 0.99979869 | 4580.04061 | 4266.46654 |
| *RDI1* | -0.093605297 | 0.58088549 | 0.99979869 | 12943.1265 | 12130.0093 |
| *YMR027W* | -0.091228101 | 0.58090214 | 0.99979869 | 24460.0507 | 22961.2167 |
| *MGE1* | -0.090412174 | 0.58127442 | 0.99979869 | 33249.4364 | 31229.674 |
| *GEM1* | -0.099773249 | 0.58128775 | 0.99979869 | 5619.93907 | 5244.41202 |
| *YMD8* | -0.094192342 | 0.58131917 | 0.99979869 | 11326.6989 | 10610.8109 |
| *SWC7* | -0.169917557 | 0.58149755 | 0.99979869 | 582.38216 | 517.675541 |
| *TED1* | -0.090832362 | 0.58169176 | 0.99979869 | 26239.3892 | 24638.2805 |
| *YML079W* | -0.094450571 | 0.5821432 | 0.99979869 | 10421.4215 | 9761.00298 |
| *tT(AGU)I1* | -0.764965948 | 0.58217775 | 0.99979869 | 24.3878627 | 14.3514011 |
| *YOR238W* | -0.103568151 | 0.58221383 | 0.99979869 | 4054.23829 | 3773.3934 |
| *TVP38* | -0.094280874 | 0.58258059 | 0.99979869 | 10503.3647 | 9838.91059 |
| *SSB1* | -0.088084863 | 0.58294577 | 0.99979869 | 520217.498 | 489405.331 |
| *PEX10* | -0.099950722 | 0.58356722 | 0.99979869 | 5205.3454 | 4856.92419 |
| *YRF1-2* | -0.183147803 | 0.58382104 | 0.99979869 | 470.197992 | 414.140433 |
| *YNR063W* | -0.115341285 | 0.58398573 | 0.99979869 | 2119.79302 | 1956.91606 |
| *YCR023C* | -0.092232017 | 0.58436201 | 0.99979869 | 14127.4011 | 13252.4939 |
| *THR1* | -0.088616333 | 0.58442457 | 0.99979869 | 63916.686 | 60108.7935 |
| *ILV6* | -0.088212763 | 0.58451251 | 0.99979869 | 100627.248 | 94658.7667 |
| *CAB2* | -0.095561732 | 0.58465093 | 0.99979869 | 8001.16998 | 7488.3561 |
| *OGG1* | -0.103473701 | 0.58539652 | 0.99979869 | 3817.18826 | 3552.99688 |
| *RSM23* | -0.094183997 | 0.58547699 | 0.99979869 | 9385.42506 | 8792.2834 |
| *YNR065C* | -0.099051534 | 0.58547772 | 0.99979869 | 5362.40324 | 5006.5888 |
| *VPS9* | -0.097357441 | 0.58574261 | 0.99979869 | 6269.63173 | 5860.49717 |
| *AVL9* | -0.089840925 | 0.58586842 | 0.99979869 | 24953.6611 | 23447.1142 |
| *AFG2* | -0.090979072 | 0.58619375 | 0.99979869 | 16940.7849 | 15905.4529 |
| *YDR182W-A* | -0.252008729 | 0.58629324 | 0.99979869 | 218.515249 | 183.492915 |
| *PTI1* | -0.096494283 | 0.58722758 | 0.99979869 | 6566.18814 | 6141.37459 |
| *IMG1* | -0.091222509 | 0.58733241 | 0.99979869 | 14777.0937 | 13871.6543 |
| *YNG2* | -0.09939056 | 0.5874307 | 0.99979869 | 4950.73612 | 4621.15117 |
| *DCW1* | -0.088746221 | 0.58803254 | 0.99979869 | 31145.2516 | 29287.1093 |
| *RHO1* | -0.087759666 | 0.58805899 | 0.99979869 | 58371.8615 | 54926.9126 |
| *CBC2* | -0.090797832 | 0.58813788 | 0.99979869 | 15519.4603 | 14572.8228 |
| *SAD1* | -0.094176762 | 0.58835406 | 0.99979869 | 8369.91446 | 7840.99053 |
| *IMG2* | -0.098230364 | 0.58847306 | 0.99979869 | 5334.11332 | 4983.0115 |
| *PEX19* | -0.091599037 | 0.5886234 | 0.99979869 | 12628.0353 | 11851.182 |
| *TIM50* | -0.088137204 | 0.58869122 | 0.99979869 | 39321.0387 | 36990.7365 |
| *HPM1* | -0.09253156 | 0.58886485 | 0.99979869 | 10509.2178 | 9856.33729 |
| *DUT1* | -0.091033411 | 0.58888525 | 0.99979869 | 14015.2169 | 13158.1846 |
| *MEF1* | -0.090697706 | 0.58959548 | 0.99979869 | 14473.7087 | 13591.802 |
| *SEC17* | -0.090189033 | 0.58971191 | 0.99979869 | 16246.2186 | 15261.69 |
| *GAS2* | -0.09805916 | 0.58977867 | 0.99979869 | 5233.63533 | 4889.72739 |

| | | | | | |
|---|---|---|---|---|---|
| *TRS31* | -0.094809371 | 0.58988783 | 0.99979869 | 7300.75056 | 6836.39245 |
| *CAB1* | -0.090830151 | 0.59009067 | 0.99979869 | 13637.6928 | 12805.5502 |
| *GYP8* | -0.094582642 | 0.59040222 | 0.99979869 | 7372.93864 | 6905.07415 |
| *BCH2* | -0.094425133 | 0.59055095 | 0.99979869 | 7478.2942 | 7004.50886 |
| *SRB6* | -0.097725513 | 0.59078357 | 0.99979869 | 5252.1701 | 4908.17919 |
| *FYV4* | -0.101817574 | 0.59092027 | 0.99979869 | 3808.40863 | 3548.89648 |
| *CUS2* | -0.095845848 | 0.59170502 | 0.99979869 | 6135.98624 | 5741.58556 |
| *NPA3* | -0.08753859 | 0.59221765 | 0.99979869 | 32770.4588 | 30841.1611 |
| *URE2* | -0.089424959 | 0.59227738 | 0.99979869 | 16559.3587 | 15564.0945 |
| *YHR140W* | -0.131852277 | 0.59261872 | 0.99979869 | 1127.69477 | 1029.20048 |
| *URB2* | -0.090986474 | 0.59311584 | 0.99979869 | 11226.2209 | 10540.079 |
| *TRM9* | -0.090942462 | 0.59335394 | 0.99979869 | 11183.2983 | 10500.1001 |
| *tG(GCC)O1* | -0.722014128 | 0.59365295 | 0.99979869 | 25.3633772 | 15.3765012 |
| *SUE1* | -0.197756471 | 0.59396355 | 0.99979869 | 360.940367 | 314.705725 |
| *DBF20* | -0.100134161 | 0.59476273 | 0.99979869 | 3947.90721 | 3683.18459 |
| *DUS3* | -0.08767549 | 0.59570599 | 0.99979869 | 21222.3181 | 19970.9998 |
| *CMD1* | -0.086472882 | 0.59621256 | 0.99979869 | 33092.3786 | 31167.1429 |
| *RPL1B* | -0.084493854 | 0.59725397 | 0.99979869 | 281868.088 | 265834.079 |
| *HRQ1* | -0.090552308 | 0.59775429 | 0.99979869 | 9730.7572 | 9138.76723 |
| *RPS1A* | -0.084204946 | 0.59823107 | 0.99979869 | 340374.57 | 321076.722 |
| *IMP2* | -0.095173321 | 0.59854242 | 0.99979869 | 5399.47279 | 5054.7685 |
| *snR47* | -0.166833371 | 0.59900053 | 0.99979869 | 531.655406 | 473.596238 |
| *YNL284C-B* | -0.095306693 | 0.59923489 | 0.99979869 | 5234.61084 | 4899.97839 |
| *RTT10* | -0.087282996 | 0.59987969 | 0.99979869 | 17079.308 | 16076.6446 |
| *PFA5* | -0.097564446 | 0.60018632 | 0.99979869 | 4233.73296 | 3956.88632 |
| *EMC6* | -0.089205158 | 0.60059547 | 0.99979869 | 10723.831 | 10080.8342 |
| *CMC1* | -0.092230435 | 0.60075559 | 0.99979869 | 6891.03447 | 6464.28112 |
| *EKI1* | -0.093710325 | 0.60194429 | 0.99979869 | 5660.91068 | 5304.89292 |
| *BUL3* | -0.114429333 | 0.6027941 | 0.99979869 | 1710.07693 | 1579.67923 |
| *RTC5* | -0.087758299 | 0.60287913 | 0.99979869 | 12589.0147 | 11846.0565 |
| *ARK1* | -0.088199932 | 0.60294346 | 0.99979869 | 11485.7078 | 10804.5549 |
| *RRP46* | -0.089205038 | 0.60323197 | 0.99979869 | 9502.4868 | 8932.72211 |
| *MRX6* | -0.092254174 | 0.60338328 | 0.99979869 | 6317.43194 | 5926.10357 |
| *BUB3* | -0.094704261 | 0.60384305 | 0.99979869 | 4902.93591 | 4591.42327 |
| *ASP1* | -0.084278914 | 0.6043857 | 0.99979869 | 34438.5886 | 32484.3965 |
| *YBR196C-A* | -0.189234105 | 0.60439089 | 0.99979869 | 371.671027 | 325.981826 |
| *FPS1* | -0.084653698 | 0.60448477 | 0.99979869 | 28170.9079 | 26565.4686 |
| *EFR3* | -0.085708644 | 0.6046328 | 0.99979869 | 18762.0705 | 17679.9011 |
| *YOR142W-B* | -0.154237187 | 0.60474569 | 0.99979869 | 626.280313 | 562.779945 |
| *YNL095C* | -0.090367307 | 0.60493057 | 0.99979869 | 7517.31478 | 7060.88936 |
| *PDB1* | -0.082989196 | 0.60510039 | 0.99979869 | 88400.149 | 83458.5232 |
| *PET100* | -0.097259216 | 0.60530675 | 0.99979869 | 3869.86605 | 3617.57819 |
| *NGL1* | -0.092595535 | 0.60531305 | 0.99979869 | 5751.63353 | 5394.07663 |
| *FRE5* | -0.088944411 | 0.60617715 | 0.99979869 | 8738.65895 | 8216.17716 |
| *YJR027W* | -0.088270537 | 0.60646392 | 0.99979869 | 9592.23414 | 9022.93092 |

| | | | | | |
|---|---|---|---|---|---|
| *SFA1* | -0.084568643 | 0.60662998 | 0.99979869 | 23172.3716 | 21853.0835 |
| *CIN10* | -0.089242858 | 0.60711377 | 0.99979869 | 8066.52945 | 7582.66531 |
| *KSS1* | -0.093181986 | 0.60722177 | 0.99979869 | 5153.64314 | 4831.29669 |
| *SPC98* | -0.096878618 | 0.60859001 | 0.99979869 | 3709.88167 | 3468.93868 |
| *RPL6A* | -0.081753764 | 0.60863595 | 0.99979869 | 170930.627 | 161513.744 |
| *MRPL7* | -0.087688762 | 0.609008 | 0.99979869 | 9429.32322 | 8873.26631 |
| *YOL118C* | -0.131042047 | 0.60906309 | 0.99979869 | 975.514506 | 890.811971 |
| *GET1* | -0.085578349 | 0.60929482 | 0.99979869 | 13992.7801 | 13186.8875 |
| *YTM1* | -0.083748491 | 0.60965197 | 0.99979869 | 23517.7037 | 22191.3666 |
| *YOR316C-A* | -0.2096515 | 0.60976401 | 0.99979869 | 280.948178 | 242.948719 |
| *YJL107C* | -0.10255551 | 0.60979071 | 0.99979869 | 2583.16241 | 2405.90989 |
| *UTR1* | -0.084856073 | 0.61029623 | 0.99979869 | 15687.2488 | 14791.1691 |
| *YBL039W-A* | -0.118734567 | 0.61040271 | 0.99979869 | 1350.11208 | 1243.4464 |
| *THI6* | -0.088098096 | 0.61055319 | 0.99979869 | 8307.48153 | 7815.36302 |
| *AIM36* | -0.092098579 | 0.61059722 | 0.99979869 | 5213.14952 | 4890.75249 |
| *IRA2* | -0.082748022 | 0.61068374 | 0.99979869 | 32886.545 | 31053.3568 |
| *RSM22* | -0.089723713 | 0.61101007 | 0.99979869 | 6613.98835 | 6215.1818 |
| *YAL064W-B* | -0.163918556 | 0.6110747 | 0.99979869 | 505.316514 | 451.044036 |
| *MET12* | -0.08393181 | 0.61117401 | 0.99979869 | 19194.2234 | 18109.418 |
| *SHH3* | -0.118020448 | 0.61154173 | 0.99979869 | 1362.79377 | 1255.7476 |
| *TIM23* | -0.082629017 | 0.61173684 | 0.99979869 | 30417.5178 | 28724.3294 |
| *NIT1* | -0.098738454 | 0.61201201 | 0.99979869 | 3075.79724 | 2872.33043 |
| *RPS4A* | -0.080607157 | 0.61232695 | 0.99979869 | 439326.86 | 415453.586 |
| *RMP1* | -0.099745789 | 0.61264029 | 0.99979869 | 2867.03713 | 2675.51121 |
| *YPL062W* | -0.270701546 | 0.61267779 | 0.99979869 | 157.057835 | 130.18771 |
| *YDR170W-A* | -0.094644035 | 0.61286821 | 0.99979869 | 3981.0747 | 3728.289 |
| *RSM24* | -0.087794567 | 0.61318658 | 0.99979869 | 7837.28354 | 7374.56999 |
| *CCT7* | -0.08143048 | 0.61325437 | 0.99979869 | 51317.9161 | 48501.5853 |
| *ELA1* | -0.091140277 | 0.6133956 | 0.99979869 | 5297.04377 | 4972.7605 |
| *YDL057W* | -0.099116046 | 0.61347048 | 0.99979869 | 2929.47006 | 2734.96702 |
| *PAU20* | -0.846000464 | 0.61348354 | 0.99979869 | 16.5837466 | 9.22590074 |
| *FOB1* | -0.086919841 | 0.61375138 | 0.99979869 | 8703.54042 | 8194.65005 |
| *NUT2* | -0.107223471 | 0.61376115 | 0.99979869 | 1958.83313 | 1818.52755 |
| *TRM13* | -0.086373685 | 0.61429621 | 0.99979869 | 9268.36332 | 8729.7523 |
| *KEX1* | -0.082513033 | 0.61431612 | 0.99979869 | 23860.1093 | 22533.75 |
| *MGM1* | -0.085039936 | 0.61446375 | 0.99979869 | 11695.4434 | 11025.9765 |
| *SSY5* | -0.086881808 | 0.61458621 | 0.99979869 | 8461.61283 | 7967.07784 |
| *IFA38* | -0.081642166 | 0.61473893 | 0.99979869 | 34263.9715 | 32378.8112 |
| *YHR045W* | -0.084454044 | 0.61551982 | 0.99979869 | 12470.0019 | 11760.9732 |
| *BDS1* | -0.088778823 | 0.61563211 | 0.99979869 | 6363.28112 | 5983.50918 |
| *SEC4* | -0.081866538 | 0.615652 | 0.99979869 | 27175.8831 | 25676.7068 |
| *IFM1* | -0.094901054 | 0.61577456 | 0.99979869 | 3670.86109 | 3437.16057 |
| *MET31* | -0.091339078 | 0.61622374 | 0.99979869 | 4824.89475 | 4528.89216 |
| *tQ(UUG)D3* | -0.928461073 | 0.61633203 | 0.99979869 | 13.6572031 | 7.17570057 |
| *YCL002C* | -0.087879616 | 0.61643786 | 0.99979869 | 6900.78962 | 6492.98392 |

| | | | | | |
|---|---|---|---|---|---|
| *CKB2* | -0.083743771 | 0.61682109 | 0.99979869 | 13546.9699 | 12782.998 |
| *SNF4* | -0.083206602 | 0.61684122 | 0.99979869 | 15461.9049 | 14595.375 |
| *NOP14* | -0.081107644 | 0.61749861 | 0.99979869 | 31316.9422 | 29604.8904 |
| *CBR1* | -0.08009567 | 0.61772756 | 0.99979869 | 62511.9451 | 59135.9735 |
| *CWC25* | -0.090061398 | 0.6177661 | 0.99979869 | 5219.97812 | 4904.07879 |
| *ISC10* | -0.097449211 | 0.6180852 | 0.99979869 | 2974.34373 | 2780.07142 |
| *YFL041W-A* | -0.14822258 | 0.61849677 | 0.99979869 | 621.40274 | 560.729745 |
| *PRP5* | -0.085120818 | 0.61907437 | 0.99979869 | 9234.22031 | 8705.14989 |
| *CDS1* | -0.08283438 | 0.61922147 | 0.99979869 | 14516.6314 | 13706.6132 |
| *YLR162W* | -0.423232738 | 0.61950325 | 0.99979869 | 60.4818994 | 45.1044036 |
| *IML3* | -0.093495171 | 0.61983737 | 0.99979869 | 3727.44093 | 3493.54108 |
| *ERV46* | -0.079370369 | 0.6198903 | 0.99979869 | 77521.2113 | 73371.5384 |
| *ASA1* | -0.090874944 | 0.62040664 | 0.99979869 | 4526.38731 | 4250.06494 |
| *YJR154W* | -0.129098161 | 0.62101779 | 0.99979869 | 909.17952 | 831.356166 |
| *YLR278C* | -0.081219132 | 0.62129035 | 0.99979869 | 19924.8838 | 18834.1638 |
| *KIN28* | -0.087591119 | 0.62164364 | 0.99979869 | 6034.53273 | 5679.05445 |
| *YMR253C* | -0.086946101 | 0.62170407 | 0.99979869 | 6489.12249 | 6109.59649 |
| *TGA1* | -0.735819064 | 0.62174223 | 0.99979869 | 20.4858046 | 12.301201 |
| *MIA40* | -0.080131496 | 0.62176625 | 0.99979869 | 29728.8046 | 28122.5956 |
| *YMR226C* | -0.079542598 | 0.62185215 | 0.99979869 | 41652.5184 | 39418.1734 |
| *YCL021W-A* | -0.108261615 | 0.62201903 | 0.99979869 | 1681.78701 | 1560.20232 |
| *BSC1* | -0.085230517 | 0.62225809 | 0.99979869 | 7968.00249 | 7510.9083 |
| *UTP20* | -0.079856288 | 0.62237932 | 0.99979869 | 31670.0784 | 29964.7005 |
| *KRE33* | -0.079674261 | 0.6225119 | 0.99979869 | 34431.76 | 32581.781 |
| *TPA1* | -0.079655734 | 0.62251401 | 0.99979869 | 34798.5535 | 32929.2899 |
| *YML003W* | -0.105024011 | 0.62303267 | 0.99979869 | 1888.59608 | 1755.99644 |
| *SPO75* | -0.092589629 | 0.62303958 | 0.99979869 | 3719.63681 | 3488.41558 |
| *BRE4* | -0.082394169 | 0.6234216 | 0.99979869 | 12464.1488 | 11772.2493 |
| *RPL37A* | -0.078114218 | 0.62425638 | 0.99979869 | 100405.806 | 95113.9111 |
| *EXO5* | -0.087111659 | 0.62464812 | 0.99979869 | 5808.21337 | 5467.88384 |
| *GRS1* | -0.077852415 | 0.62478142 | 0.99979869 | 129472.236 | 122670.651 |
| *YPL150W* | -0.083485252 | 0.62495626 | 0.99979869 | 9355.18411 | 8829.187 |
| *PRP2* | -0.083930663 | 0.62521913 | 0.99979869 | 8604.03794 | 8117.76755 |
| *GUF1* | -0.094782697 | 0.62531713 | 0.99979869 | 3064.09106 | 2869.25513 |
| *YMR317W* | -0.083534088 | 0.62578297 | 0.99979869 | 8950.34559 | 8446.82467 |
| *RPB4* | -0.079511374 | 0.62613109 | 0.99979869 | 23887.4237 | 22606.5321 |
| *NPT1* | -0.078504916 | 0.62628259 | 0.99979869 | 39772.7019 | 37666.2774 |
| *RPL19A* | -0.077244062 | 0.62637729 | 0.99979869 | 246299.854 | 233459.368 |
| *PRO3* | -0.077993232 | 0.62685587 | 0.99979869 | 53169.4426 | 50371.3678 |
| *ERG2* | -0.077855063 | 0.62687163 | 0.99979869 | 60564.8181 | 57383.0524 |
| *SUA7* | -0.079938444 | 0.62688556 | 0.99979869 | 18998.145 | 17974.1048 |
| *VAM7* | -0.090327488 | 0.62708503 | 0.99979869 | 4044.48314 | 3799.0209 |
| *PPX1* | -0.0797189 | 0.62771233 | 0.99979869 | 19063.5045 | 18038.6861 |
| *NMD2* | -0.079033371 | 0.62774181 | 0.99979869 | 24638.5699 | 23325.1273 |
| *THI74* | -0.090025716 | 0.62836336 | 0.99979869 | 4023.99734 | 3780.5691 |

| | | | | | |
|---|---|---|---|---|---|
| *PBY1* | -0.081788247 | 0.62836962 | 0.99979869 | 10827.2355 | 10230.4988 |
| *KTR3* | -0.07820279 | 0.62976853 | 0.99979869 | 28627.4487 | 27116.9725 |
| *NOP6* | -0.078423675 | 0.62979729 | 0.99979869 | 25692.1255 | 24332.8006 |
| *STE4* | -0.078591934 | 0.63023003 | 0.99979869 | 22841.6722 | 21630.6368 |
| *KAR1* | -0.085734124 | 0.63055585 | 0.99979869 | 5657.98414 | 5331.54553 |
| *YBR144C* | -0.248289854 | 0.63077259 | 0.99979869 | 166.812981 | 140.438711 |
| *GRH1* | -0.077871372 | 0.63088411 | 0.99979869 | 29421.5175 | 27875.5465 |
| *POP4* | -0.09020845 | 0.63142515 | 0.99979869 | 3706.95512 | 3482.26498 |
| *HTB2* | -0.076561719 | 0.63166199 | 0.99979869 | 68906.4427 | 65345.0047 |
| *RBS1* | -0.079686939 | 0.631695 | 0.99979869 | 14312.7488 | 13543.6223 |
| *PMT5* | -0.080966584 | 0.63221665 | 0.99979869 | 10464.3441 | 9893.24089 |
| *ARG2* | -0.087721756 | 0.63241263 | 0.99979869 | 4433.71343 | 4172.15733 |
| *PMP3* | -0.076406445 | 0.63266017 | 0.99979869 | 62278.7971 | 59066.2667 |
| *FRM2* | -0.182891105 | 0.63271512 | 0.99979869 | 325.821845 | 287.028023 |
| *CYC3* | -0.079197179 | 0.63286081 | 0.99979869 | 15064.8705 | 14260.1672 |
| *YDL176W* | -0.08557818 | 0.63291958 | 0.99979869 | 5370.20736 | 5060.9191 |
| *RPL20B* | -0.075698062 | 0.63295474 | 0.99979869 | 199162.993 | 188982.326 |
| *RPO26* | -0.077036085 | 0.63343065 | 0.99979869 | 32898.2512 | 31187.6449 |
| *YDL186W* | -0.119924385 | 0.63361787 | 0.99979869 | 1012.58406 | 931.815974 |
| *YGL118C* | -0.481008293 | 0.6336538 | 0.99979869 | 42.9226383 | 30.7530025 |
| *PRP4* | -0.084603237 | 0.63366307 | 0.99979869 | 5839.42983 | 5506.83764 |
| *PPN1* | -0.076897744 | 0.6337856 | 0.99979869 | 33895.227 | 32135.8625 |
| *YTP1* | -0.088683683 | 0.63384711 | 0.99979869 | 3953.76029 | 3718.038 |
| *TOM20* | -0.078146455 | 0.63424706 | 0.99979869 | 18600.1351 | 17619.4202 |
| *VMA11* | -0.076662968 | 0.63427199 | 0.99979869 | 36394.4952 | 34511.0194 |
| *END3* | -0.078211219 | 0.63451011 | 0.99979869 | 17820.699 | 16880.323 |
| *TDA2* | -0.088082881 | 0.63460471 | 0.99979869 | 4079.60166 | 3837.97471 |
| *RNA1* | -0.076255375 | 0.6347022 | 0.99979869 | 45126.3255 | 42803.0539 |
| *CAR1* | -0.077720982 | 0.63548018 | 0.99979869 | 19355.1833 | 18340.0656 |
| *NKP1* | -0.089163132 | 0.63606395 | 0.99979869 | 3627.93845 | 3410.50797 |
| *tC(GCA)B* | -0.835352649 | 0.63617475 | 0.99979869 | 14.6327176 | 8.20080065 |
| *tF(GAA)N* | -0.835352649 | 0.63617475 | 0.99979869 | 14.6327176 | 8.20080065 |
| *TYR1* | -0.079128217 | 0.63632907 | 0.99979869 | 12316.8462 | 11659.4883 |
| *SNQ2* | -0.075570685 | 0.63654175 | 0.99979869 | 55499.9468 | 52667.592 |
| *YNL146W* | -0.136215709 | 0.63666778 | 0.99979869 | 669.202951 | 608.909449 |
| *EBS1* | -0.077329059 | 0.63669603 | 0.99979869 | 19970.733 | 18928.473 |
| *RHO2* | -0.080740701 | 0.63689939 | 0.99979869 | 8774.75298 | 8297.16006 |
| *VOA1* | -0.075599059 | 0.63694946 | 0.99979869 | 49322.0134 | 46804.0195 |
| *TSA2* | -0.085558179 | 0.63727787 | 0.99979869 | 4775.14351 | 4500.18936 |
| *ECM9* | -0.092909942 | 0.63743322 | 0.99979869 | 2752.90194 | 2581.20201 |
| *CTP1* | -0.087868379 | 0.63755543 | 0.99979869 | 3879.62119 | 3650.38139 |
| *ENT5* | -0.079450444 | 0.63794665 | 0.99979869 | 10544.3363 | 9979.3493 |
| *COG2* | -0.087282401 | 0.63823069 | 0.99979869 | 4005.46256 | 3770.3181 |
| *BET5* | -0.083725575 | 0.6383877 | 0.99979869 | 5583.84503 | 5269.01442 |
| *RAD9* | -0.079187378 | 0.63847262 | 0.99979869 | 10817.4804 | 10239.7247 |

| | | | | | |
|---|---|---|---|---|---|
| *GLC7* | -0.075152685 | 0.63860795 | 0.99979869 | 51220.3647 | 48620.4969 |
| *TOM70* | -0.075210286 | 0.63909492 | 0.99979869 | 44130.3252 | 41888.6646 |
| *MIF2* | -0.091163973 | 0.63976978 | 0.99979869 | 2936.29866 | 2756.49412 |
| *YPT7* | -0.07656279 | 0.64055758 | 0.99979869 | 18648.9108 | 17685.0266 |
| *COS3* | -0.133185516 | 0.64092959 | 0.99979869 | 685.786698 | 625.31105 |
| *PAU13* | -0.257776927 | 0.6410425 | 0.99979869 | 143.400632 | 119.93671 |
| *YEL076C* | -0.236203672 | 0.64111211 | 0.99979869 | 172.666068 | 146.589312 |
| *NRP1* | -0.076849641 | 0.64135123 | 0.99979869 | 15989.6583 | 15160.2051 |
| *PEX14* | -0.080432532 | 0.6414136 | 0.99979869 | 7659.7399 | 7244.38228 |
| *BAP3* | -0.074707162 | 0.64177196 | 0.99979869 | 39487.8517 | 37495.0857 |
| *CCT6* | -0.073819064 | 0.64221182 | 0.99979869 | 88554.2803 | 84137.1394 |
| *DOS2* | -0.077852413 | 0.64284244 | 0.99979869 | 11325.7234 | 10730.7477 |
| *RUF5-1* | -0.332365617 | 0.64287732 | 0.99979869 | 83.8942475 | 66.6315053 |
| *MAK3* | -0.079743747 | 0.64317906 | 0.99979869 | 7928.00639 | 7501.6824 |
| *PUS7* | -0.074689834 | 0.64318634 | 0.99979869 | 32002.7289 | 30388.0668 |
| *GPI18* | -0.078535887 | 0.64344104 | 0.99979869 | 9598.08723 | 9089.56243 |
| *RAM2* | -0.076813058 | 0.64355261 | 0.99979869 | 13866.9387 | 13147.9336 |
| *TVP18* | -0.07691396 | 0.64402217 | 0.99979869 | 13119.6946 | 12438.5644 |
| *YLR307C-A* | -0.716959716 | 0.64430202 | 0.99979869 | 18.5347756 | 11.2761009 |
| *COS10* | -0.086931099 | 0.64434658 | 0.99979869 | 3591.84441 | 3381.80517 |
| *RPL36A* | -0.073683237 | 0.64463525 | 0.99979869 | 51672.0279 | 49099.2186 |
| *NAM7* | -0.074415618 | 0.64525455 | 0.99979869 | 28207.0019 | 26788.9404 |
| *ARP2* | -0.073459595 | 0.64554265 | 0.99979869 | 51911.0289 | 49333.9665 |
| *YRA1* | -0.073334223 | 0.64557052 | 0.99979869 | 58447.9516 | 55551.1985 |
| *tR(ACG)J* | -0.478664205 | 0.64568335 | 0.99979869 | 39.9960948 | 28.7028023 |
| *HPF1* | -0.072517633 | 0.64711222 | 0.99979869 | 111140.368 | 105691.919 |
| *RMD1* | -0.083339303 | 0.64734543 | 0.99979869 | 4533.21591 | 4278.76774 |
| *HTL1* | -0.089833227 | 0.64751989 | 0.99979869 | 2765.58362 | 2598.62871 |
| *NNT1* | -0.075989927 | 0.64752833 | 0.99979869 | 13267.9728 | 12587.2039 |
| *ELG1* | -0.086041825 | 0.64772151 | 0.99979869 | 3573.30964 | 3366.42867 |
| *CCA1* | -0.077308738 | 0.64776177 | 0.99979869 | 9822.45556 | 9309.95894 |
| *MRX5* | -0.090897867 | 0.6479555 | 0.99979869 | 2574.38278 | 2417.18599 |
| *MEU1* | -0.074260514 | 0.64799944 | 0.99979869 | 22227.098 | 21111.9362 |
| *ARI1* | -0.074812776 | 0.64802745 | 0.99979869 | 18031.4101 | 17120.1965 |
| *HCH1* | -0.073847975 | 0.64829271 | 0.99979869 | 25879.4243 | 24588.0506 |
| *PRP24* | -0.089981858 | 0.64855922 | 0.99979869 | 2689.49349 | 2526.8717 |
| *RMD9* | -0.077954083 | 0.64859933 | 0.99979869 | 8434.29842 | 7990.65514 |
| *TFC1* | -0.078413494 | 0.64891856 | 0.99979869 | 7737.78106 | 7328.44048 |
| *ADE5%2C7* | -0.072018809 | 0.64906643 | 0.99979869 | 119170.803 | 113367.868 |
| *NUP84* | -0.075567323 | 0.64933384 | 0.99979869 | 13173.3479 | 12501.0955 |
| *MRP13* | -0.077987832 | 0.64959736 | 0.99979869 | 8046.04365 | 7622.64421 |
| *IRT1* | -0.086384304 | 0.64969378 | 0.99979869 | 3340.16167 | 3146.03215 |
| *GYL1* | -0.077337146 | 0.65011527 | 0.99979869 | 8765.97335 | 8308.43616 |
| *FAA4* | -0.071875018 | 0.65016956 | 0.99979869 | 94705.8748 | 90103.2219 |
| *DOT1* | -0.080445677 | 0.65022903 | 0.99979869 | 5649.2045 | 5342.82163 |

| | | | | | |
|---|---|---|---|---|---|
| *BER1* | -0.079587999 | 0.65064957 | 0.99979869 | 6179.8844 | 5848.19597 |
| *SSL1* | -0.076035452 | 0.65085303 | 0.99979869 | 10836.0151 | 10279.7036 |
| *CSS2* | -0.08810623 | 0.65091812 | 0.99979869 | 2896.30257 | 2724.71602 |
| *UPC2* | -0.07413701 | 0.65132206 | 0.99979869 | 17237.3413 | 16373.9236 |
| *tR(UCU)M2* | -0.494064017 | 0.65173587 | 0.99979869 | 36.0940367 | 25.627502 |
| *tW(CCA)G1* | -0.462803646 | 0.6518177 | 0.99979869 | 40.9716093 | 29.7279024 |
| *AIM32* | -0.090783197 | 0.65193235 | 0.99979869 | 2420.25149 | 2272.64688 |
| *UBS1* | -0.082606373 | 0.65251997 | 0.99979869 | 4255.19428 | 4018.39232 |
| *YLR264C-A* | -0.073400993 | 0.652533 | 0.99979869 | 20137.5459 | 19138.6185 |
| *RVB1* | -0.071998548 | 0.6526081 | 0.99979869 | 42369.5215 | 40306.9352 |
| *PSF1* | -0.089188731 | 0.65292514 | 0.99979869 | 2609.5013 | 2453.0645 |
| *MMS22* | -0.07901613 | 0.65315065 | 0.99979869 | 6113.54941 | 5787.71506 |
| *tN(GUU)F* | -0.758537827 | 0.65363657 | 0.99979869 | 15.6082321 | 9.22590074 |
| *DIS3* | -0.072547231 | 0.65408929 | 0.99979869 | 24613.2065 | 23406.1102 |
| *TFC8* | -0.079275719 | 0.65485336 | 0.99979869 | 5612.13495 | 5312.06862 |
| *CBK1* | -0.073546194 | 0.65524999 | 0.99979869 | 15336.0636 | 14573.8479 |
| *PAL1* | -0.073263797 | 0.6552764 | 0.99979869 | 16702.7594 | 15875.725 |
| *YLR456W* | -0.088805547 | 0.65530477 | 0.99979869 | 2559.75006 | 2406.93499 |
| *BFA1* | -0.091069119 | 0.65534096 | 0.99979869 | 2251.48748 | 2113.75637 |
| *FYV6* | -0.088837515 | 0.65541373 | 0.99979869 | 2549.99492 | 2397.70909 |
| *MAL32* | -0.147782993 | 0.65547479 | 0.99979869 | 456.540789 | 412.090233 |
| *BNI5* | -0.074251491 | 0.6561003 | 0.99979869 | 12037.849 | 11433.9663 |
| *TRE2* | -0.073834277 | 0.65632739 | 0.99979869 | 13159.6907 | 12503.1457 |
| *YDR239C* | -0.075868443 | 0.65637185 | 0.99979869 | 8577.69905 | 8138.26955 |
| *SEC39* | -0.074560035 | 0.65640502 | 0.99979869 | 11031.118 | 10475.4977 |
| *CDC123* | -0.079198171 | 0.6572269 | 0.99979869 | 5270.70488 | 4989.1621 |
| *NGL3* | -0.086827467 | 0.65734988 | 0.99979869 | 2787.04494 | 2624.25621 |
| *DPB2* | -0.081247596 | 0.65760775 | 0.99979869 | 4242.51259 | 4010.19152 |
| *PAU8* | -0.665429873 | 0.65762356 | 0.99979869 | 19.5102901 | 12.301201 |
| *MET7* | -0.072902872 | 0.65762751 | 0.99979869 | 15616.0362 | 14846.5245 |
| *THI73* | -0.096998753 | 0.65790477 | 0.99979869 | 1636.91334 | 1530.47442 |
| *PTM1* | -0.071268935 | 0.65794436 | 0.99979869 | 29449.8074 | 28030.3366 |
| *NOP58* | -0.070110759 | 0.65805136 | 0.99979869 | 81150.1252 | 77300.747 |
| *NMD4* | -0.086067816 | 0.65815019 | 0.99979869 | 2884.59639 | 2717.54032 |
| *DPH1* | -0.075047086 | 0.6582603 | 0.99979869 | 9118.13409 | 8655.94509 |
| *RPS1B* | -0.069458437 | 0.65857709 | 0.99979869 | 418109.419 | 398456.402 |
| *RAD27* | -0.074533144 | 0.65890683 | 0.99979869 | 9746.36543 | 9255.62864 |
| *MCY1* | -0.074789525 | 0.65922216 | 0.99979869 | 9143.49747 | 8681.57259 |
| *MRPS9* | -0.07252073 | 0.6592977 | 0.99979869 | 15491.1704 | 14731.7133 |
| *CCT5* | -0.069954238 | 0.65989374 | 0.99979869 | 56861.765 | 54170.3887 |
| *RPN7* | -0.070363332 | 0.66035102 | 0.99979869 | 36220.8536 | 34496.668 |
| *SEA4* | -0.074804259 | 0.66046973 | 0.99979869 | 8616.71963 | 8181.32375 |
| *MEC3* | -0.081396057 | 0.66052981 | 0.99979869 | 3899.13148 | 3685.23479 |
| *FAR3* | -0.095019764 | 0.66055667 | 0.99979869 | 1717.88105 | 1608.38203 |
| *YKR041W* | -0.102055296 | 0.66062676 | 0.99979869 | 1299.38532 | 1210.6432 |

| | | | | | |
|---|---|---|---|---|---|
| *SLX9* | -0.075649344 | 0.66065595 | 0.99979869 | 7443.17568 | 7062.93956 |
| *RAD23* | -0.069882425 | 0.66097531 | 0.99979869 | 47412.9315 | 45171.0351 |
| *TRS33* | -0.074490192 | 0.66201537 | 0.99979869 | 8502.58444 | 8074.71334 |
| *DCV1* | -0.16342574 | 0.66207147 | 0.99979869 | 337.528019 | 301.379424 |
| *MRX11* | -0.092295018 | 0.66225753 | 0.99979869 | 1899.32674 | 1781.62394 |
| *RPS16B* | -0.068681478 | 0.66250108 | 0.99979869 | 199308.344 | 190042.279 |
| *tV(CAC)D* | -0.538519536 | 0.66281444 | 0.99979869 | 28.2899207 | 19.4769016 |
| *YPT1* | -0.06906595 | 0.66287119 | 0.99979869 | 71396.9312 | 68059.4697 |
| *LDB18* | -0.082043564 | 0.66307643 | 0.99979869 | 3487.46436 | 3294.67166 |
| *BUD14* | -0.070698966 | 0.66322788 | 0.99979869 | 21062.3337 | 20055.058 |
| *SUL1* | -0.082625013 | 0.66394466 | 0.99979869 | 3272.85117 | 3090.67675 |
| *OPY2* | -0.070972725 | 0.66407542 | 0.99979869 | 17569.9918 | 16726.558 |
| *CSN12* | -0.075231978 | 0.66412213 | 0.99979869 | 6938.83468 | 6586.26803 |
| *PTP3* | -0.077230656 | 0.66419434 | 0.99979869 | 5346.79501 | 5068.0948 |
| *EFM5* | -0.071329073 | 0.66428896 | 0.99979869 | 15376.0596 | 14634.3288 |
| *YLR287C* | -0.073520605 | 0.66440769 | 0.99979869 | 9086.91762 | 8635.44309 |
| *RPN11* | -0.069320571 | 0.66444634 | 0.99979869 | 38353.3283 | 36554.0438 |
| *YTH1* | -0.080041341 | 0.66461194 | 0.99979869 | 3981.0747 | 3766.2177 |
| *GOT1* | -0.074391167 | 0.6647605 | 0.99979869 | 7696.80945 | 7309.98868 |
| *IDI1* | -0.069615511 | 0.66477427 | 0.99979869 | 29872.2052 | 28464.9791 |
| *DCP1* | -0.07419573 | 0.66496243 | 0.99979869 | 7879.23067 | 7484.2557 |
| *KXD1* | -0.078897733 | 0.66514393 | 0.99979869 | 4364.4519 | 4132.17843 |
| *TUF1* | -0.068856395 | 0.66549616 | 0.99979869 | 45743.8262 | 43611.8579 |
| *RPL26B* | -0.067964036 | 0.66568502 | 0.99979869 | 182988.962 | 174568.393 |
| *YKL097C* | -0.330566929 | 0.66605707 | 0.99979869 | 72.1880735 | 57.4056046 |
| *KCH1* | -0.078745199 | 0.66676712 | 0.99979869 | 4250.3167 | 4024.54292 |
| *APT1* | -0.068548651 | 0.66687032 | 0.99979869 | 45133.1541 | 43038.8269 |
| *SRL4* | -0.343505843 | 0.66712698 | 0.99979869 | 66.3349864 | 52.2801042 |
| *IMT2* | -0.572321665 | 0.66717757 | 0.99979869 | 24.3878627 | 16.4016013 |
| *tK(UUU)G2* | -0.572321665 | 0.66717757 | 0.99979869 | 24.3878627 | 16.4016013 |
| *LOT5* | -0.071591517 | 0.66739692 | 0.99979869 | 11672.0311 | 11106.9594 |
| *FYV8* | -0.07226865 | 0.66744665 | 0.99979869 | 10015.6074 | 9526.25506 |
| *MLS1* | -0.111123435 | 0.66744992 | 0.99979869 | 902.350918 | 835.456567 |
| *CBT1* | -0.080786552 | 0.66750208 | 0.99979869 | 3495.26848 | 3304.92266 |
| *DNF2* | -0.068787914 | 0.66757091 | 0.99979869 | 33239.6813 | 31691.9941 |
| *NTG2* | -0.078775064 | 0.66768335 | 0.99979869 | 4141.05908 | 3921.00781 |
| *POP6* | -0.086503053 | 0.66890568 | 0.99979869 | 2312.94489 | 2178.33767 |
| *TAF5* | -0.068917686 | 0.66924543 | 0.99979869 | 24597.5983 | 23450.1895 |
| *GIR2* | -0.069663224 | 0.66928884 | 0.99979869 | 17913.3729 | 17068.9415 |
| *YNL033W* | -0.111290809 | 0.66967627 | 0.99979869 | 876.987541 | 811.879265 |
| *SWF1* | -0.086916622 | 0.67012901 | 0.99979869 | 2213.44241 | 2084.02847 |
| *NDL1* | -0.075893752 | 0.67030304 | 0.99979869 | 5181.93306 | 4916.37999 |
| *URB1* | -0.068834356 | 0.67079389 | 0.99979869 | 21570.5768 | 20565.5578 |
| *RRG9* | -0.078233964 | 0.67130896 | 0.99979869 | 3977.17264 | 3767.2428 |
| *PSY2* | -0.070787047 | 0.67132048 | 0.99979869 | 11246.7067 | 10708.1955 |

| | | | | | |
|---|---|---|---|---|---|
| *POP7* | -0.080569105 | 0.67139169 | 0.99979869 | 3266.02257 | 3088.62655 |
| *NTO1* | -0.078256377 | 0.67153701 | 0.99979869 | 3946.93169 | 3738.54 |
| *ACK1* | -0.072424073 | 0.67169199 | 0.99979869 | 7981.65969 | 7590.86611 |
| *DOP1* | -0.068796032 | 0.6724782 | 0.99979869 | 18618.6699 | 17751.6581 |
| *RUD3* | -0.069719127 | 0.67251462 | 0.99979869 | 13733.2932 | 13085.4025 |
| *TRM10* | -0.070714567 | 0.67275401 | 0.99979869 | 10556.0425 | 10051.1063 |
| *YDR415C* | -0.071812149 | 0.67276148 | 0.99979869 | 8478.19657 | 8066.51254 |
| *YHP1* | -0.076173016 | 0.67330443 | 0.99979869 | 4611.25707 | 4374.10205 |
| *SLU7* | -0.073649477 | 0.67342864 | 0.99979869 | 6195.49263 | 5887.14977 |
| *BDF2* | -0.066842248 | 0.67376953 | 0.99979869 | 49279.0908 | 47047.9934 |
| *TPT1* | -0.073805159 | 0.67475747 | 0.99979869 | 5802.36028 | 5512.98824 |
| *RPL35B* | -0.066020577 | 0.67524817 | 0.99979869 | 93229.9213 | 89059.67 |
| *CRR1* | -0.078959716 | 0.67526123 | 0.99979869 | 3407.47217 | 3225.98996 |
| *UBC1* | -0.068175136 | 0.67547566 | 0.99979869 | 17978.7323 | 17148.8993 |
| *RTT106* | -0.06997554 | 0.67606579 | 0.99979869 | 10389.2295 | 9897.34129 |
| *ADE1* | -0.066203285 | 0.67643504 | 0.99979869 | 50289.7238 | 48034.1396 |
| *HCR1* | -0.06591775 | 0.67681971 | 0.99979869 | 62788.0157 | 59983.7313 |
| *ISW2* | -0.067865307 | 0.67689559 | 0.99979869 | 17794.3601 | 16976.6825 |
| *YOR343C-B* | -0.077383385 | 0.67755187 | 0.99979869 | 3690.37138 | 3497.64148 |
| *SHU1* | -0.088126882 | 0.67759837 | 0.99979869 | 1843.72242 | 1734.46934 |
| *RRN6* | -0.070829608 | 0.67765917 | 0.99979869 | 8128.96238 | 7739.50562 |
| *FIG4* | -0.075089386 | 0.67814219 | 0.99979869 | 4522.48525 | 4293.11914 |
| *YNR061C* | -0.06916977 | 0.67859108 | 0.99979869 | 10844.7948 | 10337.1092 |
| *PRS3* | -0.066117937 | 0.67862111 | 0.99979869 | 35150.7142 | 33576.1281 |
| *PET10* | -0.067212117 | 0.67899742 | 0.99979869 | 18874.2547 | 18015.1088 |
| *RDS3* | -0.079576322 | 0.67957734 | 0.99979869 | 2962.63756 | 2803.64872 |
| *RPL19B* | -0.064643459 | 0.67961657 | 0.99979869 | 313003.584 | 299288.22 |
| *RAD57* | -0.072981477 | 0.67962896 | 0.99979869 | 5496.04873 | 5224.93512 |
| *TIM12* | -0.078150659 | 0.67969981 | 0.99979869 | 3296.26352 | 3122.45485 |
| *SDH7* | -0.075347529 | 0.67990564 | 0.99979869 | 4202.51649 | 3988.66442 |
| *PUS6* | -0.073809627 | 0.6800606 | 0.99979869 | 4916.59311 | 4671.38107 |
| *PRP42* | -0.071755258 | 0.68010518 | 0.99979869 | 6366.20767 | 6057.31638 |
| *RPS9B* | -0.064503762 | 0.68018087 | 0.99979869 | 336640.301 | 321920.379 |
| *MSC7* | -0.065636918 | 0.68056388 | 0.99979869 | 35924.2972 | 34326.5013 |
| *BSC6* | -0.070534888 | 0.6808793 | 0.99979869 | 7448.05325 | 7092.66747 |
| *QRI7* | -0.08298549 | 0.68098348 | 0.99979869 | 2305.14078 | 2176.28747 |
| *PGC1* | -0.06765507 | 0.68109432 | 0.99979869 | 13736.2198 | 13106.9296 |
| *YGR079W* | -0.077164983 | 0.68130208 | 0.99979869 | 3446.49275 | 3266.99396 |
| *YRB2* | -0.066832559 | 0.68152307 | 0.99979869 | 17376.8399 | 16590.2197 |
| *BNA6* | -0.066792066 | 0.68216929 | 0.99979869 | 16688.1267 | 15933.1306 |
| *TEN1* | -0.092821998 | 0.68292336 | 0.99979869 | 1386.20611 | 1299.8269 |
| *CKB1* | -0.065993512 | 0.68292553 | 0.99979869 | 21415.47 | 20457.9223 |
| *GPI15* | -0.070073724 | 0.68303357 | 0.99979869 | 7356.35489 | 7007.58416 |
| *YEN1* | -0.072672023 | 0.68311265 | 0.99979869 | 5117.5491 | 4866.15009 |
| *REV7* | -0.077831545 | 0.68315225 | 0.99979869 | 3134.32811 | 2969.71494 |

| | | | | | |
|---|---|---|---|---|---|
| *YER181C* | -0.329006257 | 0.68316308 | 0.99979869 | 64.3839574 | 51.2550041 |
| *LOT6* | -0.079779565 | 0.68343704 | 0.99979869 | 2704.12621 | 2558.6498 |
| *tN(GUU)P* | -0.150862291 | 0.68353705 | 0.99979869 | 341.430077 | 307.530025 |
| *YPD1* | -0.068492664 | 0.68358021 | 0.99979869 | 9618.57303 | 9172.59553 |
| *YNL320W* | -0.07134909 | 0.68373537 | 0.99979869 | 5928.20165 | 5642.15085 |
| *BSC2* | -0.073784614 | 0.68403347 | 0.99979869 | 4408.35005 | 4188.55893 |
| *YSY6* | -0.066003714 | 0.68415339 | 0.99979869 | 18817.6748 | 17976.155 |
| *PTC3* | -0.064557824 | 0.68436551 | 0.99979869 | 42221.2433 | 40373.5667 |
| *BTS1* | -0.075012038 | 0.68449618 | 0.99979869 | 3858.15987 | 3662.68259 |
| *ERJ5* | -0.069504122 | 0.68459294 | 0.99979869 | 7583.64977 | 7226.95558 |
| *BUD7* | -0.066045028 | 0.68459648 | 0.99979869 | 17782.6539 | 16986.9335 |
| *SWC4* | -0.069375657 | 0.68512381 | 0.99979869 | 7579.74771 | 7223.88028 |
| *MTC1* | -0.064528876 | 0.68521842 | 0.99979869 | 36799.3337 | 35189.6356 |
| *SSP120* | -0.065785796 | 0.68558416 | 0.99979869 | 17965.0751 | 17164.2758 |
| *RUF23* | -0.296200428 | 0.6857325 | 0.99979869 | 78.0411605 | 63.5562051 |
| *TAD3* | -0.068693853 | 0.6862688 | 0.99979869 | 8130.91341 | 7752.83192 |
| *PSF3* | -0.077364083 | 0.68668034 | 0.99979869 | 3014.33982 | 2856.95393 |
| *PSK2* | -0.066597327 | 0.68728111 | 0.99979869 | 12191.0048 | 11641.0365 |
| *ZRT3* | -0.065235265 | 0.68751115 | 0.99979869 | 18670.3721 | 17844.9422 |
| *MTC5* | -0.06627399 | 0.68753788 | 0.99979869 | 13128.4742 | 12539.0242 |
| *GLN4* | -0.063264663 | 0.68803599 | 0.99979869 | 70221.4362 | 67208.6367 |
| *CDC34* | -0.065247224 | 0.6882163 | 0.99979869 | 17428.5422 | 16657.8763 |
| *YIL067C* | -0.067880411 | 0.68826385 | 0.99979869 | 8642.08301 | 8244.87996 |
| *CPD1* | -0.068902082 | 0.68855416 | 0.99979869 | 7120.28038 | 6788.21274 |
| *ENV10* | -0.066487174 | 0.688657 | 0.99979869 | 11515.9487 | 10997.2737 |
| *YDC1* | -0.064978655 | 0.68869325 | 0.99979869 | 18520.1429 | 17704.5035 |
| *YMR315W-A* | -0.146815467 | 0.68909889 | 0.99979869 | 347.283164 | 313.680625 |
| *PET8* | -0.067526613 | 0.68956071 | 0.99979869 | 8705.49145 | 8307.41106 |
| *ADO1* | -0.062550419 | 0.69025238 | 0.99979869 | 101915.903 | 97591.578 |
| *ISY1* | -0.069624207 | 0.69047838 | 0.99979869 | 5944.7854 | 5664.70305 |
| *PEX7* | -0.072679607 | 0.69089513 | 0.99979869 | 4102.0385 | 3900.50581 |
| *NSE1* | -0.070237131 | 0.69097484 | 0.99979869 | 5388.74213 | 5132.67611 |
| *ATP12* | -0.070498713 | 0.69143367 | 0.99979869 | 5139.98593 | 4894.85289 |
| *RSC3* | -0.06599947 | 0.69151616 | 0.99979869 | 10964.783 | 10474.4726 |
| *PAU23* | -0.099107414 | 0.69176452 | 0.99979869 | 988.196195 | 922.590074 |
| *YDR338C* | -0.068414275 | 0.6923055 | 0.99979869 | 6637.4007 | 6329.99301 |
| *SYF2* | -0.07403605 | 0.69257787 | 0.99979869 | 3466.00304 | 3292.62146 |
| *YMR209C* | -0.067228399 | 0.69269874 | 0.99979869 | 7968.00249 | 7605.21751 |
| *RIC1* | -0.066210787 | 0.69323137 | 0.99979869 | 9502.4868 | 9076.23612 |
| *NCS6* | -0.065573214 | 0.69341614 | 0.99979869 | 10889.6684 | 10405.7909 |
| *HEM14* | -0.0675714 | 0.69357216 | 0.99979869 | 7227.58698 | 6896.87335 |
| *ICL1* | -0.068333211 | 0.69364453 | 0.99979869 | 6387.66899 | 6092.16979 |
| *tP(UGG)N2* | -0.309072428 | 0.69397759 | 0.99979869 | 67.3105009 | 54.3303043 |
| *UTP7* | -0.063512861 | 0.69397882 | 0.99979869 | 20513.119 | 19629.6415 |
| *RTF1* | -0.063005678 | 0.69437727 | 0.99979869 | 25199.4907 | 24122.6551 |

| | | | | | |
|---|---|---|---|---|---|
| *HRT1* | -0.067604651 | 0.69455332 | 0.99979869 | 6907.61822 | 6591.39353 |
| *YNR064C* | -0.089864331 | 0.69462687 | 0.99979869 | 1336.45487 | 1255.7476 |
| *ADE4* | -0.062810743 | 0.69469032 | 0.99979869 | 27022.7273 | 25871.4759 |
| *HBS1* | -0.063401297 | 0.69509821 | 0.99979869 | 19130.815 | 18308.2875 |
| *SAS4* | -0.069997395 | 0.695395 | 0.99979869 | 4833.67438 | 4604.74957 |
| *SQT1* | -0.06205087 | 0.69600487 | 0.99979869 | 37009.0693 | 35451.0361 |
| *MET3* | -0.067714913 | 0.69646331 | 0.99979869 | 6308.65231 | 6019.38768 |
| *YGL114W* | -0.064138907 | 0.69646995 | 0.99979869 | 13184.0786 | 12610.7812 |
| *RPC19* | -0.063171195 | 0.69681356 | 0.99979869 | 17859.7196 | 17094.569 |
| *GPI19* | -0.081806981 | 0.69718439 | 0.99979869 | 1876.88991 | 1773.42314 |
| *ETT1* | -0.061895938 | 0.69738183 | 0.99979869 | 32809.4794 | 31431.6187 |
| *AVT4* | -0.064218746 | 0.69740407 | 0.99979869 | 12101.2574 | 11574.405 |
| *YOR105W* | -0.105589448 | 0.69772182 | 0.99979869 | 759.9258 | 706.293956 |
| *NOP10* | -0.062003859 | 0.69785621 | 0.99979869 | 28392.3497 | 27197.9554 |
| *tV(AAC)M2* | -0.288364553 | 0.69792624 | 0.99979869 | 75.114617 | 61.5060049 |
| *POL4* | -0.070677007 | 0.69823249 | 0.99979869 | 4136.18151 | 3938.43451 |
| *HIR1* | -0.066868041 | 0.69864721 | 0.99979869 | 6612.03732 | 6312.5663 |
| *YDR248C* | -0.067884105 | 0.69873663 | 0.99979869 | 5676.51891 | 5415.60373 |
| *TRE1* | -0.063177797 | 0.69880196 | 0.99979869 | 15035.6051 | 14391.38 |
| *CST26* | -0.064011283 | 0.69912993 | 0.99979869 | 11466.1975 | 10968.5709 |
| *CTO1* | -0.074109158 | 0.69924381 | 0.99979869 | 2966.53961 | 2818.00012 |
| *OSM1* | -0.062999484 | 0.69981197 | 0.99979869 | 14747.8283 | 14117.6783 |
| *MRPL35* | -0.065804895 | 0.69991096 | 0.99979869 | 7498.78001 | 7164.42447 |
| *RPS4B* | -0.060040176 | 0.70000183 | 0.99979869 | 340681.858 | 326794.73 |
| *YPL077C* | -0.073723272 | 0.70006424 | 0.99979869 | 3004.58468 | 2854.90373 |
| *APC11* | -0.067973739 | 0.7005315 | 0.99979869 | 5281.43554 | 5038.3669 |
| *YMR244W* | -0.13697194 | 0.70088125 | 0.99979869 | 369.719998 | 336.232827 |
| *CRZ1* | -0.062315448 | 0.7012936 | 0.99979869 | 16636.4244 | 15933.1306 |
| *CAJ1* | -0.060964579 | 0.70135487 | 0.99979869 | 33207.4893 | 31833.4579 |
| *RIM21* | -0.063176261 | 0.70138101 | 0.99979869 | 12464.1488 | 11930.1148 |
| *MRPS12* | -0.061378121 | 0.7016298 | 0.99979869 | 24551.7491 | 23529.1222 |
| *HSP60* | -0.059773104 | 0.70180191 | 0.99979869 | 171221.331 | 164272.288 |
| *YLR460C* | -0.082421576 | 0.70192093 | 0.99979869 | 1686.66458 | 1593.00553 |
| *SSQ1* | -0.064184703 | 0.70197028 | 0.99979869 | 9355.18411 | 8948.09861 |
| *VPS4* | -0.061070535 | 0.70204739 | 0.99979869 | 27794.3593 | 26642.3511 |
| *ATG22* | -0.066746787 | 0.70253745 | 0.99979869 | 5827.72366 | 5564.24324 |
| *TAN1* | -0.062913059 | 0.70263167 | 0.99979869 | 12355.8667 | 11828.6298 |
| *NCP1* | -0.059884365 | 0.70284472 | 0.99979869 | 74700.9988 | 71663.7216 |
| *DCG1* | -0.089763704 | 0.70307847 | 0.99979869 | 1198.90733 | 1126.58499 |
| *ERV41* | -0.061706169 | 0.7031924 | 0.99979869 | 17781.6784 | 17037.1634 |
| *YAR010C* | -0.306979967 | 0.70352404 | 0.99979869 | 63.4084429 | 51.2550041 |
| *YMR085W* | -0.073807239 | 0.70414207 | 0.99979869 | 2738.26922 | 2601.70401 |
| *MTE1* | -0.068148344 | 0.70429116 | 0.99979869 | 4598.57538 | 4386.40325 |
| *DIG2* | -0.071366401 | 0.70435374 | 0.99979869 | 3320.65138 | 3160.38355 |
| *IRE1* | -0.062602742 | 0.7043824 | 0.99979869 | 12009.5591 | 11499.5727 |

| | | | | | |
|---|---|---|---|---|---|
| *ARV1* | -0.073333325 | 0.70448074 | 0.99979869 | 2818.26141 | 2678.58651 |
| *YDR262W* | -0.062347946 | 0.70489158 | 0.99979869 | 12482.6836 | 11954.7172 |
| *RRT12* | -0.066624857 | 0.70520238 | 0.99979869 | 5405.32588 | 5161.37891 |
| *JEM1* | -0.064972272 | 0.70586927 | 0.99979869 | 6742.75627 | 6445.82931 |
| *SEC6* | -0.062622891 | 0.70588802 | 0.99979869 | 10865.2806 | 10403.7407 |
| *tK(UUU)G1* | -0.549953175 | 0.70747147 | 0.99979869 | 19.5102901 | 13.3263011 |
| *DPB3* | -0.067564586 | 0.70787734 | 0.99979869 | 4425.90931 | 4223.41234 |
| *YLR125W* | -0.101643898 | 0.70800628 | 0.99979869 | 758.950286 | 707.319056 |
| *MDL1* | -0.060839835 | 0.70806553 | 0.99979869 | 16090.1363 | 15425.706 |
| *SNU23* | -0.07319729 | 0.70818495 | 0.99979869 | 2636.81571 | 2506.3697 |
| *SKI6* | -0.06210423 | 0.70826604 | 0.99979869 | 10751.1454 | 10298.1554 |
| *RIM2* | -0.064676833 | 0.70839787 | 0.99979869 | 6399.37516 | 6118.82239 |
| *MDM32* | -0.063485406 | 0.70876697 | 0.99979869 | 7741.68312 | 7408.39829 |
| *DBP5* | -0.059258703 | 0.70880796 | 0.99979869 | 33789.8715 | 32430.0662 |
| *NOT5* | -0.060102847 | 0.70882118 | 0.99979869 | 20382.4001 | 19550.7088 |
| *YOR268C* | -0.115066056 | 0.70919476 | 0.99979869 | 530.679891 | 489.997839 |
| *CBS1* | -0.07814719 | 0.70929954 | 0.99979869 | 1880.79197 | 1781.62394 |
| *DCD1* | -0.065629515 | 0.70969379 | 0.99979869 | 5313.62751 | 5077.32071 |
| *HAP3* | -0.065837592 | 0.70970581 | 0.99979869 | 5166.32482 | 4935.85689 |
| *MRP51* | -0.060845168 | 0.70975949 | 0.99979869 | 13997.6576 | 13419.5852 |
| *YDR102C* | -0.157556802 | 0.70989309 | 0.99979869 | 245.829656 | 220.396518 |
| *CHL1* | -0.064556946 | 0.70990023 | 0.99979869 | 6155.49653 | 5886.12467 |
| *PRP11* | -0.065301846 | 0.71029434 | 0.99979869 | 5444.34646 | 5203.40802 |
| *SEC62* | -0.059307405 | 0.71035605 | 0.99979869 | 25822.8445 | 24782.8196 |
| *BOR1* | -0.066857986 | 0.71039947 | 0.99979869 | 4458.10129 | 4256.21554 |
| *YER187W* | -0.090020266 | 0.71122199 | 0.99979869 | 1068.18838 | 1003.57298 |
| *PEX27* | -0.069419961 | 0.71177023 | 0.99979869 | 3311.87175 | 3156.28315 |
| *LAG1* | -0.059076201 | 0.71179533 | 0.99979869 | 24446.3935 | 23465.566 |
| *DTD1* | -0.064016302 | 0.71200453 | 0.99979869 | 6178.90888 | 5910.72707 |
| *TMA46* | -0.058724587 | 0.71269003 | 0.99979869 | 26724.2199 | 25658.255 |
| *RPL2A* | -0.057197001 | 0.71287198 | 0.99979869 | 312087.576 | 299956.585 |
| *AUA1* | -0.065270758 | 0.71295057 | 0.99979869 | 5003.4139 | 4782.09188 |
| *SKG6* | -0.061868889 | 0.71334195 | 0.99979869 | 8593.30728 | 8232.57876 |
| *CCT3* | -0.057735297 | 0.71347934 | 0.99979869 | 53793.7719 | 51683.4959 |
| *HHT2* | -0.057225951 | 0.71371975 | 0.99979869 | 123247.478 | 118454.415 |
| *YPL216W* | -0.064747532 | 0.71409301 | 0.99979869 | 5162.42277 | 4935.85689 |
| *SMD1* | -0.065659989 | 0.71491806 | 0.99979869 | 4484.44018 | 4284.91834 |
| *MSE1* | -0.064202464 | 0.7149639 | 0.99979869 | 5399.47279 | 5164.45421 |
| *RPL27A* | -0.056693198 | 0.71525601 | 0.99979869 | 275050.217 | 264451.219 |
| *YPL038W-A* | -0.239398054 | 0.71542988 | 0.99979869 | 95.6004216 | 80.9829065 |
| *ATG3* | -0.065132813 | 0.71557648 | 0.99979869 | 4684.42066 | 4477.63716 |
| *CNL1* | -0.066248284 | 0.71567587 | 0.99979869 | 4102.0385 | 3917.93251 |
| *PET191* | -0.067508542 | 0.7159009 | 0.99979869 | 3578.18721 | 3414.60837 |
| *VPS16* | -0.061110403 | 0.71598734 | 0.99979869 | 8840.11245 | 8473.47728 |
| *snR69* | -0.12304248 | 0.71603634 | 0.99979869 | 417.520209 | 383.387431 |

| | | | | | |
|---|---|---|---|---|---|
| *GBP2* | -0.057230218 | 0.71652064 | 0.99979869 | 44386.8855 | 42660.565 |
| *MRPL27* | -0.062711762 | 0.71670665 | 0.99979869 | 6334.9912 | 6065.51718 |
| *HSP33* | -0.4232313 | 0.71672219 | 0.99979869 | 30.2409497 | 22.5522018 |
| *CAK1* | -0.062883163 | 0.71680171 | 0.99979869 | 6140.86382 | 5878.94897 |
| *NOP19* | -0.064460022 | 0.71788372 | 0.99979869 | 4739.04947 | 4531.96746 |
| *YHL042W* | -0.091132773 | 0.71798349 | 0.99979869 | 942.347013 | 884.661371 |
| *AIM22* | -0.064299261 | 0.71820202 | 0.99979869 | 4787.8252 | 4579.12207 |
| *COG4* | -0.059565782 | 0.71841251 | 0.99979869 | 11241.8292 | 10787.1282 |
| *DXO1* | -0.065302766 | 0.71860294 | 0.99979869 | 4193.73686 | 4008.14132 |
| *ILS1* | -0.056075785 | 0.71875185 | 0.99979869 | 141756.89 | 136352.662 |
| *YHL012W* | -0.085868073 | 0.71896863 | 0.99979869 | 1132.57234 | 1067.12919 |
| *LPD1* | -0.056513081 | 0.71904479 | 0.99979869 | 53170.4182 | 51127.8917 |
| *SET2* | -0.059304108 | 0.71974585 | 0.99979869 | 11094.5265 | 10647.7145 |
| *PRI1* | -0.061961295 | 0.72005939 | 0.99979869 | 6269.63173 | 6006.06138 |
| *tI(AAU)G* | -0.309558276 | 0.72016215 | 0.99979869 | 54.6288123 | 44.0793035 |
| *snR76* | -0.110467857 | 0.72043042 | 0.99979869 | 526.777833 | 487.947639 |
| *UBC6* | -0.057856138 | 0.72099713 | 0.99979869 | 16251.0962 | 15612.2742 |
| *YDR029W* | -0.074165914 | 0.72133541 | 0.99979869 | 1929.56769 | 1832.87895 |
| *FMO1* | -0.071986861 | 0.72161528 | 0.99979869 | 2204.66278 | 2097.35477 |
| *YPR158C-D* | -0.058098368 | 0.72228128 | 0.99979869 | 13384.059 | 12855.7801 |
| *ABZ1* | -0.057676054 | 0.72241908 | 0.99979869 | 15373.1331 | 14770.6671 |
| *DAD1* | -0.06499442 | 0.72257914 | 0.99979869 | 3878.64568 | 3707.787 |
| *SRB8* | -0.060863618 | 0.72314366 | 0.99979869 | 6653.00893 | 6378.17271 |
| *DIA4* | -0.062919126 | 0.72315255 | 0.99979869 | 4878.54805 | 4670.35597 |
| *MRPL10* | -0.058785769 | 0.72315536 | 0.99979869 | 10312.1638 | 9900.41659 |
| *YLR419W* | -0.058157795 | 0.72384198 | 0.99979869 | 11744.2191 | 11280.2013 |
| *SUF6* | -0.556495371 | 0.7247159 | 0.99979869 | 16.5837466 | 11.2761009 |
| *TRT2* | -0.556495371 | 0.7247159 | 0.99979869 | 16.5837466 | 11.2761009 |
| *YML037C* | -0.077293407 | 0.72518313 | 0.99979869 | 1522.77814 | 1443.34092 |
| *YTA6* | -0.058704873 | 0.72556862 | 0.99979869 | 9154.22813 | 8789.2081 |
| *YKL065W-A* | -0.136009121 | 0.72583933 | 0.99979869 | 298.507439 | 271.651522 |
| *RSM19* | -0.058542155 | 0.72584877 | 0.99979869 | 9368.84132 | 8996.27832 |
| *LYS5* | -0.065929928 | 0.72628041 | 0.99979869 | 3209.44273 | 3066.07434 |
| *ESC8* | -0.070037251 | 0.72737673 | 0.99979869 | 2254.41402 | 2147.58467 |
| *MRPL16* | -0.059744535 | 0.72742346 | 0.99979869 | 6763.24207 | 6488.88352 |
| *RAT1* | -0.055324453 | 0.7278161 | 0.99979869 | 27084.1847 | 26065.2198 |
| *VAM3* | -0.060160836 | 0.7279435 | 0.99979869 | 6161.34962 | 5909.70197 |
| *ALO1* | -0.055484252 | 0.72801642 | 0.99979869 | 23822.0642 | 22923.288 |
| *APA2* | -0.059519234 | 0.72806158 | 0.99979869 | 6852.98941 | 6576.01702 |
| *YAR028W* | -0.05923826 | 0.72821745 | 0.99979869 | 7171.00713 | 6882.52195 |
| *TFB4* | -0.060322997 | 0.72863765 | 0.99979869 | 5841.38086 | 5602.17195 |
| *URH1* | -0.05902514 | 0.72936063 | 0.99979869 | 7091.99046 | 6807.68964 |
| *YBR220C* | -0.056304898 | 0.72942677 | 0.99979869 | 14173.2503 | 13630.7558 |
| *SYN8* | -0.057056681 | 0.7298139 | 0.99979869 | 10856.5009 | 10435.5188 |
| *TAF7* | -0.056999754 | 0.73031981 | 0.99979869 | 10677.9818 | 10264.3271 |

| | | | | | |
|---|---|---|---|---|---|
| *TYW3* | -0.058845211 | 0.73037463 | 0.99979869 | 7011.02276 | 6730.80714 |
| *YOL013W-B* | -0.078350112 | 0.73091394 | 0.99979869 | 1324.7487 | 1254.7225 |
| *VPS1* | -0.054029651 | 0.73100629 | 0.99979869 | 44895.1286 | 43244.8721 |
| *tK(UUU)P* | -0.11260884 | 0.7313047 | 0.99979869 | 447.761158 | 414.140433 |
| *YAL063C-A* | -0.209489286 | 0.73140831 | 0.99979869 | 110.233139 | 95.3343076 |
| *YMR050C* | -0.085078694 | 0.73169395 | 0.99979869 | 986.245166 | 929.765774 |
| *ZRC1* | -0.053596467 | 0.73219625 | 0.99979869 | 55635.5433 | 53606.5837 |
| *UTP15* | -0.056057556 | 0.73225602 | 0.99979869 | 12390.0097 | 11917.8136 |
| *RIM1* | -0.054349564 | 0.73257882 | 0.99979869 | 25708.7093 | 24758.2172 |
| *ELC1* | -0.070247992 | 0.73260767 | 0.99979869 | 2011.51091 | 1915.91205 |
| *COX16* | -0.066736724 | 0.7327198 | 0.99979869 | 2583.16241 | 2466.3908 |
| *PET54* | -0.061077761 | 0.73322972 | 0.99979869 | 4476.63607 | 4291.06894 |
| *UBP12* | -0.055274075 | 0.73325349 | 0.99979869 | 15083.4053 | 14516.4423 |
| *CAF130* | -0.057521517 | 0.73327204 | 0.99979869 | 7968.978 | 7657.49761 |
| *PEP8* | -0.057239058 | 0.73330288 | 0.99979869 | 8465.51488 | 8136.21935 |
| *RTG2* | -0.05455797 | 0.73351547 | 0.99979869 | 20224.3667 | 19473.8263 |
| *tG(GCC)E* | -0.322746678 | 0.73384535 | 0.99979869 | 44.8736673 | 35.8785029 |
| *YHR139C-A* | -0.133788721 | 0.73408772 | 0.99979869 | 286.801265 | 261.400521 |
| *YHR112C* | -0.056166075 | 0.73425955 | 0.99979869 | 10468.2462 | 10068.533 |
| *WBP1* | -0.053150775 | 0.73563433 | 0.99979869 | 39106.4255 | 37691.9049 |
| *IMD4* | -0.052555059 | 0.73573943 | 0.99979869 | 89580.5216 | 86375.958 |
| *RPS8A* | -0.052244565 | 0.73597615 | 0.99979869 | 213573.293 | 205977.46 |
| *FAA3* | -0.055108796 | 0.73609035 | 0.99979869 | 12710.954 | 12234.5695 |
| *RPS8B* | -0.052075842 | 0.73705718 | 0.99979869 | 159581.492 | 153923.903 |
| *tF(GAA)D* | -0.452027618 | 0.73713479 | 0.99979869 | 22.4368336 | 16.4016013 |
| *RRP8* | -0.054001161 | 0.73749111 | 0.99979869 | 17266.6068 | 16632.2488 |
| *PFA4* | -0.057725038 | 0.73784052 | 0.99979869 | 6226.70909 | 5982.48408 |
| *APP1* | -0.054454733 | 0.73792161 | 0.99979869 | 13824.0161 | 13311.9497 |
| *SMM1* | -0.054140912 | 0.7379498 | 0.99979869 | 15564.3339 | 14991.0636 |
| *RRF1* | -0.06354641 | 0.7379507 | 0.99979869 | 3012.38879 | 2882.58143 |
| *YDR365W-B* | -0.061247749 | 0.7379642 | 0.99979869 | 3805.48209 | 3647.30609 |
| *ECM10* | -0.063775054 | 0.73806114 | 0.99979869 | 2942.15175 | 2814.92482 |
| *ZWF1* | -0.052218635 | 0.73809018 | 0.99979869 | 64630.7626 | 62333.2607 |
| *YAR1* | -0.053542919 | 0.73814192 | 0.99979869 | 20099.5009 | 19367.2158 |
| *YHR022C* | -0.081529234 | 0.73834653 | 0.99979869 | 1036.97192 | 979.995678 |
| *tC(GCA)P1* | -0.313131649 | 0.73839584 | 0.99979869 | 45.8491818 | 36.9036029 |
| *YJL052C-A* | -0.313131649 | 0.73839584 | 0.99979869 | 45.8491818 | 36.9036029 |
| *MSM1* | -0.058374884 | 0.73852473 | 0.99979869 | 5449.22403 | 5233.13592 |
| *MRPL40* | -0.056546556 | 0.73895739 | 0.99979869 | 7395.37547 | 7111.11927 |
| *TYS1* | -0.052008652 | 0.73906326 | 0.99979869 | 64617.1054 | 62329.1603 |
| *TDA1* | -0.053542973 | 0.73985212 | 0.99979869 | 16751.5351 | 16141.2259 |
| *YSP2* | -0.053082554 | 0.7400106 | 0.99979869 | 20606.7684 | 19862.3392 |
| *RPO31* | -0.052779711 | 0.74016795 | 0.99979869 | 24081.5511 | 23216.4667 |
| *YRM1* | -0.055614921 | 0.74024149 | 0.99979869 | 8487.95172 | 8166.97235 |
| *SPR28* | -0.170287382 | 0.74027207 | 0.99979869 | 158.03335 | 140.438711 |

| | | | | | |
|---|---|---|---|---|---|
| *YGR210C* | -0.052243049 | 0.74044893 | 0.99979869 | 34255.1919 | 33036.9254 |
| *TIF34* | -0.051536153 | 0.74087868 | 0.99979869 | 74625.8842 | 72007.1301 |
| *YER175W-A* | -0.10991684 | 0.74118821 | 0.99979869 | 429.226383 | 397.738832 |
| *ROG1* | -0.05955611 | 0.74120572 | 0.99979869 | 4223.0023 | 4052.22062 |
| *EGD2* | -0.051190937 | 0.74125143 | 0.99979869 | 151005.743 | 145741.554 |
| *CMC4* | -0.071210383 | 0.74165116 | 0.99979869 | 1615.45202 | 1537.65012 |
| *KRI1* | -0.051607129 | 0.74210861 | 0.99979869 | 45335.0856 | 43742.0456 |
| *TCP1* | -0.051482705 | 0.74215027 | 0.99979869 | 52432.9292 | 50594.8396 |
| *RCF3* | -0.062529406 | 0.74227945 | 0.99979869 | 2979.2213 | 2852.85353 |
| *EGD1* | -0.051072479 | 0.74245021 | 0.99979869 | 98412.8299 | 94989.874 |
| *TLG2* | -0.055273161 | 0.74320651 | 0.99979869 | 7848.98972 | 7553.9625 |
| *RSM25* | -0.054949426 | 0.74338132 | 0.99979869 | 8367.96343 | 8055.23644 |
| *CNE1* | -0.054315713 | 0.74345555 | 0.99979869 | 9762.94918 | 9402.21795 |
| *ECM31* | -0.058871337 | 0.74346676 | 0.99979869 | 4289.33728 | 4117.82703 |
| *tL(UAA)B2* | -0.565893204 | 0.74350128 | 0.99979869 | 13.6572031 | 9.22590074 |
| *MTR2* | -0.052726132 | 0.7436483 | 0.99979869 | 16549.6036 | 15955.6828 |
| *YMR175W-A* | -0.187002282 | 0.74423028 | 0.99979869 | 124.865857 | 109.685709 |
| *LYS14* | -0.053963571 | 0.7442843 | 0.99979869 | 10210.7103 | 9835.83528 |
| *YMR158C-B* | -0.134498739 | 0.74433413 | 0.99979869 | 256.560315 | 233.722819 |
| *DGK1* | -0.05340513 | 0.74437858 | 0.99979869 | 12018.3387 | 11581.5807 |
| *SKT5* | -0.05271297 | 0.74513564 | 0.99979869 | 14494.1945 | 13974.1643 |
| *KRE29* | -0.062416572 | 0.74520099 | 0.99979869 | 2805.57972 | 2686.78731 |
| *ENA1* | -0.058871784 | 0.74660898 | 0.99979869 | 3894.25391 | 3738.54 |
| *snR6* | -0.124475492 | 0.74682831 | 0.99979869 | 299.482953 | 274.726822 |
| *NTF2* | -0.050608517 | 0.74685387 | 0.99979869 | 43995.7042 | 42479.1223 |
| *RIO2* | -0.052749813 | 0.74713104 | 0.99979869 | 12161.7393 | 11725.0947 |
| *PPG1* | -0.054819399 | 0.74737482 | 0.99979869 | 7038.33716 | 6775.91154 |
| *NSA2* | -0.051364248 | 0.74799494 | 0.99979869 | 20165.8359 | 19460.5 |
| *SRP40* | -0.050257558 | 0.74808671 | 0.99979869 | 48877.1788 | 47203.8086 |
| *GPX1* | -0.062303804 | 0.74818211 | 0.99979869 | 2642.6688 | 2530.9721 |
| *YGR017W* | -0.051516444 | 0.74865482 | 0.99979869 | 17337.8193 | 16729.6333 |
| *MRPL19* | -0.055180173 | 0.74912318 | 0.99979869 | 6079.4064 | 5851.27127 |
| *YND1* | -0.05260658 | 0.74964374 | 0.99979869 | 10637.0102 | 10256.1263 |
| *ARG82* | -0.058172403 | 0.75041931 | 0.99979869 | 3778.16768 | 3628.85429 |
| *SPO16* | -0.104378794 | 0.75059331 | 0.99979869 | 441.908071 | 411.065133 |
| *RCH1* | -0.059427028 | 0.75071865 | 0.99979869 | 3260.16948 | 3128.60545 |
| *MSS116* | -0.050315068 | 0.75118471 | 0.99979869 | 24993.6572 | 24137.0065 |
| *RPL16A* | -0.049058196 | 0.75120445 | 0.99979869 | 154184.945 | 149030.075 |
| *SMC2* | -0.051943078 | 0.75126626 | 0.99979869 | 11678.8597 | 11265.8499 |
| *YCR095W-A* | -0.104923018 | 0.75142745 | 0.99979869 | 432.152926 | 401.839232 |
| *ACM1* | -0.072627198 | 0.75168042 | 0.99979869 | 1272.07092 | 1209.6181 |
| *KTR7* | -0.050944274 | 0.75203254 | 0.99979869 | 16013.0706 | 15457.4841 |
| *TVP23* | -0.052804914 | 0.75208757 | 0.99979869 | 8659.64227 | 8348.41507 |
| *AHC2* | -0.051734297 | 0.75220257 | 0.99979869 | 11673.9821 | 11262.7746 |
| *TEL1* | -0.051463101 | 0.75237613 | 0.99979869 | 12641.6925 | 12198.691 |

| | | | | | |
|---|---|---|---|---|---|
| *ABD1* | -0.051586353 | 0.75283327 | 0.99979869 | 11699.3455 | 11288.4021 |
| *TRM7* | -0.051965924 | 0.75342759 | 0.99979869 | 9988.29303 | 9634.91567 |
| *YDR476C* | -0.050980014 | 0.75405219 | 0.99979869 | 13156.7641 | 12699.9649 |
| *YBR296C-A* | -0.26288729 | 0.75429615 | 0.99979869 | 56.5798414 | 47.1546038 |
| *MET18* | -0.051819308 | 0.75439439 | 0.99979869 | 9785.38601 | 9440.14665 |
| *ORC2* | -0.052582683 | 0.7544315 | 0.99979869 | 8023.60681 | 7736.43032 |
| *BIR1* | -0.053452172 | 0.75485799 | 0.99979869 | 6524.24102 | 6286.9388 |
| *YKL071W* | -0.062818497 | 0.75496353 | 0.99979869 | 2183.20146 | 2090.17907 |
| *ERV29* | -0.049225308 | 0.75562709 | 0.99979869 | 27226.6099 | 26313.294 |
| *FYV5* | -0.070823115 | 0.75568443 | 0.99979869 | 1308.16495 | 1245.4966 |
| *FIN1* | -0.05529865 | 0.75569774 | 0.99979869 | 4636.62045 | 4462.26066 |
| *TBS1* | -0.051489595 | 0.75612004 | 0.99979869 | 9619.54854 | 9282.28124 |
| *DAL5* | -0.063739306 | 0.75715751 | 0.99979869 | 1951.02901 | 1866.70725 |
| *CHS1* | -0.047925734 | 0.75777267 | 0.99979869 | 74098.1309 | 71677.0479 |
| *CSM4* | -0.20744487 | 0.75780127 | 0.99979869 | 88.7718201 | 76.8825061 |
| *KIP2* | -0.05142294 | 0.75851001 | 0.99979869 | 8478.19657 | 8181.32375 |
| *KAP120* | -0.050056389 | 0.75912632 | 0.99979869 | 12178.3231 | 11763.0234 |
| *YLR365W* | -0.188858809 | 0.75932164 | 0.99979869 | 106.331081 | 93.2841074 |
| *GMH1* | -0.051338659 | 0.75976578 | 0.99979869 | 8065.55394 | 7783.58492 |
| *RFU1* | -0.056833482 | 0.75994997 | 0.99979869 | 3346.01476 | 3216.76406 |
| *ASR1* | -0.054561872 | 0.76067309 | 0.99979869 | 4336.16198 | 4175.23263 |
| *ISD11* | -0.0512495 | 0.761096 | 0.99979869 | 7664.61747 | 7397.12219 |
| *YPR114W* | -0.048162646 | 0.7620968 | 0.99979869 | 21650.5689 | 20939.7194 |
| *YDR003W-A* | -0.066952156 | 0.76261002 | 0.99979869 | 1438.8839 | 1373.63411 |
| *NSE4* | -0.062625997 | 0.76271937 | 0.99979869 | 1889.5716 | 1809.30164 |
| *RPL14B* | -0.046574713 | 0.7637947 | 0.99979869 | 88736.7015 | 85917.7383 |
| *GYP7* | -0.047378352 | 0.76385128 | 0.99979869 | 28988.3891 | 28051.8637 |
| *SRM1* | -0.047050269 | 0.76419427 | 0.99979869 | 36779.8234 | 35599.6756 |
| *ADE6* | -0.046542523 | 0.76434254 | 0.99979869 | 73416.2462 | 71085.5652 |
| *SPB1* | -0.047219063 | 0.7643933 | 0.99979869 | 30075.1122 | 29106.6917 |
| *SAC1* | -0.046689375 | 0.765032 | 0.99979869 | 45554.5764 | 44103.9059 |
| *YLR227W-B* | -0.048857009 | 0.76555965 | 0.99979869 | 11246.7067 | 10872.2115 |
| *SLF1* | -0.053186035 | 0.76595258 | 0.99979869 | 4393.71734 | 4234.68844 |
| *SLK19* | -0.049723428 | 0.7661918 | 0.99979869 | 8362.11035 | 8078.81374 |
| *tL(GAG)G* | -0.413892343 | 0.7664352 | 0.99979869 | 20.4858046 | 15.3765012 |
| *RTT107* | -0.049590481 | 0.76706233 | 0.99979869 | 8225.53832 | 7947.60093 |
| *NSI1* | -0.048752652 | 0.76721802 | 0.99979869 | 10317.0414 | 9974.2238 |
| *ZPS1* | -0.061118912 | 0.76754538 | 0.99979869 | 1910.0574 | 1830.82875 |
| *NUP85* | -0.047141406 | 0.76757845 | 0.99979869 | 19630.2784 | 18999.2049 |
| *PPS1* | -0.047519148 | 0.76768473 | 0.99979869 | 15912.5926 | 15397.0032 |
| *RPA12* | -0.048656048 | 0.7678795 | 0.99979869 | 10154.1305 | 9817.38348 |
| *MPA43* | -0.05522283 | 0.76853967 | 0.99979869 | 3143.10774 | 3025.07034 |
| *DPH6* | -0.0492361 | 0.76873129 | 0.99979869 | 8175.78708 | 7901.47143 |
| *SEC2* | -0.047853233 | 0.76884274 | 0.99979869 | 12418.2997 | 12013.1479 |
| *FMS1* | -0.048560445 | 0.76962892 | 0.99979869 | 9294.70221 | 8987.05242 |

| | | | | | |
|---|---|---|---|---|---|
| *HUF* | -0.048942407 | 0.76986148 | 0.99979869 | 8255.77927 | 7980.40414 |
| *EFB1* | -0.044933566 | 0.77024785 | 0.99979869 | 339671.224 | 329254.971 |
| *CCC1* | -0.047586737 | 0.77036789 | 0.99979869 | 12100.2819 | 11707.668 |
| *MRPL6* | -0.048684684 | 0.77064637 | 0.99979869 | 8431.37188 | 8151.59585 |
| *POB3* | -0.045891933 | 0.77115755 | 0.99979869 | 27281.2387 | 26427.0801 |
| *MAM1* | -0.18853185 | 0.77175274 | 0.99979869 | 94.6249071 | 83.0331066 |
| *SPH1* | -0.065654573 | 0.77176251 | 0.99979869 | 1322.79767 | 1263.9484 |
| *RRG1* | -0.054285138 | 0.77211641 | 0.99979869 | 3153.8384 | 3037.37154 |
| *YDL242W* | -0.082441379 | 0.77251523 | 0.99979869 | 636.035458 | 600.708648 |
| *PCM1* | -0.045296187 | 0.77292497 | 0.99979869 | 33233.8282 | 32206.5944 |
| *TIM44* | -0.045331406 | 0.77297391 | 0.99979869 | 31794.9443 | 30811.4332 |
| *ERP1* | -0.045396377 | 0.77304231 | 0.99979869 | 29560.0406 | 28644.3716 |
| *VTA1* | -0.047303889 | 0.77305153 | 0.99979869 | 10844.7948 | 10494.9746 |
| *SYS1* | -0.048342049 | 0.7731346 | 0.99979869 | 7965.07594 | 7702.60201 |
| *YPR159C-A* | -0.286019603 | 0.77319979 | 0.99979869 | 39.9960948 | 32.8032026 |
| *LNP1* | -0.047825486 | 0.77331591 | 0.99979869 | 9054.72565 | 8759.4802 |
| *GUP1* | -0.046496527 | 0.77364328 | 0.99979869 | 14045.4579 | 13600.0028 |
| *SLD7* | -0.051578894 | 0.77373369 | 0.99979869 | 4222.02678 | 4073.74773 |
| *NUP120* | -0.046261827 | 0.77427909 | 0.99979869 | 14657.1055 | 14194.5608 |
| *WHI2* | -0.04621055 | 0.77432435 | 0.99979869 | 14937.0781 | 14466.2124 |
| *YME1* | -0.044875854 | 0.7749113 | 0.99979869 | 33235.7792 | 32217.8705 |
| *GPI2* | -0.051472158 | 0.77506217 | 0.99979869 | 4094.23438 | 3950.73572 |
| *VMA8* | -0.044524087 | 0.77513201 | 0.99979869 | 47808.9904 | 46356.0508 |
| *YLR256W-A* | -0.047913457 | 0.77515875 | 0.99979869 | 7906.54507 | 7648.27171 |
| *TRS85* | -0.049510915 | 0.77522839 | 0.99979869 | 5576.04092 | 5387.92603 |
| *KHA1* | -0.045665139 | 0.77551628 | 0.99979869 | 17216.8555 | 16680.4285 |
| *YLL066C* | -0.069439205 | 0.77595459 | 0.99979869 | 1016.48612 | 968.719577 |
| *YPL257W-B* | -0.123817303 | 0.77615968 | 0.99979869 | 223.392822 | 205.020016 |
| *NNK1* | -0.046477629 | 0.77655641 | 0.99979869 | 11041.8487 | 10691.7939 |
| *TIF1* | -0.043808098 | 0.7766435 | 0.99979869 | 111906.147 | 108559.124 |
| *DAL2* | -0.057476295 | 0.77705184 | 0.99979869 | 2064.18869 | 1983.56866 |
| *LTO1* | -0.052163362 | 0.7772598 | 0.99979869 | 3478.68473 | 3355.15257 |
| *SEC63* | -0.0440298 | 0.77764828 | 0.99979869 | 45597.499 | 44226.9179 |
| *tG(GCC)F2* | -0.276390969 | 0.77773401 | 0.99979869 | 40.9716093 | 33.8283027 |
| *YOR389W* | -0.053007889 | 0.77788138 | 0.99979869 | 3090.42996 | 2978.94084 |
| *YBL059W* | -0.056825044 | 0.77800865 | 0.99979869 | 2131.4992 | 2049.17506 |
| *VPS41* | -0.046452579 | 0.77851292 | 0.99979869 | 9652.71604 | 9346.86255 |
| *PXR1* | -0.044286398 | 0.77892364 | 0.99979869 | 26303.7731 | 25508.5904 |
| *MSN1* | -0.047883206 | 0.77931733 | 0.99979869 | 6385.71796 | 6177.25309 |
| *MRPL51* | -0.046187962 | 0.78020992 | 0.99979869 | 9324.94316 | 9031.13172 |
| *YER188W* | -0.059354902 | 0.78025889 | 0.99979869 | 1675.93392 | 1608.38203 |
| *BUD16* | -0.048949732 | 0.7804388 | 0.99979869 | 4955.61369 | 4790.29268 |
| *GTR1* | -0.046279419 | 0.78075177 | 0.99979869 | 8758.16924 | 8481.67808 |
| *tV(AAC)E2* | -0.327016099 | 0.78120198 | 0.99979869 | 28.2899207 | 22.5522018 |
| *ERG10* | -0.043015 | 0.78129597 | 0.99979869 | 70409.7105 | 68341.3723 |

| | | | | | |
|---|---|---|---|---|---|
| *MTQ1* | -0.049933779 | 0.78145861 | 0.99979869 | 4083.50372 | 3944.58511 |
| *CDC23* | -0.04916622 | 0.78158516 | 0.99979869 | 4570.28546 | 4417.15625 |
| *ARF2* | -0.043120573 | 0.78165509 | 0.99979869 | 50618.4722 | 49127.9214 |
| *SGF11* | -0.051605648 | 0.78168267 | 0.99979869 | 3255.29191 | 3140.90665 |
| *CDC43* | -0.045856263 | 0.78216879 | 0.99979869 | 9038.1419 | 8755.3798 |
| *MHF1* | -0.050827137 | 0.78240247 | 0.99979869 | 3503.07259 | 3381.80517 |
| *SUB2* | -0.042399342 | 0.78331431 | 0.99979869 | 123037.743 | 119474.389 |
| *YPT6* | -0.046815801 | 0.78335433 | 0.99979869 | 6589.60049 | 6379.19781 |
| *GUK1* | -0.042845014 | 0.78368383 | 0.99979869 | 41046.7239 | 39845.6402 |
| *STM1* | -0.042079331 | 0.78391693 | 0.99979869 | 478449.869 | 464696.319 |
| *RFC4* | -0.047561529 | 0.78394588 | 0.99979869 | 5491.17115 | 5313.09372 |
| *TDH3* | -0.041891384 | 0.78453622 | 0.99979869 | 4622088.21 | 4489806.12 |
| *UCC1* | -0.044956024 | 0.7851007 | 0.99979869 | 9771.72881 | 9471.92476 |
| *RFC5* | -0.045820936 | 0.78517137 | 0.99979869 | 7579.74771 | 7342.79189 |
| *OTU1* | -0.045403295 | 0.78536749 | 0.99979869 | 8382.59615 | 8122.89305 |
| *YCR043C* | -0.04557529 | 0.78565238 | 0.99979869 | 7857.76935 | 7613.41831 |
| *YKL133C* | -0.04749932 | 0.78580657 | 0.99979869 | 5124.3777 | 4958.4091 |
| *PRS1* | -0.04200326 | 0.78614823 | 0.99979869 | 71193.0487 | 69150.1762 |
| *YEL028W* | -0.063640734 | 0.78665295 | 0.99979869 | 1145.25403 | 1095.83199 |
| *YNR029C* | -0.045260494 | 0.78685673 | 0.99979869 | 7953.36977 | 7707.72751 |
| *YDR157W* | -0.084264693 | 0.78713081 | 0.99979869 | 495.561369 | 467.445637 |
| *YRF1-6* | -0.311795488 | 0.78720649 | 0.99979869 | 29.2654352 | 23.5773019 |
| *MME1* | -0.047359951 | 0.78737081 | 0.99979869 | 4920.49517 | 4761.58988 |
| *CTA1* | -0.050735538 | 0.78772895 | 0.99979869 | 3013.36431 | 2909.23403 |
| *PUS1* | -0.042836207 | 0.78784205 | 0.99979869 | 19200.0765 | 18638.3697 |
| *ADP1* | -0.042071364 | 0.78804494 | 0.99979869 | 34796.6024 | 33796.5246 |
| *VPS45* | -0.04414313 | 0.78822117 | 0.99979869 | 10192.1756 | 9885.04009 |
| *BUG1* | -0.042554475 | 0.78859512 | 0.99979869 | 20785.2876 | 20181.1453 |
| *VTH1* | -0.087615149 | 0.78874851 | 0.99979869 | 438.981528 | 413.115333 |
| *CAX4* | -0.048571806 | 0.78907514 | 0.99979869 | 3810.35966 | 3684.20969 |
| *YDR131C* | -0.048934266 | 0.78922719 | 0.99979869 | 3604.5261 | 3484.31518 |
| *PEX17* | -0.052361971 | 0.78949441 | 0.99979869 | 2418.30046 | 2332.10269 |
| *tV(AAC)G1* | -0.397950352 | 0.78965081 | 0.99979869 | 17.5592611 | 13.3263011 |
| *RPL38* | -0.040997806 | 0.78968349 | 0.99979869 | 188670.359 | 183384.254 |
| *PKR1* | -0.043604409 | 0.78977731 | 0.99979869 | 10937.4686 | 10611.836 |
| *CDC3* | -0.04148168 | 0.79008228 | 0.99979869 | 42674.8576 | 41465.2983 |
| *MON2* | -0.042552648 | 0.79023134 | 0.99979869 | 16808.1149 | 16319.5933 |
| *LEO1* | -0.041810698 | 0.79079238 | 0.99979869 | 25667.7377 | 24934.5344 |
| *YIG1* | -0.089341716 | 0.79087784 | 0.99979869 | 406.789549 | 382.362331 |
| *MPE1* | -0.043973852 | 0.7908849 | 0.99979869 | 8900.59435 | 8633.39289 |
| *PIH1* | -0.045434659 | 0.79101441 | 0.99979869 | 6075.50434 | 5887.14977 |
| *ALE1* | -0.041410631 | 0.79136202 | 0.99979869 | 33580.1358 | 32629.9607 |
| *MUD2* | -0.044713567 | 0.79137727 | 0.99979869 | 7044.19025 | 6829.21674 |
| *CUE5* | -0.041499204 | 0.79158415 | 0.99979869 | 29243.9739 | 28414.7492 |
| *MRS3* | -0.044770347 | 0.79203673 | 0.99979869 | 6700.80914 | 6496.05922 |

| | | | | | |
|---|---|---|---|---|---|
| *NNF1* | -0.048801134 | 0.79211032 | 0.99979869 | 3340.16167 | 3229.06526 |
| *KSH1* | -0.044379621 | 0.79216566 | 0.99979869 | 7329.04048 | 7107.01887 |
| *NEM1* | -0.044817637 | 0.79248323 | 0.99979869 | 6471.56323 | 6273.6125 |
| *ICS3* | -0.051189405 | 0.7926777 | 0.99979869 | 2500.24368 | 2413.08559 |
| *SLX1* | -0.047788551 | 0.79268582 | 0.99979869 | 3760.60842 | 3638.08019 |
| *CNN1* | -0.049610155 | 0.79272082 | 0.99979869 | 2967.51513 | 2867.20493 |
| *MCM5* | -0.042481539 | 0.79301896 | 0.99979869 | 12953.8571 | 12577.978 |
| *tD(GUC)G1* | -0.242576495 | 0.79443913 | 0.99979869 | 44.8736673 | 37.928703 |
| *RPL32* | -0.039906012 | 0.79462033 | 0.99979869 | 374464.9 | 364248.812 |
| *GAL1* | -0.055218823 | 0.79463629 | 0.99979869 | 1681.78701 | 1618.63303 |
| *DED81* | -0.03995634 | 0.79501497 | 0.99979869 | 144111.782 | 140175.26 |
| *WRS1* | -0.04006964 | 0.7960022 | 0.99979869 | 57688.0258 | 56107.8279 |
| *RPS0A* | -0.039649345 | 0.79607027 | 0.99979869 | 253332.338 | 246464.813 |
| *SDO1* | -0.042582237 | 0.79623644 | 0.99979869 | 9530.77672 | 9253.57844 |
| *RSM10* | -0.042311413 | 0.79634868 | 0.99979869 | 10388.254 | 10088.0099 |
| *CYT2* | -0.04263949 | 0.7965515 | 0.99979869 | 9141.54644 | 8875.31651 |
| *PRP38* | -0.049047554 | 0.79680659 | 0.99979869 | 2810.45729 | 2716.51522 |
| *CWH43* | -0.040093562 | 0.79725318 | 0.99979869 | 37380.7404 | 36356.1995 |
| *SKI8* | -0.042133929 | 0.79754331 | 0.99979869 | 10081.9424 | 9791.75598 |
| *RNA15* | -0.042919011 | 0.79776829 | 0.99979869 | 7751.43827 | 7524.2346 |
| *YKR078W* | -0.045978383 | 0.79781056 | 0.99979869 | 4103.01401 | 3974.31302 |
| *ATG18* | -0.042700683 | 0.79808484 | 0.99979869 | 8086.03974 | 7850.21643 |
| *YBP1* | -0.042586675 | 0.79841484 | 0.99979869 | 8183.59119 | 7945.55073 |
| *tR(ACG)D* | -0.155240031 | 0.79848102 | 0.99979869 | 107.306596 | 96.3594077 |
| *OMS1* | -0.043733276 | 0.79864819 | 0.99979869 | 6021.85105 | 5842.04537 |
| *CFT2* | -0.041610929 | 0.79878271 | 0.99979869 | 11083.7958 | 10768.6764 |
| *SEN2* | -0.044163974 | 0.79893156 | 0.99979869 | 5418.00757 | 5254.66302 |
| *MSC6* | -0.041286234 | 0.79924634 | 0.99979869 | 12151.9842 | 11809.1529 |
| *YRB30* | -0.042231609 | 0.79926776 | 0.99979869 | 8606.96449 | 8358.66607 |
| *RPT1* | -0.039271998 | 0.79958071 | 0.99979869 | 65463.852 | 63705.8697 |
| *MET30* | -0.041550332 | 0.79964408 | 0.99979869 | 10558.969 | 10259.2016 |
| *IPP1* | -0.038924645 | 0.80000995 | 0.99979869 | 152580.224 | 148518.55 |
| *YOR293C-A* | -0.099656472 | 0.80003 | 0.99979869 | 278.997149 | 260.375421 |
| *RFM1* | -0.04898096 | 0.8005786 | 0.99979869 | 2544.14183 | 2459.2151 |
| *PET309* | -0.044039356 | 0.8007575 | 0.99979869 | 5106.81844 | 4953.2836 |
| *YPT31* | -0.03955429 | 0.80115759 | 0.99979869 | 27774.849 | 27023.6884 |
| *YCG1* | -0.042198404 | 0.80167358 | 0.99979869 | 7438.29811 | 7223.88028 |
| *RNA14* | -0.040465908 | 0.80216635 | 0.99979869 | 13256.2666 | 12889.6084 |
| *CDC12* | -0.039693588 | 0.80264227 | 0.99979869 | 19353.2323 | 18828.0132 |
| *YOL097W-A* | -0.096000331 | 0.80295115 | 0.99979869 | 293.629866 | 274.726822 |
| *TOA1* | -0.039899778 | 0.80298407 | 0.99979869 | 16260.8513 | 15817.2943 |
| *PPA2* | -0.041798363 | 0.80316829 | 0.99979869 | 7579.74771 | 7363.29389 |
| *SSO1* | -0.039248806 | 0.80334058 | 0.99979869 | 24271.7764 | 23620.3561 |
| *YJL136W-A* | -0.272421576 | 0.80356623 | 0.99979869 | 32.1919787 | 26.6526021 |
| *PRP39* | -0.041517793 | 0.80444853 | 0.99979869 | 7571.9436 | 7357.14329 |

| | | | | | |
|---|---|---|---|---|---|
| *CTL1* | -0.045023294 | 0.80453098 | 0.99979869 | 3707.93064 | 3594.00089 |
| *MDM38* | -0.039010274 | 0.8046159 | 0.99979869 | 23713.7821 | 23081.1534 |
| *RPS5* | -0.037811877 | 0.80476327 | 0.99979869 | 641143.252 | 624557.601 |
| *GAL80* | -0.040038512 | 0.80503817 | 0.99979869 | 12164.6659 | 11831.7051 |
| *HTD2* | -0.046086554 | 0.80514788 | 0.99979869 | 3127.49951 | 3029.17074 |
| *BRE1* | -0.039279732 | 0.80522902 | 0.99979869 | 17760.2171 | 17283.1874 |
| *YBR298C-A* | -0.191502958 | 0.80530532 | 0.99979869 | 64.3839574 | 56.3805045 |
| *EAF6* | -0.042674809 | 0.80535243 | 0.99979869 | 5434.59131 | 5276.19012 |
| *PRP46* | -0.041602903 | 0.80535882 | 0.99979869 | 6997.36555 | 6798.46374 |
| *tP(UGG)N1* | -0.413890904 | 0.80548975 | 0.99979869 | 13.6572031 | 10.2510008 |
| *FMP49* | -0.141198611 | 0.8055547 | 0.99979869 | 120.963799 | 109.685709 |
| *PPT2* | -0.05474017 | 0.80563503 | 0.99979869 | 1385.2306 | 1333.65521 |
| *REH1* | -0.039521371 | 0.80591818 | 0.99979869 | 14233.7322 | 13849.1021 |
| *YJR128W* | -0.076646335 | 0.80625925 | 0.99979869 | 489.708282 | 464.370337 |
| *SWC5* | -0.041401761 | 0.80626214 | 0.99979869 | 6996.39004 | 6798.46374 |
| *SRP102* | -0.040246294 | 0.80644004 | 0.99979869 | 9905.3743 | 9632.86547 |
| *CUE2* | -0.045478885 | 0.8067408 | 0.99979869 | 3218.22236 | 3118.35445 |
| *MRPL38* | -0.040596092 | 0.80725477 | 0.99979869 | 8284.06919 | 8054.21134 |
| *YKR070W* | -0.039456427 | 0.8075038 | 0.99979869 | 12463.1733 | 12126.934 |
| *VMA6* | -0.037715482 | 0.80755087 | 0.99979869 | 55847.23 | 54406.1617 |
| *YEL076C-A* | -0.176396267 | 0.80773117 | 0.99979869 | 74.1391025 | 65.6064052 |
| *REC102* | -0.067659502 | 0.80816581 | 0.99979869 | 668.227437 | 637.612251 |
| *MMS1* | -0.03921911 | 0.80838284 | 0.99979869 | 12752.9011 | 12410.8867 |
| *HEM12* | -0.038076975 | 0.80839937 | 0.99979869 | 27054.9193 | 26350.1976 |
| *RPB3* | -0.038250159 | 0.80865506 | 0.99979869 | 22167.5916 | 21587.5826 |
| *MID2* | -0.038865985 | 0.80871798 | 0.99979869 | 14699.0526 | 14308.3469 |
| *YNR062C* | -0.051836875 | 0.80899811 | 0.99979869 | 1594.96622 | 1538.67522 |
| *PET130* | -0.045768325 | 0.80951967 | 0.99979869 | 2835.82067 | 2747.26822 |
| *SPT4* | -0.039682332 | 0.81022341 | 0.99979869 | 9033.26433 | 8788.183 |
| *DYN1* | -0.038053827 | 0.8102729 | 0.99979869 | 19898.5449 | 19380.5421 |
| *SIR3* | -0.040304588 | 0.81036303 | 0.99979869 | 7363.18349 | 7160.32407 |
| *SDS23* | -0.039019774 | 0.81066003 | 0.99979869 | 11214.5148 | 10915.2657 |
| *DCC1* | -0.043782574 | 0.81083301 | 0.99979869 | 3555.75037 | 3449.46178 |
| *NHA1* | -0.037193758 | 0.81097014 | 0.99979869 | 41215.4879 | 40166.4965 |
| *ALK2* | -0.04013538 | 0.81110401 | 0.99979869 | 7373.91415 | 7171.60017 |
| *snR75* | -0.140622475 | 0.81124901 | 0.99979869 | 114.135197 | 103.535108 |
| *STE50* | -0.040275428 | 0.81134256 | 0.99979869 | 6985.65938 | 6793.33824 |
| *SCS22* | -0.041509363 | 0.811389 | 0.99979869 | 5190.71269 | 5043.4924 |
| *MNE1* | -0.041058884 | 0.81155937 | 0.99979869 | 5678.46994 | 5519.13884 |
| *MDM10* | -0.041528428 | 0.81173147 | 0.99979869 | 5086.33263 | 4942.00749 |
| *IMA2* | -0.053804459 | 0.81186329 | 0.99979869 | 1301.33635 | 1253.6974 |
| *YER034W* | -0.040853694 | 0.81195462 | 0.99979869 | 5831.62572 | 5668.80345 |
| *NCW2* | -0.036729785 | 0.81199617 | 0.99979869 | 66227.6798 | 64562.8534 |
| *RPL43A* | -0.036431684 | 0.81211517 | 0.99979869 | 189740.498 | 185009.038 |
| *YGL242C* | -0.038860094 | 0.81293576 | 0.99979869 | 9826.35762 | 9565.20886 |

| | | | | | |
|---|---|---|---|---|---|
| *SUF2* | -0.138686955 | 0.81299346 | 0.99979869 | 115.110712 | 104.560208 |
| *HPC2* | -0.039332558 | 0.81352668 | 0.99979869 | 7992.39035 | 7777.43432 |
| *IES2* | -0.038025577 | 0.81352747 | 0.99979869 | 13380.157 | 13032.0973 |
| *PPZ1* | -0.037916954 | 0.81365723 | 0.99979869 | 13953.7595 | 13591.802 |
| *CDC37* | -0.03710741 | 0.81397449 | 0.99979869 | 22835.8191 | 22255.9479 |
| *YNL097C-A* | -0.051852622 | 0.81401212 | 0.99979869 | 1428.15324 | 1377.73451 |
| *LIP5* | -0.038125997 | 0.81423456 | 0.99979869 | 11862.2564 | 11552.8779 |
| *YBR259W* | -0.04194982 | 0.81443634 | 0.99979869 | 4165.44694 | 4046.07002 |
| *FDC1* | -0.038029994 | 0.81456121 | 0.99979869 | 12005.657 | 11693.3166 |
| *NMA111* | -0.036892742 | 0.81522567 | 0.99979869 | 21990.048 | 21434.8427 |
| *YBR287W* | -0.03641372 | 0.81529664 | 0.99979869 | 35494.0953 | 34609.429 |
| *TRS23* | -0.038519056 | 0.81562838 | 0.99979869 | 8992.29272 | 8755.3798 |
| *LIF1* | -0.042287359 | 0.81569729 | 0.99979869 | 3742.07365 | 3633.97979 |
| *tS(AGA)D3* | -0.375923496 | 0.81610973 | 0.99979869 | 14.6327176 | 11.2761009 |
| *SMB1* | -0.038436771 | 0.81633138 | 0.99979869 | 8772.80195 | 8542.15898 |
| *SNO2* | -0.069597787 | 0.81643144 | 0.99979869 | 544.337094 | 518.700641 |
| *COX11* | -0.039904422 | 0.81694864 | 0.99979869 | 5590.67363 | 5438.15593 |
| *RAS1* | -0.037115875 | 0.81709916 | 0.99979869 | 14523.46 | 14154.5819 |
| *ALG9* | -0.037017691 | 0.81711815 | 0.99979869 | 15330.2105 | 14941.8588 |
| *NAS2* | -0.040665972 | 0.81813778 | 0.99979869 | 4498.09739 | 4373.07695 |
| *EPT1* | -0.03744005 | 0.81832322 | 0.99979869 | 10915.0318 | 10635.4133 |
| *HHF1* | -0.035275362 | 0.81876515 | 0.99979869 | 82585.1071 | 80590.2931 |
| *MUK1* | -0.038014698 | 0.81897392 | 0.99979869 | 8325.0408 | 8108.54165 |
| *SGT1* | -0.037988542 | 0.81902533 | 0.99979869 | 8366.98792 | 8149.54565 |
| *YGR153W* | -0.051813518 | 0.8190959 | 0.99979869 | 1285.72812 | 1240.3711 |
| *DCP2* | -0.035948441 | 0.8191265 | 0.99979869 | 24812.2115 | 24201.5878 |
| *OST1* | -0.035328623 | 0.8195664 | 0.99979869 | 49505.4102 | 48307.8414 |
| *PSD1* | -0.03688336 | 0.81959896 | 0.99979869 | 12437.81 | 12123.8587 |
| *tQ(UUG)H* | -0.159794396 | 0.81963531 | 0.99979869 | 79.016675 | 70.7319056 |
| *MIX14* | -0.039589342 | 0.82001038 | 0.99979869 | 5132.18182 | 4993.2625 |
| *TMN3* | -0.036889819 | 0.82014056 | 0.99979869 | 11736.415 | 11440.1169 |
| *ORM1* | -0.037661602 | 0.82050842 | 0.99979869 | 8377.71858 | 8161.84685 |
| *BUR6* | -0.035856318 | 0.82064647 | 0.99979869 | 20467.2699 | 19964.8492 |
| *TOM1* | -0.035251344 | 0.82074369 | 0.99979869 | 37930.9305 | 37015.3389 |
| *YLR035C-A* | -0.092189638 | 0.82114297 | 0.99979869 | 254.609286 | 238.848319 |
| *FMP25* | -0.036638876 | 0.82116103 | 0.99979869 | 11924.6893 | 11625.66 |
| *TRL1* | -0.038485428 | 0.82121719 | 0.99979869 | 6257.92556 | 6093.19489 |
| *GCR2* | -0.036246629 | 0.82156078 | 0.99979869 | 13958.6371 | 13612.304 |
| *YUH1* | -0.039363866 | 0.82191722 | 0.99979869 | 4904.88694 | 4772.86598 |
| *AAT1* | -0.03751965 | 0.82193349 | 0.99979869 | 7919.22676 | 7715.92832 |
| *YKT6* | -0.034959739 | 0.8228848 | 0.99979869 | 31272.0685 | 30523.38 |
| *MIX17* | -0.03466625 | 0.82323707 | 0.99979869 | 42737.2905 | 41722.5984 |
| *NIP100* | -0.039064298 | 0.82328107 | 0.99979869 | 4891.22973 | 4760.56478 |
| *SNF8* | -0.038995567 | 0.82356871 | 0.99979869 | 4894.15628 | 4763.64008 |
| *MMO1* | -0.046632026 | 0.82373066 | 0.99979869 | 1749.09751 | 1693.46534 |

| | | | | | |
|---|---|---|---|---|---|
| *CMR3* | -0.037203609 | 0.82380108 | 0.99979869 | 7686.07879 | 7490.4063 |
| *ATE1* | -0.038069972 | 0.82387673 | 0.99979869 | 5974.05084 | 5818.46806 |
| *RPS17A* | -0.033952556 | 0.82440504 | 0.99979869 | 199887.8 | 195238.511 |
| *DUS4* | -0.036902099 | 0.82450168 | 0.99979869 | 8070.43151 | 7866.61803 |
| *AAD14* | -0.043262865 | 0.82490633 | 0.99979869 | 2397.81466 | 2326.97719 |
| *ERV1* | -0.036310212 | 0.82503596 | 0.99979869 | 9602.9648 | 9364.28925 |
| *NDC1* | -0.034925548 | 0.82557475 | 0.99979869 | 19062.529 | 18606.5916 |
| *tI(AAU)L1* | -0.343502393 | 0.82563982 | 0.99979869 | 15.6082321 | 12.301201 |
| *DID4* | -0.035243591 | 0.82601697 | 0.99979869 | 14495.17 | 14145.356 |
| *GTB1* | -0.035314265 | 0.82616214 | 0.99979869 | 13679.6399 | 13348.8533 |
| *ABF1* | -0.035455374 | 0.82637377 | 0.99979869 | 12392.9363 | 12092.0806 |
| *YJL206C* | -0.040824277 | 0.82637524 | 0.99979869 | 3150.91185 | 3062.99904 |
| *ALY1* | -0.035996892 | 0.82722417 | 0.99979869 | 9032.28881 | 8809.7101 |
| *SNU114* | -0.034958322 | 0.82726684 | 0.99979869 | 14697.1015 | 14345.2505 |
| *PRE3* | -0.0339868 | 0.82756257 | 0.99979869 | 31951.0266 | 31207.1218 |
| *FMN1* | -0.037342917 | 0.8276045 | 0.99979869 | 5811.13991 | 5662.65285 |
| *SAC6* | -0.033513829 | 0.82776076 | 0.99979869 | 70694.5607 | 69071.2435 |
| *RPL36B* | -0.033199826 | 0.82806368 | 0.99979869 | 238317.218 | 232895.563 |
| *HOT13* | -0.038126252 | 0.82816704 | 0.99979869 | 4704.90646 | 4582.19737 |
| *TFC6* | -0.035923438 | 0.82839282 | 0.99979869 | 8453.80871 | 8245.90506 |
| *RRT6* | -0.051618406 | 0.82848583 | 0.99979869 | 1073.06596 | 1035.35108 |
| *TAF10* | -0.035229136 | 0.82853247 | 0.99979869 | 11077.9427 | 10810.7055 |
| *COQ3* | -0.037256067 | 0.82935617 | 0.99979869 | 5385.81559 | 5248.51242 |
| *SYC1* | -0.038222341 | 0.82938218 | 0.99979869 | 4354.69676 | 4240.83904 |
| *ASN2* | -0.0328856 | 0.82991686 | 0.99979869 | 156069.639 | 152552.319 |
| *DIA2* | -0.037399131 | 0.83011811 | 0.99979869 | 5003.4139 | 4875.37599 |
| *LSG1* | -0.033677855 | 0.83057349 | 0.99979869 | 22872.8886 | 22345.1316 |
| *YAF9* | -0.037887456 | 0.83116377 | 0.99979869 | 4287.38625 | 4176.25773 |
| *RPS18A* | -0.032556702 | 0.83125483 | 0.99979869 | 253373.309 | 247719.535 |
| *KIC1* | -0.034204543 | 0.83129792 | 0.99979869 | 13928.3961 | 13602.053 |
| *SWC3* | -0.035319689 | 0.83153088 | 0.99979869 | 8226.51383 | 8027.55874 |
| *POR2* | -0.035212864 | 0.83214099 | 0.99979869 | 8152.37473 | 7955.80173 |
| *QNS1* | -0.033193905 | 0.83242353 | 0.99979869 | 25371.1813 | 24794.0957 |
| *TAD2* | -0.03930518 | 0.8326068 | 0.99979869 | 3152.86288 | 3068.12454 |
| *SNC2* | -0.033339926 | 0.83260812 | 0.99979869 | 21260.3631 | 20774.6783 |
| *CBP3* | -0.035272766 | 0.83265193 | 0.99979869 | 7683.15225 | 7497.582 |
| *YAR009C* | -0.036466436 | 0.83277481 | 0.99979869 | 5410.20345 | 5275.16502 |
| *STE3* | -0.059400959 | 0.83307771 | 0.99979869 | 640.913031 | 615.060049 |
| *UBC9* | -0.033881284 | 0.83326344 | 0.99979869 | 13235.7808 | 12928.5622 |
| *CDC36* | -0.034983259 | 0.83332673 | 0.99979869 | 8073.35805 | 7879.94433 |
| *YGL034C* | -0.075458103 | 0.83349865 | 0.99979869 | 342.405592 | 324.956726 |
| *BIK1* | -0.034590262 | 0.83382464 | 0.99979869 | 8982.53757 | 8769.7312 |
| *NUP100* | -0.033179441 | 0.83395071 | 0.99979869 | 19271.2891 | 18833.1387 |
| *FSH1* | -0.033486966 | 0.83404495 | 0.99979869 | 15215.0998 | 14866.0014 |
| *SUP16* | -0.265501403 | 0.83443705 | 0.99979869 | 23.4123481 | 19.4769016 |

| | | | | | |
|---|---|---|---|---|---|
| *BPH1* | -0.034054171 | 0.83495257 | 0.99979869 | 10159.9836 | 9922.96879 |
| *QDR1* | -0.038443105 | 0.83501333 | 0.99979869 | 3296.26352 | 3209.58836 |
| *LCB4* | -0.033700065 | 0.83503245 | 0.99979869 | 11940.2976 | 11664.6138 |
| *SFT2* | -0.033663251 | 0.8350676 | 0.99979869 | 12124.6698 | 11845.0314 |
| *YKL096C-B* | -0.185625957 | 0.83516884 | 0.99979869 | 47.8002108 | 42.0291034 |
| *NIT3* | -0.032750445 | 0.83534311 | 0.99979869 | 21795.9206 | 21306.7052 |
| *MTG1* | -0.035571012 | 0.83545586 | 0.99979869 | 5835.52778 | 5693.40585 |
| *FAU1* | -0.03573721 | 0.83559631 | 0.99979869 | 5540.92239 | 5405.35273 |
| *YBL111C* | -0.11941207 | 0.83594207 | 0.99979869 | 118.037255 | 108.660609 |
| *PEX1* | -0.034148361 | 0.83596133 | 0.99979869 | 8922.05567 | 8713.3507 |
| *ECM7* | -0.034807842 | 0.8362219 | 0.99979869 | 6932.98159 | 6767.71074 |
| *THO2* | -0.033046918 | 0.83643434 | 0.99979869 | 14662.9585 | 14330.8991 |
| *PRM3* | -0.048438038 | 0.83653866 | 0.99979869 | 1124.76823 | 1087.63119 |
| *TEF4* | -0.031479514 | 0.8366121 | 0.99979869 | 285896.963 | 279726.235 |
| *ADI1* | -0.033786481 | 0.83720826 | 0.99979869 | 9270.31435 | 9055.73412 |
| *YGL230C* | -0.074410266 | 0.8372263 | 0.99979869 | 334.601476 | 317.781025 |
| *YPL107W* | -0.03672316 | 0.83741938 | 0.99979869 | 4048.3852 | 3946.63531 |
| *TIP20* | -0.033943959 | 0.83751632 | 0.99979869 | 8482.09863 | 8284.85886 |
| *CNB1* | -0.032641014 | 0.83818859 | 0.99979869 | 15054.1399 | 14717.3619 |
| *YBL008W-A* | -0.198556946 | 0.83831829 | 0.99979869 | 39.9960948 | 34.8534028 |
| *GON7* | -0.033969577 | 0.83833317 | 0.99979869 | 7871.42655 | 7688.25061 |
| *YGR273C* | -0.180006928 | 0.83838162 | 0.99979869 | 48.7757253 | 43.0542034 |
| *SPP1* | -0.035701685 | 0.83839853 | 0.99979869 | 4793.67828 | 4676.50657 |
| *CYS4* | -0.031086088 | 0.83909119 | 0.99979869 | 130817.471 | 128028.85 |
| *CYM1* | -0.032650242 | 0.83913382 | 0.99979869 | 13246.5115 | 12950.0893 |
| *ESL2* | -0.032827745 | 0.83925073 | 0.99979869 | 11840.7951 | 11574.405 |
| *IES3* | -0.033504066 | 0.8392811 | 0.99979869 | 8683.05462 | 8483.72828 |
| *YJU3* | -0.031621258 | 0.84012406 | 0.99979869 | 25390.6916 | 24840.2252 |
| *ERP6* | -0.034350692 | 0.84031664 | 0.99979869 | 6065.7492 | 5923.02827 |
| *EMC5* | -0.033615063 | 0.84046309 | 0.99979869 | 7577.79668 | 7403.27279 |
| *YHL005C* | -0.098394557 | 0.84107972 | 0.99979869 | 166.812981 | 155.815212 |
| *YOR381W-A* | -0.12516104 | 0.84115502 | 0.99979869 | 99.5024796 | 91.2339073 |
| *MRPL22* | -0.032355902 | 0.84184805 | 0.99979869 | 11387.1808 | 11134.6371 |
| *OST6* | -0.032642082 | 0.84195154 | 0.99979869 | 9816.60247 | 9596.98697 |
| *YEF3* | -0.03036817 | 0.84196737 | 0.99979869 | 712020.234 | 697189.017 |
| *ZRG17* | -0.033615237 | 0.84223819 | 0.99979869 | 6661.78856 | 6508.36042 |
| *GET4* | -0.031618045 | 0.84255795 | 0.99979869 | 16329.1373 | 15975.1597 |
| *HIS6* | -0.03338268 | 0.84266402 | 0.99979869 | 6968.10012 | 6808.71474 |
| *YLR108C* | -0.03647893 | 0.84267959 | 0.99979869 | 3367.47608 | 3283.39556 |
| *ABM1* | -0.051213202 | 0.84275143 | 0.99979869 | 825.285272 | 796.502764 |
| *MSR1* | -0.03350071 | 0.84301959 | 0.99979869 | 6543.75131 | 6393.54921 |
| *LSM6* | -0.031978801 | 0.84331496 | 0.99979869 | 11807.6276 | 11548.7775 |
| *TLG1* | -0.032515166 | 0.84343227 | 0.99979869 | 9043.01947 | 8841.48821 |
| *FUM1* | -0.030719634 | 0.84345537 | 0.99979869 | 33890.3495 | 33176.339 |
| *REV1* | -0.031427721 | 0.84373953 | 0.99979869 | 15706.7591 | 15368.3004 |

| | | | | | |
|---|---|---|---|---|---|
| *ORC1* | -0.03176045 | 0.84446438 | 0.99979869 | 11662.2759 | 11408.3388 |
| *PML1* | -0.034031711 | 0.84458086 | 0.99979869 | 5115.59807 | 4996.3378 |
| *PDR11* | -0.032738046 | 0.84527126 | 0.99979869 | 7149.54582 | 6989.13236 |
| *LAS1* | -0.032378457 | 0.84530431 | 0.99979869 | 8127.98686 | 7947.60093 |
| *EMC2* | -0.031089399 | 0.84542171 | 0.99979869 | 15600.428 | 15267.8406 |
| *YGL140C* | -0.030737209 | 0.84543425 | 0.99979869 | 20741.3894 | 20304.1573 |
| *ENT3* | -0.030692873 | 0.84548568 | 0.99979869 | 21420.3475 | 20969.4473 |
| *RPP0* | -0.029451291 | 0.84664251 | 0.99979869 | 592238.759 | 580271.228 |
| *SNN1* | -0.038793654 | 0.84703701 | 0.99979869 | 2067.11524 | 2012.27146 |
| *tQ(UUG)E1* | -0.343501337 | 0.84713866 | 0.99979869 | 11.7061741 | 9.22590074 |
| *ATG11* | -0.0328419 | 0.84720696 | 0.99979869 | 6038.43479 | 5902.52627 |
| *ALG2* | -0.032047265 | 0.84723106 | 0.99979869 | 7871.42655 | 7698.50161 |
| *SMT3* | -0.030031848 | 0.84729945 | 0.99979869 | 29892.691 | 29276.8583 |
| *RTR1* | -0.036609839 | 0.84736504 | 0.99979869 | 2742.17128 | 2673.46101 |
| *CHK1* | -0.032892933 | 0.84779182 | 0.99979869 | 5728.22118 | 5599.09665 |
| *MTW1* | -0.038934507 | 0.84780033 | 0.99979869 | 1985.17202 | 1932.31365 |
| *RET3* | -0.030307793 | 0.84808043 | 0.99979869 | 18750.3643 | 18360.5676 |
| *RPL27B* | -0.029219732 | 0.84856975 | 0.99979869 | 120139.489 | 117730.694 |
| *OSH7* | -0.030890973 | 0.84894014 | 0.99979869 | 11231.0985 | 10993.1733 |
| *API2* | -0.110672605 | 0.84917869 | 0.99979869 | 115.110712 | 106.610409 |
| *TAF6* | -0.030361413 | 0.84925424 | 0.99979869 | 14906.8372 | 14596.4001 |
| *YMR099C* | -0.029714735 | 0.8494631 | 0.99979869 | 25664.8111 | 25141.6046 |
| *RFA2* | -0.030683178 | 0.85001465 | 0.99979869 | 11124.7674 | 10890.6633 |
| *EUG1* | -0.030955916 | 0.85003271 | 0.99979869 | 9687.83456 | 9482.17576 |
| *YPR090W* | -0.030151893 | 0.85026746 | 0.99979869 | 14907.8127 | 14599.4754 |
| *RRN10* | -0.0364238 | 0.85060389 | 0.99979869 | 2503.17022 | 2440.76329 |
| *HMI1* | -0.036374736 | 0.8506382 | 0.99979869 | 2517.80294 | 2455.1147 |
| *PNG1* | -0.031919487 | 0.8507669 | 0.99979869 | 6301.82371 | 6163.92679 |
| *CTR9* | -0.029311436 | 0.85118969 | 0.99979869 | 27425.6148 | 26874.0237 |
| *RDL2* | -0.030413831 | 0.85161523 | 0.99979869 | 10738.4637 | 10514.4515 |
| *ECM8* | -0.047683301 | 0.85197915 | 0.99979869 | 848.69762 | 821.105166 |
| *YCL001W-B* | -0.058990047 | 0.85257136 | 0.99979869 | 462.393876 | 443.868335 |
| *DNF1* | -0.028923163 | 0.85271515 | 0.99979869 | 30588.2329 | 29981.1021 |
| *LEA1* | -0.034129426 | 0.85319089 | 0.99979869 | 3273.82668 | 3197.28716 |
| *RUP1* | -0.029604569 | 0.85340189 | 0.99979869 | 13904.9838 | 13622.555 |
| *YPK3* | -0.031248877 | 0.85462763 | 0.99979869 | 5958.4426 | 5830.76927 |
| *MFB1* | -0.031016061 | 0.85515056 | 0.99979869 | 6184.76197 | 6053.21598 |
| *MRP17* | -0.031338649 | 0.85526885 | 0.99979869 | 5556.53063 | 5437.13083 |
| *NRK1* | -0.03210958 | 0.85540086 | 0.99979869 | 4489.31776 | 4390.50365 |
| *BBP1* | -0.032863505 | 0.8557066 | 0.99979869 | 3731.34299 | 3647.30609 |
| *YHR218W* | -0.212259919 | 0.85587522 | 0.99979869 | 27.3144062 | 23.5773019 |
| *APS3* | -0.030900554 | 0.85676695 | 0.99979869 | 5736.0253 | 5614.47315 |
| *tS(GCU)F* | -0.183724673 | 0.85681871 | 0.99979869 | 36.0940367 | 31.7781025 |
| *APM1* | -0.029373798 | 0.85681886 | 0.99979869 | 10440.9318 | 10230.4988 |
| *SEC9* | -0.028455676 | 0.85713383 | 0.99979869 | 18779.6298 | 18412.8477 |

| | | | | | |
|---|---|---|---|---|---|
| *TPD3* | -0.028002705 | 0.85725473 | 0.99979869 | 30934.5405 | 30339.8871 |
| *MDE1* | -0.028760704 | 0.85744075 | 0.99979869 | 13988.878 | 13712.7638 |
| *SPE1* | -0.02828902 | 0.85765561 | 0.99979869 | 19889.7653 | 19503.5542 |
| *SPS1* | -0.066883909 | 0.85776821 | 0.99979869 | 309.238098 | 295.228824 |
| *YOL155W-A* | -0.075746013 | 0.85783155 | 0.99979869 | 231.196938 | 219.371418 |
| *TYW1* | -0.027791294 | 0.85783698 | 0.99979869 | 36056.9672 | 35369.0281 |
| *ARC19* | -0.028197704 | 0.85796785 | 0.99979869 | 20447.7596 | 20051.9827 |
| *YDR124W* | -0.036521776 | 0.8582463 | 0.99979869 | 1906.15534 | 1858.50645 |
| *PCL10* | -0.030578324 | 0.8582911 | 0.99979869 | 5710.66192 | 5590.89585 |
| *COX10* | -0.031725316 | 0.85832956 | 0.99979869 | 4194.71238 | 4103.47563 |
| *PAN6* | -0.028333 | 0.85853583 | 0.99979869 | 16293.0433 | 15976.1848 |
| *HSK3* | -0.030696547 | 0.85875005 | 0.99979869 | 5347.77052 | 5235.18612 |
| *YJR112W-A* | -0.030898089 | 0.85888191 | 0.99979869 | 5010.2425 | 4904.07879 |
| *15S_RRNA* | -0.110947186 | 0.85914232 | 0.99979869 | 98.5269651 | 91.2339073 |
| *RSC4* | -0.028565035 | 0.8591482 | 0.99979869 | 12589.0147 | 12342.205 |
| *FHL1* | -0.02878584 | 0.85950929 | 0.99979869 | 10572.6262 | 10363.7618 |
| *THO1* | -0.028046626 | 0.85958556 | 0.99979869 | 17303.6763 | 16970.5319 |
| *ECM32* | -0.027817191 | 0.85964813 | 0.99979869 | 21300.3592 | 20893.5899 |
| *tK(UUU)L* | -0.104792015 | 0.85968069 | 0.99979869 | 110.233139 | 102.510008 |
| *MRPL28* | -0.029611403 | 0.85992827 | 0.99979869 | 7016.87584 | 6874.32115 |
| *VID27* | -0.02809773 | 0.86005367 | 0.99979869 | 15336.0636 | 15040.2684 |
| *YCR085W* | -0.085059942 | 0.86049034 | 0.99979869 | 170.715039 | 160.940713 |
| *SPS22* | -0.079675467 | 0.86088666 | 0.99979869 | 196.078416 | 185.543115 |
| *HDA1* | -0.028444813 | 0.86097953 | 0.99979869 | 10767.7291 | 10557.5057 |
| *VMA10* | -0.026992903 | 0.86121556 | 0.99979869 | 46225.7304 | 45368.8794 |
| *OCA1* | -0.028481331 | 0.86132036 | 0.99979869 | 10152.1795 | 9953.72179 |
| *YLR001C* | -0.028754243 | 0.86230795 | 0.99979869 | 8062.62739 | 7903.52163 |
| *MGS1* | -0.031702712 | 0.86273825 | 0.99979869 | 3376.25571 | 3302.87246 |
| *HIM1* | -0.036158094 | 0.86355241 | 0.99979869 | 1681.78701 | 1640.16013 |
| *PSD2* | -0.027183466 | 0.86412404 | 0.99979869 | 16373.0355 | 16067.4187 |
| *BET3* | -0.028072594 | 0.86433027 | 0.99979869 | 9051.7991 | 8877.36671 |
| *SIR4* | -0.027640419 | 0.86465259 | 0.99979869 | 10998.9261 | 10790.2035 |
| *CDC8* | -0.029780482 | 0.86468816 | 0.99979869 | 4761.4863 | 4664.20537 |
| *SUI3* | -0.026075284 | 0.86478756 | 0.99979869 | 90070.2299 | 88456.9112 |
| *NAB2* | -0.026417548 | 0.86487115 | 0.99979869 | 34093.2565 | 33474.6432 |
| *GAT3* | -0.191501569 | 0.86500287 | 0.99979869 | 29.2654352 | 25.627502 |
| *IMT4* | -0.191501569 | 0.86500287 | 0.99979869 | 29.2654352 | 25.627502 |
| *TFC7* | -0.02823634 | 0.86501642 | 0.99979869 | 7854.8428 | 7702.60201 |
| *HUA2* | -0.035385431 | 0.86544226 | 0.99979869 | 1734.46479 | 1692.44024 |
| *THI13* | -0.060920206 | 0.86605482 | 0.99979869 | 333.625961 | 319.831226 |
| *APS1* | -0.028071553 | 0.8660581 | 0.99979869 | 7662.66645 | 7515.0087 |
| *tV(CAC)H* | -0.276388034 | 0.86608216 | 0.99979869 | 13.6572031 | 11.2761009 |
| *NUP170* | -0.026357903 | 0.86637604 | 0.99979869 | 23897.1789 | 23464.5409 |
| *HER2* | -0.031148574 | 0.86663204 | 0.99979869 | 3138.23017 | 3071.19985 |
| *LCD1* | -0.029439436 | 0.86690741 | 0.99979869 | 4560.53032 | 4468.41126 |

| | | | | | |
|---|---|---|---|---|---|
| *BUB1* | -0.032526428 | 0.86747332 | 0.99979869 | 2383.18194 | 2330.05249 |
| *MID1* | -0.027248439 | 0.86799365 | 0.99979869 | 9286.8981 | 9113.13973 |
| *SME1* | -0.027769144 | 0.86814157 | 0.99979869 | 7215.8808 | 7078.31606 |
| *USA1* | -0.026752315 | 0.8684359 | 0.99979869 | 11702.272 | 11487.2715 |
| *YJL213W* | -0.027762225 | 0.86850555 | 0.99979869 | 7011.02276 | 6877.39645 |
| *RCF2* | -0.025571606 | 0.86944875 | 0.99979869 | 30408.7382 | 29874.4917 |
| *RPL22A* | -0.024975129 | 0.8697821 | 0.99979869 | 202147.091 | 198677.722 |
| *YBR182C-A* | -0.086010731 | 0.87027428 | 0.99979869 | 141.449603 | 133.263011 |
| *VPS74* | -0.025542759 | 0.87033842 | 0.99979869 | 24465.9038 | 24036.5467 |
| *RPL34B* | -0.024858109 | 0.87053528 | 0.99979869 | 153026.034 | 150411.91 |
| *SET3* | -0.027349738 | 0.87061895 | 0.99979869 | 6892.00999 | 6762.58524 |
| *CLB5* | -0.030327246 | 0.87079494 | 0.99979869 | 3033.85011 | 2970.74004 |
| *PAD1* | -0.028640657 | 0.8708974 | 0.99979869 | 4435.66446 | 4348.47455 |
| *LCL2* | -0.027930307 | 0.87130858 | 0.99979869 | 5315.57854 | 5213.65902 |
| *VHC1* | -0.025865771 | 0.87150591 | 0.99979869 | 13986.927 | 13738.3913 |
| *TRX1* | -0.024876638 | 0.87177865 | 0.99979869 | 48775.7253 | 47941.8806 |
| *YJL127C-A* | -0.027547714 | 0.87207428 | 0.99979869 | 5702.8578 | 5594.99625 |
| *SAW1* | -0.029495864 | 0.87210864 | 0.99979869 | 3375.28019 | 3306.97286 |
| *SWD3* | -0.027568208 | 0.87244537 | 0.99979869 | 5510.68145 | 5406.37783 |
| *SIP18* | -0.069168594 | 0.8724521 | 0.99979869 | 220.466278 | 210.145517 |
| *YBR013C* | -0.03257376 | 0.87245892 | 0.99979869 | 1961.75967 | 1917.96225 |
| *PRP18* | -0.031472831 | 0.87263827 | 0.99979869 | 2293.4346 | 2243.94408 |
| *COG8* | -0.025926392 | 0.87294309 | 0.99979869 | 10916.9828 | 10722.5469 |
| *NPL3* | -0.024447957 | 0.87300407 | 0.99979869 | 95775.0387 | 94165.6935 |
| *SCD6* | -0.025178041 | 0.87313719 | 0.99979869 | 19319.0893 | 18984.8535 |
| *TIF11* | -0.024452157 | 0.8732021 | 0.99979869 | 78371.8599 | 77054.7229 |
| *CDC4* | -0.025631262 | 0.87365946 | 0.99979869 | 12012.4856 | 11800.9521 |
| *YGL138C* | -0.25039317 | 0.87404354 | 0.99979869 | 14.6327176 | 12.301201 |
| *YIL060W* | -0.031466852 | 0.8742834 | 0.99979869 | 2151.985 | 2105.55557 |
| *YDR249C* | -0.029959927 | 0.87463079 | 0.99979869 | 2728.51407 | 2672.43591 |
| *GLR1* | -0.024430165 | 0.87472734 | 0.99979869 | 35715.5371 | 35115.8284 |
| *ERV15* | -0.027359762 | 0.87482607 | 0.99979869 | 4977.07501 | 4883.57679 |
| *VMA21* | -0.025242175 | 0.87496948 | 0.99979869 | 13081.6495 | 12854.755 |
| *BOS1* | -0.026139398 | 0.87535465 | 0.99979869 | 7481.22075 | 7346.89229 |
| *AVT2* | -0.03046356 | 0.87539673 | 0.99979869 | 2415.37392 | 2364.90589 |
| *MXR1* | -0.025591512 | 0.87569218 | 0.99979869 | 9484.92754 | 9318.15974 |
| *CSF1* | -0.025107421 | 0.87590186 | 0.99979869 | 12542.19 | 12325.8034 |
| *RPA135* | -0.023906346 | 0.87591591 | 0.99979869 | 83057.2561 | 81692.2757 |
| *NUP82* | -0.025004323 | 0.87604578 | 0.99979869 | 13248.4625 | 13020.8212 |
| *NDT80* | -0.02970555 | 0.87621369 | 0.99979869 | 2663.1546 | 2608.87971 |
| *MSY1* | -0.027136146 | 0.87655614 | 0.99979869 | 4737.09844 | 4648.82887 |
| *SEC5* | -0.025228142 | 0.87656031 | 0.99979869 | 10557.018 | 10374.0128 |
| *YDR209C* | -0.02461285 | 0.87685764 | 0.99979869 | 16009.1686 | 15738.3616 |
| *YNL146C-A* | -0.046362559 | 0.87696838 | 0.99979869 | 534.581949 | 517.675541 |
| *GPN3* | -0.025716027 | 0.87713322 | 0.99979869 | 7632.4255 | 7497.582 |

| | | | | | |
|---|---|---|---|---|---|
| *NIT2* | -0.027950569 | 0.87734977 | 0.99979869 | 3630.86499 | 3561.19768 |
| *SPO14* | -0.024486199 | 0.87762791 | 0.99979869 | 15575.0646 | 15312.945 |
| *YJU2* | -0.028867809 | 0.87770903 | 0.99979869 | 2916.78837 | 2859.00413 |
| *RPL23A* | -0.023450317 | 0.87772526 | 0.99979869 | 156689.091 | 154162.751 |
| *SEC10* | -0.024868149 | 0.87843005 | 0.99979869 | 10368.7437 | 10191.545 |
| *PRP6* | -0.024956323 | 0.87855517 | 0.99979869 | 9681.00596 | 9514.97896 |
| *MRX4* | -0.025203337 | 0.87900567 | 0.99979869 | 8062.62739 | 7922.99853 |
| *MRPS18* | -0.024387456 | 0.87900649 | 0.99979869 | 13346.014 | 13122.3061 |
| *MSF1* | -0.026633478 | 0.8791979 | 0.99979869 | 4612.23258 | 4527.86706 |
| *TRM3* | -0.023867307 | 0.87921107 | 0.99979869 | 21344.2574 | 20994.0497 |
| *RSC6* | -0.023876953 | 0.87932414 | 0.99979869 | 20518.9721 | 20182.1704 |
| *RAD6* | -0.024437503 | 0.87937968 | 0.99979869 | 12126.6208 | 11922.9391 |
| *SWI1* | -0.02493226 | 0.87962923 | 0.99979869 | 8657.69124 | 8509.35578 |
| *SAK1* | -0.023853604 | 0.8799528 | 0.99979869 | 18251.8764 | 17952.5777 |
| *YKE2* | -0.024542104 | 0.8801656 | 0.99979869 | 10145.3509 | 9974.2238 |
| *tA(AGC)L* | -0.139972124 | 0.88030293 | 0.99979869 | 42.9226383 | 38.9538031 |
| *GCG1* | -0.024815097 | 0.88062963 | 0.99979869 | 8245.04861 | 8104.44125 |
| *YEL025C* | -0.024090143 | 0.88072023 | 0.99979869 | 12792.8972 | 12581.0533 |
| *YLR146W-A* | -0.030403192 | 0.88122091 | 0.99979869 | 1930.54321 | 1890.28455 |
| *UTP10* | -0.023150573 | 0.88129806 | 0.99979869 | 34063.0155 | 33520.7727 |
| *VPS24* | -0.024406832 | 0.88207857 | 0.99979869 | 8703.54042 | 8557.53548 |
| *CHS5* | -0.022812705 | 0.8822691 | 0.99979869 | 48035.3098 | 47281.7162 |
| *ERD1* | -0.025714622 | 0.88229196 | 0.99979869 | 4953.66266 | 4866.15009 |
| *EAF3* | -0.022904685 | 0.88414247 | 0.99979869 | 20615.5481 | 20290.831 |
| *KTR6* | -0.02297186 | 0.88466843 | 0.99979869 | 16924.2012 | 16656.8512 |
| *RAP1* | -0.022970079 | 0.88469014 | 0.99979869 | 16878.352 | 16611.7468 |
| *SIT4* | -0.022734187 | 0.88476455 | 0.99979869 | 21870.0597 | 21528.1268 |
| *OPY1* | -0.024628751 | 0.88485674 | 0.99979869 | 5976.00186 | 5874.84857 |
| *LSB5* | -0.022726126 | 0.88486344 | 0.99979869 | 21532.5317 | 21195.9944 |
| *VPS75* | -0.023018573 | 0.88511151 | 0.99979869 | 14809.2857 | 14574.873 |
| *APM3* | -0.023462496 | 0.88526335 | 0.99979869 | 10272.1677 | 10106.4617 |
| *IDS2* | -0.024768433 | 0.88574453 | 0.99979869 | 5272.65591 | 5182.90601 |
| *CHZ1* | -0.022431646 | 0.88597495 | 0.99979869 | 23794.7498 | 23427.6373 |
| *TFA1* | -0.022588441 | 0.88691997 | 0.99979869 | 15724.3183 | 15480.0363 |
| *HOC1* | -0.022022126 | 0.8875853 | 0.99979869 | 27360.2554 | 26945.7807 |
| *AIM6* | -0.022365068 | 0.88789702 | 0.99979869 | 16132.0834 | 15883.9258 |
| *RTT102* | -0.023196598 | 0.88800307 | 0.99979869 | 8534.77641 | 8398.64497 |
| *YIR021W-A* | -0.11402414 | 0.88820161 | 0.99979869 | 56.5798414 | 52.2801042 |
| *CAN1* | -0.02212848 | 0.88837915 | 0.99979869 | 18809.8707 | 18523.5585 |
| *YMR310C* | -0.023118833 | 0.88874461 | 0.99979869 | 8172.86053 | 8042.93524 |
| *SPR1* | -0.03173537 | 0.88951324 | 0.99979869 | 1214.51556 | 1188.09099 |
| *PXA1* | -0.026264071 | 0.88957134 | 0.99979869 | 2768.51017 | 2718.56542 |
| *ATG16* | -0.031357081 | 0.88967415 | 0.99979869 | 1261.34026 | 1234.2205 |
| *MDL2* | -0.021971116 | 0.89008737 | 0.99979869 | 15342.8922 | 15111.0003 |
| *TMA22* | -0.021553347 | 0.89022619 | 0.99979869 | 24933.1753 | 24563.4482 |

| | | | | | |
|---|---|---|---|---|---|
| *LPL1* | -0.022851666 | 0.89048123 | 0.99979869 | 7748.51172 | 7626.74461 |
| *SLN1* | -0.021729825 | 0.89051713 | 0.99979869 | 18124.084 | 17853.143 |
| *RAD17* | -0.024991531 | 0.89156223 | 0.99979869 | 3331.38204 | 3274.16966 |
| *MRPL37* | -0.022578439 | 0.89160905 | 0.99979869 | 7893.86338 | 7771.28372 |
| *TOM22* | -0.021246113 | 0.8916895 | 0.99979869 | 25592.6231 | 25218.4871 |
| *YDR366C* | -0.052205223 | 0.89203568 | 0.99979869 | 284.850236 | 274.726822 |
| *ICE2* | -0.021552406 | 0.89271145 | 0.99979869 | 13754.7545 | 13550.798 |
| *YHR086W-A* | -0.098393788 | 0.89278575 | 0.99979869 | 70.2370444 | 65.6064052 |
| *YEL075C* | -0.062221037 | 0.89283708 | 0.99979869 | 187.298785 | 179.392514 |
| *CDC11* | -0.021156215 | 0.89290839 | 0.99979869 | 20380.4491 | 20083.7608 |
| *ICS2* | -0.022985296 | 0.89300301 | 0.99979869 | 5676.51891 | 5586.79545 |
| *SPT2* | -0.021548393 | 0.89350872 | 0.99979869 | 11991.0243 | 11813.2533 |
| *SPT8* | -0.021305228 | 0.89358498 | 0.99979869 | 14698.0771 | 14482.614 |
| *RAD24* | -0.024153617 | 0.89369575 | 0.99979869 | 3648.42425 | 3587.85029 |
| *YLR012C* | -0.191500269 | 0.89379355 | 0.99979869 | 17.5592611 | 15.3765012 |
| *RCY1* | -0.022905841 | 0.89383206 | 0.99979869 | 5450.19955 | 5364.34873 |
| *TAF8* | -0.021767303 | 0.8939929 | 0.99979869 | 9502.4868 | 9360.18885 |
| *FPR3* | -0.020235275 | 0.8945202 | 0.99979869 | 103588.91 | 102146.098 |
| *SHE10* | -0.020563962 | 0.89471757 | 0.99979869 | 29641.0083 | 29221.5029 |
| *YDR010C* | -0.067872556 | 0.89481364 | 0.99979869 | 148.278205 | 141.463811 |
| *MDJ2* | -0.027596568 | 0.89482048 | 0.99979869 | 1718.85656 | 1686.28963 |
| *RAD1* | -0.020949337 | 0.89500314 | 0.99979869 | 15774.0696 | 15546.6678 |
| *KEL3* | -0.02046567 | 0.89525195 | 0.99979869 | 29213.7329 | 28802.237 |
| *CSE2* | -0.025246164 | 0.8956643 | 0.99979869 | 2510.97434 | 2467.4159 |
| *SGF73* | -0.021193479 | 0.89566949 | 0.99979869 | 11173.5432 | 11010.6 |
| *GSH1* | -0.02038619 | 0.89575309 | 0.99979869 | 28131.8873 | 27737.158 |
| *UTR2* | -0.019858386 | 0.89649777 | 0.99979869 | 98710.3618 | 97360.9305 |
| *FSH3* | -0.022430761 | 0.89658534 | 0.99979869 | 5182.90857 | 5102.94821 |
| *SSB2* | -0.019722063 | 0.89662333 | 0.99979869 | 365650.151 | 360685.564 |
| *YBR292C* | -0.043649635 | 0.89671743 | 0.99979869 | 394.10786 | 382.362331 |
| *BSD2* | -0.020655618 | 0.8968505 | 0.99979869 | 14514.6803 | 14308.3469 |
| *TPM2* | -0.020281158 | 0.89688835 | 0.99979869 | 22894.3499 | 22574.754 |
| *YDR444W* | -0.021215126 | 0.89748718 | 0.99979869 | 8491.85378 | 8367.89197 |
| *AIM34* | -0.02498931 | 0.89762379 | 0.99979869 | 2396.83914 | 2355.67999 |
| *DCR2* | -0.021375209 | 0.89851087 | 0.99979869 | 6882.25484 | 6781.03704 |
| *SSM4* | -0.019716101 | 0.89876745 | 0.99979869 | 32804.6018 | 32359.3343 |
| *FYV12* | -0.078886007 | 0.89885069 | 0.99979869 | 98.5269651 | 93.2841074 |
| *MSH6* | -0.020946991 | 0.89908433 | 0.99979869 | 8154.32576 | 8036.78464 |
| *PAU3* | -0.061736107 | 0.89922303 | 0.99979869 | 165.837466 | 158.890513 |
| *NHP6B* | -0.020364737 | 0.89933728 | 0.99979869 | 11887.6198 | 11720.9943 |
| *CUE1* | -0.019947405 | 0.8994707 | 0.99979869 | 17751.4375 | 17507.6843 |
| *SSN8* | -0.0223973 | 0.89958963 | 0.99979869 | 4132.27945 | 4068.62222 |
| *PHS1* | -0.020041979 | 0.89963178 | 0.99979869 | 15306.7981 | 15095.6238 |
| *PIK1* | -0.019754041 | 0.90047582 | 0.99979869 | 17565.1142 | 17326.2416 |
| *ATG10* | -0.034264771 | 0.9006579 | 0.99979869 | 676.031553 | 660.164453 |

| | | | | | |
|---|---|---|---|---|---|
| *GPR1* | -0.019797717 | 0.90093391 | 0.99979869 | 14996.5845 | 14792.1942 |
| *ADK1* | -0.018876619 | 0.90155759 | 0.99979869 | 98624.5166 | 97342.4787 |
| *COG5* | -0.021259712 | 0.90172127 | 0.99979869 | 5277.53348 | 5200.33271 |
| *MNT3* | -0.019900176 | 0.90179495 | 0.99979869 | 11501.316 | 11343.7575 |
| *SEC11* | -0.019658476 | 0.90206248 | 0.99979869 | 13665.0072 | 13480.0661 |
| *RNH1* | -0.020951196 | 0.90236948 | 0.99979869 | 5671.64134 | 5589.87075 |
| *TAF2* | -0.019306542 | 0.90243979 | 0.99979869 | 18847.9158 | 18597.3657 |
| *RAS2* | -0.018788283 | 0.9025052 | 0.99979869 | 59301.5268 | 58534.2398 |
| *YPL034W* | -0.023627485 | 0.90253249 | 0.99979869 | 2475.85582 | 2435.63779 |
| *RRP6* | -0.019284302 | 0.902872 | 0.99979869 | 17317.3335 | 17087.3933 |
| *ZAP1* | -0.022359013 | 0.9032193 | 0.99979869 | 3251.38985 | 3201.38756 |
| *VPH1* | -0.018557633 | 0.90322523 | 0.99979869 | 95772.1122 | 94548.0558 |
| *YFR012W-A* | -0.074320062 | 0.90328457 | 0.99979869 | 101.453509 | 96.3594077 |
| *AIM11* | -0.022927346 | 0.90332247 | 0.99979869 | 2786.06943 | 2742.14272 |
| *tK(UUU)O* | -0.092855485 | 0.90341435 | 0.99979869 | 63.4084429 | 59.4558047 |
| *APC5* | -0.020865301 | 0.90357637 | 0.99979869 | 5252.1701 | 5176.75541 |
| *BRO1* | -0.019214357 | 0.90371106 | 0.99979869 | 15378.0107 | 15174.5565 |
| *IRC3* | -0.020688317 | 0.90375355 | 0.99979869 | 5577.01643 | 5497.61174 |
| *CTK2* | -0.022763378 | 0.90407136 | 0.99979869 | 2775.33877 | 2731.89172 |
| *tI(UAU)D* | -0.162931506 | 0.90446523 | 0.99979869 | 19.5102901 | 17.4267014 |
| *HAT2* | -0.019114441 | 0.90544494 | 0.99979869 | 11902.2525 | 11745.5967 |
| *REF2* | -0.021056933 | 0.90554756 | 0.99979869 | 4127.40188 | 4067.59712 |
| *NSE3* | -0.020378168 | 0.90617376 | 0.99979869 | 5073.65095 | 5002.4884 |
| *TAT2* | -0.018450958 | 0.90650532 | 0.99979869 | 20070.2354 | 19815.1846 |
| *RSM28* | -0.01911959 | 0.90658079 | 0.99979869 | 9755.14506 | 9626.71487 |
| *SKI3* | -0.018664979 | 0.90667028 | 0.99979869 | 14567.3581 | 14380.1039 |
| *YNR021W* | -0.018013173 | 0.90695622 | 0.99979869 | 42275.8722 | 41751.3012 |
| *PMS1* | -0.020834289 | 0.90725722 | 0.99979869 | 3901.08251 | 3845.15041 |
| *PAR32* | -0.018141569 | 0.90731176 | 0.99979869 | 25948.6859 | 25624.4267 |
| *MIM1* | -0.019203643 | 0.90756376 | 0.99979869 | 7969.95351 | 7864.56783 |
| *AEP1* | -0.019448796 | 0.90758399 | 0.99979869 | 6863.72006 | 6771.81114 |
| *DSS1* | -0.019937418 | 0.90759713 | 0.99979869 | 5367.28081 | 5293.61682 |
| *BUD20* | -0.018214883 | 0.90764776 | 0.99979869 | 20331.6733 | 20076.5851 |
| *ROY1* | -0.019071157 | 0.90838153 | 0.99979869 | 7772.89958 | 7670.82391 |
| *YKL102C* | -0.073667016 | 0.90849764 | 0.99979869 | 91.6983636 | 87.133507 |
| *NAT3* | -0.019564713 | 0.90867716 | 0.99979869 | 5723.34361 | 5646.25125 |
| *GLN3* | -0.019070804 | 0.90991768 | 0.99979869 | 6426.68957 | 6342.29421 |
| *SLD2* | -0.021169281 | 0.9109054 | 0.99979869 | 2743.14679 | 2703.18892 |
| *PTC7* | -0.018088141 | 0.91128145 | 0.99979869 | 10252.6575 | 10124.9135 |
| *SSY1* | -0.018316159 | 0.91212918 | 0.99979869 | 7609.98866 | 7513.9836 |
| *KES1* | -0.016971044 | 0.91252188 | 0.99979869 | 36708.6109 | 36279.317 |
| *CUS1* | -0.019157018 | 0.91267186 | 0.99979869 | 4643.44905 | 4582.19737 |
| *AHP1* | -0.016544908 | 0.91311619 | 0.99979869 | 374886.323 | 370611.608 |
| *PSF2* | -0.020349142 | 0.91312211 | 0.99979869 | 2972.3927 | 2930.76113 |
| *CCT4* | -0.016434556 | 0.91447389 | 0.99979869 | 67897.7607 | 67128.6789 |

| | | | | | |
|---|---|---|---|---|---|
| *ABP140* | -0.016494362 | 0.91474266 | 0.99979869 | 42010.5322 | 41532.9549 |
| *RPL24B* | -0.0162643 | 0.91475158 | 0.99979869 | 186991.498 | 184895.252 |
| *VPS68* | -0.016728242 | 0.91526796 | 0.99979869 | 19364.9385 | 19141.6938 |
| *TIM13* | -0.017249374 | 0.91530177 | 0.99979869 | 10362.8906 | 10239.7247 |
| *CDC50* | -0.017378019 | 0.91544291 | 0.99979869 | 9075.21145 | 8966.55042 |
| *PXL1* | -0.017016097 | 0.91677327 | 0.99979869 | 9750.26749 | 9635.94077 |
| *COA4* | -0.018020407 | 0.91754844 | 0.99979869 | 4765.38836 | 4706.23448 |
| *tX(XXX)D* | -0.130100568 | 0.91794652 | 0.99979869 | 22.4368336 | 20.5020016 |
| *MDM1* | -0.016776243 | 0.91808524 | 0.99979869 | 9496.63372 | 9386.84145 |
| *SXM1* | -0.015884855 | 0.91837806 | 0.99979869 | 30978.4387 | 30639.2163 |
| *YPR097W* | -0.016116037 | 0.91889031 | 0.99979869 | 16351.5742 | 16169.9287 |
| *ATG29* | -0.020564528 | 0.91918018 | 0.99979869 | 1940.29835 | 1912.83675 |
| *SNA3* | -0.015743368 | 0.91950658 | 0.99979869 | 25493.1206 | 25216.4369 |
| *SSK2* | -0.015995774 | 0.91969245 | 0.99979869 | 15445.3212 | 15275.0163 |
| *YIL054W* | -0.052204851 | 0.92018051 | 0.99979869 | 142.425118 | 137.363411 |
| *UBP14* | -0.01581422 | 0.9204144 | 0.99979869 | 16273.533 | 16096.1215 |
| *YMR130W* | -0.016430987 | 0.92126992 | 0.99979869 | 7421.71436 | 7337.66639 |
| *PPM1* | -0.016397807 | 0.92129341 | 0.99979869 | 7572.91911 | 7487.331 |
| *WTM1* | -0.015089068 | 0.92129951 | 0.99979869 | 79282.9905 | 78458.085 |
| *TAZ1* | -0.016687019 | 0.92153197 | 0.99979869 | 6068.67574 | 5998.88568 |
| *INA17* | -0.017279542 | 0.92174493 | 0.99979869 | 4379.08462 | 4326.94745 |
| *ALG13* | -0.016920766 | 0.92180063 | 0.99979869 | 5167.30034 | 5107.04861 |
| *CPR4* | -0.015379025 | 0.92189681 | 0.99979869 | 20474.0985 | 20257.0027 |
| *VFA1* | -0.020204784 | 0.92217268 | 0.99979869 | 1773.48537 | 1748.82074 |
| *TOH1* | -0.015160019 | 0.92250933 | 0.99979869 | 25023.8981 | 24762.3176 |
| *YGP1* | -0.014869815 | 0.92266211 | 0.99979869 | 60531.6506 | 59910.9492 |
| *YHR180W* | -0.021881138 | 0.92297173 | 0.99979869 | 1244.75651 | 1226.0197 |
| *RAD3* | -0.015696669 | 0.92302788 | 0.99979869 | 10042.9218 | 9934.24489 |
| *DUG2* | -0.015562494 | 0.92307732 | 0.99979869 | 11390.1074 | 11267.9001 |
| *OCA6* | -0.016866264 | 0.92348545 | 0.99979869 | 4431.7624 | 4380.25265 |
| *IOC2* | -0.015170088 | 0.92373433 | 0.99979869 | 15788.7023 | 15623.5503 |
| *DAL3* | -0.02749316 | 0.92389853 | 0.99979869 | 588.235247 | 577.131346 |
| *REC107* | -0.015786665 | 0.92401473 | 0.99979869 | 7807.04259 | 7722.07892 |
| *AST1* | -0.015298639 | 0.92412318 | 0.99979869 | 12058.3348 | 11931.1399 |
| *TSC3* | -0.01491558 | 0.92423385 | 0.99979869 | 20479.9515 | 20269.3039 |
| *COX14* | -0.015682832 | 0.92429974 | 0.99979869 | 8082.13768 | 7994.75554 |
| *FAP1* | -0.015097235 | 0.92441666 | 0.99979869 | 14412.2513 | 14262.2174 |
| *OST5* | -0.015386966 | 0.92448636 | 0.99979869 | 10144.3753 | 10036.7549 |
| *CSS3* | -0.018330864 | 0.92451356 | 0.99979869 | 2423.17803 | 2392.58359 |
| *SWD2* | -0.016215808 | 0.92483645 | 0.99979869 | 5290.21517 | 5231.08572 |
| *YOL024W* | -0.072858561 | 0.9249507 | 0.99979869 | 61.4574139 | 58.4307047 |
| *ICL2* | -0.016884766 | 0.92500883 | 0.99979869 | 3790.84937 | 3746.7408 |
| *PAU11* | -0.150857698 | 0.92501403 | 0.99979869 | 13.6572031 | 12.301201 |
| *VAS1* | -0.014299482 | 0.92510701 | 0.99979869 | 131842.737 | 130542.395 |
| *TPK3* | -0.015429423 | 0.92518478 | 0.99979869 | 8555.26222 | 8464.25138 |

| | | | | | |
|---|---|---|---|---|---|
| *RBK1* | -0.01510606 | 0.925408 | 0.99979869 | 11163.788 | 11047.5036 |
| *ARR2* | -0.024557234 | 0.92575284 | 0.99979869 | 756.999257 | 744.222659 |
| *PTC1* | -0.015570632 | 0.92594717 | 0.99979869 | 6793.48302 | 6720.55614 |
| *CSM1* | -0.017959147 | 0.9259761 | 0.99979869 | 2432.93318 | 2402.83459 |
| *JHD1* | -0.016222333 | 0.92611515 | 0.99979869 | 4562.48135 | 4511.46546 |
| *YEA4* | -0.017001394 | 0.92637273 | 0.99979869 | 3212.36927 | 3174.73495 |
| *AOS1* | -0.015352962 | 0.926396 | 0.99979869 | 7427.56745 | 7348.94249 |
| *ERG20* | -0.014118046 | 0.92658581 | 0.99979869 | 58068.4765 | 57502.9891 |
| *TAF3* | -0.01561783 | 0.92679051 | 0.99979869 | 5857.96461 | 5794.89076 |
| *24-May* | -0.019145287 | 0.92781279 | 0.99979869 | 1617.40305 | 1596.08083 |
| *DAS1* | -0.01561227 | 0.92868863 | 0.99979869 | 4660.0328 | 4609.87507 |
| *GYP6* | -0.014123021 | 0.92923924 | 0.99979869 | 14520.5334 | 14379.0788 |
| *MRPS8* | -0.015015379 | 0.9292692 | 0.99979869 | 6132.08419 | 6068.59248 |
| *VMA4* | -0.013481534 | 0.92947342 | 0.99979869 | 104212.264 | 103242.955 |
| *SMC3* | -0.014959063 | 0.9298477 | 0.99979869 | 5867.71975 | 5807.19196 |
| *DAP2* | -0.013628271 | 0.93019387 | 0.99979869 | 26311.5773 | 26064.1947 |
| *YGL159W* | -0.015200079 | 0.93031117 | 0.99979869 | 4798.55586 | 4748.26358 |
| *IPK1* | -0.017923821 | 0.9309429 | 0.99979869 | 1768.6078 | 1746.77054 |
| *PNP1* | -0.014157552 | 0.93119191 | 0.99979869 | 8763.04681 | 8677.47219 |
| *BNI4* | -0.014322792 | 0.93144042 | 0.99979869 | 7260.75447 | 7189.02687 |
| *YOS1* | -0.014136904 | 0.93165784 | 0.99979869 | 8172.86053 | 8093.16515 |
| *YHR177W* | -0.016380225 | 0.9317993 | 0.99979869 | 2557.79904 | 2528.9219 |
| *EFM1* | -0.013681269 | 0.93241841 | 0.99979869 | 11123.7919 | 11018.8008 |
| *MRE11* | -0.01484107 | 0.93278424 | 0.99979869 | 4347.86815 | 4303.37014 |
| *YPL191C* | -0.015610166 | 0.9331353 | 0.99979869 | 2995.80505 | 2963.56434 |
| *CRF1* | -0.013989992 | 0.93363982 | 0.99979869 | 6559.35954 | 6496.05922 |
| *UPS3* | -0.016327753 | 0.93381455 | 0.99979869 | 2229.05065 | 2203.96518 |
| *ATM1* | -0.013282948 | 0.93383093 | 0.99979869 | 12866.0608 | 12748.1446 |
| *MRPL3* | -0.013324741 | 0.93465537 | 0.99979869 | 9905.3743 | 9814.30818 |
| *PUS9* | -0.013411759 | 0.93471058 | 0.99979869 | 8923.03119 | 8840.46311 |
| *TDA8* | -0.020697441 | 0.93491122 | 0.99979869 | 850.648649 | 838.531867 |
| *UFD2* | -0.01280717 | 0.93494968 | 0.99979869 | 19928.7858 | 19752.6535 |
| *THP2* | -0.01578702 | 0.93500294 | 0.99979869 | 2408.54532 | 2382.33259 |
| *FMP52* | -0.012628839 | 0.93529018 | 0.99979869 | 26270.6056 | 26041.6425 |
| *RDL1* | -0.012651115 | 0.93539292 | 0.99979869 | 23424.0543 | 23219.542 |
| *POP5* | -0.013585682 | 0.93569696 | 0.99979869 | 6402.3017 | 6342.29421 |
| *YRF1-3* | -0.076566649 | 0.93571981 | 0.99979869 | 39.9960948 | 37.928703 |
| *KTI12* | -0.013131062 | 0.93576804 | 0.99979869 | 9533.70327 | 9447.32235 |
| *YNL247W* | -0.012214422 | 0.93649163 | 0.99979869 | 54333.2314 | 53875.1599 |
| *MIC26* | -0.012918053 | 0.93670197 | 0.99979869 | 9762.94918 | 9675.91967 |
| *NMD5* | -0.012522876 | 0.93679447 | 0.99979869 | 16874.4499 | 16728.6082 |
| *CGI121* | -0.013988344 | 0.93748019 | 0.99979869 | 3932.29897 | 3894.35521 |
| *PDR8* | -0.013205417 | 0.93751028 | 0.99979869 | 6378.88936 | 6320.7671 |
| *SDC1* | -0.013126829 | 0.93767159 | 0.99979869 | 6607.15975 | 6547.31422 |
| *RUB1* | -0.012749597 | 0.9377993 | 0.99979869 | 9163.00776 | 9082.38672 |

| | | | | | |
|---|---|---|---|---|---|
| *EPO1* | -0.012564394 | 0.93784429 | 0.99979869 | 11318.8948 | 11220.7455 |
| *BEM4* | -0.013025546 | 0.93859336 | 0.99979869 | 6135.01073 | 6079.86859 |
| *YIL089W* | -0.018279139 | 0.93860959 | 0.99979869 | 1050.62912 | 1037.40128 |
| *COY1* | -0.01210208 | 0.93903539 | 0.99979869 | 16073.5525 | 15939.2812 |
| *MSH4* | -0.017577662 | 0.93978294 | 0.99979869 | 1123.79271 | 1110.18339 |
| *CDC19* | -0.011384649 | 0.93997877 | 0.99979869 | 2073341.95 | 2057044.76 |
| *SUF9* | -0.10903824 | 0.94014105 | 0.99979869 | 16.5837466 | 15.3765012 |
| *snR68* | -0.020588308 | 0.94044038 | 0.99979869 | 662.37435 | 652.988752 |
| *AEP2* | -0.012613499 | 0.94059989 | 0.99979869 | 6060.87163 | 6008.11158 |
| *FAR10* | -0.012755396 | 0.94087987 | 0.99979869 | 5225.83121 | 5179.83071 |
| *PEX3* | -0.011852572 | 0.94103798 | 0.99979869 | 12420.2507 | 12318.6277 |
| *YCT1* | -0.012921688 | 0.94104525 | 0.99979869 | 4581.01612 | 4540.16826 |
| *PBN1* | -0.011818712 | 0.94127828 | 0.99979869 | 12147.1066 | 12048.0013 |
| *YNL122C* | -0.013396843 | 0.94139397 | 0.99979869 | 3395.766 | 3364.37847 |
| *PUP3* | -0.011232909 | 0.94181828 | 0.99979869 | 41608.6202 | 41285.9058 |
| *RRP5* | -0.011186006 | 0.94181908 | 0.99979869 | 54201.537 | 53782.9009 |
| *tF(GAA)P1* | -0.030650161 | 0.94191141 | 0.99979869 | 228.270394 | 223.471818 |
| *FAS2* | -0.01101889 | 0.94216567 | 0.99979869 | 154153.729 | 152980.811 |
| *DBP6* | -0.011688725 | 0.94217092 | 0.99979869 | 11277.9232 | 11186.9172 |
| *ADE3* | -0.011003558 | 0.94233811 | 0.99979869 | 116390.587 | 115506.227 |
| *IOC3* | -0.011495421 | 0.94268021 | 0.99979869 | 12943.1265 | 12840.4036 |
| *YER039C-A* | -0.021767679 | 0.94320041 | 0.99979869 | 498.487913 | 491.022939 |
| *LIP1* | -0.011340778 | 0.94374733 | 0.99979869 | 11760.8029 | 11668.7142 |
| *RPL17A* | -0.010697295 | 0.94378534 | 0.99979869 | 198565.978 | 197099.068 |
| *RPL7A* | -0.010654814 | 0.94392907 | 0.99979869 | 312805.555 | 310503.84 |
| *MSB1* | -0.011306348 | 0.94449182 | 0.99979869 | 9964.88068 | 9887.09029 |
| *SKP1* | -0.010623862 | 0.94489164 | 0.99979869 | 44798.5527 | 44469.8666 |
| *JJJ2* | -0.012394776 | 0.94521949 | 0.99979869 | 3625.0119 | 3594.00089 |
| *SEC59* | -0.011792809 | 0.94535777 | 0.99979869 | 5197.54129 | 5155.22831 |
| *RPL33B* | -0.010406728 | 0.94555681 | 0.99979869 | 90375.5659 | 89725.9851 |
| *YMR321C* | -0.036183275 | 0.94567349 | 0.99979869 | 135.596516 | 132.237911 |
| *PEX25* | -0.011173387 | 0.94588897 | 0.99979869 | 8246.99963 | 8183.37395 |
| *YER076C* | -0.012267242 | 0.94612554 | 0.99979869 | 3474.78267 | 3445.36137 |
| *PCL6* | -0.010827615 | 0.94701601 | 0.99979869 | 9476.14791 | 9405.29325 |
| *RSM7* | -0.010488702 | 0.94750919 | 0.99979869 | 13803.5303 | 13703.5379 |
| *YKL023W* | -0.010750463 | 0.94754414 | 0.99979869 | 9097.64828 | 9030.10662 |
| *YPL278C* | -0.023927069 | 0.94760615 | 0.99979869 | 319.968758 | 314.705725 |
| *ECM23* | -0.088931026 | 0.94834019 | 0.99979869 | 18.5347756 | 17.4267014 |
| *BUL2* | -0.010371951 | 0.94856113 | 0.99979869 | 11666.178 | 11582.6058 |
| *TPP1* | -0.012637519 | 0.94891515 | 0.99979869 | 2186.12801 | 2167.06157 |
| *SRB4* | -0.010466554 | 0.94916948 | 0.99979869 | 8527.94781 | 8466.30158 |
| *MCT1* | -0.010463658 | 0.94929019 | 0.99979869 | 8298.7019 | 8238.72936 |
| *SEC18* | -0.009904883 | 0.94937448 | 0.99979869 | 23355.7683 | 23195.9647 |
| *TFC4* | -0.010406523 | 0.94947337 | 0.99979869 | 8498.68238 | 8437.59877 |
| *CCT2* | -0.009679739 | 0.94953587 | 0.99979869 | 62556.8187 | 62138.4917 |

| | | | | | |
|---|---|---|---|---|---|
| *HPA3* | -0.010853144 | 0.94967261 | 0.99979869 | 5219.97812 | 5180.85581 |
| *PEX31* | -0.010549646 | 0.95047655 | 0.99979869 | 5838.45432 | 5795.91586 |
| *PGM1* | -0.00950475 | 0.95122399 | 0.99979869 | 26718.3668 | 26542.9164 |
| *YFL064C* | -0.031562983 | 0.95163899 | 0.99979869 | 141.449603 | 138.388511 |
| *GTT2* | -0.011592476 | 0.95183313 | 0.99979869 | 2509.02331 | 2488.943 |
| *ALA1* | -0.009182999 | 0.95198533 | 0.99979869 | 81344.2526 | 80828.1163 |
| *RPL42B* | -0.00911988 | 0.95204002 | 0.99979869 | 211293.515 | 209962.024 |
| *tH(GUG)G1* | -0.045825655 | 0.95215396 | 0.99979869 | 62.4329284 | 60.4809048 |
| *MFT1* | -0.009595592 | 0.95280838 | 0.99979869 | 10151.204 | 10083.9095 |
| *EXO84* | -0.009547928 | 0.95327181 | 0.99979869 | 9442.98042 | 9380.69085 |
| *BET1* | -0.010030902 | 0.95337064 | 0.99979869 | 5324.35817 | 5287.46622 |
| *YML100W-A* | -0.015150091 | 0.95348322 | 0.99979869 | 786.264692 | 778.050962 |
| *SVP26* | -0.0096776 | 0.95360747 | 0.99979869 | 7253.92587 | 7205.42847 |
| *YPR137C-B* | -0.017450547 | 0.95362474 | 0.99979869 | 521.900261 | 515.625341 |
| *SLM4* | -0.009825679 | 0.95412816 | 0.99979869 | 5533.11828 | 5495.56154 |
| *CAF4* | -0.009437601 | 0.95413193 | 0.99979869 | 8579.65008 | 8523.70718 |
| *RPP1* | -0.00935448 | 0.95434672 | 0.99979869 | 9071.30939 | 9012.67992 |
| *ALG6* | -0.008969708 | 0.95453696 | 0.99979869 | 18208.9538 | 18096.0917 |
| *TAP42* | -0.00963441 | 0.9554276 | 0.99979869 | 5099.98984 | 5066.0446 |
| *tW(CCA)K* | -0.07285656 | 0.95548516 | 0.99979869 | 20.4858046 | 19.4769016 |
| *LIP2* | -0.009390278 | 0.95581434 | 0.99979869 | 5955.51606 | 5916.87767 |
| *TRS65* | -0.009227006 | 0.95657202 | 0.99979869 | 5967.22223 | 5929.17887 |
| *RRP42* | -0.008753472 | 0.95670872 | 0.99979869 | 11047.7018 | 10980.8721 |
| *SRP21* | -0.008715202 | 0.95671537 | 0.99979869 | 11864.2074 | 11792.7513 |
| *SPO21* | -0.015103408 | 0.95685891 | 0.99979869 | 636.035458 | 629.41145 |
| *RPN5* | -0.008311894 | 0.95686752 | 0.99979869 | 44029.8472 | 43776.899 |
| *SUP2* | -0.053998706 | 0.9571922 | 0.99979869 | 35.1185222 | 33.8283027 |
| *YPR147C* | -0.008754709 | 0.95735747 | 0.99979869 | 8812.79805 | 8759.4802 |
| *GEA2* | -0.008379604 | 0.95741879 | 0.99979869 | 19407.8611 | 19295.4588 |
| *SGN1* | -0.008921718 | 0.9577802 | 0.99979869 | 6294.99511 | 6256.1858 |
| *RCE1* | -0.009424865 | 0.95925953 | 0.99979869 | 3128.47502 | 3108.10345 |
| *UGA4* | -0.012388862 | 0.95935291 | 0.99979869 | 966.734876 | 958.468576 |
| *GPI1* | -0.008521559 | 0.95959511 | 0.99979869 | 6415.95891 | 6378.17271 |
| *DUR3* | -0.009761608 | 0.95999029 | 0.99979869 | 2332.45518 | 2316.72618 |
| *YPL039W* | -0.008875848 | 0.96003584 | 0.99979869 | 4103.01401 | 4077.84813 |
| *YFL012W* | -0.026501537 | 0.96038483 | 0.99979869 | 133.645487 | 131.21281 |
| *YMR1* | -0.008327091 | 0.96044293 | 0.99979869 | 6539.84925 | 6502.20982 |
| *LYP1* | -0.007517075 | 0.96109652 | 0.99979869 | 37599.2556 | 37403.8518 |
| *ERP2* | -0.007655121 | 0.96110492 | 0.99979869 | 19251.7788 | 19149.8946 |
| *YPR196W* | -0.009318246 | 0.96205926 | 0.99979869 | 2255.38954 | 2240.86878 |
| *SNM1* | -0.008882208 | 0.96265711 | 0.99979869 | 2663.1546 | 2646.80841 |
| *TRP1* | -0.007529774 | 0.96302504 | 0.99979869 | 9841.96585 | 9790.73088 |
| *YCK1* | -0.007079195 | 0.96337817 | 0.99979869 | 36625.6921 | 36446.4083 |
| *VBA5* | -0.010000099 | 0.96340232 | 0.99979869 | 1420.34912 | 1410.53771 |
| *YJR061W* | -0.007842743 | 0.96366866 | 0.99979869 | 5139.01042 | 5111.14901 |

| | | | | | |
|---|---|---|---|---|---|
| *ABZ2* | -0.007374544 | 0.96381816 | 0.99979869 | 9714.17345 | 9664.64357 |
| *PRP16* | -0.007429908 | 0.96389774 | 0.99979869 | 8498.68238 | 8455.02547 |
| *YPR1* | -0.007087969 | 0.96392651 | 0.99979869 | 20085.8437 | 19987.4014 |
| *RRN9* | -0.007458579 | 0.96475123 | 0.99979869 | 6188.66403 | 6156.75109 |
| *SCM3* | -0.008421533 | 0.96482335 | 0.99979869 | 2569.50521 | 2554.5494 |
| *SNF12* | -0.007478156 | 0.96502131 | 0.99979869 | 5601.40429 | 5572.44404 |
| *PEP1* | -0.006749943 | 0.96507259 | 0.99979869 | 36960.2936 | 36787.7666 |
| *UFD1* | -0.00706744 | 0.96511211 | 0.99979869 | 10652.6184 | 10600.5599 |
| *BUD8* | -0.007188771 | 0.96520422 | 0.99979869 | 8083.1132 | 8042.93524 |
| *WSS1* | -0.007623088 | 0.96561598 | 0.99979869 | 4147.88768 | 4126.02783 |
| *YAE1* | -0.007742135 | 0.96628626 | 0.99979869 | 3266.02257 | 3248.54216 |
| *PRM9* | -0.007850758 | 0.96638032 | 0.99979869 | 2951.9069 | 2935.88663 |
| *ERV2* | -0.006747964 | 0.96660515 | 0.99979869 | 11082.8203 | 11031.102 |
| *YMR295C* | -0.006419172 | 0.96676524 | 0.99979869 | 37910.4447 | 37742.1348 |
| *IRC21* | -0.007372213 | 0.96722564 | 0.99979869 | 3739.1471 | 3720.0882 |
| *YPT10* | -0.006951777 | 0.96731431 | 0.99979869 | 5877.4749 | 5849.22107 |
| *SNU71* | -0.006599483 | 0.96745025 | 0.99979869 | 10492.634 | 10444.7447 |
| *GCD11* | -0.006163524 | 0.96775224 | 0.99979869 | 80640.9066 | 80297.1145 |
| *tF(GAA)M* | -0.038092566 | 0.96776879 | 0.99979869 | 39.9960948 | 38.9538031 |
| *TRK1* | -0.006349051 | 0.96779224 | 0.99979869 | 18322.1135 | 18241.656 |
| *EMC4* | -0.006392659 | 0.96825802 | 0.99979869 | 11707.1496 | 11655.3879 |
| *ERV14* | -0.006180082 | 0.96842345 | 0.99979869 | 22434.8826 | 22338.981 |
| *LCB3* | -0.006265039 | 0.96864284 | 0.99979869 | 13546.9699 | 13488.2669 |
| *CIN8* | -0.006444112 | 0.96877527 | 0.99979869 | 8176.76259 | 8140.31975 |
| *YPT32* | -0.006367719 | 0.96914264 | 0.99979869 | 8184.56671 | 8148.52055 |
| *MRPS35* | -0.006383779 | 0.96915111 | 0.99979869 | 7904.59404 | 7869.69333 |
| *IPL1* | -0.009181979 | 0.9691822 | 0.99979869 | 1039.89846 | 1033.30088 |
| *SCY1* | -0.006362994 | 0.96927445 | 0.99979869 | 7832.40597 | 7797.93632 |
| *MED1* | -0.006352457 | 0.96956371 | 0.99979869 | 7146.61927 | 7115.21967 |
| *APS2* | -0.006444492 | 0.96989438 | 0.99979869 | 5525.31416 | 5500.68704 |
| *PXA2* | -0.006919358 | 0.97063095 | 0.99979869 | 2801.67766 | 2788.27222 |
| *PBP2* | -0.006400745 | 0.97094725 | 0.99979869 | 4341.03955 | 4321.82194 |
| *UTP30* | -0.00620654 | 0.97097432 | 0.99979869 | 5578.96746 | 5555.01734 |
| *RKR1* | -0.00578797 | 0.9709761 | 0.99979869 | 14039.6048 | 13983.3902 |
| *TSA1* | -0.005491443 | 0.97105831 | 0.99979869 | 336036.457 | 334759.758 |
| *YER156C* | -0.005563153 | 0.97136685 | 0.99979869 | 28871.3273 | 28760.2079 |
| *EMP70* | -0.005510317 | 0.97150246 | 0.99979869 | 35569.2099 | 35433.6094 |
| *ARO8* | -0.005434826 | 0.97161251 | 0.99979869 | 67544.6244 | 67290.6447 |
| *KTR4* | -0.005503436 | 0.97162715 | 0.99979869 | 30881.8627 | 30764.2786 |
| *AIM24* | -0.005754836 | 0.97187499 | 0.99979869 | 9145.44849 | 9109.03933 |
| *ADH6* | -0.005283935 | 0.97239741 | 0.99979869 | 68106.5208 | 67857.525 |
| *AIM4* | -0.006972211 | 0.97268182 | 0.99979869 | 1878.84094 | 1869.78255 |
| *FPR4* | -0.005166725 | 0.97304794 | 0.99979869 | 60256.5555 | 60041.1369 |
| *OLA1* | -0.004959095 | 0.97395446 | 0.99979869 | 130976.48 | 130527.019 |
| *ROX1* | -0.005133763 | 0.97426202 | 0.99979869 | 13948.8819 | 13899.332 |

| | | | | | |
|---|---|---|---|---|---|
| *TSC13* | -0.004972524 | 0.97430666 | 0.99979869 | 33961.562 | 33844.7043 |
| *BMH2* | -0.004781051 | 0.97506226 | 0.99979869 | 59492.7277 | 59295.8891 |
| *DPS1* | -0.004625496 | 0.97572013 | 0.99979869 | 118520.135 | 118140.734 |
| *CKA1* | -0.00479315 | 0.97578961 | 0.99979869 | 16375.962 | 16321.6435 |
| *POL30* | -0.004939236 | 0.97587206 | 0.99979869 | 9070.33388 | 9039.33252 |
| *DCN1* | -0.00562004 | 0.97651723 | 0.99979869 | 2566.57867 | 2556.5996 |
| *MOB1* | -0.00463881 | 0.9770795 | 0.99979869 | 10736.5127 | 10702.0449 |
| *IAH1* | -0.0049499 | 0.97741264 | 0.99979869 | 4515.65665 | 4500.18936 |
| *NMA1* | -0.004432673 | 0.9777662 | 0.99979869 | 14055.213 | 14012.093 |
| *SEC12* | -0.004465671 | 0.97806377 | 0.99979869 | 9809.77387 | 9779.45478 |
| *KTR1* | -0.004162654 | 0.97844615 | 0.99979869 | 37369.0342 | 37261.3629 |
| *CDC26* | -0.005064417 | 0.97845903 | 0.99979869 | 2829.96758 | 2820.05033 |
| *tT(AGU)D* | -0.043942721 | 0.97848821 | 0.99979869 | 12.6816886 | 12.301201 |
| *YPR108W-A* | -0.01810624 | 0.97850992 | 0.99979869 | 80.967704 | 79.9578064 |
| *APC1* | -0.004424005 | 0.97863374 | 0.99979869 | 7817.77325 | 7793.83592 |
| *RPS0B* | -0.003934876 | 0.97927165 | 0.99979869 | 256896.868 | 256197.113 |
| *PRP22* | -0.004186017 | 0.9797457 | 0.99979869 | 7994.34138 | 7971.17824 |
| *APN2* | -0.004518244 | 0.97982775 | 0.99979869 | 3837.67407 | 3825.67351 |
| *RPT6* | -0.003846971 | 0.9799961 | 0.99979869 | 47390.4947 | 47264.2895 |
| *SAY1* | -0.004347782 | 0.98030464 | 0.99979869 | 4264.94942 | 4252.11514 |
| *BOP2* | -0.005463305 | 0.9807061 | 0.99979869 | 1238.90342 | 1234.2205 |
| *SLX4* | -0.004300525 | 0.98079154 | 0.99979869 | 3848.40473 | 3836.94961 |
| *RCF1* | -0.003939531 | 0.98110127 | 0.99979869 | 7215.8808 | 7196.20257 |
| *HEH2* | -0.003758416 | 0.98117873 | 0.99979869 | 13583.064 | 13547.7227 |
| *LAT1* | -0.003525428 | 0.98152098 | 0.99979869 | 95138.0277 | 94905.8158 |
| *SEO1* | -0.007361378 | 0.98162459 | 0.99979869 | 439.957042 | 437.717735 |
| *tE(UUC)C* | -0.019616266 | 0.98186671 | 0.99979869 | 47.8002108 | 47.1546038 |
| *YLR342W-A* | -0.005091472 | 0.98247086 | 0.99979869 | 1131.59683 | 1127.61009 |
| *RTC1* | -0.003536774 | 0.98265977 | 0.99979869 | 9523.94812 | 9500.62756 |
| *ANP1* | -0.003258295 | 0.98325997 | 0.99979869 | 26628.6195 | 26568.5439 |
| *RCO1* | -0.003421149 | 0.98369545 | 0.99979869 | 6705.68671 | 6689.80313 |
| *NAM2* | -0.003376293 | 0.98377324 | 0.99979869 | 7365.13452 | 7347.91739 |
| *ECM18* | -0.003947905 | 0.98456547 | 0.99979869 | 1856.40411 | 1851.33075 |
| *GDH2* | -0.002965316 | 0.98531607 | 0.99979869 | 11087.6979 | 11064.9303 |
| *YKR018C* | -0.002837005 | 0.98532237 | 0.99979869 | 35610.1815 | 35540.2198 |
| *SNX3* | -0.00301907 | 0.98581297 | 0.99979869 | 5811.13991 | 5798.99116 |
| *BGL2* | -0.002690756 | 0.98584563 | 0.99979869 | 168763.034 | 168448.546 |
| *DNL4* | -0.003002752 | 0.98607397 | 0.99979869 | 5153.64314 | 5142.92711 |
| *BOP3* | -0.002902359 | 0.98613663 | 0.99979869 | 6867.62212 | 6853.81915 |
| *ESC2* | -0.002910123 | 0.98626308 | 0.99979869 | 6071.60229 | 6059.36658 |
| *YFT2* | -0.002886342 | 0.9862644 | 0.99979869 | 6597.4046 | 6584.21783 |
| *SUP61* | -0.014198344 | 0.986485 | 0.99979869 | 50.7267543 | 50.229904 |
| *TOP1* | -0.002618567 | 0.98665289 | 0.99979869 | 20562.8703 | 20525.5789 |
| *YCK3* | -0.002631961 | 0.98708235 | 0.99979869 | 9656.6181 | 9639.01607 |
| *MDM36* | -0.002970479 | 0.987259 | 0.99979869 | 2965.5641 | 2959.46394 |

| | | | | | |
|---|---|---|---|---|---|
| *CDC10* | -0.002483583 | 0.98741778 | 0.99979869 | 17478.2934 | 17448.2285 |
| *CDA2* | -0.015930633 | 0.98762369 | 0.99979869 | 33.1674932 | 32.8032026 |
| *NSG2* | -0.002464424 | 0.9877696 | 0.99979869 | 11492.5364 | 11472.9201 |
| *ASI2* | -0.00252674 | 0.9879699 | 0.99979869 | 6628.62107 | 6617.02103 |
| *YOL036W* | -0.002335896 | 0.98827202 | 0.99979869 | 14267.8752 | 14244.7907 |
| *tS(AGA)H* | -0.021575542 | 0.98830664 | 0.99979869 | 15.6082321 | 15.3765012 |
| *RPL21A* | -0.002166783 | 0.98858344 | 0.99979869 | 259178.596 | 258789.591 |
| *ILM1* | -0.002306525 | 0.98873425 | 0.99979869 | 9013.75404 | 8999.35362 |
| *YLR173W* | -0.002321445 | 0.98890023 | 0.99979869 | 6939.8102 | 6928.65145 |
| *APL4* | -0.002154827 | 0.98906363 | 0.99979869 | 18279.1908 | 18251.907 |
| *RLM1* | -0.00206281 | 0.98955491 | 0.99979869 | 17242.2189 | 17217.581 |
| *UTR4* | -0.002069291 | 0.98964779 | 0.99979869 | 13278.7035 | 13259.6696 |
| *ANR2* | -0.002130309 | 0.98984095 | 0.99979869 | 6741.78075 | 6731.83224 |
| *RTT109* | -0.002207469 | 0.99046359 | 0.99979869 | 3093.3565 | 3088.62655 |
| *GDH3* | -0.002013147 | 0.9905099 | 0.99979869 | 5986.73252 | 5978.38368 |
| *MET1* | -0.00205333 | 0.9906982 | 0.99979869 | 4257.1453 | 4251.09004 |
| *GAS4* | -0.003758118 | 0.99093071 | 0.99979869 | 402.887491 | 401.839232 |
| *DNF3* | -0.001724865 | 0.99137665 | 0.99979869 | 13103.1108 | 13087.4527 |
| *YOR131C* | -0.001802498 | 0.99138912 | 0.99979869 | 6869.57315 | 6860.99485 |
| *DSF1* | -0.003758069 | 0.9914739 | 0.99979869 | 345.332135 | 344.433627 |
| *YPT53* | -0.002041886 | 0.9921473 | 0.99979869 | 1726.66068 | 1724.21834 |
| *OAF1* | -0.001581611 | 0.99216767 | 0.99979869 | 11075.0162 | 11062.8801 |
| *MRPL20* | -0.001522384 | 0.99262839 | 0.99979869 | 8041.16607 | 8032.68424 |
| *VPS54* | -0.001384627 | 0.99310635 | 0.99979869 | 12133.4494 | 12121.8085 |
| *ADE16* | -0.001234046 | 0.9935668 | 0.99979869 | 55553.6001 | 55506.0941 |
| *CTS2* | -0.00137506 | 0.99363421 | 0.99979869 | 5067.79786 | 5062.9693 |
| *HEL1* | -0.00134916 | 0.99374266 | 0.99979869 | 5148.76556 | 5143.95221 |
| *FZF1* | -0.001310411 | 0.99416355 | 0.99979869 | 3764.51048 | 3761.0922 |
| *SRP68* | -0.00109063 | 0.99440823 | 0.99979869 | 24621.0106 | 24602.402 |
| *NUP192* | -0.001084585 | 0.99445554 | 0.99979869 | 22377.3273 | 22360.5081 |
| *SEN54* | -0.001169464 | 0.99453383 | 0.99979869 | 5518.48556 | 5514.01334 |
| *ATG20* | -0.001040463 | 0.9948548 | 0.99979869 | 10817.4804 | 10809.6804 |
| *SPP382* | -0.00113775 | 0.99503818 | 0.99979869 | 3276.75323 | 3274.16966 |
| *CDC40* | -0.001010639 | 0.99525713 | 0.99979869 | 5729.19669 | 5725.18396 |
| *PDC1* | -0.000894219 | 0.99527544 | 0.99979869 | 2925849.93 | 2924036.48 |
| *DMA2* | -0.000884733 | 0.99576849 | 0.99979869 | 6954.44291 | 6950.17855 |
| *MRPL49* | -0.000847767 | 0.99598035 | 0.99979869 | 6339.86878 | 6336.14361 |
| *GAA1* | -0.000690857 | 0.99653717 | 0.99979869 | 13765.4852 | 13758.8933 |
| *RPS28A* | -0.000653199 | 0.99656509 | 0.99979869 | 153496.232 | 153426.729 |
| *FIP1* | -0.00068813 | 0.99671142 | 0.99979869 | 6892.9855 | 6889.69765 |
| *YOR012W* | -0.000766642 | 0.99688556 | 0.99979869 | 2215.39344 | 2214.21618 |
| *INP1* | -0.000688983 | 0.99710891 | 0.99979869 | 2617.30542 | 2616.05541 |
| *PSO2* | -0.000639677 | 0.99716037 | 0.99979869 | 3678.6652 | 3677.03399 |
| *PDR18* | -0.00057608 | 0.99734202 | 0.99979869 | 4922.4462 | 4920.48039 |
| *SLM2* | -0.000489072 | 0.99767172 | 0.99979869 | 6611.06181 | 6608.82023 |

| | | | | | |
|---|---|---|---|---|---|
| *NIF3* | -0.00042304 | 0.99783862 | 0.99979869 | 21976.3908 | 21969.945 |
| *SKI2* | -0.000205738 | 0.9989559 | 0.99999915 | 18200.1741 | 18197.5767 |
| *CAB4* | -1.32E-05 | 0.99993545 | 0.99999915 | 9214.71002 | 9214.62464 |
| *TRS20* | 7.18E-06 | 0.99996709 | 0.99999915 | 4659.05728 | 4659.07987 |
| *AVO1* | 1.77E-07 | 0.99999915 | 0.99999915 | 7038.33716 | 7038.33716 |

**Supplementary Table 1**

**Upregulated** (significant)

| ID | log2FoldChange | *p* value | padj | WT | *ykl075c*Δ |
|---|---|---|---|---|---|
| *INH1* | 1.184460297 | 0.01797334 | 0.99979869 | 2551.94595 | 5800.01626 |
| *TMA10* | 0.968830784 | 0.02247536 | 0.99979869 | 12642.668 | 24744.8909 |
| *YCR062W* | 0.989148038 | 0.02251539 | 0.99979869 | 7276.3627 | 14443.6602 |
| *CHA4* | 0.989410647 | 0.0232281 | 0.99979869 | 5562.38371 | 11043.4032 |
| *CYC7* | 1.029834879 | 0.02401514 | 0.99979869 | 2825.09001 | 5768.23816 |
| *FIG1* | 1.90718932 | 0.02476688 | 0.99979869 | 133.645487 | 501.27394 |
| *URC2* | 0.961183689 | 0.02493333 | 0.99979869 | 5179.98203 | 10084.9346 |
| *GPH1* | 0.909605359 | 0.02531816 | 0.99979869 | 15482.3907 | 29084.1395 |
| *STP4* | 0.886078595 | 0.02594774 | 0.99979869 | 33541.1153 | 61988.827 |
| *GIP2* | 1.013992581 | 0.02649721 | 0.99979869 | 2183.20146 | 4408.95545 |
| *YAT1* | 0.986516483 | 0.02657154 | 0.99979869 | 2727.53856 | 5404.32763 |
| *PLB3* | 0.880779365 | 0.02680435 | 0.99979869 | 18671.3476 | 34380.8316 |
| *RTN2* | 0.928456327 | 0.02748513 | 0.99979869 | 4417.12968 | 8406.84577 |
| *RTC3* | 0.867829138 | 0.02774935 | 0.99979869 | 17012.973 | 31047.2062 |
| *CYT1* | 0.882088021 | 0.02831601 | 0.99979869 | 8026.53336 | 14793.2193 |
| *FMP48* | 0.878571708 | 0.02844585 | 0.99979869 | 8333.82043 | 15322.1709 |
| *YRO2* | 0.83800556 | 0.02888159 | 0.99979869 | 43920.5896 | 78511.3902 |
| *PHM7* | 1.065101634 | 0.02899198 | 0.99979869 | 1206.71144 | 2524.8215 |
| *FRE7* | 0.857294229 | 0.02998617 | 0.99979869 | 8087.01526 | 14650.7304 |
| *CYC1* | 0.903234482 | 0.03025057 | 0.99979869 | 3561.60346 | 6661.10033 |
| *YGL081W* | 1.135660891 | 0.0310327 | 0.99979869 | 751.14617 | 1650.41113 |
| *UGA1* | 0.809162078 | 0.03230584 | 0.99979869 | 13942.0533 | 24429.16 |
| *OM14* | 0.802837145 | 0.0339299 | 0.99979869 | 8921.08016 | 15563.0694 |
| *YCL042W* | 0.847204172 | 0.03429647 | 0.99979869 | 3702.07755 | 6660.07523 |
| *IMD2* | 0.779023023 | 0.03441463 | 0.99979869 | 17501.7058 | 30032.3571 |
| *SDH1* | 0.788717247 | 0.03516419 | 0.99979869 | 8926.93325 | 15421.6056 |
| *SPI1* | 0.757191104 | 0.03570701 | 0.99979869 | 28854.7436 | 48770.1615 |
| *TPO2* | 0.747992559 | 0.03620353 | 0.99979869 | 42972.3895 | 72170.1211 |
| *tD(GUC)N* | 1.363221566 | 0.03789327 | 0.99979869 | 263.388917 | 677.591154 |
| *YNL190W* | 0.732574419 | 0.0379108 | 0.99979869 | 33723.5365 | 56035.0458 |
| *PAN1* | 0.725855711 | 0.0384879 | 0.99979869 | 37466.5856 | 61965.2497 |
| *SSA4* | 0.730736723 | 0.03869501 | 0.99979869 | 21015.509 | 34874.9299 |
| *USV1* | 0.757145726 | 0.03880483 | 0.99979869 | 7443.17568 | 12580.0282 |
| *POR1* | 0.715918058 | 0.03894178 | 0.99979869 | 99452.7284 | 163353.798 |
| *YNL194C* | 0.792623103 | 0.04056881 | 0.99979869 | 3138.23017 | 5436.10573 |
| *YNR014W* | 0.717062666 | 0.04160516 | 0.99979869 | 11110.1347 | 18263.1831 |

| | | | | | |
|---|---|---|---|---|---|
| *RPL15A* | 0.682808339 | 0.04232428 | 0.99979869 | 450649.657 | 723410.052 |
| *SED1* | 0.67929313 | 0.04278159 | 0.99979869 | 311445.688 | 498734.767 |
| *TSL1* | 0.679495673 | 0.04296954 | 0.99979869 | 116051.108 | 185864.996 |
| *COX26* | 1.05777735 | 0.04493196 | 0.99979869 | 514.096145 | 1070.20449 |
| *ASG7* | 0.770986038 | 0.04511263 | 0.99979869 | 2531.46014 | 4319.77174 |
| *SRO9* | 0.700293758 | 0.04602777 | 0.99979869 | 6329.13812 | 10283.804 |
| *MSC1* | 0.658682729 | 0.04751023 | 0.99979869 | 18194.3211 | 28722.2792 |
| *CRC1* | 1.307598045 | 0.04798266 | 0.99979869 | 219.490764 | 543.303043 |
| *GAD1* | 0.647803909 | 0.04818775 | 0.99979869 | 28268.4594 | 44290.4741 |
| *PNS1* | 0.652196787 | 0.04910277 | 0.99979869 | 13998.6332 | 21999.6729 |
| *FMP45* | 0.775035194 | 0.05020687 | 0.99979869 | 1693.49318 | 2897.95793 |
| *EGO4* | 0.644320738 | 0.05086433 | 0.99979869 | 11572.5286 | 18087.8909 |
| *CWP2* | 0.61659139 | 0.05135159 | 0.99979869 | 267695.813 | 410443.922 |
| *PIL1* | 0.612425227 | 0.05220235 | 0.99979869 | 119860.492 | 183245.866 |
| *FMP16* | 0.757443732 | 0.05226614 | 0.99979869 | 1722.75862 | 2912.30933 |
| *SNT1* | 0.62065099 | 0.05283237 | 0.99979869 | 20355.0857 | 31297.3306 |
| *DCS2* | 0.652835654 | 0.0532434 | 0.99979869 | 5779.92345 | 9087.51223 |
| *BAT2* | 0.606446154 | 0.05353892 | 0.99979869 | 62383.1772 | 94978.5979 |
| *LSP1* | 0.599977938 | 0.05435797 | 0.99979869 | 91247.6759 | 138303.428 |
| *HER1* | 0.62165402 | 0.05469684 | 0.99979869 | 10780.4108 | 16587.1444 |
| *COG3* | 0.626091189 | 0.05503953 | 0.99979869 | 8607.94 | 13285.2971 |
| *YDR222W* | 0.633736747 | 0.05573187 | 0.99979869 | 6233.53769 | 9671.81927 |
| *TIR3* | 0.6528796 | 0.05666502 | 0.99979869 | 3900.107 | 6132.14869 |
| *ICY1* | 0.625965804 | 0.0577662 | 0.99979869 | 5668.71479 | 8748.2041 |
| *QCR7* | 0.59766639 | 0.0577801 | 0.99979869 | 13916.6899 | 21059.6561 |
| *XBP1* | 0.638475388 | 0.05779612 | 0.99979869 | 4427.86034 | 6892.77295 |
| *AGA2* | 0.610866193 | 0.05813654 | 0.99979869 | 7854.8428 | 11995.7212 |
| *YML122C* | 1.144559814 | 0.05937945 | 0.99979869 | 253.633772 | 560.729745 |
| *ALD3* | 0.685849912 | 0.05940049 | 0.99979869 | 2135.40125 | 3435.11037 |
| *YOR376W* | 1.581391802 | 0.05952734 | 0.99979869 | 89.7473346 | 268.576221 |
| *TFS1* | 0.587450226 | 0.06027678 | 0.99979869 | 11906.1545 | 17890.0466 |
| *PML39* | 0.630067053 | 0.06113185 | 0.99979869 | 3726.46541 | 5767.21306 |
| *YIL082W-A* | 0.579749615 | 0.06244533 | 0.99979869 | 10300.4577 | 15394.953 |
| *PGM2* | 0.551221741 | 0.06425425 | 0.99979869 | 40183.3935 | 58881.7487 |
| *YHR202W* | 0.612804346 | 0.06520884 | 0.99979869 | 3488.43987 | 5334.62083 |
| *DED1* | 0.538924171 | 0.06731384 | 0.99979869 | 31391.0813 | 45607.7277 |
| *AMA1* | 1.046263784 | 0.06765927 | 0.99979869 | 280.948178 | 580.206646 |
| *GDB1* | 0.566227044 | 0.06801471 | 0.99979869 | 6701.78466 | 9922.96879 |
| *TOS8* | 0.609186901 | 0.06812453 | 0.99979869 | 2993.85402 | 4566.82086 |
| *LSC2* | 0.538788735 | 0.0682687 | 0.99979869 | 21225.2446 | 30835.0105 |
| *YIR018C-A* | 0.783560246 | 0.06840237 | 0.99979869 | 762.852344 | 1313.1532 |
| *SDH2* | 0.558408491 | 0.06849569 | 0.99979869 | 7890.93684 | 11620.5345 |
| *GUT2* | 0.556094223 | 0.06858507 | 0.99979869 | 8392.3513 | 12339.1297 |
| *OM45* | 0.531843184 | 0.07033918 | 0.99979869 | 18283.0929 | 26433.2307 |
| *VPS73* | 0.572307512 | 0.0718791 | 0.99979869 | 4038.63006 | 6005.03628 |

| | | | | | |
|---|---|---|---|---|---|
| *YMR196W* | 0.535744043 | 0.07198272 | 0.99979869 | 10282.8984 | 14907.0054 |
| *STF2* | 0.51510401 | 0.07212593 | 0.99979869 | 51774.4569 | 73990.6988 |
| *UIP4* | 0.60265018 | 0.07254769 | 0.99979869 | 2503.17022 | 3801.0711 |
| *DDR2* | 0.554922857 | 0.07284985 | 0.99979869 | 5242.41496 | 7701.57691 |
| *UGP1* | 0.507138093 | 0.0736388 | 0.99979869 | 80616.5188 | 114574.411 |
| *HSP42* | 0.503128823 | 0.07450598 | 0.99979869 | 100716.02 | 142743.136 |
| *MFA1* | 0.509702093 | 0.07452699 | 0.99979869 | 28318.2106 | 40318.2113 |
| *REC8* | 0.989068347 | 0.07489692 | 0.99979869 | 289.727808 | 575.081146 |
| *RPM2* | 0.537335726 | 0.0750163 | 0.99979869 | 6489.12249 | 9417.59445 |
| *SCW4* | 0.500692316 | 0.07505412 | 0.99979869 | 116008.185 | 164139.025 |
| *IGD1* | 0.518033329 | 0.07545476 | 0.99979869 | 12177.3476 | 17437.9775 |
| *COR1* | 0.508406965 | 0.07577687 | 0.99979869 | 20449.7106 | 29089.265 |
| *YJR115W* | 0.559979658 | 0.07645334 | 0.99979869 | 3587.94235 | 5289.51642 |
| *GIS1* | 0.518791526 | 0.07746502 | 0.99979869 | 8551.36016 | 12251.9962 |
| *YAP5* | 0.54701554 | 0.07806699 | 0.99979869 | 4018.14425 | 5870.74817 |
| *MTL1* | 0.516020233 | 0.0783598 | 0.99979869 | 8383.57167 | 11988.5455 |
| *HXT6* | 0.814540365 | 0.07849048 | 0.99979869 | 511.169601 | 899.012772 |
| *YLR297W* | 0.538019087 | 0.07895957 | 0.99979869 | 4466.88092 | 6485.80822 |
| *QCR2* | 0.496064383 | 0.07970235 | 0.99979869 | 17392.4481 | 24529.6199 |
| *ACH1* | 0.506288637 | 0.0797246 | 0.99979869 | 10046.8239 | 14270.4182 |
| *VID24* | 0.515660104 | 0.07973312 | 0.99979869 | 7195.395 | 10286.8793 |
| *TMA17* | 0.505912714 | 0.08001655 | 0.99979869 | 9773.67984 | 13878.83 |
| *ECM21* | 0.500786372 | 0.08103495 | 0.99979869 | 10415.5684 | 14737.8639 |
| *QCR9* | 0.520091143 | 0.08135594 | 0.99979869 | 5446.29749 | 7810.23752 |
| *REE1* | 0.484739164 | 0.08140512 | 0.99979869 | 27920.2007 | 39069.6394 |
| *GSY2* | 0.478546628 | 0.08187033 | 0.99979869 | 57227.583 | 79737.4099 |
| *BTN2* | 0.473042863 | 0.08357425 | 0.99979869 | 55410.1995 | 76911.2089 |
| *YPL247C* | 0.480468706 | 0.08398662 | 0.99979869 | 18673.2987 | 26052.9186 |
| *COX4* | 0.50420627 | 0.08406945 | 0.99979869 | 6443.27331 | 9138.76723 |
| *MAL31* | 0.522813007 | 0.08420393 | 0.99979869 | 4141.05908 | 5949.68087 |
| *YHR219W* | 1.027860097 | 0.08531419 | 0.99979869 | 212.662162 | 433.617335 |
| *BNR1* | 0.50661261 | 0.08555832 | 0.99979869 | 5298.9948 | 7528.335 |
| *HOP1* | 0.635580566 | 0.0863209 | 0.99979869 | 1107.20896 | 1720.11794 |
| *DBP1* | 0.563959344 | 0.08633859 | 0.99979869 | 2062.23767 | 3048.64764 |
| *GRE3* | 0.47102294 | 0.08717148 | 0.99979869 | 17636.3268 | 24445.5617 |
| *COS8* | 0.484586882 | 0.08718443 | 0.99979869 | 8710.36903 | 12187.4149 |
| *MFA2* | 0.461125578 | 0.08759226 | 0.99979869 | 47170.0284 | 64934.9647 |
| *RPL15B* | 0.480678963 | 0.08765493 | 0.99979869 | 9579.55245 | 13367.3051 |
| *HSP82* | 0.457255309 | 0.08773524 | 0.99979869 | 138025.547 | 189498.976 |
| *YJR149W* | 0.56441699 | 0.08783875 | 0.99979869 | 1935.42078 | 2862.07943 |
| *AIM17* | 0.460693262 | 0.08794242 | 0.99979869 | 41902.2501 | 57665.98 |
| *YPC1* | 0.489075562 | 0.08910774 | 0.99979869 | 6167.20271 | 8655.94509 |
| *NAT4* | 0.637050948 | 0.08986196 | 0.99979869 | 996.000311 | 1548.92622 |
| *PHO87* | 0.469320621 | 0.09003144 | 0.99979869 | 11451.5648 | 15854.1979 |
| *PRX1* | 0.472166099 | 0.09005791 | 0.99979869 | 9993.1706 | 13862.4284 |

| | | | | | |
|---|---|---|---|---|---|
| *VRP1* | 0.458982424 | 0.09012194 | 0.99979869 | 22560.724 | 31011.3277 |
| *SDH4* | 0.474889595 | 0.09014482 | 0.99979869 | 8833.28385 | 12276.5986 |
| *SOL4* | 0.472429501 | 0.09031508 | 0.99979869 | 9553.21356 | 13254.5441 |
| *HSP26* | 0.464516393 | 0.09059533 | 0.99979869 | 13521.6066 | 18657.8466 |
| *YJL218W* | 0.557710459 | 0.09071796 | 0.99979869 | 1867.13476 | 2748.29332 |
| *MGA1* | 0.462814458 | 0.09093386 | 0.99979869 | 14083.5029 | 19410.27 |
| *MUP3* | 0.488590078 | 0.09131033 | 0.99979869 | 5215.10055 | 7317.16438 |
| *NCA3* | 0.510231842 | 0.09182706 | 0.99979869 | 3288.4594 | 4683.68227 |
| *YBR238C* | 0.471387953 | 0.09199336 | 0.99979869 | 8155.30127 | 11306.8539 |
| *YLR149C* | 0.495393442 | 0.0925237 | 0.99979869 | 4133.25496 | 5826.66886 |
| *QCR10* | 0.497836533 | 0.09261597 | 0.99979869 | 3918.64177 | 5533.49024 |
| *GSM1* | 0.533663342 | 0.09267646 | 0.99979869 | 2270.99777 | 3287.49596 |
| *SDS24* | 0.448474492 | 0.09321498 | 0.99979869 | 26195.491 | 35746.265 |
| *KIN82* | 0.48021677 | 0.09363216 | 0.99979869 | 5385.81559 | 7512.9585 |
| *TIP1* | 0.437353864 | 0.09502693 | 0.99979869 | 79402.0032 | 107519.672 |
| *NCE102* | 0.436241642 | 0.09530985 | 0.99979869 | 89457.6068 | 121042.793 |
| *SCW10* | 0.43614637 | 0.09601376 | 0.99979869 | 52921.662 | 71602.2156 |
| *FIG2* | 0.464422975 | 0.09607196 | 0.99979869 | 6868.59764 | 9477.05026 |
| *PHO3* | 0.435500794 | 0.09626668 | 0.99979869 | 52446.5864 | 70927.6998 |
| *RGI1* | 0.440305596 | 0.09628544 | 0.99979869 | 25069.7473 | 34016.9211 |
| *CLN3* | 0.443397751 | 0.09779659 | 0.99979869 | 13931.3227 | 18943.8495 |
| *CIK1* | 0.475168328 | 0.09798061 | 0.99979869 | 4493.21982 | 6245.9348 |
| *ALD6* | 0.426419742 | 0.09852514 | 0.99979869 | 158925.946 | 213579.602 |
| *PHO92* | 1.385639379 | 0.09888882 | 0.99979869 | 72.1880735 | 188.618415 |
| *KNS1* | 0.440287601 | 0.09936463 | 0.99979869 | 13063.1148 | 17725.0055 |
| *OPI1* | 0.440070393 | 0.10006685 | 0.99979869 | 11918.8362 | 16169.9287 |
| *COX7* | 0.458326676 | 0.10022036 | 0.99979869 | 5850.16049 | 8037.80974 |
| *GOR1* | 0.447323123 | 0.10064151 | 0.99979869 | 8051.89673 | 10978.8219 |
| *YNL195C* | 0.525140606 | 0.10078722 | 0.99979869 | 1872.01234 | 2693.96301 |
| *YPR010C-A* | 0.477123598 | 0.10088091 | 0.99979869 | 3663.05697 | 5098.84781 |
| *HCM1* | 0.466981665 | 0.10095747 | 0.99979869 | 4485.4157 | 6199.80529 |
| *NQM1* | 0.537972136 | 0.1011573 | 0.99979869 | 1620.32959 | 2352.60469 |
| *CUR1* | 0.446992646 | 0.10140733 | 0.99979869 | 7570.96808 | 10320.7076 |
| *STF1* | 0.455285954 | 0.10168771 | 0.99979869 | 5707.73538 | 7825.61402 |
| *PBP1* | 0.4224992 | 0.10182802 | 0.99979869 | 38936.686 | 52184.7699 |
| *PRY2* | 0.425587495 | 0.10206395 | 0.99979869 | 23859.1338 | 32045.6537 |
| *YJL144W* | 0.430690988 | 0.10276987 | 0.99979869 | 13731.3422 | 18508.182 |
| *AQY3* | 0.483604237 | 0.10294006 | 0.99979869 | 2971.41719 | 4154.73063 |
| *MED2* | 0.429956195 | 0.10388024 | 0.99979869 | 12168.5679 | 16393.4005 |
| *YJR146W* | 0.528748127 | 0.10555522 | 0.99979869 | 1561.79872 | 2253.16998 |
| *RIP1* | 0.438867722 | 0.10568423 | 0.99979869 | 6876.40175 | 9321.23504 |
| *ISF1* | 0.47364086 | 0.10595318 | 0.99979869 | 3062.14003 | 4252.11514 |
| *YHR054C* | 0.965776948 | 0.10659729 | 0.99979869 | 186.323271 | 363.910529 |
| *MSA1* | 0.429861798 | 0.10670048 | 0.99979869 | 8757.19372 | 11796.8517 |
| *KGD1* | 0.417754053 | 0.10679527 | 0.99979869 | 17048.0915 | 22773.6234 |

| | | | | | |
|---|---|---|---|---|---|
| *MIG2* | 0.428752788 | 0.10715526 | 0.99979869 | 8760.12026 | 11791.7262 |
| *PCL1* | 0.470878118 | 0.10767889 | 0.99979869 | 2976.29476 | 4125.00273 |
| *BMS1* | 0.406410022 | 0.10816771 | 0.99979869 | 46103.7911 | 61105.1908 |
| *GSY1* | 0.402734409 | 0.10970172 | 0.99979869 | 47969.9503 | 63416.7915 |
| *FUS1* | 0.444485716 | 0.10976618 | 0.99979869 | 4470.78298 | 6083.96899 |
| *KSP1* | 0.40511777 | 0.11026397 | 0.99979869 | 26688.1259 | 35340.3253 |
| *PRM1* | 0.465410012 | 0.11033535 | 0.99979869 | 2914.83734 | 4024.54292 |
| *PRT1* | 0.398697779 | 0.11091283 | 0.99979869 | 67510.4814 | 89000.2142 |
| *RPL10* | 0.395243264 | 0.11112807 | 0.99979869 | 722531.403 | 950247.274 |
| *IRC7* | 0.401658293 | 0.11202833 | 0.99979869 | 25225.8296 | 33323.9535 |
| *COX8* | 0.421944027 | 0.11205356 | 0.99979869 | 7298.79953 | 9778.42968 |
| *PTR2* | 0.403319447 | 0.11322376 | 0.99979869 | 16754.4616 | 22158.5634 |
| *DHH1* | 0.40413329 | 0.11361683 | 0.99979869 | 14789.7754 | 19571.2108 |
| *NDI1* | 0.404155876 | 0.11363192 | 0.99979869 | 14732.2201 | 19495.3534 |
| *PNC1* | 0.391298345 | 0.11428505 | 0.99979869 | 69516.1392 | 91175.4766 |
| *PRB1* | 0.391529667 | 0.11437266 | 0.99979869 | 61344.2542 | 80470.3564 |
| *YBR241C* | 0.424051595 | 0.11461512 | 0.99979869 | 5653.10656 | 7584.71551 |
| *SSA2* | 0.386525948 | 0.11528156 | 0.99979869 | 468879.096 | 612938.092 |
| *OSW5* | 0.437619713 | 0.11575921 | 0.99979869 | 3816.21275 | 5168.55461 |
| *FLO11* | 0.505610959 | 0.11594795 | 0.99979869 | 1486.68411 | 2110.68107 |
| *YMR122W-A* | 0.38427177 | 0.11688921 | 0.99979869 | 141979.308 | 185311.442 |
| *YHL050C* | 0.421639203 | 0.11706115 | 0.99979869 | 5192.66372 | 6955.30405 |
| *ATP2* | 0.383359465 | 0.11789919 | 0.99979869 | 80889.6628 | 105510.476 |
| *FAR1* | 0.394829577 | 0.11799335 | 0.99979869 | 14769.2896 | 19418.4708 |
| *HAP4* | 0.384113664 | 0.11892009 | 0.99979869 | 39104.4745 | 51033.5825 |
| *YJR096W* | 0.399860091 | 0.12014661 | 0.99979869 | 8729.87932 | 11518.0245 |
| *CCR4* | 0.387677011 | 0.12053978 | 0.99979869 | 17172.9574 | 22467.1185 |
| *RPG1* | 0.377763711 | 0.12067345 | 0.99979869 | 82570.4743 | 107285.949 |
| *TPS2* | 0.376618566 | 0.12089411 | 0.99979869 | 113990.821 | 147993.699 |
| *YKR075C* | 0.408912418 | 0.12092592 | 0.99979869 | 5905.76482 | 7840.99053 |
| *FUN19* | 0.407280482 | 0.12116402 | 0.99979869 | 6120.37801 | 8116.74245 |
| *RSP5* | 0.382151732 | 0.12139788 | 0.99979869 | 25142.9109 | 32768.3492 |
| *YPK9* | 0.387695888 | 0.12147195 | 0.99979869 | 14830.747 | 19403.0943 |
| *CDC6* | 0.489811093 | 0.12152183 | 0.99979869 | 1516.92506 | 2130.15797 |
| *ATP1* | 0.376458288 | 0.12157645 | 0.99979869 | 70001.9454 | 90873.0721 |
| *YOX1* | 0.444295818 | 0.12278969 | 0.99979869 | 2567.55418 | 3493.54108 |
| *YBR284W* | 0.484063821 | 0.12313816 | 0.99979869 | 1547.16601 | 2163.98627 |
| *SWH1* | 0.378298075 | 0.12360689 | 0.99979869 | 23834.7459 | 30980.5747 |
| *SFK1* | 0.383471302 | 0.12364891 | 0.99979869 | 14763.4365 | 19258.5552 |
| *UBI4* | 0.374746518 | 0.12370115 | 0.99979869 | 38971.8045 | 50531.2834 |
| *DOT6* | 0.372366162 | 0.12439459 | 0.99979869 | 48649.8839 | 62975.9984 |
| *UTH1* | 0.367747924 | 0.12501913 | 0.99979869 | 244686.353 | 315727.75 |
| *AGP2* | 0.40313269 | 0.12533473 | 0.99979869 | 5364.35427 | 7093.69257 |
| *NUP60* | 0.377744916 | 0.12548878 | 0.99979869 | 17578.7714 | 22840.2549 |
| *RIB4* | 0.370995267 | 0.12625939 | 0.99979869 | 32417.3226 | 41923.518 |

| | | | | | |
|---|---|---|---|---|---|
| *AZF1* | 0.398487081 | 0.1266525 | 0.99979869 | 5698.95574 | 7511.9334 |
| *HOS3* | 0.394101084 | 0.12667735 | 0.99979869 | 6564.23711 | 8626.21719 |
| *NNR1* | 0.389840029 | 0.12733554 | 0.99979869 | 7322.21188 | 9593.91167 |
| *PST1* | 0.366107752 | 0.12755092 | 0.99979869 | 53619.1548 | 69108.1471 |
| *PYC1* | 0.372499066 | 0.12782372 | 0.99979869 | 19033.2635 | 24640.3307 |
| *OST3* | 0.371390613 | 0.12900991 | 0.99979869 | 17274.4109 | 22346.1567 |
| *YBR137W* | 0.393116519 | 0.12922149 | 0.99979869 | 5770.1683 | 7577.5398 |
| *YGL036W* | 0.373703871 | 0.12979471 | 0.99979869 | 13012.388 | 16859.821 |
| *OPT1* | 0.368928512 | 0.13031113 | 0.99979869 | 17428.5422 | 22507.0974 |
| *DCS1* | 0.375101952 | 0.13059562 | 0.99979869 | 10925.7625 | 14169.9584 |
| *YIA6* | 0.442588543 | 0.13109771 | 0.99979869 | 2038.82532 | 2770.84552 |
| *MDH2* | 0.366898997 | 0.13123846 | 0.99979869 | 18011.8998 | 23227.7428 |
| *MDG1* | 0.370431512 | 0.13138884 | 0.99979869 | 13356.7446 | 17266.7858 |
| *HSP104* | 0.355752472 | 0.13169116 | 0.99979869 | 243854.239 | 312048.666 |
| *PAC1* | 0.413461834 | 0.1317321 | 0.99979869 | 3144.08325 | 4187.53383 |
| *ATP7* | 0.369085218 | 0.13204255 | 0.99979869 | 13521.6066 | 17463.605 |
| *ATP19* | 0.380391113 | 0.13359196 | 0.99979869 | 6664.71511 | 8675.42199 |
| *HXT11* | 0.521808955 | 0.13386881 | 0.99979869 | 897.473346 | 1288.5508 |
| *SNF2* | 0.357508105 | 0.1342623 | 0.99979869 | 28753.2901 | 36839.0216 |
| *VNX1* | 0.365647265 | 0.13449234 | 0.99979869 | 12467.0754 | 16063.3183 |
| *YDL124W* | 0.352919383 | 0.13484036 | 0.99979869 | 59858.5456 | 76447.8637 |
| *MCR1* | 0.357176104 | 0.13542135 | 0.99979869 | 22334.4046 | 28608.4931 |
| *YAP6* | 0.460043137 | 0.13548798 | 0.99979869 | 1499.3658 | 2062.50136 |
| *SPO11* | 0.569682031 | 0.13598139 | 0.99979869 | 629.206856 | 933.866175 |
| *GLK1* | 0.348907979 | 0.13602562 | 0.99979869 | 151133.536 | 192483.042 |
| *AFR1* | 0.375394613 | 0.13718945 | 0.99979869 | 6366.20767 | 8258.20626 |
| *YOL164W-A* | 1.038056001 | 0.13743058 | 0.99979869 | 106.331081 | 218.346317 |
| *YGR130C* | 0.351023422 | 0.13754754 | 0.99979869 | 31939.3204 | 40737.4773 |
| *YER079W* | 0.371574025 | 0.13771691 | 0.99979869 | 7089.06392 | 9171.57043 |
| *AGP3* | 0.456729374 | 0.13850109 | 0.99979869 | 1453.51661 | 1994.84476 |
| *FAT3* | 0.399938471 | 0.13901292 | 0.99979869 | 3163.59354 | 4174.20753 |
| *PFK27* | 0.361661242 | 0.13906355 | 0.99979869 | 10033.1667 | 12891.6586 |
| *FRT2* | 0.382994065 | 0.13976077 | 0.99979869 | 4481.51364 | 5844.09557 |
| *PRM6* | 1.332295068 | 0.1399122 | 0.99979869 | 51.7022688 | 130.18771 |
| *EIS1* | 0.345691234 | 0.1401287 | 0.99979869 | 39604.9134 | 50328.3136 |
| *MDH1* | 0.345768646 | 0.14055367 | 0.99979869 | 34387.8619 | 43701.0416 |
| *YGR067C* | 0.513070175 | 0.14152948 | 0.99979869 | 841.869019 | 1201.4173 |
| *TOS3* | 0.371581759 | 0.1426723 | 0.99979869 | 5367.28081 | 6944.02795 |
| *SRL1* | 0.339465589 | 0.1427777 | 0.99979869 | 68606.9597 | 86807.5251 |
| *FPK1* | 0.360392498 | 0.14330847 | 0.99979869 | 7749.48724 | 9948.59629 |
| *OPI10* | 0.353654259 | 0.14338534 | 0.99979869 | 10790.166 | 13787.5961 |
| *GLG1* | 0.362391346 | 0.14342239 | 0.99979869 | 7079.30877 | 9100.83853 |
| *YOR29-13* | 0.383442658 | 0.14370067 | 0.99979869 | 3750.85328 | 4892.80269 |
| *RIE1* | 0.340355924 | 0.14417181 | 0.99979869 | 35043.4076 | 44367.3566 |
| *EMI1* | 0.368330605 | 0.14440747 | 0.99979869 | 5423.86065 | 7001.43356 |

| | | | | | |
|---|---|---|---|---|---|
| *PUF3* | 0.356482542 | 0.14494183 | 0.99979869 | 8188.46876 | 10483.6985 |
| *YNL144C* | 0.353358381 | 0.14502578 | 0.99979869 | 9473.22137 | 12102.3316 |
| *YOL163W* | 0.712946121 | 0.14587017 | 0.99979869 | 277.04612 | 454.119336 |
| *HSP150* | 0.332504137 | 0.14598081 | 0.99979869 | 428038.206 | 538984.297 |
| *UME6* | 0.348238094 | 0.14610779 | 0.99979869 | 11507.1691 | 14648.6802 |
| *GRX2* | 0.344181911 | 0.14613151 | 0.99979869 | 15362.4024 | 19501.504 |
| *HFA1* | 0.347627008 | 0.14698593 | 0.99979869 | 10975.5137 | 13965.9635 |
| *CTT1* | 0.354296244 | 0.14737818 | 0.99979869 | 7591.45389 | 9704.62247 |
| *YLR177W* | 0.337632223 | 0.14748083 | 0.99979869 | 24287.3847 | 30691.4964 |
| *YDR215C* | 0.542762158 | 0.14793638 | 0.99979869 | 626.280313 | 912.339073 |
| *REC104* | 0.647291036 | 0.14815762 | 0.99979869 | 358.013824 | 560.729745 |
| *ECM29* | 0.338353532 | 0.1488692 | 0.99979869 | 17742.6578 | 22432.2651 |
| *PIR3* | 0.404523085 | 0.14889621 | 0.99979869 | 2179.29941 | 2884.63163 |
| *SPO12* | 0.344139895 | 0.14941904 | 0.99979869 | 10693.59 | 13574.3753 |
| *YGR127W* | 0.364743664 | 0.14983031 | 0.99979869 | 4653.20419 | 5991.70998 |
| *BNS1* | 0.434100057 | 0.1500638 | 0.99979869 | 1461.32073 | 1974.34276 |
| *SST2* | 0.336065537 | 0.1507973 | 0.99979869 | 16553.5057 | 20895.6401 |
| *SEC31* | 0.328187106 | 0.15130779 | 0.99979869 | 45156.5665 | 56691.1098 |
| *YMR206W* | 0.530166633 | 0.15174149 | 0.99979869 | 643.839574 | 929.765774 |
| *PHO23* | 0.345519798 | 0.15199469 | 0.99979869 | 8190.41979 | 10406.816 |
| *SNX4* | 0.34182858 | 0.15247691 | 0.99979869 | 9501.51129 | 12041.8507 |
| *HSP30* | 0.322718949 | 0.15323058 | 0.99979869 | 133118.71 | 166489.58 |
| *MMF1* | 0.322094679 | 0.15437568 | 0.99979869 | 79554.1835 | 99454.1848 |
| *SDP1* | 0.357699647 | 0.1546213 | 0.99979869 | 4606.3795 | 5902.52627 |
| *MHP1* | 0.326583655 | 0.1549272 | 0.99979869 | 24698.0763 | 30972.3739 |
| *YBL005W-B* | 0.326850311 | 0.15502048 | 0.99979869 | 23415.2747 | 29369.1173 |
| *KAR2* | 0.320135415 | 0.15507462 | 0.99979869 | 137743.624 | 171965.664 |
| *GND2* | 0.426750261 | 0.1552081 | 0.99979869 | 1429.12875 | 1921.03755 |
| *STH1* | 0.325139724 | 0.15561237 | 0.99979869 | 26293.0425 | 32939.5409 |
| *SGA1* | 0.461887121 | 0.15666022 | 0.99979869 | 992.098253 | 1366.45841 |
| *HXT13* | 0.566579182 | 0.15793307 | 0.99979869 | 478.977622 | 709.369257 |
| *XKS1* | 0.336306239 | 0.15806967 | 0.99979869 | 8343.57557 | 10533.9284 |
| *UPF3* | 0.336846697 | 0.15810449 | 0.99979869 | 8112.37863 | 10245.8753 |
| *BAR1* | 0.316915827 | 0.15947122 | 0.99979869 | 46000.3865 | 57301.0444 |
| *RSF2* | 0.321800075 | 0.15954038 | 0.99979869 | 20377.5225 | 25469.6366 |
| *NTH1* | 0.317859698 | 0.159556 | 0.99979869 | 36367.1808 | 45330.9507 |
| *SYM1* | 0.341926783 | 0.16065245 | 0.99979869 | 5688.22509 | 7209.52888 |
| *UGA2* | 0.339338587 | 0.16099779 | 0.99979869 | 6131.10867 | 7756.93232 |
| *CAR2* | 0.322555655 | 0.16105091 | 0.99979869 | 15475.5621 | 19352.8644 |
| *BBC1* | 0.316114378 | 0.16113171 | 0.99979869 | 34072.7707 | 42419.6665 |
| *AIM3* | 0.326895247 | 0.16125941 | 0.99979869 | 10886.7419 | 13655.3582 |
| *MYO2* | 0.314129328 | 0.16144134 | 0.99979869 | 48343.5724 | 60103.668 |
| *APE1* | 0.321542485 | 0.16150359 | 0.99979869 | 15832.6004 | 19785.4567 |
| *REC114* | 0.384639076 | 0.16224794 | 0.99979869 | 2112.96442 | 2758.54432 |
| *TPO1* | 0.31765488 | 0.16237037 | 0.99979869 | 20929.6637 | 26084.6967 |

| | | | | | |
|---|---|---|---|---|---|
| *EDC2* | 0.321503397 | 0.1629636 | 0.99979869 | 13494.2922 | 16862.8963 |
| *SNA2* | 0.386843697 | 0.16312019 | 0.99979869 | 2004.68231 | 2621.18091 |
| *ZNF1* | 0.531193857 | 0.16331461 | 0.99979869 | 551.165696 | 796.502764 |
| *SWI4* | 0.336968782 | 0.16362646 | 0.99979869 | 5848.20946 | 7386.87119 |
| *FUS2* | 0.464231201 | 0.16382653 | 0.99979869 | 873.085483 | 1204.4926 |
| *BCY1* | 0.313113207 | 0.16384576 | 0.99979869 | 30107.3042 | 37404.8769 |
| *SOD2* | 0.310070231 | 0.1650713 | 0.99979869 | 39867.3268 | 49426.2255 |
| *IRC4* | 0.416642852 | 0.16535201 | 0.99979869 | 1324.7487 | 1768.29764 |
| *YBL086C* | 0.325349774 | 0.16548744 | 0.99979869 | 8605.98897 | 10783.0278 |
| *SSA1* | 0.304950694 | 0.16601404 | 0.99979869 | 297205.127 | 367160.096 |
| *SCM4* | 0.345247533 | 0.1665535 | 0.99979869 | 4024.97285 | 5113.19921 |
| *TPK1* | 0.321774961 | 0.1671954 | 0.99979869 | 9229.34274 | 11535.4512 |
| *RPT4* | 0.306275203 | 0.16805485 | 0.99979869 | 39948.2945 | 49396.4976 |
| *UBX6* | 0.323289675 | 0.16837689 | 0.99979869 | 7894.8389 | 9877.86439 |
| *YLL067C* | 0.708060038 | 0.16867053 | 0.99979869 | 223.392822 | 364.935629 |
| *YIL055C* | 0.363970581 | 0.16878646 | 0.99979869 | 2526.58257 | 3251.61746 |
| *SIM1* | 0.304405282 | 0.16893965 | 0.99979869 | 48270.4088 | 59609.5698 |
| *YJL070C* | 0.331281895 | 0.16908747 | 0.99979869 | 5508.73042 | 6930.70165 |
| *FRK1* | 0.322711748 | 0.16916428 | 0.99979869 | 7729.97695 | 9667.71887 |
| *FAA1* | 0.304619703 | 0.16928337 | 0.99979869 | 41101.3527 | 50763.9812 |
| *YPR015C* | 0.459046468 | 0.16961006 | 0.99979869 | 838.942475 | 1153.23759 |
| *YJR120W* | 0.402234187 | 0.16977251 | 0.99979869 | 1438.8839 | 1901.56065 |
| *PAB1* | 0.300436911 | 0.17003508 | 0.99979869 | 166002.328 | 204434.684 |
| *CHS2* | 0.309447764 | 0.17020581 | 0.99979869 | 17079.308 | 21165.2414 |
| *GPG1* | 0.423336721 | 0.17039348 | 0.99979869 | 1137.44991 | 1525.34892 |
| *ERO1* | 0.305443677 | 0.17053908 | 0.99979869 | 26418.8839 | 32648.4125 |
| *SCH9* | 0.310375831 | 0.17087308 | 0.99979869 | 14531.2641 | 18019.2092 |
| *ARG5%2C6* | 0.316154942 | 0.17106891 | 0.99979869 | 9565.89525 | 11909.6128 |
| *ATP10* | 0.366253021 | 0.17145671 | 0.99979869 | 2276.85086 | 2934.86153 |
| *ACE2* | 0.317031193 | 0.17172683 | 0.99979869 | 8708.418 | 10848.6342 |
| *HXT5* | 0.343782825 | 0.17247259 | 0.99979869 | 3424.05592 | 4345.39925 |
| *HAC1* | 0.29750292 | 0.17252445 | 0.99979869 | 155009.255 | 190508.7 |
| *RBA50* | 0.309378827 | 0.17356004 | 0.99979869 | 12110.0371 | 15006.4401 |
| *FMP33* | 0.377451223 | 0.17393894 | 0.99979869 | 1821.28558 | 2365.93099 |
| *MNN5* | 0.30138013 | 0.17427848 | 0.99979869 | 24603.4514 | 30319.3851 |
| *AXL2* | 0.321203252 | 0.17442051 | 0.99979869 | 6190.61506 | 7734.38012 |
| *SDH3* | 0.306048954 | 0.1744919 | 0.99979869 | 14380.0593 | 17778.3107 |
| *STE2* | 0.295799741 | 0.17457382 | 0.99979869 | 94392.7346 | 115873.213 |
| *TPS1* | 0.294799939 | 0.17523808 | 0.99979869 | 106954.435 | 131202.559 |
| *MPT5* | 0.310092155 | 0.17551093 | 0.99979869 | 9879.0354 | 12247.8958 |
| *SAP4* | 0.328846514 | 0.17572599 | 0.99979869 | 4521.50974 | 5679.05445 |
| *YPL119C-A* | 0.593482943 | 0.17584659 | 0.99979869 | 342.405592 | 516.650441 |
| *YAR035C-A* | 1.978419922 | 0.17626636 | 0.99979869 | 11.7061741 | 46.1295037 |
| *TDA9* | 0.302475565 | 0.17637315 | 0.99979869 | 16194.5163 | 19972.0249 |
| *NIS1* | 0.302737078 | 0.17659415 | 0.99979869 | 15417.0313 | 19016.6316 |

| | | | | | |
|---|---|---|---|---|---|
| *DEP1* | 0.303898464 | 0.17666162 | 0.99979869 | 13837.6733 | 17082.2678 |
| *GIP3* | 0.302744401 | 0.1774531 | 0.99979869 | 14086.4295 | 17375.4464 |
| *ADY3* | 0.35807212 | 0.17794346 | 0.99979869 | 2246.60991 | 2879.50613 |
| *VHS1* | 0.302010389 | 0.17795674 | 0.99979869 | 14253.2424 | 17572.2656 |
| *YMR279C* | 0.330975489 | 0.17802854 | 0.99979869 | 3926.44589 | 4938.93219 |
| *HXT9* | 0.569814412 | 0.17895657 | 0.99979869 | 373.622056 | 554.579144 |
| *YJL045W* | 0.435884059 | 0.18056763 | 0.99979869 | 876.012026 | 1185.01569 |
| *YER121W* | 0.467790351 | 0.18087542 | 0.99979869 | 678.958096 | 938.991675 |
| *COX19* | 0.310672445 | 0.18204107 | 0.99979869 | 6503.75521 | 8066.51254 |
| *SLM1* | 0.296810455 | 0.18229393 | 0.99979869 | 14464.9291 | 17769.0848 |
| *SRT1* | 0.732843849 | 0.18234243 | 0.99979869 | 179.494669 | 298.304124 |
| *YML119W* | 0.309021765 | 0.18260518 | 0.99979869 | 6781.77685 | 8401.72027 |
| *TPH3* | 0.300911259 | 0.18269194 | 0.99979869 | 10302.4087 | 12691.7641 |
| *COS1* | 0.321036321 | 0.18270372 | 0.99979869 | 4407.37454 | 5505.81254 |
| *POG1* | 0.324661186 | 0.18314057 | 0.99979869 | 3921.56831 | 4911.25449 |
| *KIN1* | 0.292514155 | 0.18339894 | 0.99979869 | 19713.1971 | 24144.1822 |
| *ESBP6* | 0.30813253 | 0.18406964 | 0.99979869 | 6546.67785 | 8105.46635 |
| *MIG1* | 0.297505907 | 0.18456839 | 0.99979869 | 11206.7106 | 13773.2447 |
| *LEE1* | 0.439349452 | 0.18501152 | 0.99979869 | 802.848439 | 1088.65629 |
| *STE6* | 0.284990745 | 0.18517286 | 0.99979869 | 56879.3243 | 69301.891 |
| *YHR097C* | 0.288515664 | 0.18673568 | 0.99979869 | 20453.6126 | 24981.689 |
| *FAR11* | 0.298883832 | 0.18676757 | 0.99979869 | 8818.65114 | 10848.6342 |
| *GCY1* | 0.304770697 | 0.18696048 | 0.99979869 | 6554.48197 | 8096.24045 |
| *ATG14* | 0.364735855 | 0.18702927 | 0.99979869 | 1692.51767 | 2179.36277 |
| *MAL11* | 0.326307585 | 0.18718585 | 0.99979869 | 3333.33307 | 4179.33303 |
| *HXT12* | 0.398458742 | 0.18771778 | 0.99979869 | 1123.79271 | 1481.26962 |
| *YMR045C* | 0.289322011 | 0.1877304 | 0.99979869 | 16546.6771 | 20221.1242 |
| *ARA1* | 0.283862957 | 0.18789955 | 0.99979869 | 35179.0041 | 42828.6814 |
| *MPO1* | 0.346316827 | 0.18864632 | 0.99979869 | 2166.61772 | 2754.44392 |
| *MEF2* | 0.313584074 | 0.1887933 | 0.99979869 | 4448.34615 | 5528.36474 |
| *YLR345W* | 0.294098974 | 0.18883158 | 0.99979869 | 10239.9758 | 12555.4258 |
| *SET1* | 0.292365972 | 0.18967084 | 0.99979869 | 10857.4765 | 13296.5732 |
| *SPO1* | 0.350591801 | 0.18970131 | 0.99979869 | 1979.31893 | 2523.7964 |
| *STE11* | 0.302000153 | 0.19038131 | 0.99979869 | 6279.38688 | 7741.55582 |
| *APJ1* | 0.286743901 | 0.19079178 | 0.99979869 | 15498.9745 | 18906.9459 |
| *SPS19* | 0.35463314 | 0.19082121 | 0.99979869 | 1822.2611 | 2330.05249 |
| *GIC2* | 0.289392295 | 0.19140698 | 0.99979869 | 11735.4395 | 14342.1752 |
| *SVS1* | 0.285339172 | 0.19180544 | 0.99979869 | 15999.4134 | 19498.4287 |
| *HED1* | 0.654372818 | 0.19330278 | 0.99979869 | 221.441793 | 348.534028 |
| *CCW12* | 0.27332385 | 0.19366527 | 0.99979869 | 999111.226 | 1207514.59 |
| *ZTA1* | 0.291985433 | 0.19388962 | 0.99979869 | 8413.81262 | 10301.2307 |
| *MDY2* | 0.282086027 | 0.19486503 | 0.99979869 | 16066.7239 | 19536.3574 |
| *MSH5* | 0.326488824 | 0.19545031 | 0.99979869 | 2679.73835 | 3360.27807 |
| *YOR338W* | 0.347131088 | 0.19632812 | 0.99979869 | 1839.82036 | 2340.30349 |
| *RGC1* | 0.278769858 | 0.19669237 | 0.99979869 | 18780.6053 | 22783.8744 |

| | | | | | |
|---|---|---|---|---|---|
| *HSL1* | 0.280735913 | 0.19723748 | 0.99979869 | 14459.076 | 17565.0899 |
| *YHL008C* | 0.284171996 | 0.19757578 | 0.99979869 | 10676.0308 | 13000.3192 |
| *VIK1* | 0.31873262 | 0.19803836 | 0.99979869 | 2964.58858 | 3697.536 |
| *YBR225W* | 0.292127401 | 0.19833173 | 0.99979869 | 6613.01284 | 8097.26555 |
| *ZRG8* | 0.298963546 | 0.19833526 | 0.99979869 | 5056.09169 | 6220.3073 |
| *ROD1* | 0.288707404 | 0.19857538 | 0.99979869 | 7686.07879 | 9388.89165 |
| *YDR391C* | 0.285742886 | 0.19902764 | 0.99979869 | 8800.11636 | 10727.6724 |
| *UBP11* | 0.300122981 | 0.19951709 | 0.99979869 | 4659.05728 | 5736.46006 |
| *BXI1* | 0.279425938 | 0.19952407 | 0.99979869 | 13172.3724 | 15987.4609 |
| *DSE4* | 0.269959717 | 0.19961362 | 0.99979869 | 59880.9824 | 72202.9243 |
| *YNL058C* | 0.279217075 | 0.19964059 | 0.99979869 | 13274.8014 | 16109.4478 |
| *YDR034W-B* | 0.38235646 | 0.19974664 | 0.99979869 | 1119.89065 | 1459.74252 |
| *UBP9* | 0.300502425 | 0.20091313 | 0.99979869 | 4389.81528 | 5406.37783 |
| *YER158C* | 0.283580323 | 0.20095836 | 0.99979869 | 8896.6923 | 10829.1573 |
| *COX13* | 0.279231978 | 0.20119728 | 0.99979869 | 11692.5169 | 14189.4353 |
| *VMA2* | 0.266770158 | 0.20170962 | 0.99979869 | 101183.291 | 121734.735 |
| *YIR016W* | 0.28255195 | 0.20197923 | 0.99979869 | 8893.76575 | 10817.8812 |
| *HSP31* | 0.291498051 | 0.20259313 | 0.99979869 | 5644.32693 | 6908.14945 |
| *CHC1* | 0.269742351 | 0.20265736 | 0.99979869 | 29329.8191 | 35359.8022 |
| *IVY1* | 0.278750578 | 0.20270704 | 0.99979869 | 10843.8192 | 13155.1093 |
| *IST2* | 0.267286946 | 0.20278896 | 0.99979869 | 51217.4381 | 61642.3432 |
| *HSP78* | 0.267571682 | 0.20294233 | 0.99979869 | 45005.3617 | 54176.5393 |
| *RMR1* | 0.315746065 | 0.20310215 | 0.99979869 | 2797.7756 | 3482.26498 |
| *COX5A* | 0.276906467 | 0.20382033 | 0.99979869 | 11445.7117 | 13867.5539 |
| *SLA1* | 0.266461413 | 0.20384837 | 0.99979869 | 48280.1639 | 58073.9698 |
| *YSC84* | 0.2738836 | 0.20452949 | 0.99979869 | 13851.3305 | 16747.06 |
| *YHR210C* | 0.354867154 | 0.20472281 | 0.99979869 | 1434.00632 | 1833.90405 |
| *EDC1* | 0.292052128 | 0.20492638 | 0.99979869 | 5056.09169 | 6190.57939 |
| *RGS2* | 0.306273106 | 0.20531153 | 0.99979869 | 3290.41043 | 4068.62222 |
| *ATP20* | 0.298490598 | 0.20541145 | 0.99979869 | 4042.53211 | 4971.7354 |
| *MSC3* | 0.270817476 | 0.20548005 | 0.99979869 | 17148.5695 | 20689.595 |
| *PIN4* | 0.268142668 | 0.20676281 | 0.99979869 | 20411.6655 | 24580.8749 |
| *TKL2* | 0.327190565 | 0.20695493 | 0.99979869 | 2072.96833 | 2600.67891 |
| *HYP2* | 0.260360906 | 0.20761368 | 0.99979869 | 211712.987 | 253585.158 |
| *TOS6* | 0.274991919 | 0.20794485 | 0.99979869 | 9867.32923 | 11939.3407 |
| *MSB2* | 0.266121669 | 0.20862787 | 0.99979869 | 21131.5952 | 25412.231 |
| *FEX2* | 0.411883579 | 0.20866737 | 0.99979869 | 758.950286 | 1009.72358 |
| *KAP95* | 0.265034328 | 0.20886873 | 0.99979869 | 23943.028 | 28771.484 |
| *TPO3* | 0.26227545 | 0.20897624 | 0.99979869 | 40406.7864 | 48462.6315 |
| *YFL051C* | 0.599832649 | 0.20930928 | 0.99979869 | 244.854141 | 371.08623 |
| *YOR387C* | 0.448127033 | 0.21003685 | 0.99979869 | 565.798414 | 771.900362 |
| *DAN4* | 0.309734828 | 0.21007652 | 0.99979869 | 2706.07724 | 3354.12747 |
| *GEX2* | 0.346088891 | 0.21011653 | 0.99979869 | 1479.85551 | 1881.05865 |
| *ALK1* | 0.270742714 | 0.21030564 | 0.99979869 | 11357.9154 | 13702.5128 |
| *AHC1* | 0.269791014 | 0.21095015 | 0.99979869 | 11661.3004 | 14059.2476 |

| | | | | | |
|---|---|---|---|---|---|
| *TFG1* | 0.261193548 | 0.21109892 | 0.99979869 | 32959.7086 | 39501.2066 |
| *YBR139W* | 0.264051916 | 0.21191991 | 0.99979869 | 18593.3065 | 22327.7049 |
| *FLC1* | 0.260835326 | 0.2122135 | 0.99979869 | 28900.5928 | 34627.8808 |
| *SDH8* | 0.273373669 | 0.21285485 | 0.99979869 | 8159.20333 | 9861.46279 |
| *MMP1* | 0.271783788 | 0.21454268 | 0.99979869 | 8163.10539 | 9855.31219 |
| *EDE1* | 0.255819893 | 0.21554823 | 0.99979869 | 45612.1318 | 54461.5171 |
| *YAK1* | 0.256323173 | 0.21576352 | 0.99979869 | 37855.8159 | 45216.1395 |
| *ATP14* | 0.261451503 | 0.21757379 | 0.99979869 | 14095.2091 | 16895.6995 |
| *KRE1* | 0.253976003 | 0.21807559 | 0.99979869 | 41059.4056 | 48962.8803 |
| *MSS4* | 0.261626064 | 0.21831458 | 0.99979869 | 13056.2861 | 15652.2531 |
| *ATG2* | 0.272757959 | 0.21891963 | 0.99979869 | 6334.9912 | 7653.39721 |
| *PUF2* | 0.263986423 | 0.21908394 | 0.99979869 | 10267.2902 | 12328.8787 |
| *ERG5* | 0.256691345 | 0.22047783 | 0.99979869 | 17794.3601 | 21259.5506 |
| *TGL5* | 0.263252129 | 0.22078315 | 0.99979869 | 9725.87963 | 11672.8146 |
| *FMT1* | 0.315873436 | 0.22097378 | 0.99979869 | 1945.17593 | 2421.28639 |
| *HMX1* | 0.345195339 | 0.22117994 | 0.99979869 | 1274.99746 | 1619.65813 |
| *YBR085C-A* | 0.25424325 | 0.22144253 | 0.99979869 | 22022.24 | 26266.1394 |
| *QCR6* | 0.254103483 | 0.22155695 | 0.99979869 | 22155.8855 | 26422.9797 |
| *YFR016C* | 0.252395647 | 0.22173323 | 0.99979869 | 28762.0697 | 34260.8949 |
| *MAL33* | 0.332667275 | 0.22194659 | 0.99979869 | 1484.73308 | 1869.78255 |
| *SPO22* | 0.289825941 | 0.22272038 | 0.99979869 | 3194.81001 | 3905.63131 |
| *FDH1* | 0.838869956 | 0.22277709 | 0.99979869 | 91.6983636 | 164.016013 |
| *GAB1* | 0.269326155 | 0.22278876 | 0.99979869 | 6306.70128 | 7601.11711 |
| *TPO4* | 0.249533475 | 0.22310305 | 0.99979869 | 42484.6323 | 50506.681 |
| *CIT1* | 0.250588199 | 0.22353916 | 0.99979869 | 30311.1867 | 36060.9707 |
| *IML2* | 0.255648715 | 0.22366623 | 0.99979869 | 14734.1711 | 17590.7174 |
| *RPN4* | 0.253220936 | 0.22481468 | 0.99979869 | 17233.4393 | 20539.9303 |
| *ROG3* | 0.270704204 | 0.22528373 | 0.99979869 | 5403.37485 | 6518.61142 |
| *PRR2* | 0.384302553 | 0.22534964 | 0.99979869 | 801.872924 | 1046.62718 |
| *YIL024C* | 0.355511286 | 0.22556068 | 0.99979869 | 1067.21287 | 1365.43331 |
| *GSP2* | 0.266673563 | 0.22565898 | 0.99979869 | 6331.08914 | 7616.49361 |
| *SIR1* | 0.298353017 | 0.22613875 | 0.99979869 | 2446.59038 | 3008.66874 |
| *SWI5* | 0.255647887 | 0.22635187 | 0.99979869 | 11939.322 | 14254.0166 |
| *COQ1* | 0.258194361 | 0.22665727 | 0.99979869 | 9668.32427 | 11563.1289 |
| *GAS1* | 0.243502554 | 0.22687035 | 0.99979869 | 224741.958 | 266063.701 |
| *YNL115C* | 0.255471183 | 0.22705147 | 0.99979869 | 11535.459 | 13770.1694 |
| *GPT2* | 0.254906789 | 0.22810425 | 0.99979869 | 11235.0006 | 13406.2589 |
| *TDA10* | 0.264229795 | 0.2281308 | 0.99979869 | 6410.10582 | 7698.50161 |
| *GOS1* | 0.257918381 | 0.22891294 | 0.99979869 | 8679.15256 | 10378.1132 |
| *YKR015C* | 0.317875575 | 0.22946391 | 0.99979869 | 1639.83988 | 2044.04956 |
| *SVL3* | 0.248600438 | 0.23006401 | 0.99979869 | 17682.1759 | 21007.376 |
| *WSC4* | 0.255062996 | 0.23079761 | 0.99979869 | 9412.73947 | 11233.0467 |
| *BNI1* | 0.244483855 | 0.2309587 | 0.99979869 | 29962.9281 | 35496.1405 |
| *NAM8* | 0.258708508 | 0.23136353 | 0.99979869 | 7351.47732 | 8795.3587 |
| *HBT1* | 0.262989551 | 0.23154429 | 0.99979869 | 5938.93231 | 7126.49577 |

| | | | | | |
|---|---|---|---|---|---|
| *SSN2* | 0.257004817 | 0.2321087 | 0.99979869 | 7821.67531 | 9346.86255 |
| *YBR285W* | 0.445147668 | 0.23259745 | 0.99979869 | 465.320419 | 633.511851 |
| *VRG4* | 0.241994202 | 0.23307391 | 0.99979869 | 35654.0797 | 42165.4417 |
| *CPS1* | 0.243778046 | 0.23338757 | 0.99979869 | 23808.407 | 28191.2773 |
| *RPO21* | 0.240009028 | 0.23386929 | 0.99979869 | 53023.1155 | 62620.2887 |
| *MNR2* | 0.244245969 | 0.23400629 | 0.99979869 | 20541.409 | 24330.7504 |
| *KCC4* | 0.268321963 | 0.23426095 | 0.99979869 | 4405.42351 | 5305.91802 |
| *ULA1* | 0.261431991 | 0.23438051 | 0.99979869 | 5713.58846 | 6848.69365 |
| *SFL1* | 0.256097583 | 0.235355 | 0.99979869 | 7076.38223 | 8450.92507 |
| *PPH22* | 0.245581121 | 0.23590717 | 0.99979869 | 14607.3542 | 17318.0408 |
| *PHR1* | 0.254941621 | 0.23591802 | 0.99979869 | 7339.77114 | 8758.4551 |
| *HEM1* | 0.241852876 | 0.2360571 | 0.99979869 | 23175.2981 | 27405.0256 |
| *APE3* | 0.23607743 | 0.23623711 | 0.99979869 | 206763.226 | 243522.775 |
| *ECM38* | 0.24964563 | 0.23624334 | 0.99979869 | 10051.7015 | 11950.6168 |
| *DIP5* | 0.242033042 | 0.23649003 | 0.99979869 | 21360.8411 | 25262.5664 |
| *PTA1* | 0.24490303 | 0.2372299 | 0.99979869 | 14041.5558 | 16639.4245 |
| *COX6* | 0.249888137 | 0.23726618 | 0.99979869 | 9318.11456 | 11080.3068 |
| *YEL014C* | 0.398432155 | 0.23831816 | 0.99979869 | 621.40274 | 819.054965 |
| *YIL108W* | 0.245515894 | 0.23978035 | 0.99979869 | 11074.0407 | 13128.4567 |
| *HMS1* | 0.302559855 | 0.24034591 | 0.99979869 | 1763.73023 | 2175.26237 |
| *HXT7* | 2.121591076 | 0.24034842 | 0.99979869 | 1519.8516 | 6613.94573 |
| *FKS1* | 0.233058062 | 0.24093708 | 0.99979869 | 115015.111 | 135179.948 |
| *BAG7* | 0.27719803 | 0.24108797 | 0.99979869 | 2877.76779 | 3487.39048 |
| *MTF2* | 0.267546639 | 0.24147581 | 0.99979869 | 3700.12652 | 4454.05986 |
| *RRT8* | 0.250750682 | 0.24174489 | 0.99979869 | 7109.54972 | 8459.12587 |
| *HRK1* | 0.235416197 | 0.24180741 | 0.99979869 | 33708.9038 | 39683.6744 |
| *LPP1* | 0.280446597 | 0.24188922 | 0.99979869 | 2623.15851 | 3186.01105 |
| *DAL82* | 0.272021345 | 0.24352733 | 0.99979869 | 3107.98922 | 3752.8914 |
| *PDE1* | 0.244371123 | 0.24353488 | 0.99979869 | 9641.98538 | 11421.6651 |
| *PHO91* | 0.235196801 | 0.24389932 | 0.99979869 | 25392.6426 | 29888.8431 |
| *GUD1* | 0.266775985 | 0.24444077 | 0.99979869 | 3513.80325 | 4227.51274 |
| *TDA6* | 0.240699509 | 0.24518109 | 0.99979869 | 11596.9164 | 13702.5128 |
| *YHL044W* | 0.319573695 | 0.24570118 | 0.99979869 | 1274.02194 | 1589.93023 |
| *PCA1* | 0.247204546 | 0.24575046 | 0.99979869 | 7247.09727 | 8601.61479 |
| *YOL162W* | 0.669976248 | 0.24600354 | 0.99979869 | 138.52306 | 220.396518 |
| *HMG1* | 0.234180396 | 0.24639289 | 0.99979869 | 21873.9618 | 25728.987 |
| *ATH1* | 0.248003266 | 0.24661682 | 0.99979869 | 6694.95606 | 7950.67623 |
| *BLM10* | 0.231528768 | 0.24702924 | 0.99979869 | 33086.5255 | 38846.1676 |
| *SEC16* | 0.233930073 | 0.24712046 | 0.99979869 | 20918.9331 | 24601.3769 |
| *MEP3* | 0.240911884 | 0.24823881 | 0.99979869 | 9506.38886 | 11234.0718 |
| *ALY2* | 0.243635584 | 0.24824119 | 0.99979869 | 7953.36977 | 9416.56935 |
| *UGX2* | 0.361531429 | 0.24842316 | 0.99979869 | 775.534032 | 996.39728 |
| *VPS17* | 0.237037068 | 0.24889997 | 0.99979869 | 12478.7816 | 14707.1109 |
| *HSF1* | 0.233613637 | 0.24906651 | 0.99979869 | 17877.2788 | 21019.6772 |
| *TPS3* | 0.230139745 | 0.24910821 | 0.99979869 | 32049.5536 | 37592.4702 |

| | | | | | |
|---|---|---|---|---|---|
| *SSE1* | 0.225969329 | 0.24963262 | 0.99979869 | 224459.059 | 262518.905 |
| *BIO2* | 0.245221101 | 0.25009523 | 0.99979869 | 6717.39289 | 7961.95234 |
| *DGR2* | 0.251491163 | 0.25030204 | 0.99979869 | 5015.12008 | 5970.18288 |
| *BUD3* | 0.240007123 | 0.25052059 | 0.99979869 | 8969.85588 | 10593.3842 |
| *YNL092W* | 0.292346436 | 0.25058398 | 0.99979869 | 1787.14258 | 2188.58867 |
| *SPG1* | 0.539679133 | 0.25104868 | 0.99979869 | 234.123481 | 340.333227 |
| *FLO10* | 0.297260584 | 0.25155826 | 0.99979869 | 1624.23165 | 1995.86986 |
| *ATG7* | 0.263847286 | 0.25166739 | 0.99979869 | 3222.12441 | 3868.72771 |
| *YBL081W* | 0.250557028 | 0.25202193 | 0.99979869 | 4932.20134 | 5867.67287 |
| *CDC20* | 0.237795356 | 0.25244863 | 0.99979869 | 9455.66211 | 11150.0136 |
| *POM33* | 0.231368968 | 0.2527438 | 0.99979869 | 16818.8456 | 19744.4527 |
| *GIP4* | 0.2416268 | 0.25286575 | 0.99979869 | 7273.43616 | 8599.56459 |
| *URA8* | 0.228726279 | 0.25295873 | 0.99979869 | 24210.319 | 28369.6448 |
| *YHR033W* | 0.262754235 | 0.25302036 | 0.99979869 | 3221.1489 | 3864.62731 |
| *PRR1* | 0.240290136 | 0.25306915 | 0.99979869 | 7790.45885 | 9202.32343 |
| *CHS6* | 0.251108119 | 0.2531495 | 0.99979869 | 4665.88588 | 5552.96714 |
| *URA1* | 0.224343195 | 0.25334376 | 0.99979869 | 84739.0431 | 98995.9651 |
| *SKY1* | 0.232065334 | 0.25343392 | 0.99979869 | 14636.6196 | 17190.9284 |
| *WSC2* | 0.232362441 | 0.25370573 | 0.99979869 | 13888.4 | 16315.4929 |
| *LOH1* | 0.424890696 | 0.25407689 | 0.99979869 | 442.883586 | 594.558047 |
| *YPL261C* | 0.485403771 | 0.25484793 | 0.99979869 | 302.409497 | 423.366334 |
| *ULP2* | 0.238682137 | 0.25521125 | 0.99979869 | 7790.45885 | 9192.07243 |
| *CCW14* | 0.222411571 | 0.25586844 | 0.99979869 | 92784.1112 | 108249.544 |
| *XYL2* | 0.281306375 | 0.25601813 | 0.99979869 | 2000.78025 | 2431.53739 |
| *AIM19* | 0.252787301 | 0.25668522 | 0.99979869 | 3985.95227 | 4749.28868 |
| *UBC8* | 0.240545637 | 0.25692893 | 0.99979869 | 6537.89822 | 7724.12912 |
| *HST3* | 0.238450485 | 0.25772379 | 0.99979869 | 7086.13737 | 8359.69117 |
| *DAK2* | 0.302665944 | 0.2577356 | 0.99979869 | 1375.47545 | 1696.54064 |
| *SKG3* | 0.241877149 | 0.25781995 | 0.99979869 | 5923.32408 | 7004.50886 |
| *PMD1* | 0.234740631 | 0.25897321 | 0.99979869 | 8380.64512 | 9861.46279 |
| *NDE1* | 0.221317375 | 0.25930325 | 0.99979869 | 49521.9939 | 57732.6115 |
| *ATP17* | 0.231289741 | 0.25940407 | 0.99979869 | 10513.1198 | 12341.1799 |
| *CST9* | 0.254469854 | 0.25945956 | 0.99979869 | 3528.43597 | 4209.06094 |
| *UBR2* | 0.232387935 | 0.25988947 | 0.99979869 | 9411.76395 | 11056.7295 |
| *YFL040W* | 0.335613965 | 0.2599141 | 0.99979869 | 893.571288 | 1127.61009 |
| *TVP15* | 0.233544705 | 0.26055879 | 0.99979869 | 8405.03298 | 9881.96479 |
| *PRY3* | 0.220475971 | 0.26102124 | 0.99979869 | 44013.2635 | 51280.6316 |
| *ZIM17* | 0.240242418 | 0.26123729 | 0.99979869 | 5677.49443 | 6706.20474 |
| *CLG1* | 0.22379879 | 0.26150103 | 0.99979869 | 20112.1826 | 23487.0931 |
| *TBF1* | 0.23926312 | 0.26166945 | 0.99979869 | 5855.03807 | 6911.22475 |
| *ATP3* | 0.223630627 | 0.26171895 | 0.99979869 | 20161.9338 | 23542.4485 |
| *KIN3* | 0.23827198 | 0.26186555 | 0.99979869 | 6098.91669 | 7194.15237 |
| *SCW11* | 0.220631794 | 0.26234482 | 0.99979869 | 32729.4872 | 38137.8234 |
| *URA4* | 0.220823233 | 0.26252485 | 0.99979869 | 30412.6402 | 35442.8353 |
| *DSD1* | 0.226691901 | 0.26267618 | 0.99979869 | 12861.1832 | 15049.4943 |

| | | | | | |
|---|---|---|---|---|---|
| *ECM30* | 0.224236403 | 0.26288664 | 0.99979869 | 16600.3304 | 19391.8182 |
| *NTH2* | 0.230009805 | 0.26291302 | 0.99979869 | 9561.01767 | 11213.5698 |
| *SEC23* | 0.218360235 | 0.26302592 | 0.99979869 | 57865.5695 | 67321.3977 |
| *XRN1* | 0.217788688 | 0.26379171 | 0.99979869 | 59163.9793 | 68804.7175 |
| *SFC1* | 0.584481664 | 0.26380327 | 0.99979869 | 173.641582 | 260.375421 |
| *JIP4* | 0.223670623 | 0.26387839 | 0.99979869 | 16349.6231 | 19091.4639 |
| *DIA3* | 0.341876571 | 0.26436128 | 0.99979869 | 799.921895 | 1013.82398 |
| *CUZ1* | 0.224522993 | 0.26466977 | 0.99979869 | 13925.4696 | 16270.3885 |
| *OYE3* | 0.373900696 | 0.26476315 | 0.99979869 | 594.088334 | 769.850161 |
| *VHR2* | 0.228808495 | 0.26488382 | 0.99979869 | 9426.39667 | 11046.4785 |
| *ETP1* | 0.244445918 | 0.26492446 | 0.99979869 | 4282.50868 | 5073.2203 |
| *LRE1* | 0.230748869 | 0.26496829 | 0.99979869 | 8209.93008 | 9633.89057 |
| *HEF3* | 0.268864809 | 0.26499081 | 0.99979869 | 2209.54036 | 2662.18491 |
| *CLN2* | 0.228893698 | 0.26501939 | 0.99979869 | 9302.50633 | 10901.9394 |
| *GLC3* | 0.219829641 | 0.26514595 | 0.99979869 | 25761.3871 | 30001.6041 |
| *YPL067C* | 0.23795654 | 0.26528262 | 0.99979869 | 5504.82836 | 6491.95882 |
| *RPE1* | 0.218681621 | 0.26542308 | 0.99979869 | 31587.1597 | 36757.0136 |
| *STP22* | 0.228285748 | 0.26544002 | 0.99979869 | 9519.07055 | 11151.0387 |
| *MCM3* | 0.22069055 | 0.26557621 | 0.99979869 | 21064.2847 | 24546.0215 |
| *TEC1* | 0.226555059 | 0.26601794 | 0.99979869 | 10559.9445 | 12355.5313 |
| *tG(GCC)G1* | 0.812287971 | 0.26660771 | 0.99979869 | 74.1391025 | 130.18771 |
| *PPE1* | 0.262104693 | 0.26718099 | 0.99979869 | 2466.10067 | 2957.41374 |
| *CDC15* | 0.234593666 | 0.26849546 | 0.99979869 | 5779.92345 | 6800.51394 |
| *NDD1* | 0.229730409 | 0.26868599 | 0.99979869 | 7432.44502 | 8715.4009 |
| *GGA1* | 0.223108201 | 0.26899036 | 0.99979869 | 11931.5179 | 13927.0097 |
| *AKL1* | 0.216958511 | 0.27065142 | 0.99979869 | 22185.1509 | 25785.3675 |
| *ASE1* | 0.235191919 | 0.27073824 | 0.99979869 | 5229.73327 | 6155.72599 |
| *CAT2* | 0.265488254 | 0.27110148 | 0.99979869 | 2142.22986 | 2575.05141 |
| *PTK2* | 0.215006222 | 0.27122745 | 0.99979869 | 30161.933 | 35009.218 |
| *ATG33* | 0.242174904 | 0.27156659 | 0.99979869 | 3907.91111 | 4622.17627 |
| *MCM21* | 0.240721611 | 0.2721146 | 0.99979869 | 4053.26277 | 4789.26758 |
| *YDL199C* | 0.250736688 | 0.27222567 | 0.99979869 | 2972.3927 | 3536.59528 |
| *EMI2* | 0.212742959 | 0.27281585 | 0.99979869 | 41816.4048 | 48460.5813 |
| *BFR2* | 0.219970587 | 0.27286635 | 0.99979869 | 12539.2635 | 14604.6009 |
| *RRN5* | 0.227169117 | 0.27416219 | 0.99979869 | 6919.32439 | 8099.31575 |
| *MYO3* | 0.216675406 | 0.27427044 | 0.99979869 | 16313.5291 | 18957.1758 |
| *JSN1* | 0.213782115 | 0.27482829 | 0.99979869 | 23939.126 | 27762.7855 |
| *HOR7* | 0.20947334 | 0.27507453 | 0.99979869 | 97400.2459 | 112620.57 |
| *PMC1* | 0.211975465 | 0.27561496 | 0.99979869 | 31408.6406 | 36379.7768 |
| *PLB1* | 0.21221542 | 0.27563762 | 0.99979869 | 29600.0367 | 34290.6228 |
| *UBP15* | 0.217988907 | 0.2757904 | 0.99979869 | 12589.9902 | 14643.5547 |
| *CDC27* | 0.224977406 | 0.27635445 | 0.99979869 | 7206.12566 | 8422.22227 |
| *RTS3* | 0.218430152 | 0.27649181 | 0.99979869 | 11549.1162 | 13437.0119 |
| *MLP2* | 0.211861617 | 0.276562 | 0.99979869 | 28064.5768 | 32503.8734 |
| *HXT15* | 0.47488592 | 0.27691281 | 0.99979869 | 272.168547 | 378.26193 |

| | | | | | |
|---|---|---|---|---|---|
| *YET3* | 0.212255482 | 0.27711515 | 0.99979869 | 24154.7147 | 27983.182 |
| *ASG1* | 0.219993822 | 0.27729787 | 0.99979869 | 9675.15287 | 11268.9252 |
| *PIR1* | 0.207454267 | 0.27740422 | 0.99979869 | 161241.817 | 186177.652 |
| *21S_RRNA* | 1.474005834 | 0.2776468 | 0.99979869 | 4737.09844 | 13159.2097 |
| *MPC3* | 1.467942122 | 0.27775685 | 0.99979869 | 883.816143 | 2444.8637 |
| *PIN3* | 0.212363708 | 0.27788077 | 0.99979869 | 21718.855 | 25163.1317 |
| *RGA1* | 0.216117066 | 0.27789155 | 0.99979869 | 13242.6094 | 15382.6518 |
| *LAM4* | 0.218046532 | 0.27827093 | 0.99979869 | 10768.7046 | 12525.6979 |
| *MEO1* | 0.210389609 | 0.27835723 | 0.99979869 | 30172.6637 | 34909.7833 |
| *YOR114W* | 0.338799456 | 0.27857809 | 0.99979869 | 718.954191 | 909.263773 |
| *NTE1* | 0.216124158 | 0.27880446 | 0.99979869 | 12455.3692 | 14468.2626 |
| *YMR31* | 0.230378638 | 0.27881757 | 0.99979869 | 5045.36103 | 5918.92787 |
| *YGR174W-A* | 0.350602608 | 0.27916039 | 0.99979869 | 639.937516 | 815.979665 |
| *LAM1* | 0.221944836 | 0.27922178 | 0.99979869 | 7714.36871 | 8997.30342 |
| *SLD5* | 0.25796248 | 0.27965688 | 0.99979869 | 2185.15249 | 2612.98011 |
| *MPC1* | 0.223099773 | 0.28055161 | 0.99979869 | 6824.69948 | 7966.05274 |
| *YOR343C* | 0.400769715 | 0.28066364 | 0.99979869 | 422.397781 | 557.654444 |
| *EMP46* | 0.251235853 | 0.28105562 | 0.99979869 | 2485.61096 | 2958.43884 |
| *PRY1* | 0.208619089 | 0.28106641 | 0.99979869 | 30581.4043 | 35339.3002 |
| *ADR1* | 1.404733089 | 0.28108329 | 0.99979869 | 5673.59237 | 15021.8166 |
| *YAP1802* | 0.216705933 | 0.28111126 | 0.99979869 | 10311.1883 | 11982.3949 |
| *YER053C-A* | 1.401188307 | 0.28118964 | 0.99979869 | 2253.43851 | 5951.73107 |
| *PER33* | 0.215100538 | 0.28126966 | 0.99979869 | 11784.2152 | 13678.9355 |
| *FYV10* | 0.234185252 | 0.28135475 | 0.99979869 | 4054.23829 | 4768.76558 |
| *PGD1* | 0.219060647 | 0.2822896 | 0.99979869 | 8163.10539 | 9501.65266 |
| *APC9* | 0.227906519 | 0.28245884 | 0.99979869 | 5044.38551 | 5907.65177 |
| *PCL8* | 0.23199448 | 0.28256538 | 0.99979869 | 4262.02288 | 5005.5637 |
| *YOR034C-A* | 0.401601361 | 0.28258235 | 0.99979869 | 413.618151 | 546.378344 |
| *PIG1* | 0.256373661 | 0.28284205 | 0.99979869 | 2148.08294 | 2565.8255 |
| *YLR030W* | 0.278505868 | 0.2831324 | 0.99979869 | 1429.12875 | 1733.44424 |
| *YFL015C* | 0.571959422 | 0.28412285 | 0.99979869 | 159.984379 | 237.823219 |
| *YFL067W* | 0.293216327 | 0.2842528 | 0.99979869 | 1139.40094 | 1396.18631 |
| *SYP1* | 0.209352432 | 0.28427763 | 0.99979869 | 18441.1262 | 21321.0566 |
| *MSN4* | 1.336445836 | 0.28433902 | 0.99979869 | 5940.88334 | 15002.3397 |
| *YNG1* | 0.244590067 | 0.28484409 | 0.99979869 | 2740.22025 | 3246.49196 |
| *SPO13* | 0.3809103 | 0.28492728 | 0.99979869 | 473.124535 | 616.085149 |
| *ATP16* | 0.209319526 | 0.28526511 | 0.99979869 | 16985.6586 | 19637.8423 |
| *HSP12* | 1.314019767 | 0.28536329 | 0.99979869 | 4511.75459 | 11217.6702 |
| *RNY1* | 0.229193309 | 0.28575061 | 0.99979869 | 4367.37844 | 5119.34981 |
| *ATG13* | 0.21713942 | 0.28615206 | 0.99979869 | 7853.86729 | 9129.54133 |
| *YHL026C* | 0.237502093 | 0.28629543 | 0.99979869 | 3248.46331 | 3829.77391 |
| *RAD7* | 0.219690486 | 0.28648568 | 0.99979869 | 6638.37621 | 7730.27972 |
| *RNP1* | 0.61012305 | 0.28674825 | 0.99979869 | 133.645487 | 203.994916 |
| *BUL1* | 0.210243168 | 0.28713479 | 0.99979869 | 13213.344 | 15286.2924 |
| *SRV2* | 0.203193701 | 0.28715312 | 0.99979869 | 50625.3008 | 58282.0652 |

| | | | | | |
|---|---|---|---|---|---|
| *YNL054W-B* | 1.273148917 | 0.28726328 | 0.99979869 | 10992.0975 | 26566.4937 |
| *BOI1* | 0.20682348 | 0.28738361 | 0.99979869 | 20221.4402 | 23338.4536 |
| *CIN5* | 1.262275043 | 0.2876407 | 0.99979869 | 2401.71671 | 5761.06246 |
| *YLR415C* | 0.405069494 | 0.28827972 | 0.99979869 | 386.303744 | 511.524941 |
| *SOD1* | 0.200472481 | 0.28847075 | 0.99979869 | 208412.821 | 239481.831 |
| *HXT4* | 1.244848186 | 0.28852826 | 0.99979869 | 12869.9629 | 30500.8278 |
| *SHC1* | 0.34995646 | 0.28888314 | 0.99979869 | 591.161791 | 753.44856 |
| *CYS3* | 0.200420432 | 0.28940392 | 0.99979869 | 114678.559 | 131769.44 |
| *PAM1* | 0.204711663 | 0.28961878 | 0.99979869 | 22978.2442 | 26481.4104 |
| *YDR210W-D* | 0.238768136 | 0.28990646 | 0.99979869 | 2912.88632 | 3437.16057 |
| *MCH2* | 0.431654582 | 0.29001267 | 0.99979869 | 319.968758 | 431.567134 |
| *FUN14* | 0.215224971 | 0.29018618 | 0.99979869 | 7526.09441 | 8736.928 |
| *PEX18* | 0.322475965 | 0.29026185 | 0.99979869 | 759.9258 | 950.267776 |
| *SOM1* | 0.291692499 | 0.2907258 | 0.99979869 | 1082.8211 | 1325.45441 |
| *YBR221W-A* | 0.882769314 | 0.29098973 | 0.99979869 | 51.7022688 | 95.3343076 |
| *PRD1* | 0.202748939 | 0.2910101 | 0.99979869 | 28911.3234 | 33273.7236 |
| *MTH1* | 1.164173779 | 0.29193719 | 0.99979869 | 4854.16018 | 10878.3621 |
| *FUS3* | 0.209150834 | 0.29207911 | 0.99979869 | 10886.7419 | 12585.1537 |
| *PET122* | 0.223717018 | 0.29217368 | 0.99979869 | 4585.89369 | 5355.12283 |
| *AVT6* | 0.209083682 | 0.2934705 | 0.99979869 | 10160.9591 | 11745.5967 |
| *SPT23* | 0.206502229 | 0.2935732 | 0.99979869 | 12901.1793 | 14886.5034 |
| *HXK1* | 1.118654775 | 0.29387973 | 0.99979869 | 31205.7335 | 67761.1656 |
| *ALD4* | 1.108031617 | 0.29428899 | 0.99979869 | 14526.3865 | 31311.682 |
| *SAF1* | 0.213979333 | 0.29463162 | 0.99979869 | 6843.23426 | 7937.34993 |
| *SSE2* | 1.09073484 | 0.29498409 | 0.99979869 | 19042.0432 | 40556.0345 |
| *SAE3* | 0.86002684 | 0.29576488 | 0.99979869 | 53.6532978 | 97.3845078 |
| *CWP1* | 1.061934261 | 0.29610516 | 0.99979869 | 17648.0329 | 36844.1471 |
| *AGA1* | 1.057496964 | 0.29627192 | 0.99979869 | 15664.8119 | 32603.3081 |
| *DSC2* | 0.21256556 | 0.29707411 | 0.99979869 | 6801.28714 | 7880.96943 |
| *AIM7* | 0.207569189 | 0.29722448 | 0.99979869 | 9529.80121 | 11004.4494 |
| *SRB2* | 0.206879528 | 0.29754273 | 0.99979869 | 9927.81113 | 11458.5687 |
| *YPR053C* | 0.223532897 | 0.29755702 | 0.99979869 | 4029.85042 | 4705.20938 |
| *SSK1* | 0.203866165 | 0.29757659 | 0.99979869 | 13221.1481 | 15227.8617 |
| *EDS1* | 0.235932536 | 0.29765823 | 0.99979869 | 2723.6365 | 3207.53816 |
| *STE24* | 0.198888878 | 0.2979163 | 0.99979869 | 27439.272 | 31495.1749 |
| *DSC3* | 0.214622899 | 0.29791864 | 0.99979869 | 5894.05865 | 6839.46775 |
| *VID28* | 0.207092454 | 0.29793752 | 0.99979869 | 9564.91973 | 11041.353 |
| *AHA1* | 0.198159606 | 0.29795486 | 0.99979869 | 32538.2864 | 37329.0195 |
| *GID7* | 0.209237811 | 0.29795717 | 0.99979869 | 8132.86444 | 9402.21795 |
| *CYR1* | 0.198676388 | 0.29802752 | 0.99979869 | 28366.0108 | 32554.1033 |
| *SWI3* | 0.204230942 | 0.29810148 | 0.99979869 | 12302.2134 | 14173.0337 |
| *ROT2* | 0.200633078 | 0.29832058 | 0.99979869 | 19017.6553 | 21855.1337 |
| *UBC5* | 0.229832027 | 0.29856138 | 0.99979869 | 3187.98141 | 3738.54 |
| *MSA2* | 0.238108796 | 0.29870224 | 0.99979869 | 2526.58257 | 2979.96594 |
| *YGR053C* | 0.253388734 | 0.29879474 | 0.99979869 | 1800.79978 | 2146.55957 |

| | | | | | |
|---|---|---|---|---|---|
| *LAS17* | 0.20091848 | 0.29903918 | 0.99979869 | 17123.2061 | 19681.9216 |
| *GAC1* | 0.218237832 | 0.29930045 | 0.99979869 | 4766.36388 | 5544.76634 |
| *RGT1* | 0.204042189 | 0.29950887 | 0.99979869 | 11529.6059 | 13281.1967 |
| *YHR050W-A* | 0.275518428 | 0.3002313 | 0.99979869 | 1220.36865 | 1477.16922 |
| *OSH2* | 0.197219159 | 0.30060629 | 0.99979869 | 28120.1812 | 32239.3976 |
| *QCR8* | 0.20127764 | 0.30073765 | 0.99979869 | 14357.6225 | 16507.1866 |
| *SSD1* | 0.195560408 | 0.30075205 | 0.99979869 | 43877.667 | 50247.3307 |
| *YMR103C* | 0.21523973 | 0.30084256 | 0.99979869 | 5224.85569 | 6065.51718 |
| *AAD15* | 1.233802429 | 0.30093961 | 0.99979869 | 20.4858046 | 48.1797038 |
| *RUF21* | 0.305313708 | 0.30112287 | 0.99979869 | 826.260787 | 1020.99968 |
| *NBP1* | 0.236095416 | 0.30130412 | 0.99979869 | 2543.16632 | 2995.34244 |
| *SLS1* | 0.221511494 | 0.3013303 | 0.99979869 | 3971.31955 | 4630.37707 |
| *ERR3* | 1.607582746 | 0.30191472 | 0.99979869 | 9.75514506 | 29.7279024 |
| *CMP2* | 0.198911326 | 0.30202124 | 0.99979869 | 17796.3111 | 20427.1693 |
| *YJL225C* | 0.290012864 | 0.30249189 | 0.99979869 | 978.44105 | 1196.2918 |
| *MET13* | 0.209830384 | 0.30317725 | 0.99979869 | 6418.88545 | 7423.77479 |
| *RAD51* | 0.20991462 | 0.30318865 | 0.99979869 | 6385.71796 | 7385.84609 |
| *PAI3* | 0.272212277 | 0.304432 | 0.99979869 | 1222.31968 | 1476.14412 |
| *ODC1* | 0.219610912 | 0.30492927 | 0.99979869 | 3916.69074 | 4560.67026 |
| *KIP1* | 0.209611537 | 0.30569241 | 0.99979869 | 5973.07532 | 6907.12435 |
| *RIM15* | 0.201006312 | 0.30584088 | 0.99979869 | 10804.7987 | 12420.1126 |
| *MGT1* | 0.228967732 | 0.30632637 | 0.99979869 | 2817.28589 | 3301.84736 |
| *YLL058W* | 0.205865261 | 0.30633293 | 0.99979869 | 7277.33822 | 8393.51947 |
| *MSC2* | 0.208118705 | 0.30658374 | 0.99979869 | 6296.94614 | 7274.11018 |
| *KDX1* | 0.244972711 | 0.30663746 | 0.99979869 | 1909.08189 | 2262.39588 |
| *YCR099C* | 0.293698059 | 0.30673873 | 0.99979869 | 896.497831 | 1098.90729 |
| *SEC61* | 0.191392764 | 0.30697964 | 0.99979869 | 57180.7583 | 65292.7246 |
| *PMA1* | 0.189346286 | 0.30700811 | 0.99979869 | 990988.117 | 1129970.9 |
| *VAB2* | 0.229859391 | 0.3071503 | 0.99979869 | 2708.02827 | 3175.76005 |
| *SUR7* | 0.193454064 | 0.30722252 | 0.99979869 | 28130.9118 | 32167.6406 |
| *ALP1* | 0.3476369 | 0.30732133 | 0.99979869 | 517.998203 | 659.139353 |
| *NGR1* | 0.196772277 | 0.30787827 | 0.99979869 | 15048.2868 | 17247.3089 |
| *YPS6* | 0.233652234 | 0.30789797 | 0.99979869 | 2411.47186 | 2835.42683 |
| *SEC14* | 0.195626792 | 0.30800206 | 0.99979869 | 17430.4932 | 19961.7739 |
| *SHH4* | 0.267288176 | 0.30961978 | 0.99979869 | 1241.82997 | 1494.59592 |
| *SRX1* | 0.271226783 | 0.3097482 | 0.99979869 | 1169.64189 | 1411.56281 |
| *NPR3* | 0.199271272 | 0.30999131 | 0.99979869 | 10178.5184 | 11686.1409 |
| *MUB1* | 0.204125274 | 0.31044541 | 0.99979869 | 6972.00218 | 8031.65914 |
| *CYC8* | 0.190383334 | 0.311484 | 0.99979869 | 33660.128 | 38408.4499 |
| *GPI10* | 0.201606078 | 0.31216841 | 0.99979869 | 7702.66254 | 8857.88981 |
| *MDJ1* | 0.190032373 | 0.31243977 | 0.99979869 | 32100.2803 | 36619.6502 |
| *TCB1* | 0.189259809 | 0.31262483 | 0.99979869 | 39134.7154 | 44620.5564 |
| *PKH1* | 0.200171125 | 0.31283726 | 0.99979869 | 8210.9056 | 9432.97095 |
| *YDR018C* | 0.296884767 | 0.31303563 | 0.99979869 | 813.579098 | 999.47258 |
| *HAA1* | 0.192414388 | 0.3131717 | 0.99979869 | 18304.5542 | 20916.1421 |

| | | | | | |
|---|---|---|---|---|---|
| *EGH1* | 0.210223374 | 0.31355762 | 0.99979869 | 4621.01222 | 5345.89693 |
| *YBR056W* | 0.191959623 | 0.3143195 | 0.99979869 | 17435.3708 | 19916.6695 |
| *IMA1* | 0.233704419 | 0.31493141 | 0.99979869 | 2164.66669 | 2545.3235 |
| *FDH2* | 0.403976117 | 0.31508937 | 0.99979869 | 319.968758 | 423.366334 |
| *TUP1* | 0.187673817 | 0.31548694 | 0.99979869 | 41715.9268 | 47511.3386 |
| *GRE1* | 0.277793645 | 0.31575462 | 0.99979869 | 1002.82891 | 1215.7687 |
| *PSK1* | 0.1922095 | 0.31633219 | 0.99979869 | 14747.8283 | 16849.57 |
| *CSI1* | 0.259553969 | 0.31646296 | 0.99979869 | 1296.45878 | 1552.00152 |
| *PDC5* | 0.21546202 | 0.31658365 | 0.99979869 | 3514.77877 | 4080.92343 |
| *tR(UCU)E* | 1.319458351 | 0.31676088 | 0.99979869 | 15.6082321 | 38.9538031 |
| *CWC21* | 0.200824658 | 0.31693921 | 0.99979869 | 6783.72788 | 7796.91122 |
| *RGM1* | 0.207359874 | 0.31741851 | 0.99979869 | 4742.95153 | 5476.08464 |
| *TFB6* | 0.207836209 | 0.31780161 | 0.99979869 | 4597.59987 | 5310.01842 |
| *MLF3* | 0.189288488 | 0.31797294 | 0.99979869 | 19866.3529 | 22651.6365 |
| *TGL4* | 0.188176981 | 0.31813723 | 0.99979869 | 24048.3836 | 27398.875 |
| *MBF1* | 0.184032063 | 0.31845646 | 0.99979869 | 124035.694 | 140911.282 |
| *PCD1* | 0.22738999 | 0.31897721 | 0.99979869 | 2368.54922 | 2772.89572 |
| *TPK2* | 0.191699872 | 0.31899271 | 0.99979869 | 13150.9111 | 15019.7664 |
| *ATG19* | 0.197403145 | 0.32015946 | 0.99979869 | 7532.92302 | 8637.49329 |
| *SOH1* | 0.206872062 | 0.32133984 | 0.99979869 | 4383.96219 | 5059.894 |
| *YBR184W* | 0.278582409 | 0.32176783 | 0.99979869 | 935.518411 | 1134.78579 |
| *FEX1* | 0.269883301 | 0.32224061 | 0.99979869 | 1045.75155 | 1260.8731 |
| *SNG1* | 0.20505514 | 0.32224922 | 0.99979869 | 4637.59596 | 5345.89693 |
| *RNQ1* | 0.185379709 | 0.3223463 | 0.99979869 | 27158.3239 | 30882.1651 |
| *DPM1* | 0.183714369 | 0.32268786 | 0.99979869 | 41474.9747 | 47107.4492 |
| *FCP1* | 0.18669271 | 0.3227607 | 0.99979869 | 19980.4881 | 22740.8202 |
| *RRD2* | 0.206737954 | 0.32294995 | 0.99979869 | 4239.58604 | 4892.80269 |
| *FUN12* | 0.181329991 | 0.32327884 | 0.99979869 | 153373.317 | 173914.379 |
| *HOF1* | 0.190281871 | 0.32410597 | 0.99979869 | 11358.8909 | 12960.3403 |
| *PMT4* | 0.181989474 | 0.32563625 | 0.99979869 | 44416.151 | 50387.7694 |
| *RUF5-2* | 0.310811122 | 0.32574344 | 0.99979869 | 627.255827 | 778.050962 |
| *NST1* | 0.182415823 | 0.32577674 | 0.99979869 | 37315.3809 | 42344.8342 |
| *ALT1* | 0.185804279 | 0.32580405 | 0.99979869 | 17739.7313 | 20178.07 |
| *FKS3* | 0.216179292 | 0.32643015 | 0.99979869 | 2839.72273 | 3298.77206 |
| *KGD2* | 0.184563807 | 0.32678445 | 0.99979869 | 20000.9739 | 22730.5692 |
| *GIC1* | 0.237105886 | 0.32735926 | 0.99979869 | 1692.51767 | 1994.84476 |
| *YGR025W* | 0.244090682 | 0.32744882 | 0.99979869 | 1481.80653 | 1754.97134 |
| *FLO1* | 0.214789265 | 0.32747411 | 0.99979869 | 2906.05771 | 3372.57927 |
| *SSH4* | 0.204288718 | 0.32802418 | 0.99979869 | 4159.59385 | 4792.34288 |
| *AVO2* | 0.212118425 | 0.32837815 | 0.99979869 | 3107.0137 | 3599.12639 |
| *YJL163C* | 0.20287279 | 0.32864631 | 0.99979869 | 4349.81918 | 5006.5888 |
| *SDS22* | 0.19413378 | 0.32881168 | 0.99979869 | 6818.8464 | 7801.01162 |
| *MNN4* | 0.188850128 | 0.32883135 | 0.99979869 | 10213.6369 | 11642.0616 |
| *PET9* | 0.179777389 | 0.32927025 | 0.99979869 | 51205.7319 | 58001.1877 |
| *UBP7* | 0.190072669 | 0.32989804 | 0.99979869 | 8748.41409 | 9980.3744 |

| | | | | | |
|---|---|---|---|---|---|
| *OKP1* | 0.207337453 | 0.32990801 | 0.99979869 | 3547.94626 | 4096.29993 |
| *YGL194C-A* | 0.236177142 | 0.3302112 | 0.99979869 | 1662.27672 | 1957.94116 |
| *COX12* | 0.197039365 | 0.33022632 | 0.99979869 | 5505.80387 | 6311.5412 |
| *tS(AGA)D2* | 1.135661455 | 0.33136281 | 0.99979869 | 21.4613191 | 47.1546038 |
| *PDA1* | 0.177557599 | 0.3314008 | 0.99979869 | 103229.921 | 116749.673 |
| *TEA1* | 0.190749159 | 0.33427617 | 0.99979869 | 7020.7779 | 8013.20734 |
| *BRP1* | 0.375206511 | 0.33434264 | 0.99979869 | 341.430077 | 442.843235 |
| *YPR172W* | 0.190729764 | 0.33439925 | 0.99979869 | 6999.31658 | 7988.60494 |
| *SOR1* | 0.650955043 | 0.33482356 | 0.99979869 | 84.869762 | 133.263011 |
| *FAB1* | 0.181679947 | 0.33527364 | 0.99979869 | 15799.4329 | 17919.7745 |
| *YLR326W* | 0.214179104 | 0.33537763 | 0.99979869 | 2581.21138 | 2994.31734 |
| *YCP4* | 0.176636341 | 0.33703476 | 0.99979869 | 38207.9767 | 43184.3911 |
| *GLO4* | 0.203578057 | 0.33718313 | 0.99979869 | 3501.12156 | 4031.71862 |
| *STE12* | 0.18327966 | 0.3372893 | 0.99979869 | 11458.3934 | 13010.5702 |
| *YPI1* | 0.185211639 | 0.33764977 | 0.99979869 | 9352.25757 | 10633.3631 |
| *CSI2* | 0.195174961 | 0.33773936 | 0.99979869 | 4916.59311 | 5628.82455 |
| *RCK1* | 0.332057583 | 0.33777819 | 0.99979869 | 468.246963 | 589.432547 |
| *MRH1* | 0.17364479 | 0.33800577 | 0.99979869 | 241757.858 | 272679.697 |
| *URA2* | 0.174937901 | 0.33805658 | 0.99979869 | 63854.253 | 72086.0629 |
| *YGR109W-B* | 0.193949055 | 0.33896758 | 0.99979869 | 5056.09169 | 5783.61466 |
| *SGO1* | 0.198998475 | 0.33930379 | 0.99979869 | 3991.80536 | 4582.19737 |
| *TDA3* | 0.177924875 | 0.33934218 | 0.99979869 | 20877.9615 | 23618.3059 |
| *SEC24* | 0.174573244 | 0.33967815 | 0.99979869 | 51399.8593 | 58011.4387 |
| *COQ9* | 0.189580756 | 0.33968366 | 0.99979869 | 6303.77474 | 7189.02687 |
| *YMR007W* | 0.328627409 | 0.3398454 | 0.99979869 | 475.075564 | 596.608248 |
| *YLR446W* | 0.202810763 | 0.34063384 | 0.99979869 | 3358.69644 | 3865.65241 |
| *AST2* | 0.200712945 | 0.3406648 | 0.99979869 | 3624.03639 | 4164.98163 |
| *ATG1* | 0.193605796 | 0.34083738 | 0.99979869 | 4894.15628 | 5597.04645 |
| *AIM9* | 0.176603481 | 0.3410214 | 0.99979869 | 22996.779 | 25991.4126 |
| *COS9* | 0.191179323 | 0.34169006 | 0.99979869 | 5415.08102 | 6182.37859 |
| *SCD5* | 0.178371448 | 0.34255107 | 0.99979869 | 15038.5316 | 17017.6865 |
| *YKL063C* | 0.184767492 | 0.34276319 | 0.99979869 | 7837.28354 | 8908.11971 |
| *RAD2* | 0.181307546 | 0.3428147 | 0.99979869 | 10524.826 | 11934.2152 |
| *CBP2* | 0.190523234 | 0.34295581 | 0.99979869 | 5412.15448 | 6176.22799 |
| *EST3* | 0.203341077 | 0.3433547 | 0.99979869 | 3130.42605 | 3604.25189 |
| *YOR292C* | 0.199139859 | 0.34375744 | 0.99979869 | 3607.45264 | 4141.40433 |
| *ATG26* | 0.176975545 | 0.34466178 | 0.99979869 | 15811.1391 | 17874.6701 |
| *EXG1* | 0.171005405 | 0.34467979 | 0.99979869 | 107768.014 | 121329.821 |
| *RSF1* | 0.212674973 | 0.34481831 | 0.99979869 | 2315.87144 | 2683.71201 |
| *RTK1* | 0.181240808 | 0.34506237 | 0.99979869 | 9519.07055 | 10793.2788 |
| *ARC18* | 0.176752482 | 0.34566188 | 0.99979869 | 15214.1242 | 17197.079 |
| *YGR111W* | 0.181782695 | 0.34639817 | 0.99979869 | 8573.79699 | 9725.12448 |
| *YOR020W-A* | 0.189954805 | 0.34647292 | 0.99979869 | 5062.92029 | 5775.41386 |
| *PIB1* | 0.194684334 | 0.34664227 | 0.99979869 | 4050.33623 | 4635.50257 |
| *PIG2* | 0.183748479 | 0.34673385 | 0.99979869 | 7284.16682 | 8273.58276 |

| | | | | | |
|---|---|---|---|---|---|
| *COQ4* | 0.17964739 | 0.34713004 | 0.99979869 | 10028.2891 | 11358.1089 |
| *MEK1* | 0.344672008 | 0.34734269 | 0.99979869 | 393.132346 | 499.22374 |
| *ARD1* | 0.175940497 | 0.34750344 | 0.99979869 | 15024.8744 | 16973.6072 |
| *AHT1* | 0.435583855 | 0.34846247 | 0.99979869 | 210.711133 | 284.977823 |
| *TMN2* | 0.185786956 | 0.34937847 | 0.99979869 | 5858.94012 | 6664.17563 |
| *HXT17* | 0.257807102 | 0.35019518 | 0.99979869 | 941.371498 | 1125.55989 |
| *YER078W-A* | 0.725756079 | 0.35037617 | 0.99979869 | 59.5063849 | 98.4096079 |
| *PSP1* | 0.17542193 | 0.3506106 | 0.99979869 | 13182.1275 | 14886.5034 |
| *RME1* | 0.173383809 | 0.35167471 | 0.99979869 | 16255.9737 | 18331.8648 |
| *YOR342C* | 0.17711399 | 0.35191695 | 0.99979869 | 10200.9552 | 11533.401 |
| *SHP1* | 0.169219708 | 0.35256837 | 0.99979869 | 37233.4377 | 41867.1375 |
| *MSL5* | 0.178457173 | 0.3540771 | 0.99979869 | 8242.12206 | 9327.38564 |
| *YGR126W* | 0.213328129 | 0.35409341 | 0.99979869 | 1992.97614 | 2310.57558 |
| *YBR027C* | 0.501214727 | 0.35421836 | 0.99979869 | 143.400632 | 202.969816 |
| *COX5B* | 0.176604563 | 0.35422411 | 0.99979869 | 9619.54854 | 10872.2115 |
| *UBX4* | 0.178710283 | 0.35446561 | 0.99979869 | 7956.29631 | 9005.50422 |
| *YLR031W* | 0.266808453 | 0.35497891 | 0.99979869 | 800.89741 | 963.594077 |
| *EGT2* | 0.165680973 | 0.35565213 | 0.99979869 | 131332.542 | 147315.083 |
| *CTH1* | 0.174017573 | 0.35574749 | 0.99979869 | 11620.3288 | 13110.0049 |
| *ECM4* | 0.196833389 | 0.35592161 | 0.99979869 | 3096.28304 | 3548.89648 |
| *YGR237C* | 0.172105716 | 0.35648381 | 0.99979869 | 14064.9681 | 15847.0222 |
| *PEX30* | 0.172754926 | 0.3567852 | 0.99979869 | 12709.9785 | 14326.7987 |
| *TFB2* | 0.181413555 | 0.35680536 | 0.99979869 | 6089.16155 | 6905.07415 |
| *RIB2* | 0.177583967 | 0.35681995 | 0.99979869 | 7941.66359 | 8981.92692 |
| *HST2* | 0.170564481 | 0.3571241 | 0.99979869 | 16858.8417 | 18974.6025 |
| *RAX1* | 0.174257492 | 0.35779697 | 0.99979869 | 10217.5389 | 11529.3006 |
| *BLS1* | 0.250475791 | 0.35822914 | 0.99979869 | 967.71039 | 1151.18739 |
| *PEX22* | 0.186688135 | 0.35829203 | 0.99979869 | 4414.20314 | 5024.0155 |
| *NET1* | 0.166352085 | 0.35870949 | 0.99979869 | 37464.6346 | 42043.4548 |
| *GNA1* | 0.177595566 | 0.35893009 | 0.99979869 | 7339.77114 | 8301.26046 |
| *DSE3* | 0.170662963 | 0.35911906 | 0.99979869 | 14455.174 | 16270.3885 |
| *YKL162C* | 0.215054142 | 0.35957405 | 0.99979869 | 1781.28949 | 2067.62686 |
| *LAS21* | 0.171163039 | 0.36045008 | 0.99979869 | 12497.3163 | 14071.5488 |
| *YDL022C-A* | 1.10715508 | 0.36054148 | 0.99979869 | 19.5102901 | 42.0291034 |
| *PDR17* | 0.17455149 | 0.36056638 | 0.99979869 | 8790.36121 | 9920.91859 |
| *STI1* | 0.16341329 | 0.36113017 | 0.99979869 | 103365.517 | 115762.502 |
| *SIP5* | 0.171938139 | 0.36124884 | 0.99979869 | 10962.832 | 12350.4058 |
| *ICR1* | 0.196001399 | 0.36157943 | 0.99979869 | 2879.71882 | 3298.77206 |
| *POL2* | 0.168871411 | 0.36161458 | 0.99979869 | 15834.5515 | 17800.8629 |
| *MSN2* | 0.172689859 | 0.36183311 | 0.99979869 | 9871.23129 | 11126.4363 |
| *PEP7* | 0.198070248 | 0.36214908 | 0.99979869 | 2670.95872 | 3064.02414 |
| *TDP1* | 0.184892793 | 0.36242389 | 0.99979869 | 4355.67227 | 4951.2334 |
| *RMD5* | 0.17944956 | 0.3630151 | 0.99979869 | 5701.88229 | 6457.10542 |
| *ART5* | 0.175253095 | 0.36329601 | 0.99979869 | 7472.44112 | 8437.59877 |
| *SMC1* | 0.171448069 | 0.36356908 | 0.99979869 | 10297.5311 | 11596.9572 |

| | | | | | |
|---|---|---|---|---|---|
| *COX9* | 0.17024782 | 0.36359148 | 0.99979869 | 11708.1251 | 13174.5863 |
| *SUR1* | 0.166768491 | 0.36396535 | 0.99979869 | 18740.6092 | 21037.1039 |
| *YEL068C* | 0.193280463 | 0.36418617 | 0.99979869 | 3013.36431 | 3445.36137 |
| *GPI12* | 0.180419762 | 0.36443199 | 0.99979869 | 5188.76166 | 5879.97407 |
| *PMR1* | 0.165335419 | 0.3644765 | 0.99979869 | 23871.8155 | 26770.4886 |
| *AZR1* | 0.191031357 | 0.3652788 | 0.99979869 | 3196.76104 | 3649.35629 |
| *TOP2* | 0.165508439 | 0.36588894 | 0.99979869 | 20130.7173 | 22577.8293 |
| *MMR1* | 0.165356396 | 0.36674556 | 0.99979869 | 19249.8277 | 21587.5826 |
| *CMK1* | 0.167275196 | 0.36773224 | 0.99979869 | 13335.2833 | 14974.662 |
| *CDC55* | 0.167911121 | 0.36814717 | 0.99979869 | 12030.0449 | 13514.9195 |
| *NCR1* | 0.166856082 | 0.36871861 | 0.99979869 | 13281.63 | 14910.0807 |
| *MEC1* | 0.172438607 | 0.36884858 | 0.99979869 | 7580.72323 | 8543.18408 |
| *ENT2* | 0.162188329 | 0.36912488 | 0.99979869 | 31466.1959 | 35210.1376 |
| *YMR087W* | 0.192868497 | 0.36919751 | 0.99979869 | 2801.67766 | 3202.41266 |
| *SER3* | 0.17568937 | 0.36956028 | 0.99979869 | 5915.51997 | 6681.60233 |
| *SIP2* | 0.173990254 | 0.37035904 | 0.99979869 | 6441.32228 | 7266.93448 |
| *tD(GUC)I1* | 1.153993276 | 0.37045477 | 0.99979869 | 16.5837466 | 36.9036029 |
| *NVJ1* | 0.166869339 | 0.3707512 | 0.99979869 | 11823.2358 | 13272.9959 |
| *BUD4* | 0.163034488 | 0.37091578 | 0.99979869 | 21005.7539 | 23518.8712 |
| *SUM1* | 0.163332053 | 0.37100121 | 0.99979869 | 19658.5683 | 22015.0494 |
| *HAL1* | 0.209548161 | 0.37123114 | 0.99979869 | 1739.34236 | 2011.24636 |
| *CRG1* | 0.184370045 | 0.37152789 | 0.99979869 | 3657.20388 | 4155.75573 |
| *ACA1* | 0.176583262 | 0.37161744 | 0.99979869 | 5295.09274 | 5984.53428 |
| *FMP46* | 0.173673824 | 0.3717955 | 0.99979869 | 6289.14202 | 7093.69257 |
| *PAT1* | 0.161005312 | 0.37205796 | 0.99979869 | 30416.5423 | 34007.6952 |
| *ATG8* | 0.180162613 | 0.37229454 | 0.99979869 | 4328.35786 | 4904.07879 |
| *ZPR1* | 0.159570591 | 0.37232475 | 0.99979869 | 49406.8832 | 55185.2378 |
| *COA2* | 0.184436509 | 0.37241246 | 0.99979869 | 3583.06478 | 4071.69752 |
| *NRD1* | 0.166158917 | 0.37272808 | 0.99979869 | 11569.602 | 12981.8674 |
| *tK(CUU)M* | 0.541751014 | 0.37286904 | 0.99979869 | 106.331081 | 154.790112 |
| *YAR064W* | 0.253851939 | 0.37370325 | 0.99979869 | 803.823953 | 958.468576 |
| *SWT21* | 0.184617078 | 0.37403982 | 0.99979869 | 3445.51724 | 3915.88231 |
| *AFG3* | 0.162766418 | 0.37423781 | 0.99979869 | 16825.6742 | 18835.1889 |
| *YDR461C-A* | 0.190333242 | 0.37427254 | 0.99979869 | 2795.82457 | 3190.11145 |
| *JHD2* | 0.204531947 | 0.37432235 | 0.99979869 | 1878.84094 | 2165.01137 |
| *OXR1* | 0.169176941 | 0.37458605 | 0.99979869 | 7954.34528 | 8943.99821 |
| *YGR054W* | 0.15996156 | 0.37466497 | 0.99979869 | 29552.2364 | 33017.4485 |
| *FCY21* | 0.167873635 | 0.37478925 | 0.99979869 | 8828.40628 | 9917.84329 |
| *GCN1* | 0.158042123 | 0.37513217 | 0.99979869 | 57851.9123 | 64549.5271 |
| *KAR4* | 0.180089647 | 0.37562388 | 0.99979869 | 4031.80145 | 4567.84596 |
| *DDC1* | 0.18693591 | 0.3760432 | 0.99979869 | 3045.55629 | 3466.88848 |
| *MLP1* | 0.158871844 | 0.3762054 | 0.99979869 | 33589.891 | 37500.2112 |
| *CAP2* | 0.160791642 | 0.37620826 | 0.99979869 | 20602.8664 | 23031.9486 |
| *RHO5* | 0.165300077 | 0.37766086 | 0.99979869 | 9979.5134 | 11191.0176 |
| *DDI3* | 0.344853791 | 0.37811999 | 0.99979869 | 318.017729 | 403.889432 |

| | | | | | |
|---|---|---|---|---|---|
| *MSL1* | 0.197005789 | 0.3789497 | 0.99979869 | 2136.37677 | 2448.9641 |
| *NPC2* | 0.158316429 | 0.3793516 | 0.99979869 | 26573.9907 | 29656.1454 |
| *YMR147W* | 0.183409755 | 0.37950975 | 0.99979869 | 3254.31639 | 3695.48579 |
| *AIM39* | 0.175525736 | 0.38017561 | 0.99979869 | 4515.65665 | 5099.87291 |
| *DDL1* | 0.176962865 | 0.38021683 | 0.99979869 | 4207.39406 | 4756.46438 |
| *VTC5* | 0.165563462 | 0.38123282 | 0.99979869 | 8371.86549 | 9389.91675 |
| *SMK1* | 0.188479398 | 0.38132244 | 0.99979869 | 2645.59534 | 3014.81934 |
| *ACF4* | 0.178444982 | 0.3821108 | 0.99979869 | 3779.1432 | 4276.71754 |
| *DOA4* | 0.16850458 | 0.38215734 | 0.99979869 | 6454.00397 | 7253.60818 |
| *GFA1* | 0.155629521 | 0.38225489 | 0.99979869 | 42660.2249 | 47519.5394 |
| *YDL218W* | 0.233801308 | 0.38293824 | 0.99979869 | 983.318622 | 1156.31289 |
| *ARE2* | 0.157111226 | 0.38346806 | 0.99979869 | 23160.6654 | 25825.3464 |
| *MIR1* | 0.155590202 | 0.38365632 | 0.99979869 | 34616.1322 | 38558.1145 |
| *YPR084W* | 0.168847327 | 0.38373945 | 0.99979869 | 6011.12039 | 6757.45974 |
| *SEF1* | 0.167653868 | 0.38395548 | 0.99979869 | 6470.58772 | 7267.95958 |
| *YLR285C-A* | 0.194105352 | 0.38426174 | 0.99979869 | 2145.1564 | 2454.0896 |
| *YMR315W* | 0.155784569 | 0.38453661 | 0.99979869 | 28986.438 | 32291.6777 |
| *PIB2* | 0.156954356 | 0.385413 | 0.99979869 | 20038.0435 | 22341.0312 |
| *BFR1* | 0.153761643 | 0.3855017 | 0.99979869 | 53985.9483 | 60057.5385 |
| *RRT13* | 0.233926338 | 0.38555894 | 0.99979869 | 957.955245 | 1126.58499 |
| *WHI3* | 0.160107999 | 0.38557219 | 0.99979869 | 12093.4533 | 13512.8693 |
| *JEN1* | 0.184672516 | 0.38687705 | 0.99979869 | 2738.26922 | 3112.20385 |
| *AGX1* | 0.189508076 | 0.38739922 | 0.99979869 | 2335.38173 | 2663.21001 |
| *GMC1* | 0.172666484 | 0.38749333 | 0.99979869 | 4392.74182 | 4951.2334 |
| *VAC8* | 0.156343162 | 0.38809193 | 0.99979869 | 18108.4758 | 20181.1453 |
| *ATP18* | 0.160742964 | 0.38822415 | 0.99979869 | 9879.0354 | 11043.4032 |
| *DIT2* | 0.281793922 | 0.38872312 | 0.99979869 | 516.047174 | 627.36125 |
| *HUL4* | 0.178938848 | 0.38873756 | 0.99979869 | 3256.26742 | 3686.25989 |
| *BNA3* | 0.159747967 | 0.38894937 | 0.99979869 | 10628.2305 | 11872.7091 |
| *ATG34* | 0.194357802 | 0.38898057 | 0.99979869 | 1997.85371 | 2285.97318 |
| *PRP40* | 0.1596685 | 0.38908407 | 0.99979869 | 10653.5939 | 11900.3868 |
| *YLR049C* | 0.161197356 | 0.38952958 | 0.99979869 | 8929.85979 | 9985.4999 |
| *EAF1* | 0.158049357 | 0.38955162 | 0.99979869 | 12642.668 | 14106.4022 |
| *NAB6* | 0.152835474 | 0.38963466 | 0.99979869 | 37876.3017 | 42109.0612 |
| *POS5* | 0.165907358 | 0.3901644 | 0.99979869 | 6031.60619 | 6766.68564 |
| *VPS30* | 0.167676478 | 0.39087625 | 0.99979869 | 5294.11722 | 5946.60557 |
| *LCB5* | 0.161748944 | 0.39113205 | 0.99979869 | 7981.65969 | 8928.62171 |
| *GTT1* | 0.159621602 | 0.39145412 | 0.99979869 | 9605.89134 | 10729.7226 |
| *YPR127W* | 0.158911452 | 0.39176797 | 0.99979869 | 10201.9307 | 11389.887 |
| *FMP10* | 0.163654548 | 0.39225071 | 0.99979869 | 6613.01284 | 7407.37319 |
| *YNR071C* | 0.300125559 | 0.39233579 | 0.99979869 | 420.446752 | 517.675541 |
| *HUL5* | 0.159667565 | 0.39268312 | 0.99979869 | 9074.23594 | 10136.1896 |
| *CSH1* | 0.170111596 | 0.39272677 | 0.99979869 | 4427.86034 | 4981.9864 |
| *SIS1* | 0.149843134 | 0.39306744 | 0.99979869 | 94118.6151 | 104419.77 |
| *SHM1* | 0.150696228 | 0.39337246 | 0.99979869 | 49045.9428 | 54446.1406 |

| | | | | | |
|---|---|---|---|---|---|
| *YIL092W* | 0.170708685 | 0.39366683 | 0.99979869 | 4208.36958 | 4736.98748 |
| *MCM22* | 0.195829422 | 0.39372182 | 0.99979869 | 1810.55492 | 2073.77747 |
| *PRM2* | 0.20884491 | 0.39396212 | 0.99979869 | 1357.91619 | 1569.42823 |
| *YMR247W-A* | 0.289121451 | 0.39398508 | 0.99979869 | 461.418361 | 563.805045 |
| *FIR1* | 0.16456314 | 0.3946449 | 0.99979869 | 5784.80102 | 6483.75802 |
| *STB3* | 0.157292754 | 0.39470561 | 0.99979869 | 10623.353 | 11847.0816 |
| *ZEO1* | 0.148308113 | 0.39502199 | 0.99979869 | 310285.801 | 343880.073 |
| *YHL037C* | 0.64138651 | 0.39526227 | 0.99979869 | 62.4329284 | 97.3845078 |
| *RIM11* | 0.157294811 | 0.39600003 | 0.99979869 | 10001.9502 | 11154.114 |
| *NGG1* | 0.154502077 | 0.39703792 | 0.99979869 | 13331.3812 | 14838.3237 |
| *ASH1* | 0.150092204 | 0.3972485 | 0.99979869 | 33066.0397 | 36691.4072 |
| *YJL007C* | 0.564129212 | 0.3974 | 0.99979869 | 83.8942475 | 124.03711 |
| *UBP2* | 0.152011759 | 0.39769857 | 0.99979869 | 19282.0197 | 21424.5917 |
| *RTS2* | 0.184625981 | 0.39771442 | 0.99979869 | 2320.74901 | 2637.58251 |
| *MEI4* | 0.237240414 | 0.39799577 | 0.99979869 | 817.481156 | 963.594077 |
| *ERT1* | 0.176483018 | 0.3980681 | 0.99979869 | 3025.07048 | 3418.70877 |
| *FAR7* | 0.185280958 | 0.39823015 | 0.99979869 | 2259.2916 | 2568.9008 |
| *TKL1* | 0.147017306 | 0.39894416 | 0.99979869 | 166166.214 | 183992.138 |
| *HMG2* | 0.151167147 | 0.39900405 | 0.99979869 | 20557.9927 | 22828.9788 |
| *SPO20* | 0.3620815 | 0.39913977 | 0.99979869 | 244.854141 | 314.705725 |
| *YNL140C* | 0.217135182 | 0.39943902 | 0.99979869 | 1102.33139 | 1281.3751 |
| *RFS1* | 0.15031784 | 0.39961772 | 0.99979869 | 23566.4794 | 26154.4035 |
| *HFD1* | 0.154324342 | 0.39974474 | 0.99979869 | 11758.8519 | 13086.4276 |
| *GLO2* | 0.151620297 | 0.40015979 | 0.99979869 | 17157.3491 | 19058.6607 |
| *SNF1* | 0.151661906 | 0.40024393 | 0.99979869 | 16926.1522 | 18802.3857 |
| *SDD1* | 0.256772194 | 0.40115314 | 0.99979869 | 619.451711 | 740.122259 |
| *YER152C* | 0.149641391 | 0.40120675 | 0.99979869 | 23758.6558 | 26355.3231 |
| *SBE22* | 0.150765136 | 0.40122739 | 0.99979869 | 18557.2124 | 20601.4363 |
| *AIM18* | 0.160456178 | 0.4013112 | 0.99979869 | 6275.48482 | 7013.73476 |
| *OSH6* | 0.154733085 | 0.40139083 | 0.99979869 | 10335.5762 | 11505.7233 |
| *HAP2* | 0.163008181 | 0.40268003 | 0.99979869 | 5126.32873 | 5739.53536 |
| *YDR114C* | 0.595951784 | 0.40324286 | 0.99979869 | 71.2125589 | 107.635509 |
| *MYO1* | 0.149010973 | 0.40335461 | 0.99979869 | 22454.3929 | 24897.6308 |
| *BDH1* | 0.147785031 | 0.40359704 | 0.99979869 | 30499.461 | 33789.3489 |
| *GZF3* | 0.160569887 | 0.40360262 | 0.99979869 | 5822.84609 | 6508.36042 |
| *YPR096C* | 0.322009635 | 0.40510261 | 0.99979869 | 318.993243 | 398.763932 |
| *ATP15* | 0.152991752 | 0.40530966 | 0.99979869 | 10478.9768 | 11651.2875 |
| *YAL064W* | 0.845255416 | 0.40537129 | 0.99979869 | 30.2409497 | 54.3303043 |
| *NKP2* | 0.209033372 | 0.40633875 | 0.99979869 | 1190.1277 | 1375.68431 |
| *CHM7* | 0.165765028 | 0.40634948 | 0.99979869 | 4061.06689 | 4555.54476 |
| *ACP1* | 0.146808738 | 0.4065609 | 0.99979869 | 28665.4938 | 31736.0734 |
| *PCS60* | 0.14788403 | 0.40666862 | 0.99979869 | 21448.6374 | 23763.8701 |
| *KIN2* | 0.148848853 | 0.40757211 | 0.99979869 | 16478.391 | 18269.3337 |
| *BDP1* | 0.149834753 | 0.40826622 | 0.99979869 | 13466.9778 | 14940.8337 |
| *AFG1* | 0.148794884 | 0.40934301 | 0.99979869 | 14770.2651 | 16374.9487 |

| | | | | | |
|---|---|---|---|---|---|
| *tL(CAA)K* | 1.178446234 | 0.40976485 | 0.99979869 | 12.6816886 | 28.7028023 |
| *CLB2* | 0.153216639 | 0.41007045 | 0.99979869 | 8372.84101 | 9310.98404 |
| *YEF1* | 0.180277459 | 0.41074948 | 0.99979869 | 2192.95661 | 2484.8426 |
| *MPM1* | 0.15230217 | 0.41091479 | 0.99979869 | 8843.039 | 9827.63448 |
| *ANY1* | 0.154874779 | 0.41150683 | 0.99979869 | 6905.66719 | 7688.25061 |
| *HYR1* | 0.146337165 | 0.41199724 | 0.99979869 | 19007.9002 | 21037.1039 |
| *PET127* | 0.156498889 | 0.41210093 | 0.99979869 | 6014.04693 | 6703.12943 |
| *PRE4* | 0.143882687 | 0.41214629 | 0.99979869 | 36835.4278 | 40698.5234 |
| *YLR257W* | 0.142642319 | 0.41216435 | 0.99979869 | 70316.0611 | 77623.6535 |
| *VCX1* | 0.145336236 | 0.41244416 | 0.99979869 | 22840.6966 | 25261.5413 |
| *LCP5* | 0.149459377 | 0.41267262 | 0.99979869 | 11178.4207 | 12398.5855 |
| *YNL134C* | 0.143952547 | 0.41272258 | 0.99979869 | 33066.0397 | 36535.592 |
| *LCB1* | 0.142769956 | 0.41341118 | 0.99979869 | 48583.5489 | 53637.3367 |
| *STP3* | 0.145101567 | 0.41404691 | 0.99979869 | 20845.7695 | 23051.4255 |
| *MEI5* | 0.617812949 | 0.41430295 | 0.99979869 | 61.4574139 | 94.3092075 |
| *APE4* | 0.146247747 | 0.41430489 | 0.99979869 | 16278.4106 | 18015.1088 |
| *LAG2* | 0.16544241 | 0.41608486 | 0.99979869 | 3393.81497 | 3806.1966 |
| *HXT2* | 0.141050168 | 0.41633496 | 0.99979869 | 70128.7623 | 77331.5 |
| *IXR1* | 0.14362167 | 0.41672311 | 0.99979869 | 23131.4 | 25552.6697 |
| *PKP1* | 0.156170325 | 0.41716868 | 0.99979869 | 5343.86846 | 5954.80638 |
| *PUS5* | 0.173710183 | 0.41731231 | 0.99979869 | 2445.61487 | 2758.54432 |
| *ATO2* | 0.17058206 | 0.41738118 | 0.99979869 | 2717.78341 | 3058.89864 |
| *PAC11* | 0.157424869 | 0.41745909 | 0.99979869 | 4913.66657 | 5480.18504 |
| *CSM2* | 0.213282865 | 0.41753538 | 0.99979869 | 992.098253 | 1150.16229 |
| *MIC60* | 0.146357618 | 0.41761448 | 0.99979869 | 13010.437 | 14399.5808 |
| *CIP1* | 0.141508251 | 0.41767619 | 0.99979869 | 40786.2615 | 44989.5924 |
| *CPR6* | 0.140788563 | 0.4178564 | 0.99979869 | 56812.0138 | 62635.6652 |
| *YCR102C* | 0.210771125 | 0.41807044 | 0.99979869 | 1030.14332 | 1192.1914 |
| *UBX5* | 0.148214765 | 0.41839795 | 0.99979869 | 9899.52121 | 10970.6211 |
| *WSC3* | 0.156347343 | 0.41874926 | 0.99979869 | 5074.62646 | 5655.47715 |
| *HFM1* | 0.158690566 | 0.41927152 | 0.99979869 | 4385.91322 | 4895.87799 |
| *HSV2* | 0.152375332 | 0.41935231 | 0.99979869 | 6533.02065 | 7260.78388 |
| *RIM4* | 0.156434885 | 0.41948014 | 0.99979869 | 4956.58921 | 5524.26434 |
| *ATG9* | 0.156115482 | 0.41957573 | 0.99979869 | 5040.48345 | 5616.52335 |
| *YLR112W* | 0.174040342 | 0.41984459 | 0.99979869 | 2331.47967 | 2630.40681 |
| *PRP45* | 0.154130413 | 0.42006712 | 0.99979869 | 5645.30245 | 6281.8133 |
| *OSH3* | 0.141204646 | 0.42198838 | 0.99979869 | 25708.7093 | 28352.2181 |
| *IKS1* | 0.15105547 | 0.42223799 | 0.99979869 | 6610.08629 | 7339.71659 |
| *ALG12* | 0.143532772 | 0.42279281 | 0.99979869 | 14735.1466 | 16276.5391 |
| *SCO2* | 0.157309627 | 0.42312523 | 0.99979869 | 4331.28441 | 4830.27159 |
| *YGL258W-A* | 0.53250452 | 0.42399457 | 0.99979869 | 82.918733 | 119.93671 |
| *ECM3* | 0.144030141 | 0.42427346 | 0.99979869 | 12530.4838 | 13846.0268 |
| *CDC48* | 0.1372322 | 0.42438224 | 0.99979869 | 172724.598 | 189961.296 |
| *YBR209W* | 0.242990546 | 0.42468256 | 0.99979869 | 610.672081 | 722.695558 |
| *GYP1* | 0.149673863 | 0.42478569 | 0.99979869 | 6800.31162 | 7543.7115 |

| | | | | | |
|---|---|---|---|---|---|
| *SOL2* | 0.141483677 | 0.42502077 | 0.99979869 | 18319.1869 | 20206.7728 |
| *GPC1* | 0.151752878 | 0.42514125 | 0.99979869 | 5766.26625 | 6405.85041 |
| *CYB2* | 0.169657636 | 0.42570656 | 0.99979869 | 2467.07619 | 2774.94592 |
| *MHT1* | 0.16758751 | 0.42710197 | 0.99979869 | 2592.91756 | 2912.30933 |
| *PMA2* | 0.165329483 | 0.42734656 | 0.99979869 | 2802.65318 | 3142.95685 |
| *SMF2* | 0.142728571 | 0.42735007 | 0.99979869 | 12746.0725 | 14071.5488 |
| *MIC27* | 0.145453093 | 0.42738425 | 0.99979869 | 9110.32997 | 10076.7338 |
| *YMR182W-A* | 0.299383831 | 0.42881822 | 0.99979869 | 330.699418 | 406.964732 |
| *MSH2* | 0.145075632 | 0.42915186 | 0.99979869 | 8822.55319 | 9755.87748 |
| *TOS2* | 0.157686411 | 0.42918015 | 0.99979869 | 3741.09813 | 4173.18243 |
| *TAF12* | 0.139647012 | 0.42985874 | 0.99979869 | 18361.134 | 20227.2748 |
| *AEP3* | 0.161436746 | 0.43076795 | 0.99979869 | 3079.6993 | 3444.33627 |
| *NRG1* | 0.150674622 | 0.43229745 | 0.99979869 | 5120.47564 | 5684.17995 |
| *SAC7* | 0.143100308 | 0.43328011 | 0.99979869 | 9187.39562 | 10145.4155 |
| *SEC7* | 0.135476274 | 0.4334664 | 0.99979869 | 43608.425 | 47901.9017 |
| *AGC1* | 0.145079774 | 0.43353339 | 0.99979869 | 7497.80449 | 8291.00946 |
| *MRPL4* | 0.14223582 | 0.43368976 | 0.99979869 | 9943.41936 | 10973.6964 |
| *POX1* | 0.195947107 | 0.43398176 | 0.99979869 | 1158.91123 | 1327.50461 |
| *ATG39* | 0.214711207 | 0.43480411 | 0.99979869 | 832.113874 | 965.644277 |
| *PCL2* | 0.154933352 | 0.43480471 | 0.99979869 | 3819.13929 | 4252.11514 |
| *VPS28* | 0.155187695 | 0.43542533 | 0.99979869 | 3725.4899 | 4148.58003 |
| *SNT2* | 0.143264235 | 0.43553986 | 0.99979869 | 8266.50992 | 9129.54133 |
| *SYT1* | 0.141323867 | 0.43566389 | 0.99979869 | 10135.5957 | 11178.7164 |
| *YPL257W* | 0.193616009 | 0.43566588 | 0.99979869 | 1193.05424 | 1364.40821 |
| *YMR262W* | 0.147541541 | 0.43591768 | 0.99979869 | 5734.07427 | 6351.52011 |
| *ALD2* | 0.151743482 | 0.43615555 | 0.99979869 | 4384.9377 | 4871.27559 |
| *CAT8* | 0.159965185 | 0.43619479 | 0.99979869 | 2978.24579 | 3327.47487 |
| *AGE2* | 0.141367168 | 0.43651014 | 0.99979869 | 9716.12448 | 10716.3963 |
| *SPT6* | 0.134095752 | 0.43656734 | 0.99979869 | 49742.4602 | 54587.6045 |
| *MTC4* | 0.143679538 | 0.43674389 | 0.99979869 | 7609.01315 | 8405.82067 |
| *MVB12* | 0.164114856 | 0.43699648 | 0.99979869 | 2516.82743 | 2820.05033 |
| *GCD2* | 0.134685802 | 0.43701509 | 0.99979869 | 35119.4977 | 38556.0643 |
| *SNC1* | 0.143311614 | 0.437203 | 0.99979869 | 7744.60966 | 8553.43508 |
| *YOL160W* | 0.494221748 | 0.43755933 | 0.99979869 | 91.6983636 | 129.16261 |
| *CLB6* | 0.254671598 | 0.43786062 | 0.99979869 | 482.879681 | 576.106246 |
| *VPS34* | 0.142720144 | 0.43803792 | 0.99979869 | 7945.56565 | 8771.7814 |
| *TRX3* | 0.148549623 | 0.43817543 | 0.99979869 | 5052.18963 | 5600.12175 |
| *ITC1* | 0.137886726 | 0.4383628 | 0.99979869 | 14196.6626 | 15620.475 |
| *HEM15* | 0.136750991 | 0.43839822 | 0.99979869 | 17381.7175 | 19109.9157 |
| *UBC13* | 0.139236355 | 0.43846739 | 0.99979869 | 11552.0428 | 12722.5171 |
| *RBH1* | 0.16851132 | 0.43849578 | 0.99979869 | 2132.47471 | 2396.68399 |
| *YJR039W* | 0.148510012 | 0.43868116 | 0.99979869 | 5001.46287 | 5543.74124 |
| *KIP3* | 0.144991916 | 0.4393227 | 0.99979869 | 6274.5093 | 6937.87735 |
| *CIS3* | 0.13171909 | 0.44000136 | 0.99979869 | 135373.124 | 148314.555 |
| *MTC7* | 0.135718133 | 0.44006818 | 0.99979869 | 18975.7082 | 20847.4604 |

| | | | | | |
|---|---|---|---|---|---|
| *YDL206W* | 0.149165098 | 0.44013634 | 0.99979869 | 4640.52251 | 5146.00241 |
| *COX23* | 0.151863008 | 0.44046251 | 0.99979869 | 3969.36853 | 4409.98055 |
| *GDS1* | 0.135435702 | 0.4405912 | 0.99979869 | 19362.9874 | 21268.7765 |
| *SIC1* | 0.135239092 | 0.44087694 | 0.99979869 | 19769.777 | 21712.6448 |
| *POL31* | 0.140951152 | 0.44105331 | 0.99979869 | 8449.90665 | 9317.13464 |
| *PSP2* | 0.135941579 | 0.44151581 | 0.99979869 | 16301.8229 | 17912.5988 |
| *NEJ1* | 0.148197322 | 0.44186358 | 0.99979869 | 4718.56367 | 5229.03552 |
| *PRP3* | 0.142702736 | 0.44210522 | 0.99979869 | 6925.17748 | 7645.19641 |
| *LUC7* | 0.152091685 | 0.44218731 | 0.99979869 | 3787.92283 | 4209.06094 |
| *PAU4* | 0.667140878 | 0.44221921 | 0.99979869 | 43.8981528 | 69.7068056 |
| *UMP1* | 0.134799519 | 0.44254041 | 0.99979869 | 19061.5534 | 20928.4433 |
| *BCK2* | 0.137816202 | 0.44276251 | 0.99979869 | 11286.7028 | 12418.0624 |
| *GRR1* | 0.13695337 | 0.44280088 | 0.99979869 | 12721.6847 | 13988.5157 |
| *YAP3* | 0.146333276 | 0.44290567 | 0.99979869 | 5169.25137 | 5721.08356 |
| *SEC20* | 0.142824481 | 0.4433768 | 0.99979869 | 6585.69843 | 7271.03488 |
| *TMA108* | 0.134265507 | 0.44408308 | 0.99979869 | 18964.002 | 20813.6321 |
| *AMN1* | 0.131396651 | 0.44434705 | 0.99979869 | 47590.4752 | 52128.3894 |
| *PRC1* | 0.130249053 | 0.44467368 | 0.99979869 | 106520.331 | 116584.632 |
| *CCP1* | 0.131478462 | 0.44535046 | 0.99979869 | 38511.3617 | 42185.9437 |
| *YBL101W-C* | 0.340019083 | 0.44595812 | 0.99979869 | 214.613191 | 271.651522 |
| *tS(AGA)B* | 0.984068551 | 0.44628694 | 0.99979869 | 16.5837466 | 32.8032026 |
| *YJR151W-A* | 0.984068551 | 0.44628694 | 0.99979869 | 16.5837466 | 32.8032026 |
| *MYO5* | 0.131797521 | 0.44650867 | 0.99979869 | 29156.1776 | 31945.1938 |
| *COP1* | 0.130060065 | 0.44654074 | 0.99979869 | 68326.0115 | 74771.8251 |
| *ATP5* | 0.134184221 | 0.44714217 | 0.99979869 | 15472.6356 | 16980.7829 |
| *LSO2* | 0.135274279 | 0.44722188 | 0.99979869 | 12848.5016 | 14111.5277 |
| *TOF2* | 0.142922744 | 0.44738931 | 0.99979869 | 5807.23785 | 6412.00101 |
| *RAD52* | 0.136748169 | 0.44775276 | 0.99979869 | 10231.1961 | 11248.4232 |
| *TIM11* | 0.135823738 | 0.4483255 | 0.99979869 | 11216.4658 | 12323.7532 |
| *LAA1* | 0.133236677 | 0.44837646 | 0.99979869 | 17032.4833 | 18680.3988 |
| *HUA1* | 0.136350536 | 0.44891611 | 0.99979869 | 10189.249 | 11199.2184 |
| *UBC4* | 0.129414682 | 0.44894878 | 0.99979869 | 59285.9186 | 64849.8814 |
| *YNL011C* | 0.149395173 | 0.44901224 | 0.99979869 | 3782.06974 | 4194.70953 |
| *BSC4* | 0.529212652 | 0.44902193 | 0.99979869 | 73.163588 | 105.585308 |
| *HYM1* | 0.146512198 | 0.44950406 | 0.99979869 | 4370.30499 | 4837.44729 |
| *HPA2* | 0.2712024 | 0.44950739 | 0.99979869 | 374.59757 | 452.069136 |
| *YPF1* | 0.131452531 | 0.44964177 | 0.99979869 | 23186.0288 | 25397.8796 |
| *PRP8* | 0.131265629 | 0.44965205 | 0.99979869 | 24390.7892 | 26714.1081 |
| *YDL114W* | 0.667634225 | 0.45021011 | 0.99979869 | 41.9471238 | 66.6315053 |
| *MED8* | 0.143512226 | 0.45052539 | 0.99979869 | 5129.25527 | 5665.72815 |
| *PYK2* | 0.134969909 | 0.45057928 | 0.99979869 | 11262.315 | 12366.8074 |
| *LSM12* | 0.133589621 | 0.45098136 | 0.99979869 | 13515.7535 | 14827.0476 |
| *KEL2* | 0.144378085 | 0.45126624 | 0.99979869 | 4765.38836 | 5266.96422 |
| *NPR2* | 0.147415332 | 0.45139544 | 0.99979869 | 3991.80536 | 4421.25665 |
| *STB2* | 0.149266972 | 0.45141902 | 0.99979869 | 3629.88948 | 4025.56802 |

| | | | | | |
|---|---|---|---|---|---|
| *SKN1* | 0.132309522 | 0.45154725 | 0.99979869 | 16345.7211 | 17915.6741 |
| *AYR1* | 0.132204819 | 0.45199958 | 0.99979869 | 16170.1285 | 17721.9302 |
| *INA1* | 0.128011385 | 0.45247724 | 0.99979869 | 67458.7791 | 73718.0222 |
| *SKN7* | 0.133334681 | 0.45251096 | 0.99979869 | 12904.1059 | 14153.5568 |
| *UBA1* | 0.127714839 | 0.45262927 | 0.99979869 | 83149.93 | 90846.4194 |
| *SOK2* | 0.132281859 | 0.45281435 | 0.99979869 | 15105.8421 | 16556.3914 |
| *CLA4* | 0.135485998 | 0.45292199 | 0.99979869 | 9490.78063 | 10425.2678 |
| *IDP1* | 0.130554553 | 0.45301201 | 0.99979869 | 21467.1722 | 23500.4194 |
| *TOS4* | 0.141744643 | 0.45332807 | 0.99979869 | 5378.98699 | 5934.30437 |
| *GET2* | 0.131485196 | 0.45375088 | 0.99979869 | 16483.2686 | 18056.1128 |
| *DMA1* | 0.132792042 | 0.454912 | 0.99979869 | 12301.2379 | 13487.2418 |
| *VMS1* | 0.135001874 | 0.45628213 | 0.99979869 | 8740.60997 | 9598.01207 |
| *NEL1* | 0.175406802 | 0.45657499 | 0.99979869 | 1424.25118 | 1608.38203 |
| *GAL3* | 0.178068886 | 0.45676504 | 0.99979869 | 1336.45487 | 1512.02262 |
| *YGL041C-B* | 0.42998452 | 0.45699805 | 0.99979869 | 114.135197 | 153.765012 |
| *SSU72* | 0.147830684 | 0.45719957 | 0.99979869 | 3484.53782 | 3860.52691 |
| *DDI2* | 0.306266583 | 0.45744169 | 0.99979869 | 259.486859 | 320.856326 |
| *YKL018C-A* | 0.144121163 | 0.45753867 | 0.99979869 | 4196.66341 | 4637.55277 |
| *YPS1* | 0.126985314 | 0.45785252 | 0.99979869 | 40950.1479 | 44717.9409 |
| *MUP1* | 0.127825171 | 0.45786556 | 0.99979869 | 29123.9856 | 31822.1818 |
| *CAT5* | 0.139518197 | 0.45801947 | 0.99979869 | 5559.45717 | 6123.94789 |
| *NRT1* | 0.135093359 | 0.45840287 | 0.99979869 | 7991.41483 | 8775.8818 |
| *MGR2* | 0.132078079 | 0.45847978 | 0.99979869 | 11356.9399 | 12445.7401 |
| *RQC2* | 0.127269685 | 0.45851646 | 0.99979869 | 32862.1572 | 35892.8543 |
| *AIM44* | 0.128792663 | 0.45856064 | 0.99979869 | 20551.1641 | 22470.1938 |
| *YER189W* | 0.273876558 | 0.45862009 | 0.99979869 | 343.381106 | 415.165533 |
| *NUF2* | 0.142534086 | 0.45887821 | 0.99979869 | 4466.88092 | 4930.73139 |
| *SUN4* | 0.126256512 | 0.45996241 | 0.99979869 | 41102.3282 | 44861.4549 |
| *MDS3* | 0.127560567 | 0.46003181 | 0.99979869 | 24978.0489 | 27287.1391 |
| *PAC10* | 0.131442648 | 0.46034294 | 0.99979869 | 11318.8948 | 12398.5855 |
| *MDM31* | 0.13451211 | 0.46035863 | 0.99979869 | 7891.91235 | 8663.12079 |
| *DAD3* | 0.144730895 | 0.46090432 | 0.99979869 | 3785.9718 | 4185.48363 |
| *HOS2* | 0.138179911 | 0.46101114 | 0.99979869 | 5653.10656 | 6221.3324 |
| *YPS5* | 0.332294491 | 0.46127698 | 0.99979869 | 206.809075 | 260.375421 |
| *YFL019C* | 0.375079957 | 0.46129153 | 0.99979869 | 154.131292 | 199.894516 |
| *YOL131W* | 0.419453833 | 0.46136147 | 0.99979869 | 118.037255 | 157.865413 |
| *YBL010C* | 0.150228733 | 0.4621398 | 0.99979869 | 2870.93919 | 3186.01105 |
| *DLS1* | 0.135060421 | 0.46216707 | 0.99979869 | 7042.23922 | 7733.35502 |
| *ECM16* | 0.125932822 | 0.46306252 | 0.99979869 | 30309.2357 | 33073.829 |
| *CRN1* | 0.125028451 | 0.46308252 | 0.99979869 | 44775.1403 | 48828.5922 |
| *HIR2* | 0.13778313 | 0.463099 | 0.99979869 | 5500.9263 | 6052.19088 |
| *YHC1* | 0.139118881 | 0.46334519 | 0.99979869 | 4982.9281 | 5487.36074 |
| *SMA2* | 0.175692377 | 0.46338655 | 0.99979869 | 1315.96907 | 1486.39512 |
| *MRX7* | 0.168866812 | 0.46418581 | 0.99979869 | 1537.41086 | 1728.31874 |
| *OCA5* | 0.135289105 | 0.46441108 | 0.99979869 | 6433.51817 | 7066.01486 |

| | | | | | |
|---|---|---|---|---|---|
| *SPF1* | 0.123749968 | 0.46479988 | 0.99979869 | 71166.7098 | 77540.6204 |
| *PDC6* | 0.293194119 | 0.46506891 | 0.99979869 | 276.070605 | 338.283027 |
| *RPS11A* | 0.122876134 | 0.46520563 | 0.99979869 | 186586.66 | 203174.836 |
| *CST6* | 0.127345067 | 0.46526557 | 0.99979869 | 16915.4215 | 18476.4039 |
| *MIT1* | 0.125485291 | 0.46537343 | 0.99979869 | 27007.1191 | 29461.3764 |
| *YFL066C* | 0.196111661 | 0.46539941 | 0.99979869 | 865.281367 | 991.271779 |
| *HEK2* | 0.124653993 | 0.46539951 | 0.99979869 | 36800.3092 | 40121.3921 |
| *TCB3* | 0.123433408 | 0.46568558 | 0.99979869 | 72407.5642 | 78875.3007 |
| *SIN3* | 0.124614149 | 0.4659436 | 0.99979869 | 34616.1322 | 37739.0595 |
| *VPS60* | 0.131617436 | 0.4660883 | 0.99979869 | 8624.52375 | 9448.34745 |
| *PEP4* | 0.123004448 | 0.46659134 | 0.99979869 | 81239.8726 | 88470.2375 |
| *ASF2* | 0.133563004 | 0.46673445 | 0.99979869 | 6949.56534 | 7623.66931 |
| *RPB2* | 0.123138705 | 0.46775693 | 0.99979869 | 53998.63 | 58809.9917 |
| *GRX1* | 0.124931207 | 0.46799258 | 0.99979869 | 24369.3279 | 26573.6694 |
| *TGL3* | 0.128611372 | 0.46827794 | 0.99979869 | 11408.6421 | 12472.3927 |
| *YDL159W-B* | 0.360251413 | 0.46829992 | 0.99979869 | 162.910923 | 209.120417 |
| *SNO4* | 1.393457444 | 0.46849582 | 0.99979869 | 5.85308704 | 15.3765012 |
| *SEN1* | 0.124566293 | 0.46875782 | 0.99979869 | 25165.3477 | 27434.7535 |
| *FIS1* | 0.132425128 | 0.46934193 | 0.99979869 | 7082.23531 | 7763.08292 |
| *MET32* | 0.164315408 | 0.46948894 | 0.99979869 | 1630.08474 | 1826.72835 |
| *INP52* | 0.132860676 | 0.46966529 | 0.99979869 | 6738.85421 | 7388.92139 |
| *NBA1* | 0.125120381 | 0.47017573 | 0.99979869 | 19100.574 | 20831.0588 |
| *HSX1* | 1.071531239 | 0.47027945 | 0.99979869 | 11.7061741 | 24.602402 |
| *VMA5* | 0.122355464 | 0.47062869 | 0.99979869 | 48615.7409 | 52918.7415 |
| *MRP8* | 0.124053957 | 0.47161204 | 0.99979869 | 22126.62 | 24113.4292 |
| *RAD54* | 0.131749814 | 0.47179179 | 0.99979869 | 6957.36946 | 7622.64421 |
| *YKL068W-A* | 0.136760579 | 0.47194592 | 0.99979869 | 4724.41675 | 5194.18211 |
| *SLI15* | 0.144315598 | 0.47217749 | 0.99979869 | 3129.45054 | 3458.68768 |
| *YPL264C* | 0.174623325 | 0.47379368 | 0.99979869 | 1211.58902 | 1367.48351 |
| *PFK26* | 0.131753464 | 0.47429795 | 0.99979869 | 6430.59162 | 7045.51286 |
| *GLO1* | 0.122175356 | 0.47439791 | 0.99979869 | 29815.6254 | 32450.5682 |
| *AQY1* | 0.166197139 | 0.47444174 | 0.99979869 | 1467.17382 | 1646.31073 |
| *COS7* | 0.140779969 | 0.47485883 | 0.99979869 | 3531.36251 | 3893.33011 |
| *YHR182W* | 0.138709073 | 0.47501279 | 0.99979869 | 3929.37243 | 4325.92235 |
| *VPS8* | 0.127783753 | 0.47545326 | 0.99979869 | 9200.07731 | 10052.1314 |
| *VPS52* | 0.133829227 | 0.47600016 | 0.99979869 | 5217.05158 | 5724.15886 |
| *PUN1* | 0.121520975 | 0.4760523 | 0.99979869 | 31042.8226 | 33770.8971 |
| *RFA3* | 0.129035106 | 0.47634702 | 0.99979869 | 7748.51172 | 8473.47728 |
| *PCP1* | 0.130390289 | 0.47639437 | 0.99979869 | 6793.48302 | 7436.07599 |
| *VPS15* | 0.130340353 | 0.47661844 | 0.99979869 | 6775.92376 | 7416.59909 |
| *RMA1* | 0.139394109 | 0.47662531 | 0.99979869 | 3665.98351 | 4037.86922 |
| *MRX9* | 0.12948169 | 0.4770561 | 0.99979869 | 7235.39109 | 7914.79773 |
| *YBR138C* | 0.14655318 | 0.4773555 | 0.99979869 | 2613.40336 | 2892.83243 |
| *RVS167* | 0.119947329 | 0.47758088 | 0.99979869 | 52448.5374 | 56995.5645 |
| *USO1* | 0.121676222 | 0.47795018 | 0.99979869 | 23884.4972 | 25986.2871 |

| | | | | | |
|---|---|---|---|---|---|
| *CLU1* | 0.119379954 | 0.47823969 | 0.99979869 | 67775.8213 | 73622.6879 |
| *YOR161C-C* | 0.330822299 | 0.47848872 | 0.99979869 | 188.2743 | 236.798119 |
| *RPT3* | 0.120094992 | 0.47862544 | 0.99979869 | 39920.0046 | 43385.3108 |
| *COQ6* | 0.124194231 | 0.47909001 | 0.99979869 | 12751.9256 | 13898.3069 |
| *YGL149W* | 0.176882843 | 0.47923341 | 0.99979869 | 1094.52728 | 1237.2958 |
| *LAP2* | 0.122014398 | 0.4800953 | 0.99979869 | 18188.468 | 19793.6575 |
| *YLR410W-B* | 0.127455843 | 0.4806435 | 0.99979869 | 7823.62634 | 8546.25938 |
| *DEF1* | 0.11841609 | 0.48113424 | 0.99979869 | 69399.0775 | 75335.6301 |
| *INO1* | 0.172997474 | 0.4813156 | 0.99979869 | 1164.76432 | 1313.1532 |
| *FLO5* | 0.136798366 | 0.48164084 | 0.99979869 | 3815.23723 | 4194.70953 |
| *YIR014W* | 0.147823872 | 0.48271884 | 0.99979869 | 2299.28769 | 2547.3737 |
| *ECM11* | 0.299946415 | 0.48278323 | 0.99979869 | 233.147967 | 287.028023 |
| *SPC42* | 0.139309959 | 0.48301564 | 0.99979869 | 3265.04705 | 3596.05109 |
| *CPR1* | 0.116725731 | 0.48345597 | 0.99979869 | 357273.408 | 387381.221 |
| *tR(UCU)K* | 0.644110033 | 0.4834831 | 0.99979869 | 38.0450657 | 59.4558047 |
| *PAN3* | 0.124709214 | 0.48380854 | 0.99979869 | 9456.63762 | 10310.4566 |
| *YFR032C-B* | 0.175074701 | 0.48502781 | 0.99979869 | 1075.01699 | 1213.7185 |
| *ERD2* | 0.120304678 | 0.48587578 | 0.99979869 | 17418.787 | 18933.5985 |
| *RPR2* | 0.134176663 | 0.48602203 | 0.99979869 | 4044.48314 | 4438.68335 |
| *SAS3* | 0.122976084 | 0.48642039 | 0.99979869 | 10587.2589 | 11529.3006 |
| *YBR090C* | 0.135912661 | 0.48651961 | 0.99979869 | 3633.79154 | 3992.76482 |
| *TUB1* | 0.11690346 | 0.48751863 | 0.99979869 | 47569.0139 | 51584.0612 |
| *tG(GCC)P1* | 0.569030743 | 0.48771049 | 0.99979869 | 49.7512398 | 73.8072059 |
| *YIP5* | 0.118585567 | 0.48839212 | 0.99979869 | 21802.7492 | 23670.586 |
| *YIM1* | 0.122950038 | 0.48857174 | 0.99979869 | 9651.74052 | 10510.3511 |
| *ROM1* | 0.122814414 | 0.48898097 | 0.99979869 | 9650.76501 | 10508.3009 |
| *LST8* | 0.121733321 | 0.48974155 | 0.99979869 | 10798.9456 | 11749.6971 |
| *YMR155W* | 0.130483613 | 0.48985923 | 0.99979869 | 4677.59206 | 5120.37491 |
| *DSE2* | 0.115966635 | 0.48994022 | 0.99979869 | 52860.2045 | 57284.6428 |
| *INO80* | 0.117252242 | 0.49036347 | 0.99979869 | 27084.1847 | 29377.3181 |
| *YDL180W* | 0.126001717 | 0.49067738 | 0.99979869 | 6464.73463 | 7054.73876 |
| *NPR1* | 0.125753674 | 0.49112554 | 0.99979869 | 6520.33896 | 7114.19457 |
| *TES1* | 0.12548407 | 0.49125397 | 0.99979869 | 6655.93548 | 7260.78388 |
| *COS2* | 0.277981032 | 0.49176177 | 0.99979869 | 266.31546 | 322.906526 |
| *DSN1* | 0.123815085 | 0.4927373 | 0.99979869 | 7453.90634 | 8121.86795 |
| *YKR011C* | 0.123135802 | 0.49339137 | 0.99979869 | 7821.67531 | 8518.58168 |
| *VID22* | 0.121041221 | 0.49357639 | 0.99979869 | 9974.63582 | 10847.6091 |
| *CHL4* | 0.131832346 | 0.49365277 | 0.99979869 | 3953.76029 | 4332.07295 |
| *YHL018W* | 0.174429758 | 0.49380993 | 0.99979869 | 1002.82891 | 1131.71049 |
| *YMR181C* | 0.122418453 | 0.49388048 | 0.99979869 | 8314.31014 | 9050.60862 |
| *SCS3* | 0.116097952 | 0.4942888 | 0.99979869 | 26211.0993 | 28407.5735 |
| *SCC2* | 0.124573551 | 0.49585793 | 0.99979869 | 6277.43585 | 6843.56815 |
| *YGL176C* | 0.130965869 | 0.49622657 | 0.99979869 | 3943.02963 | 4317.72154 |
| *YBR012W-B* | 0.152295668 | 0.49661912 | 0.99979869 | 1660.32569 | 1845.18015 |
| *YHR213W-B* | 0.257943225 | 0.49705868 | 0.99979869 | 311.189127 | 372.11133 |

| | | | | | |
|---|---|---|---|---|---|
| *SUP51* | 0.705403548 | 0.49796272 | 0.99979869 | 28.2899207 | 46.1295037 |
| *YJL132W* | 0.134802225 | 0.49877939 | 0.99979869 | 3067.99312 | 3368.47887 |
| *YBL113C* | 0.126018665 | 0.49942964 | 0.99979869 | 5068.77337 | 5531.44004 |
| *MRC1* | 0.122815023 | 0.49968763 | 0.99979869 | 6550.57991 | 7132.64637 |
| *ETR1* | 0.116853731 | 0.50016329 | 0.99979869 | 13754.7545 | 14915.2062 |
| *tD(GUC)I2* | 0.919528454 | 0.50024601 | 0.99979869 | 14.6327176 | 27.6777022 |
| *ESP1* | 0.122042812 | 0.50048859 | 0.99979869 | 6871.52418 | 7478.1051 |
| *LDH1* | 0.12265126 | 0.5006117 | 0.99979869 | 6463.75912 | 7037.31206 |
| *SPC110* | 0.119139976 | 0.50078642 | 0.99979869 | 9391.27815 | 10199.7458 |
| *BUB2* | 0.14044062 | 0.50145839 | 0.99979869 | 2312.94489 | 2549.4239 |
| *VAC17* | 0.124309459 | 0.50150719 | 0.99979869 | 5471.66086 | 5964.03228 |
| *CIN1* | 0.12844353 | 0.50159288 | 0.99979869 | 4097.16093 | 4478.66226 |
| *CRM1* | 0.113110049 | 0.50180207 | 0.99979869 | 33361.6206 | 36082.4978 |
| *DOG2* | 0.119217495 | 0.50182607 | 0.99979869 | 8911.32501 | 9678.99497 |
| *YLR283W* | 0.130438962 | 0.50185156 | 0.99979869 | 3627.93845 | 3971.23772 |
| *PBP4* | 0.118481931 | 0.50191038 | 0.99979869 | 9760.99815 | 10596.4595 |
| *YCL048W-A* | 0.330264624 | 0.50207204 | 0.99979869 | 163.886437 | 206.045116 |
| *YDR098C-B* | 0.165505984 | 0.5024307 | 0.99979869 | 1115.98859 | 1251.6472 |
| *BST1* | 0.121888488 | 0.50254099 | 0.99979869 | 6535.94719 | 7112.14437 |
| *FES1* | 0.112697927 | 0.50265537 | 0.99979869 | 35655.0552 | 38551.9639 |
| *YLR042C* | 0.143127302 | 0.5030286 | 0.99979869 | 2051.50701 | 2265.47118 |
| *PEX8* | 0.120314037 | 0.50314655 | 0.99979869 | 7488.04935 | 8139.29465 |
| *CBP1* | 0.124367871 | 0.50364147 | 0.99979869 | 5151.69211 | 5615.49825 |
| *MDM34* | 0.117840474 | 0.50442911 | 0.99979869 | 9539.55636 | 10351.4606 |
| *CSM3* | 0.135660105 | 0.50501793 | 0.99979869 | 2661.20357 | 2923.58543 |
| *MAP2* | 0.111136694 | 0.50560153 | 0.99979869 | 51310.112 | 55418.9606 |
| *SPG5* | 0.121799736 | 0.50568951 | 0.99979869 | 5997.46318 | 6525.78712 |
| *KKQ8* | 0.118698972 | 0.50629101 | 0.99979869 | 7983.61072 | 8668.24629 |
| *LDS2* | 0.137759694 | 0.50642081 | 0.99979869 | 2388.05951 | 2627.33151 |
| *TAH1* | 0.121424868 | 0.50655555 | 0.99979869 | 6043.31237 | 6573.96682 |
| *COS5* | 0.130768962 | 0.50662355 | 0.99979869 | 3258.21845 | 3567.34828 |
| *PTH1* | 0.129403856 | 0.50695283 | 0.99979869 | 3479.66024 | 3806.1966 |
| *NUP116* | 0.111866295 | 0.50786129 | 0.99979869 | 26113.5478 | 28218.9551 |
| *YCK2* | 0.111361211 | 0.50799914 | 0.99979869 | 31118.9127 | 33616.107 |
| *UBP6* | 0.111819087 | 0.50849333 | 0.99979869 | 24835.6238 | 26837.1201 |
| *NCL1* | 0.111626076 | 0.5088617 | 0.99979869 | 25510.6798 | 27562.891 |
| *CPR5* | 0.110405651 | 0.50919915 | 0.99979869 | 41109.1568 | 44378.6327 |
| *NTG1* | 0.121223386 | 0.50972007 | 0.99979869 | 5622.86561 | 6115.74709 |
| *YER084W* | 0.254620887 | 0.50987987 | 0.99979869 | 295.580895 | 352.634428 |
| *LCL1* | 0.124501776 | 0.50999529 | 0.99979869 | 4382.01116 | 4776.96638 |
| *SUT1* | 0.128380257 | 0.51037908 | 0.99979869 | 3446.49275 | 3767.2428 |
| *STB1* | 0.128657499 | 0.51060469 | 0.99979869 | 3381.13328 | 3696.51089 |
| *RAD4* | 0.120074451 | 0.51064597 | 0.99979869 | 6043.31237 | 6567.81622 |
| *IDP2* | 0.147771853 | 0.51102193 | 0.99979869 | 1599.84379 | 1772.39804 |
| *BIO4* | 0.148716649 | 0.51170154 | 0.99979869 | 1544.23946 | 1711.91714 |

| | | | | | |
|---|---|---|---|---|---|
| *MRL1* | 0.118440837 | 0.51182989 | 0.99979869 | 6788.60545 | 7369.44449 |
| *ERP4* | 0.117815072 | 0.51229507 | 0.99979869 | 7121.25589 | 7727.20442 |
| *PTP1* | 0.118596921 | 0.51287406 | 0.99979869 | 6471.56323 | 7026.03596 |
| *PFS2* | 0.118671604 | 0.51314627 | 0.99979869 | 6373.03627 | 6919.42555 |
| *ERS1* | 0.126800257 | 0.51342235 | 0.99979869 | 3546.97074 | 3872.82811 |
| *ISW1* | 0.111499755 | 0.51393328 | 0.99979869 | 17050.0425 | 18420.0234 |
| *CKI1* | 0.115996981 | 0.51407707 | 0.99979869 | 8165.05642 | 8848.66391 |
| *MRK1* | 0.132817864 | 0.51412395 | 0.99979869 | 2607.55027 | 2859.00413 |
| *LSM8* | 0.142175847 | 0.51440177 | 0.99979869 | 1827.13867 | 2016.37186 |
| *TIF4632* | 0.109870514 | 0.51549102 | 0.99979869 | 23326.5029 | 25172.3576 |
| *DIT1* | 0.169097116 | 0.51656547 | 0.99979869 | 906.252976 | 1018.94948 |
| *PMP1* | 0.107398965 | 0.51670727 | 0.99979869 | 69402.004 | 74765.6745 |
| *PAN2* | 0.112730994 | 0.51721218 | 0.99979869 | 11112.0857 | 12015.1981 |
| *ATG31* | 0.133615472 | 0.51746764 | 0.99979869 | 2395.86363 | 2628.35661 |
| *CEF1* | 0.116390434 | 0.5187283 | 0.99979869 | 6671.54371 | 7232.08108 |
| *RPS27A* | 0.106699901 | 0.51914827 | 0.99979869 | 67715.3394 | 72913.3186 |
| *RDH54* | 0.110797917 | 0.51916621 | 0.99979869 | 14035.7027 | 15156.1047 |
| *ADD37* | 0.118826889 | 0.51921761 | 0.99979869 | 5298.01928 | 5752.86166 |
| *SCP160* | 0.106610021 | 0.51926566 | 0.99979869 | 71299.3797 | 76767.6949 |
| *SLG1* | 0.108659416 | 0.51963161 | 0.99979869 | 23012.3872 | 24812.5475 |
| *GPD1* | 0.106530152 | 0.51985702 | 0.99979869 | 65306.7941 | 70311.6146 |
| *GAL11* | 0.109294017 | 0.52019197 | 0.99979869 | 18317.2359 | 19758.8041 |
| *SGS1* | 0.113611663 | 0.52025089 | 0.99979869 | 8602.08691 | 9306.88364 |
| *SMC4* | 0.110557106 | 0.52128311 | 0.99979869 | 12985.0736 | 14019.2687 |
| *BEM2* | 0.106663246 | 0.52190286 | 0.99979869 | 39413.7126 | 42438.1183 |
| *YGR016W* | 0.155686209 | 0.52197369 | 0.99979869 | 1153.05815 | 1284.4504 |
| *SMD2* | 0.125401154 | 0.52202473 | 0.99979869 | 3256.26742 | 3551.97178 |
| *RNR2* | 0.105864275 | 0.52302523 | 0.99979869 | 52655.3465 | 56664.4572 |
| *AXL1* | 0.10943602 | 0.52304389 | 0.99979869 | 14546.8723 | 15693.2572 |
| *UBX7* | 0.108881083 | 0.52417333 | 0.99979869 | 15212.1732 | 16404.6766 |
| *GAS5* | 0.105547802 | 0.52527423 | 0.99979869 | 42111.9857 | 45308.3985 |
| *FUN30* | 0.106983082 | 0.52534549 | 0.99979869 | 22742.1697 | 24492.7163 |
| *YIL165C* | 0.143039266 | 0.5254866 | 0.99979869 | 1554.97012 | 1717.04264 |
| *RFX1* | 0.11753706 | 0.52568754 | 0.99979869 | 4967.31987 | 5388.95113 |
| *DIG1* | 0.106167907 | 0.52588632 | 0.99979869 | 28703.5388 | 30895.4914 |
| *ARC15* | 0.106481735 | 0.52616017 | 0.99979869 | 24782.946 | 26681.3049 |
| *ARB1* | 0.104288667 | 0.52668213 | 0.99979869 | 82572.4254 | 88762.391 |
| *tN(GUU)G* | 0.946000649 | 0.52722438 | 0.99979869 | 11.7061741 | 22.5522018 |
| *RPN8* | 0.105175755 | 0.52733658 | 0.99979869 | 37043.2123 | 39844.6151 |
| *PUB1* | 0.104323799 | 0.5275948 | 0.99979869 | 61522.7734 | 66136.382 |
| *YCR024C-B* | 0.104029999 | 0.52761363 | 0.99979869 | 81395.9549 | 87482.041 |
| *PSY3* | 0.157105571 | 0.52777646 | 0.99979869 | 1053.55567 | 1174.76469 |
| *YPL277C* | 0.149532331 | 0.52789692 | 0.99979869 | 1264.2668 | 1402.33691 |
| *SPG4* | 0.244133224 | 0.52798488 | 0.99979869 | 291.678837 | 345.458728 |
| *YGR266W* | 0.110290389 | 0.52836767 | 0.99979869 | 9564.91973 | 10324.808 |

| | | | | | |
|---|---|---|---|---|---|
| *NFT1* | 0.123757729 | 0.52867085 | 0.99979869 | 3148.96083 | 3431.00997 |
| *MCM4* | 0.107020369 | 0.5287428 | 0.99979869 | 16999.3158 | 18308.2875 |
| *YHC3* | 0.121836091 | 0.528794 | 0.99979869 | 3491.36642 | 3799.0209 |
| *SKO1* | 0.10996197 | 0.52906876 | 0.99979869 | 9720.02654 | 10489.8491 |
| *DIE2* | 0.111585117 | 0.52910653 | 0.99979869 | 7865.57346 | 8498.07968 |
| *PRK1* | 0.113356133 | 0.52927009 | 0.99979869 | 6466.68566 | 6995.28296 |
| *RAV2* | 0.112712021 | 0.52940665 | 0.99979869 | 6872.4997 | 7430.95049 |
| *CDC1* | 0.107486015 | 0.52959792 | 0.99979869 | 14457.125 | 15575.3706 |
| *PRM10* | 0.124837946 | 0.52966416 | 0.99979869 | 2928.49455 | 3193.18675 |
| *ORM2* | 0.104252352 | 0.53087313 | 0.99979869 | 34996.5829 | 37619.1228 |
| *CAD1* | 0.109724577 | 0.5316111 | 0.99979869 | 9017.65609 | 9730.24998 |
| *SFH1* | 0.106355101 | 0.53164338 | 0.99979869 | 16210.1245 | 17450.2787 |
| *NIP1* | 0.104084069 | 0.53188008 | 0.99979869 | 32956.7821 | 35422.3333 |
| *AUR1* | 0.104796188 | 0.53214501 | 0.99979869 | 24088.3797 | 25903.254 |
| *TMA64* | 0.109811905 | 0.53283385 | 0.99979869 | 8488.92723 | 9160.29433 |
| *YEL043W* | 0.105403792 | 0.53319087 | 0.99979869 | 18313.3338 | 19701.3985 |
| *VHS3* | 0.107201488 | 0.53348384 | 0.99979869 | 12196.8579 | 13137.6826 |
| *CBP4* | 0.11236134 | 0.53389801 | 0.99979869 | 6184.76197 | 6685.70273 |
| *COX15* | 0.105236546 | 0.53410673 | 0.99979869 | 17827.5276 | 19176.5472 |
| *YKR005C* | 0.151431013 | 0.53414596 | 0.99979869 | 1130.62131 | 1255.7476 |
| *SET6* | 0.123750965 | 0.53426004 | 0.99979869 | 2858.2575 | 3114.25405 |
| *AAD6* | 0.12188994 | 0.5349911 | 0.99979869 | 3103.11164 | 3376.67967 |
| *NYV1* | 0.105915935 | 0.5350118 | 0.99979869 | 14316.6509 | 15407.2542 |
| *MHO1* | 0.113153653 | 0.5352672 | 0.99979869 | 5532.14276 | 5983.50918 |
| *SMC6* | 0.112512094 | 0.53527258 | 0.99979869 | 5856.98909 | 6332.04321 |
| *MCK1* | 0.103661646 | 0.53527891 | 0.99979869 | 26031.6046 | 27970.8808 |
| *MSS11* | 0.11235789 | 0.53563102 | 0.99979869 | 5879.42593 | 6355.62051 |
| *RAD16* | 0.107169361 | 0.53575139 | 0.99979869 | 10920.8849 | 11763.0234 |
| *CDC14* | 0.105434125 | 0.53645265 | 0.99979869 | 14481.5128 | 15579.471 |
| *LSC1* | 0.103271321 | 0.53656766 | 0.99979869 | 26179.8828 | 28122.5956 |
| *ERG13* | 0.101782389 | 0.53662853 | 0.99979869 | 57911.4187 | 62144.6423 |
| *PRM15* | 0.107240492 | 0.536797 | 0.99979869 | 10276.0698 | 11069.0307 |
| *SMY2* | 0.103868418 | 0.53709848 | 0.99979869 | 20398.0083 | 21920.7401 |
| *NPL4* | 0.106567627 | 0.5372641 | 0.99979869 | 11190.1269 | 12048.0013 |
| *CSR2* | 0.119804412 | 0.53742176 | 0.99979869 | 3325.52895 | 3613.47779 |
| *PEX2* | 0.107357271 | 0.53751053 | 0.99979869 | 9776.60638 | 10531.8782 |
| *EOS1* | 0.107012645 | 0.53766207 | 0.99979869 | 10225.3431 | 11012.6502 |
| *SEG1* | 0.103689528 | 0.53789423 | 0.99979869 | 20079.9906 | 21576.3065 |
| *YIL166C* | 0.131830292 | 0.53821654 | 0.99979869 | 1919.81255 | 2103.50537 |
| *MOH1* | 0.143116893 | 0.53835119 | 0.99979869 | 1342.30796 | 1482.29472 |
| *TAM41* | 0.111140136 | 0.53867373 | 0.99979869 | 6027.70413 | 6510.41062 |
| *PBI2* | 0.106075135 | 0.53876852 | 0.99979869 | 11271.0946 | 12131.0344 |
| *TMS1* | 0.105743545 | 0.53941 | 0.99979869 | 11544.2387 | 12422.1628 |
| *STE23* | 0.103240152 | 0.53968288 | 0.99979869 | 19665.3969 | 21124.2374 |
| *RPS7B* | 0.100168285 | 0.54037695 | 0.99979869 | 104611.249 | 112132.623 |

| | | | | | |
|---|---|---|---|---|---|
| *PHM8* | 0.105699595 | 0.54046302 | 0.99979869 | 11030.1425 | 11868.6087 |
| *YJL193W* | 0.117406885 | 0.54066994 | 0.99979869 | 3592.81993 | 3897.43051 |
| *VPS55* | 0.106652288 | 0.54080233 | 0.99979869 | 9381.52301 | 10101.3362 |
| *YCR006C* | 0.133814067 | 0.54117792 | 0.99979869 | 1724.70965 | 1892.33475 |
| *PTR3* | 0.112148685 | 0.54158593 | 0.99979869 | 5090.23469 | 5501.71214 |
| *VPS53* | 0.105782912 | 0.54167222 | 0.99979869 | 10272.1677 | 11053.6542 |
| *SEC3* | 0.102579192 | 0.54181109 | 0.99979869 | 19909.2756 | 21376.412 |
| *SMY1* | 0.105984214 | 0.54201567 | 0.99979869 | 9807.82284 | 10555.4555 |
| *DUG1* | 0.100088727 | 0.5420962 | 0.99979869 | 65090.2299 | 69766.2614 |
| *YHR213W-A* | 1.071530995 | 0.54227673 | 0.99979869 | 7.80411605 | 16.4016013 |
| *RPS11B* | 0.099288312 | 0.54241342 | 0.99979869 | 190864.291 | 204462.362 |
| *NAT1* | 0.100660564 | 0.5424639 | 0.99979869 | 39886.8371 | 42769.2256 |
| *UBP8* | 0.112101267 | 0.54248414 | 0.99979869 | 4993.65876 | 5397.15193 |
| *YSW1* | 0.152601453 | 0.54255402 | 0.99979869 | 1012.58406 | 1125.55989 |
| *MGR1* | 0.10758505 | 0.54355234 | 0.99979869 | 7478.2942 | 8057.28664 |
| *YEL009C-A* | 0.338469377 | 0.54402491 | 0.99979869 | 119.988284 | 151.714812 |
| *PPQ1* | 0.103194035 | 0.54413919 | 0.99979869 | 14419.0799 | 15488.2371 |
| *AVT3* | 0.102798888 | 0.54443203 | 0.99979869 | 15449.2232 | 16590.2197 |
| *tV(UAC)D* | 0.451803041 | 0.54446493 | 0.99979869 | 61.4574139 | 84.0582067 |
| *PES4* | 0.288154261 | 0.54456337 | 0.99979869 | 174.617097 | 213.220817 |
| *TYE7* | 0.100946641 | 0.54510903 | 0.99979869 | 24861.9627 | 26663.8782 |
| *YPR063C* | 0.101561564 | 0.54528629 | 0.99979869 | 19988.2922 | 21446.1188 |
| *RPS10A* | 0.098342755 | 0.54561012 | 0.99979869 | 219760.981 | 235263.544 |
| *ART10* | 0.113774603 | 0.54588903 | 0.99979869 | 4070.82203 | 4404.85505 |
| *LSB1* | 0.099686721 | 0.54664315 | 0.99979869 | 35489.2177 | 38028.1377 |
| *ERP3* | 0.112403821 | 0.54694754 | 0.99979869 | 4379.08462 | 4733.91218 |
| *DPL1* | 0.100074055 | 0.54715904 | 0.99979869 | 27777.7756 | 29773.0068 |
| *YHR214W* | 0.142242656 | 0.54773901 | 0.99979869 | 1242.80548 | 1371.58391 |
| *DER1* | 0.112597503 | 0.54792936 | 0.99979869 | 4223.0023 | 4565.79576 |
| *NUP49* | 0.100960112 | 0.54864484 | 0.99979869 | 18034.3367 | 19341.5883 |
| *YEL008W* | 0.20159055 | 0.54936672 | 0.99979869 | 413.618151 | 475.646438 |
| *PSR1* | 0.101933659 | 0.54947701 | 0.99979869 | 13557.7006 | 14550.2706 |
| *MAG1* | 0.113783524 | 0.55026627 | 0.99979869 | 3708.90615 | 4013.26682 |
| *NUS1* | 0.102739693 | 0.55093844 | 0.99979869 | 10795.0435 | 11591.8317 |
| *SKG1* | 0.103260684 | 0.55147182 | 0.99979869 | 9694.66316 | 10413.9917 |
| *GEP5* | 0.11274208 | 0.55187835 | 0.99979869 | 3837.67407 | 4149.60513 |
| *tV(AAC)G3* | 0.551524288 | 0.55222016 | 0.99979869 | 37.0695512 | 54.3303043 |
| *tV(AAC)L* | 0.455674503 | 0.55227705 | 0.99979869 | 57.5553559 | 78.9327063 |
| *YGL235W* | 0.295202345 | 0.55230325 | 0.99979869 | 157.057835 | 192.718815 |
| *INP53* | 0.099017711 | 0.55311708 | 0.99979869 | 22009.5583 | 23573.2015 |
| *LRG1* | 0.106892885 | 0.55465614 | 0.99979869 | 5660.91068 | 6096.27019 |
| *YKR051W* | 0.109012175 | 0.55507501 | 0.99979869 | 4669.78794 | 5036.3167 |
| *YEL077C* | 0.106491361 | 0.55509439 | 0.99979869 | 5806.26234 | 6251.0603 |
| *tR(ACG)K* | 0.581726185 | 0.55514795 | 0.99979869 | 32.1919787 | 48.1797038 |
| *SPC34* | 0.112320702 | 0.55530872 | 0.99979869 | 3673.78763 | 3971.23772 |

| | | | | | |
|---|---|---|---|---|---|
| *SMX3* | 0.106602505 | 0.5555637 | 0.99979869 | 5668.71479 | 6103.44589 |
| *GPP2* | 0.097843356 | 0.55587956 | 0.99979869 | 25333.1362 | 27110.8219 |
| *SYH1* | 0.097532873 | 0.55593789 | 0.99979869 | 28534.7748 | 30530.5557 |
| *YLR296W* | 0.502165559 | 0.55659363 | 0.99979869 | 44.8736673 | 63.5562051 |
| *MMT2* | 0.102288623 | 0.55698235 | 0.99979869 | 8827.43077 | 9476.02516 |
| *DCI1* | 0.132970581 | 0.55750236 | 0.99979869 | 1457.41867 | 1598.13103 |
| *FAT1* | 0.098570937 | 0.55763405 | 0.99979869 | 17220.7576 | 18438.4752 |
| *SAE2* | 0.143333054 | 0.55772909 | 0.99979869 | 1089.6497 | 1203.4675 |
| *IMP2'* | 0.097354073 | 0.55839917 | 0.99979869 | 23386.9848 | 25019.6177 |
| *YPL060C-A* | 0.283034446 | 0.55850984 | 0.99979869 | 166.812981 | 202.969816 |
| *UTP14* | 0.096055291 | 0.55938745 | 0.99979869 | 36698.8557 | 39225.4546 |
| *BUD31* | 0.126751755 | 0.55943215 | 0.99979869 | 1750.07302 | 1910.78655 |
| *GLG2* | 0.115280601 | 0.55970011 | 0.99979869 | 2851.4289 | 3088.62655 |
| *KEG1* | 0.112746343 | 0.55999089 | 0.99979869 | 3265.04705 | 3530.44468 |
| *RIM20* | 0.100976577 | 0.56031915 | 0.99979869 | 9272.26538 | 9944.49589 |
| *PTC5* | 0.099100802 | 0.56054061 | 0.99979869 | 12639.7415 | 13538.4968 |
| *SEC26* | 0.095158452 | 0.56055752 | 0.99979869 | 53758.6534 | 57424.0564 |
| *tE(UUC)E2* | 0.605867979 | 0.56098441 | 0.99979869 | 28.2899207 | 43.0542034 |
| *MIC10* | 0.098489173 | 0.56131317 | 0.99979869 | 13705.0033 | 14673.2826 |
| *YKR045C* | 0.112777493 | 0.56161821 | 0.99979869 | 3161.64251 | 3418.70877 |
| *YKU70* | 0.106726324 | 0.56190099 | 0.99979869 | 4740.02499 | 5103.97331 |
| *YPR064W* | 0.230337178 | 0.5621209 | 0.99979869 | 268.266489 | 314.705725 |
| *GEP7* | 0.123783055 | 0.56240334 | 0.99979869 | 1877.86542 | 2046.09976 |
| *GPB1* | 0.100554432 | 0.56241326 | 0.99979869 | 9024.4847 | 9675.91967 |
| *YPK2* | 0.096402181 | 0.56260968 | 0.99979869 | 21653.4955 | 23149.8351 |
| *GPI11* | 0.104280785 | 0.56311676 | 0.99979869 | 5693.10266 | 6119.84749 |
| *CRD1* | 0.110957875 | 0.56325361 | 0.99979869 | 3415.27629 | 3688.31009 |
| *CIR2* | 0.097414046 | 0.56329434 | 0.99979869 | 15411.1782 | 16487.7097 |
| *IRC24* | 0.098253308 | 0.56383616 | 0.99979869 | 12418.2997 | 13293.4979 |
| *MRN1* | 0.097027345 | 0.56429839 | 0.99979869 | 15811.1391 | 16911.076 |
| *MCH1* | 0.101501453 | 0.56468936 | 0.99979869 | 7295.87299 | 7827.66422 |
| *YER188C-A* | 0.250172446 | 0.56519203 | 0.99979869 | 214.613191 | 255.24992 |
| *HRP1* | 0.094385088 | 0.56533631 | 0.99979869 | 37387.569 | 39915.347 |
| *PIC2* | 0.095194116 | 0.56541202 | 0.99979869 | 25304.8463 | 27030.8641 |
| *HUR1* | 0.126398247 | 0.56705484 | 0.99979869 | 1610.57445 | 1758.04664 |
| *YIR035C* | 0.095905651 | 0.56717018 | 0.99979869 | 17174.9084 | 18355.4421 |
| *PRE1* | 0.094188481 | 0.56722969 | 0.99979869 | 31552.0412 | 33680.6883 |
| *TSC10* | 0.096907699 | 0.56734175 | 0.99979869 | 13357.7201 | 14285.7947 |
| *YAR066W* | 0.196156677 | 0.56763353 | 0.99979869 | 389.230288 | 445.918536 |
| *DCK1* | 0.096246466 | 0.56849295 | 0.99979869 | 14395.6676 | 15388.8024 |
| *YKE4* | 0.097964629 | 0.56851668 | 0.99979869 | 10286.8005 | 11009.5749 |
| *FDO1* | 0.116249832 | 0.56856755 | 0.99979869 | 2354.89202 | 2552.4992 |
| *SOE1* | 0.299799431 | 0.56873994 | 0.99979869 | 136.572031 | 168.116413 |
| *RTT101* | 0.102242344 | 0.57040413 | 0.99979869 | 5621.8901 | 6034.76418 |
| *SUP54* | 0.919528496 | 0.57058971 | 0.99979869 | 9.75514506 | 18.4518015 |

| | | | | | |
|---|---|---|---|---|---|
| *ENA5* | 0.109657245 | 0.57067039 | 0.99979869 | 3205.54067 | 3458.68768 |
| *VPS3* | 0.099816891 | 0.57068804 | 0.99979869 | 7242.21969 | 7761.03272 |
| *RCR2* | 0.099716562 | 0.57112667 | 0.99979869 | 7218.80735 | 7735.40522 |
| *PTK1* | 0.096201548 | 0.57141086 | 0.99979869 | 12216.3682 | 13058.7499 |
| *SML1* | 0.092264467 | 0.57163549 | 0.99979869 | 48924.0035 | 52155.042 |
| *MRP49* | 0.104873511 | 0.57165143 | 0.99979869 | 4339.08852 | 4666.25557 |
| *APQ12* | 0.102172199 | 0.57171843 | 0.99979869 | 5453.12609 | 5853.32147 |
| *VTI1* | 0.097151086 | 0.57195167 | 0.99979869 | 10017.5585 | 10715.3712 |
| *SLA2* | 0.0921296 | 0.57224139 | 0.99979869 | 47889.9581 | 51047.9339 |
| *YGL082W* | 0.095835369 | 0.57228637 | 0.99979869 | 12516.8266 | 13376.531 |
| *WAR1* | 0.099042556 | 0.57258291 | 0.99979869 | 7448.05325 | 7977.32884 |
| *YFR045W* | 0.102243436 | 0.57279221 | 0.99979869 | 5258.9987 | 5645.22615 |
| *SUP56* | 0.563384597 | 0.57294303 | 0.99979869 | 31.2164642 | 46.1295037 |
| *ADA2* | 0.101090285 | 0.57298397 | 0.99979869 | 5843.33189 | 6267.4619 |
| *JID1* | 0.123388759 | 0.57342287 | 0.99979869 | 1652.52157 | 1800.07574 |
| *SEC22* | 0.095689845 | 0.57345428 | 0.99979869 | 12072.9675 | 12900.8845 |
| *BUD9* | 0.096403253 | 0.57363374 | 0.99979869 | 10489.7075 | 11214.5949 |
| *GPI8* | 0.098255375 | 0.57415965 | 0.99979869 | 7773.8751 | 8321.76246 |
| *FMP40* | 0.101036773 | 0.57451523 | 0.99979869 | 5617.98804 | 6025.53828 |
| *IQG1* | 0.09391163 | 0.57459008 | 0.99979869 | 16908.5929 | 18045.8618 |
| *IMH1* | 0.092716311 | 0.57537621 | 0.99979869 | 23013.3627 | 24540.896 |
| *VAN1* | 0.09251589 | 0.57637189 | 0.99979869 | 22435.8581 | 23921.7355 |
| *YJL068C* | 0.09416766 | 0.57714453 | 0.99979869 | 13385.0345 | 14287.8449 |
| *CAF40* | 0.094435586 | 0.57778666 | 0.99979869 | 12191.0048 | 13015.6957 |
| *MRS2* | 0.111293704 | 0.57785109 | 0.99979869 | 2583.16241 | 2790.32242 |
| *YDL086W* | 0.092921458 | 0.57799698 | 0.99979869 | 17174.9084 | 18317.5134 |
| *FMP30* | 0.099426347 | 0.57924614 | 0.99979869 | 5745.78044 | 6155.72599 |
| *HAL5* | 0.090406667 | 0.57924884 | 0.99979869 | 42885.5687 | 45658.9827 |
| *SNP1* | 0.105430698 | 0.57932095 | 0.99979869 | 3528.43597 | 3795.9456 |
| *YIL086C* | 0.243711215 | 0.57955293 | 0.99979869 | 207.78459 | 246.02402 |
| *STV1* | 0.091584302 | 0.57977868 | 0.99979869 | 22509.9972 | 23985.2917 |
| *IRC23* | 0.112237282 | 0.57989555 | 0.99979869 | 2386.10848 | 2579.15181 |
| *SRO7* | 0.097041235 | 0.58002471 | 0.99979869 | 7327.08946 | 7836.89013 |
| *TAX4* | 0.097683285 | 0.58052022 | 0.99979869 | 6665.69062 | 7132.64637 |
| *SMA1* | 0.319458087 | 0.58078875 | 0.99979869 | 109.257625 | 136.338311 |
| *MPC54* | 0.142068846 | 0.58089888 | 0.99979869 | 895.522317 | 988.196479 |
| *DAM1* | 0.098291551 | 0.58119705 | 0.99979869 | 6095.99015 | 6525.78712 |
| *tD(GUC)J1* | 0.978421895 | 0.58144383 | 0.99979869 | 7.80411605 | 15.3765012 |
| *MRX8* | 0.104805902 | 0.58158849 | 0.99979869 | 3511.85222 | 3776.4687 |
| *WHI5* | 0.097626828 | 0.58189884 | 0.99979869 | 6413.03236 | 6862.01995 |
| *HLR1* | 0.125863653 | 0.58198981 | 0.99979869 | 1372.54891 | 1497.67122 |
| *YPR158W-B* | 0.143320542 | 0.58206593 | 0.99979869 | 860.403794 | 950.267776 |
| *YSP3* | 0.105206482 | 0.58209049 | 0.99979869 | 3386.98637 | 3643.20569 |
| *VMA1* | 0.088562955 | 0.58225827 | 0.99979869 | 148259.67 | 157646.041 |
| *BCK1* | 0.093708902 | 0.58230664 | 0.99979869 | 11019.4119 | 11758.923 |

| | | | | | |
|---|---|---|---|---|---|
| *FRA1* | 0.089667071 | 0.58236627 | 0.99979869 | 40404.8353 | 42995.7727 |
| *YLL006W-A* | 0.860027651 | 0.58241911 | 0.99979869 | 10.7306596 | 19.4769016 |
| *PIM1* | 0.089888264 | 0.58248195 | 0.99979869 | 34735.145 | 36968.1842 |
| *TOP3* | 0.109140158 | 0.58281747 | 0.99979869 | 2659.25254 | 2868.23003 |
| *SCL1* | 0.089702373 | 0.58303995 | 0.99979869 | 35477.5116 | 37753.4109 |
| *PHO2* | 0.096214684 | 0.58306704 | 0.99979869 | 7291.97093 | 7794.86102 |
| *GPI13* | 0.091388872 | 0.58363673 | 0.99979869 | 17172.9574 | 18295.9863 |
| *ECM13* | 0.160514811 | 0.58372459 | 0.99979869 | 597.990392 | 668.365253 |
| *YDL144C* | 0.09279721 | 0.58388186 | 0.99979869 | 12086.6247 | 12889.6084 |
| *NAP1* | 0.088499169 | 0.58411299 | 0.99979869 | 74286.4052 | 78986.0115 |
| *NVJ2* | 0.090900988 | 0.58419524 | 0.99979869 | 18979.6102 | 20213.9485 |
| *YDR042C* | 0.209472209 | 0.58425346 | 0.99979869 | 291.678837 | 337.257927 |
| *MIS1* | 0.088565826 | 0.58436681 | 0.99979869 | 64139.1033 | 68199.9084 |
| *FUR1* | 0.08932703 | 0.58462464 | 0.99979869 | 34580.0382 | 36788.7917 |
| *YGR050C* | 0.133590798 | 0.58472168 | 0.99979869 | 1065.26184 | 1168.61409 |
| *GLT1* | 0.089498802 | 0.5856656 | 0.99979869 | 27631.4484 | 29399.8703 |
| *YGL108C* | 0.096393314 | 0.58575232 | 0.99979869 | 6509.6083 | 6959.40446 |
| *TRX2* | 0.088207466 | 0.58576719 | 0.99979869 | 63511.8474 | 67516.1667 |
| *CTK3* | 0.100506959 | 0.58632851 | 0.99979869 | 4327.38235 | 4639.60297 |
| *MEX67* | 0.090154547 | 0.58651127 | 0.99979869 | 19745.3891 | 21018.6521 |
| *ASK10* | 0.0905604 | 0.58697688 | 0.99979869 | 16823.7232 | 17913.6239 |
| *KRE9* | 0.089051437 | 0.58732847 | 0.99979869 | 27702.6609 | 29466.5019 |
| *YKL077W* | 0.088748201 | 0.58746215 | 0.99979869 | 31524.7268 | 33524.8731 |
| *RPS30A* | 0.087121651 | 0.58779516 | 0.99979869 | 146424.727 | 155539.461 |
| *MSS51* | 0.090063266 | 0.58842093 | 0.99979869 | 17353.4275 | 18471.2784 |
| *RPS29A* | 0.087341345 | 0.58907325 | 0.99979869 | 63558.6721 | 67525.3926 |
| *YMR160W* | 0.096140191 | 0.58915432 | 0.99979869 | 6010.14487 | 6424.30221 |
| *ATG38* | 0.106429251 | 0.58944455 | 0.99979869 | 2748.99988 | 2959.46394 |
| *YDR306C* | 0.09372951 | 0.58946331 | 0.99979869 | 7989.46381 | 8525.75738 |
| *YPR117W* | 0.090163 | 0.58968951 | 0.99979869 | 15383.8638 | 16375.9738 |
| *SCS2* | 0.087099355 | 0.59075867 | 0.99979869 | 52774.3593 | 56058.6231 |
| *VBA4* | 0.088043391 | 0.59082195 | 0.99979869 | 28810.8454 | 30623.8398 |
| *ASI1* | 0.09120203 | 0.59084548 | 0.99979869 | 11368.6461 | 12110.5324 |
| *PMT2* | 0.086441932 | 0.59138766 | 0.99979869 | 91636.9062 | 97295.3241 |
| *KAP104* | 0.087823851 | 0.59166649 | 0.99979869 | 28771.8248 | 30577.7103 |
| *GLO3* | 0.087664635 | 0.59174741 | 0.99979869 | 30792.1154 | 32721.1946 |
| *PST2* | 0.087287798 | 0.59175652 | 0.99979869 | 38016.7758 | 40387.9181 |
| *YAR062W* | 0.543599997 | 0.59185048 | 0.99979869 | 30.2409497 | 44.0793035 |
| *PUS4* | 0.090675234 | 0.5923913 | 0.99979869 | 11610.5737 | 12363.7321 |
| *PER1* | 0.091513348 | 0.59255756 | 0.99979869 | 9861.47614 | 10507.2758 |
| *RPL34A* | 0.08605838 | 0.59269313 | 0.99979869 | 98309.4254 | 104352.113 |
| *tI(AAU)E2* | 0.808497465 | 0.59304552 | 0.99979869 | 11.7061741 | 20.5020016 |
| *YOR186W* | 0.219488055 | 0.59327241 | 0.99979869 | 243.878627 | 283.952723 |
| *DAP1* | 0.091580141 | 0.59383548 | 0.99979869 | 9201.05282 | 9804.05718 |
| *MKK2* | 0.096205545 | 0.59387754 | 0.99979869 | 5202.41886 | 5561.16794 |

| | | | | | |
|---|---|---|---|---|---|
| *YJR056C* | 0.096800519 | 0.59435749 | 0.99979869 | 4858.06224 | 5195.20721 |
| *FSH2* | 0.09391757 | 0.59460804 | 0.99979869 | 6487.17147 | 6923.52595 |
| *CUB1* | 0.087816726 | 0.59499804 | 0.99979869 | 20278.02 | 21550.679 |
| *SKI7* | 0.095950584 | 0.59518399 | 0.99979869 | 5142.91248 | 5496.58664 |
| *MED11* | 0.099724218 | 0.59520501 | 0.99979869 | 3756.70636 | 4025.56802 |
| *TDA4* | 0.093116593 | 0.5952808 | 0.99979869 | 6988.58592 | 7454.52779 |
| *YOR032W-A* | 0.287567412 | 0.59539231 | 0.99979869 | 126.816886 | 154.790112 |
| *AIM29* | 0.088519819 | 0.59573821 | 0.99979869 | 15481.4152 | 16461.0571 |
| *CKS1* | 0.091848491 | 0.59606556 | 0.99979869 | 8047.01916 | 8575.98728 |
| *tR(UCU)G3* | 0.362762136 | 0.59620191 | 0.99979869 | 74.1391025 | 95.3343076 |
| *SHY1* | 0.093174725 | 0.59624353 | 0.99979869 | 6711.5398 | 7159.29897 |
| *STR3* | 0.115430273 | 0.59682251 | 0.99979869 | 1641.79091 | 1778.54864 |
| *YBR062C* | 0.09236602 | 0.59697897 | 0.99979869 | 7242.21969 | 7721.05382 |
| *MOT3* | 0.088789745 | 0.59734713 | 0.99979869 | 13049.4575 | 13877.8049 |
| *YBP2* | 0.093085875 | 0.59822265 | 0.99979869 | 6350.59943 | 6773.86134 |
| *GCD7* | 0.086938435 | 0.59829266 | 0.99979869 | 20350.2081 | 21614.2352 |
| *COX20* | 0.091888574 | 0.59831739 | 0.99979869 | 7344.64872 | 7827.66422 |
| *TRK2* | 0.090364735 | 0.5984558 | 0.99979869 | 9130.81578 | 9721.02408 |
| *SDH5* | 0.09428166 | 0.5985117 | 0.99979869 | 5523.36313 | 5896.37567 |
| *GLE1* | 0.088986081 | 0.59889085 | 0.99979869 | 11459.3689 | 12188.44 |
| *FKH2* | 0.088634488 | 0.59909149 | 0.99979869 | 12187.1027 | 12959.3152 |
| *RGA2* | 0.090211699 | 0.59939307 | 0.99979869 | 8981.56206 | 9561.10846 |
| *NUM1* | 0.085953182 | 0.59995282 | 0.99979869 | 25145.8374 | 26689.5057 |
| *SWD1* | 0.093118935 | 0.60014244 | 0.99979869 | 5952.58952 | 6349.46991 |
| *MET28* | 0.109075409 | 0.60016058 | 0.99979869 | 2034.92326 | 2194.73928 |
| *CTF18* | 0.094019182 | 0.60061135 | 0.99979869 | 5339.96641 | 5699.55645 |
| *tE(UUC)K* | 0.434101267 | 0.60112214 | 0.99979869 | 47.8002108 | 64.5813052 |
| *TAO3* | 0.086061164 | 0.6012012 | 0.99979869 | 21199.8812 | 22502.997 |
| *NHX1* | 0.08674104 | 0.60152777 | 0.99979869 | 16566.1873 | 17592.7676 |
| *PTP2* | 0.090678338 | 0.60233462 | 0.99979869 | 7447.07774 | 7930.17423 |
| *YJR015W* | 0.085348755 | 0.60257881 | 0.99979869 | 24325.4297 | 25807.9197 |
| *tL(CAA)G1* | 0.763409652 | 0.60265684 | 0.99979869 | 12.6816886 | 21.5271017 |
| *YBR200W-A* | 0.763409652 | 0.60265684 | 0.99979869 | 12.6816886 | 21.5271017 |
| *SRL3* | 0.091610971 | 0.60293728 | 0.99979869 | 6472.53875 | 6896.87335 |
| *VMA13* | 0.083456171 | 0.60325928 | 0.99979869 | 83554.7685 | 88530.7184 |
| *RNH203* | 0.095689487 | 0.60330044 | 0.99979869 | 4274.70457 | 4567.84596 |
| *VPS72* | 0.087048364 | 0.60351785 | 0.99979869 | 13349.916 | 14180.2094 |
| *CDC39* | 0.084480327 | 0.603561 | 0.99979869 | 32671.9318 | 34642.2322 |
| *DLD1* | 0.086120317 | 0.60378512 | 0.99979869 | 16698.8573 | 17726.0306 |
| *HSE1* | 0.087304594 | 0.60412226 | 0.99979869 | 12155.8863 | 12914.2108 |
| *PTC6* | 0.091499683 | 0.60416384 | 0.99979869 | 6298.89717 | 6711.33024 |
| *RTT105* | 0.107236863 | 0.60489388 | 0.99979869 | 2063.21318 | 2222.41698 |
| *GTS1* | 0.086081074 | 0.6049503 | 0.99979869 | 15493.1214 | 16445.6806 |
| *CAM1* | 0.082769674 | 0.60580035 | 0.99979869 | 88679.1462 | 93915.5691 |
| *HXT14* | 0.388721188 | 0.6061449 | 0.99979869 | 59.5063849 | 77.9076062 |

| | | | | | |
|---|---|---|---|---|---|
| *CYB5* | 0.090361705 | 0.60619435 | 0.99979869 | 6756.41347 | 7193.12727 |
| *GYP5* | 0.086757121 | 0.60634806 | 0.99979869 | 12043.7021 | 12790.1737 |
| *YKL222C* | 0.09591592 | 0.6065565 | 0.99979869 | 3891.32736 | 4158.83103 |
| *YHL041W* | 0.535478731 | 0.60817246 | 0.99979869 | 28.2899207 | 41.0040033 |
| *PTC4* | 0.088820799 | 0.60830604 | 0.99979869 | 7686.07879 | 8174.14805 |
| *tK(CUU)G3* | 0.837066575 | 0.60846913 | 0.99979869 | 9.75514506 | 17.4267014 |
| *YLR464W* | 0.837066575 | 0.60846913 | 0.99979869 | 9.75514506 | 17.4267014 |
| *COS12* | 0.245178423 | 0.60911986 | 0.99979869 | 167.788495 | 198.869416 |
| *DUF1* | 0.084055998 | 0.60978468 | 0.99979869 | 19576.6251 | 20751.101 |
| *YOL057W* | 0.083943017 | 0.61024887 | 0.99979869 | 19529.8004 | 20699.846 |
| *PPZ2* | 0.086099328 | 0.61039604 | 0.99979869 | 11052.5794 | 11732.2704 |
| *BCH1* | 0.083394506 | 0.61046611 | 0.99979869 | 23507.9486 | 24906.8567 |
| *SPP41* | 0.083982968 | 0.61082033 | 0.99979869 | 18341.6237 | 19441.0231 |
| *OXA1* | 0.083307431 | 0.6117216 | 0.99979869 | 21421.323 | 22694.6907 |
| *RET2* | 0.081855026 | 0.61211662 | 0.99979869 | 43461.1223 | 45998.2909 |
| *AIM2* | 0.085876939 | 0.61251656 | 0.99979869 | 10346.3069 | 10980.8721 |
| *CDC25* | 0.08296972 | 0.61320522 | 0.99979869 | 21075.9909 | 22323.6045 |
| *GIS3* | 0.086716131 | 0.6136817 | 0.99979869 | 8481.12312 | 9006.52932 |
| *YJL043W* | 0.162677869 | 0.61420446 | 0.99979869 | 448.736673 | 502.29904 |
| *FBP1* | 0.125336282 | 0.61483235 | 0.99979869 | 975.514506 | 1064.05388 |
| *RSE1* | 0.083937896 | 0.61487848 | 0.99979869 | 13778.1669 | 14603.5758 |
| *BNA2* | 0.135987727 | 0.61496744 | 0.99979869 | 747.244112 | 821.105166 |
| *SPT7* | 0.083413216 | 0.61589253 | 0.99979869 | 14742.9507 | 15620.475 |
| *DET1* | 0.085299463 | 0.61609833 | 0.99979869 | 9663.4467 | 10252.0259 |
| *PRE5* | 0.080951839 | 0.61640292 | 0.99979869 | 38262.6055 | 40470.9512 |
| *SNF3* | 0.088901492 | 0.61663316 | 0.99979869 | 5717.49052 | 6080.89369 |
| *HRA1* | 0.129053556 | 0.61726585 | 0.99979869 | 863.330338 | 944.117175 |
| *KOG1* | 0.083911387 | 0.61774909 | 0.99979869 | 11657.3983 | 12355.5313 |
| *IST1* | 0.086431202 | 0.61806887 | 0.99979869 | 7422.68988 | 7880.96943 |
| *YIP4* | 0.081980885 | 0.6181674 | 0.99979869 | 19052.7738 | 20166.7939 |
| *RRI1* | 0.092756991 | 0.61834298 | 0.99979869 | 3817.18826 | 4070.67242 |
| *tK(CUU)P* | 0.782025353 | 0.61965852 | 0.99979869 | 10.7306596 | 18.4518015 |
| *SWA2* | 0.084881407 | 0.61996388 | 0.99979869 | 8708.418 | 9236.15174 |
| *YPL225W* | 0.080268645 | 0.62017107 | 0.99979869 | 32432.9308 | 34288.5726 |
| *SPA2* | 0.080803125 | 0.62025443 | 0.99979869 | 24506.8754 | 25918.6305 |
| *YBR219C* | 0.607584818 | 0.62044898 | 0.99979869 | 19.5102901 | 29.7279024 |
| *tS(AGA)J* | 0.541017025 | 0.62159656 | 0.99979869 | 25.3633772 | 36.9036029 |
| *HST4* | 0.103968207 | 0.62276756 | 0.99979869 | 1844.69793 | 1982.54356 |
| *IZH3* | 0.084194405 | 0.623071 | 0.99979869 | 8519.16818 | 9031.13172 |
| *YPR148C* | 0.079600193 | 0.6232859 | 0.99979869 | 30432.1505 | 32158.4147 |
| *YLR281C* | 0.105026691 | 0.62348811 | 0.99979869 | 1744.21994 | 1875.93315 |
| *EMT4* | 0.878886562 | 0.62418651 | 0.99979869 | 7.80411605 | 14.3514011 |
| *ARH1* | 0.085531514 | 0.62421207 | 0.99979869 | 6701.78466 | 7111.11927 |
| *YNL184C* | 0.122928994 | 0.62462949 | 0.99979869 | 941.371498 | 1025.10008 |
| *ADH7* | 0.105496609 | 0.62483727 | 0.99979869 | 1678.86047 | 1806.22634 |

| | | | | | |
|---|---|---|---|---|---|
| *GAT2* | 0.080763144 | 0.62487889 | 0.99979869 | 16256.9492 | 17192.9786 |
| *ARR1* | 0.086490074 | 0.6249885 | 0.99979869 | 5802.36028 | 6160.85149 |
| *GAL4* | 0.088469597 | 0.62499767 | 0.99979869 | 4707.83301 | 5005.5637 |
| *YHL017W* | 0.081040969 | 0.62540735 | 0.99979869 | 14453.2229 | 15288.3426 |
| *YMR254C* | 0.173225447 | 0.62569541 | 0.99979869 | 347.283164 | 391.588231 |
| *BMH1* | 0.077246546 | 0.62650875 | 0.99979869 | 203926.43 | 215142.88 |
| *NCE101* | 0.093150308 | 0.62682077 | 0.99979869 | 3101.16061 | 3307.99796 |
| *RRD1* | 0.080841098 | 0.62697343 | 0.99979869 | 13709.8809 | 14500.0407 |
| *ELP6* | 0.083403827 | 0.62748508 | 0.99979869 | 8026.53336 | 8504.23028 |
| *SLT2* | 0.078325332 | 0.6276598 | 0.99979869 | 33884.4964 | 35774.9678 |
| *PHO85* | 0.080454955 | 0.6278551 | 0.99979869 | 14304.9447 | 15125.3517 |
| *IFH1* | 0.079530263 | 0.62843748 | 0.99979869 | 18068.4797 | 19092.489 |
| *ENV7* | 0.089460557 | 0.62900218 | 0.99979869 | 3900.107 | 4149.60513 |
| *snR7-L* | 0.282427384 | 0.62953889 | 0.99979869 | 105.355567 | 128.13751 |
| *SED5* | 0.081617009 | 0.63032413 | 0.99979869 | 9593.20965 | 10151.5661 |
| *CNM67* | 0.088282074 | 0.63035094 | 0.99979869 | 4176.1776 | 4439.70845 |
| *AIM10* | 0.085192894 | 0.63070027 | 0.99979869 | 5638.47385 | 5981.45898 |
| *MRP4* | 0.080013696 | 0.63076437 | 0.99979869 | 13231.8788 | 13986.4655 |
| *AFI1* | 0.089508673 | 0.63165755 | 0.99979869 | 3656.22837 | 3890.25481 |
| *EPL1* | 0.080559714 | 0.6317033 | 0.99979869 | 11031.118 | 11664.6138 |
| *PHD1* | 0.079231724 | 0.63222641 | 0.99979869 | 14761.4855 | 15594.8475 |
| *ABF2* | 0.077374745 | 0.63244292 | 0.99979869 | 29556.1385 | 31184.5696 |
| *ECM2* | 0.08685607 | 0.63275416 | 0.99979869 | 4475.66055 | 4753.38908 |
| *OAR1* | 0.111433047 | 0.63285358 | 0.99979869 | 1217.4421 | 1315.2034 |
| *PDI1* | 0.075618703 | 0.6330326 | 0.99979869 | 238528.905 | 251364.791 |
| *HNT1* | 0.07670343 | 0.63343939 | 0.99979869 | 38558.1864 | 40663.67 |
| *ESL1* | 0.079565039 | 0.63454421 | 0.99979869 | 11702.272 | 12365.7823 |
| *LPX1* | 0.081221572 | 0.63559304 | 0.99979869 | 8104.57452 | 8573.93708 |
| *GGA2* | 0.076609902 | 0.63567332 | 0.99979869 | 29037.1648 | 30620.7645 |
| *NUT1* | 0.078731565 | 0.63579277 | 0.99979869 | 13255.2911 | 13998.7667 |
| *SOP4* | 0.077534709 | 0.63599699 | 0.99979869 | 18710.3682 | 19743.4276 |
| *SPT21* | 0.090003532 | 0.63646101 | 0.99979869 | 3179.20178 | 3383.85537 |
| *CAP1* | 0.077428697 | 0.6371175 | 0.99979869 | 17618.7675 | 18590.19 |
| *YLR036C* | 0.090151285 | 0.63753308 | 0.99979869 | 3078.72378 | 3277.24496 |
| *tV(AAC)O* | 0.283683883 | 0.63783675 | 0.99979869 | 98.5269651 | 119.93671 |
| *SFP1* | 0.07861974 | 0.63829396 | 0.99979869 | 11682.7617 | 12337.0795 |
| *HRD3* | 0.078065399 | 0.63835831 | 0.99979869 | 13320.6506 | 14061.2978 |
| *ARC1* | 0.074594278 | 0.63847674 | 0.99979869 | 104596.616 | 110147.004 |
| *HRR25* | 0.075846323 | 0.63905544 | 0.99979869 | 27997.2663 | 29508.531 |
| *SSL2* | 0.077344853 | 0.63910224 | 0.99979869 | 15458.0029 | 16309.3423 |
| *NUP1* | 0.076063478 | 0.63915896 | 0.99979869 | 24794.6522 | 26136.9768 |
| *PXP1* | 0.082738023 | 0.63921569 | 0.99979869 | 5753.58456 | 6093.19489 |
| *VPS35* | 0.076102457 | 0.63929685 | 0.99979869 | 23968.3914 | 25266.6668 |
| *RPN6* | 0.074979711 | 0.63952936 | 0.99979869 | 46449.1232 | 48927.0018 |
| *PHA2* | 0.087314308 | 0.64079795 | 0.99979869 | 3546.97074 | 3768.2679 |

| | | | | | |
|---|---|---|---|---|---|
| *YBR053C* | 0.078544797 | 0.64105197 | 0.99979869 | 10206.8083 | 10777.9023 |
| *STT4* | 0.075608281 | 0.64178308 | 0.99979869 | 22742.1697 | 23965.8148 |
| *ESC1* | 0.076292831 | 0.64183284 | 0.99979869 | 17404.1543 | 18349.2915 |
| *RCN2* | 0.075640602 | 0.64186925 | 0.99979869 | 22221.2449 | 23417.3863 |
| *PET20* | 0.082717599 | 0.64217562 | 0.99979869 | 5262.90076 | 5573.46914 |
| *DUO1* | 0.08614228 | 0.64268774 | 0.99979869 | 3737.19607 | 3967.13732 |
| *MPS3* | 0.082003966 | 0.64269951 | 0.99979869 | 5622.86561 | 5951.73107 |
| *MTR10* | 0.077683044 | 0.64313041 | 0.99979869 | 10955.0279 | 11561.0787 |
| *FUB1* | 0.081223124 | 0.64326361 | 0.99979869 | 6077.45537 | 6429.42771 |
| *MPD1* | 0.08101585 | 0.64412229 | 0.99979869 | 6062.82266 | 6413.02611 |
| *SSC1* | 0.072957141 | 0.64415615 | 0.99979869 | 225677.477 | 237383.451 |
| *RPS21B* | 0.073008315 | 0.64426955 | 0.99979869 | 169342.49 | 178132.666 |
| *SPR3* | 0.167885132 | 0.64431942 | 0.99979869 | 324.846331 | 364.935629 |
| *YCR025C* | 0.156627608 | 0.64521211 | 0.99979869 | 385.32823 | 429.516934 |
| *STE13* | 0.075271094 | 0.64579755 | 0.99979869 | 17656.8126 | 18602.4912 |
| *TIR2* | 0.101843606 | 0.6459466 | 0.99979869 | 1470.10036 | 1577.62903 |
| *YAR068W* | 0.145988963 | 0.64625045 | 0.99979869 | 460.442847 | 509.474741 |
| *TRM8* | 0.075676485 | 0.64629169 | 0.99979869 | 14911.7147 | 15714.7843 |
| *YIL015C-A* | 0.096333164 | 0.64645593 | 0.99979869 | 1856.40411 | 1984.59376 |
| *SAP190* | 0.07451908 | 0.64691475 | 0.99979869 | 21163.7872 | 22285.6758 |
| *CEM1* | 0.07860787 | 0.64812187 | 0.99979869 | 7338.79563 | 7749.75662 |
| *MRS6* | 0.073406327 | 0.64875064 | 0.99979869 | 29229.3411 | 30755.0527 |
| *OXP1* | 0.076119831 | 0.6492378 | 0.99979869 | 11020.3874 | 11617.4592 |
| *SUC2* | 0.075781343 | 0.64974654 | 0.99979869 | 11568.6265 | 12192.5404 |
| *YMR272W-B* | 0.361037755 | 0.64990634 | 0.99979869 | 52.6777833 | 67.6566054 |
| *DNA2* | 0.076577232 | 0.65031357 | 0.99979869 | 9463.46622 | 9979.3493 |
| *SPC105* | 0.0770362 | 0.65033629 | 0.99979869 | 8673.29947 | 9149.01823 |
| *PSE1* | 0.072267526 | 0.65048815 | 0.99979869 | 49583.4513 | 52130.4396 |
| *tN(GUU)K* | 0.298941294 | 0.65084516 | 0.99979869 | 79.9921895 | 98.4096079 |
| *TRR2* | 0.078456187 | 0.65113718 | 0.99979869 | 6689.10297 | 7062.93956 |
| *UTP11* | 0.075862774 | 0.65195918 | 0.99979869 | 10053.6525 | 10596.4595 |
| *CDC13* | 0.079761963 | 0.65214252 | 0.99979869 | 5455.07712 | 5765.16286 |
| *XDJ1* | 0.075388919 | 0.65231416 | 0.99979869 | 10925.7625 | 11511.8739 |
| *RAV1* | 0.075633732 | 0.65246263 | 0.99979869 | 10273.1433 | 10826.082 |
| *YGL188C-A* | 0.111291663 | 0.65257041 | 0.99979869 | 977.465535 | 1055.85308 |
| *YGL006W-A* | 0.134211163 | 0.65259755 | 0.99979869 | 549.214667 | 602.758848 |
| *MEP2* | 0.088688424 | 0.65295377 | 0.99979869 | 2527.55809 | 2687.81241 |
| *TRS120* | 0.07477322 | 0.65360162 | 0.99979869 | 11702.272 | 12324.7783 |
| *HNT2* | 0.080535348 | 0.65445464 | 0.99979869 | 4673.69 | 4942.00749 |
| *DOT5* | 0.076552167 | 0.65456062 | 0.99979869 | 7832.40597 | 8259.23136 |
| *IPT1* | 0.072340554 | 0.65462005 | 0.99979869 | 24293.2377 | 25542.4187 |
| *YBR016W* | 0.072751403 | 0.65496415 | 0.99979869 | 19575.6496 | 20588.11 |
| *YOR394C-A* | 0.425168468 | 0.65532862 | 0.99979869 | 35.1185222 | 47.1546038 |
| *MDN1* | 0.070706975 | 0.65580828 | 0.99979869 | 70152.1747 | 73675.9931 |
| *RAD30* | 0.077431222 | 0.65620894 | 0.99979869 | 6425.71405 | 6780.01194 |

| | | | | | |
|---|---|---|---|---|---|
| *TIM18* | 0.08524503 | 0.6566578 | 0.99979869 | 2961.66204 | 3141.93175 |
| *SIN4* | 0.074427672 | 0.65744814 | 0.99979869 | 10188.2735 | 10727.6724 |
| *ASF1* | 0.07414757 | 0.65760397 | 0.99979869 | 10742.3657 | 11308.9041 |
| *HOL1* | 0.072370896 | 0.65829517 | 0.99979869 | 16726.1717 | 17586.617 |
| *ARG81* | 0.074881304 | 0.65854256 | 0.99979869 | 8807.92048 | 9277.15574 |
| *YCL011C-A* | 0.086435958 | 0.65857932 | 0.99979869 | 2629.98711 | 2792.37262 |
| *BOL1* | 0.078920894 | 0.65861904 | 0.99979869 | 4938.05443 | 5215.70922 |
| *GAT4* | 0.200813299 | 0.6591885 | 0.99979869 | 187.298785 | 215.271017 |
| *DSK2* | 0.06985316 | 0.65952124 | 0.99979869 | 68033.3572 | 71408.4717 |
| *ERF2* | 0.083484014 | 0.65972007 | 0.99979869 | 3165.54457 | 3354.12747 |
| *AAD10* | 0.09948386 | 0.65988217 | 0.99979869 | 1346.21002 | 1442.31582 |
| *ECM33* | 0.069090129 | 0.66077816 | 0.99979869 | 185335.074 | 194426.632 |
| *MCP1* | 0.082128847 | 0.66082399 | 0.99979869 | 3439.66415 | 3641.15549 |
| *RPL17B* | 0.069173601 | 0.66083935 | 0.99979869 | 136299.862 | 142994.286 |
| *DOC1* | 0.090397929 | 0.66095912 | 0.99979869 | 2000.78025 | 2130.15797 |
| *tT(AGU)N1* | 0.595094126 | 0.66102535 | 0.99979869 | 15.6082321 | 23.5773019 |
| *JJJ1* | 0.073948324 | 0.66114242 | 0.99979869 | 9302.50633 | 9791.75598 |
| *RFT1* | 0.073535058 | 0.66174096 | 0.99979869 | 9818.5535 | 10331.9837 |
| *RRN7* | 0.085738228 | 0.66239297 | 0.99979869 | 2562.67661 | 2719.59052 |
| *TUB2* | 0.068949535 | 0.66266238 | 0.99979869 | 87895.808 | 92198.5265 |
| *YBR197C* | 0.076157461 | 0.66275268 | 0.99979869 | 6072.5778 | 6401.75001 |
| *TDA7* | 0.072069633 | 0.66288376 | 0.99979869 | 13030.9228 | 13698.4124 |
| *IKI3* | 0.069309993 | 0.66376552 | 0.99979869 | 41877.8622 | 43938.8648 |
| *YIF1* | 0.071631604 | 0.66383461 | 0.99979869 | 13813.2854 | 14516.4423 |
| *THI11* | 0.118900323 | 0.66400199 | 0.99979869 | 701.39493 | 761.649361 |
| *FRS1* | 0.06857433 | 0.66425946 | 0.99979869 | 88187.4869 | 92480.429 |
| *STU1* | 0.076285732 | 0.66458215 | 0.99979869 | 5613.11047 | 5917.90277 |
| *NEO1* | 0.07175495 | 0.66480953 | 0.99979869 | 12496.3408 | 13133.5822 |
| *RPS27B* | 0.068149564 | 0.66544755 | 0.99979869 | 119635.148 | 125422.02 |
| *TWF1* | 0.071894915 | 0.66548183 | 0.99979869 | 11559.8469 | 12150.5113 |
| *APL3* | 0.073138079 | 0.66566921 | 0.99979869 | 8745.48755 | 9200.27323 |
| *LSB6* | 0.075105734 | 0.66586937 | 0.99979869 | 6289.14202 | 6625.22183 |
| *INM2* | 0.075751405 | 0.66640643 | 0.99979869 | 5658.95965 | 5964.03228 |
| *YGR015C* | 0.085070988 | 0.66642713 | 0.99979869 | 2482.68442 | 2633.48211 |
| *PIS1* | 0.068731467 | 0.66691487 | 0.99979869 | 36750.558 | 38543.7631 |
| *VPS21* | 0.069766756 | 0.66758093 | 0.99979869 | 19170.8111 | 20120.6644 |
| *BIM1* | 0.071404819 | 0.66791217 | 0.99979869 | 11268.1681 | 11839.9059 |
| *NFS1* | 0.069601977 | 0.66836742 | 0.99979869 | 18994.2429 | 19933.0711 |
| *HST1* | 0.070418752 | 0.66871145 | 0.99979869 | 13940.1023 | 14637.4041 |
| *YNL046W* | 0.072739385 | 0.66882314 | 0.99979869 | 8140.66855 | 8561.63588 |
| *KEX2* | 0.06946166 | 0.66891471 | 0.99979869 | 19068.3821 | 20008.9285 |
| *APD1* | 0.068656611 | 0.66922337 | 0.99979869 | 26968.0985 | 28282.5113 |
| *TCO89* | 0.071393239 | 0.66950882 | 0.99979869 | 10312.1638 | 10835.3079 |
| *GMC2* | 0.252102871 | 0.66999257 | 0.99979869 | 102.429023 | 121.98691 |
| *YPR204W* | 0.081406425 | 0.67001595 | 0.99979869 | 2982.14785 | 3155.25805 |

| | | | | | |
|---|---|---|---|---|---|
| *LDB19* | 0.070102824 | 0.67014553 | 0.99979869 | 13815.2364 | 14503.116 |
| *YPL229W* | 0.084084933 | 0.67045991 | 0.99979869 | 2455.37001 | 2602.72911 |
| *YMR178W* | 0.070466876 | 0.67084038 | 0.99979869 | 11921.7628 | 12518.5222 |
| *YLL032C* | 0.074115834 | 0.67116547 | 0.99979869 | 5980.87944 | 6296.1647 |
| *FMP32* | 0.074706306 | 0.67238842 | 0.99979869 | 5310.70097 | 5592.94605 |
| *SLX5* | 0.071370949 | 0.67252219 | 0.99979869 | 8864.50032 | 9314.05934 |
| *BDF1* | 0.067216631 | 0.67294522 | 0.99979869 | 39868.3024 | 41769.753 |
| *tA(UGC)E* | 0.545463335 | 0.67303457 | 0.99979869 | 17.5592611 | 25.627502 |
| *SLI1* | 0.081901433 | 0.67307502 | 0.99979869 | 2704.12621 | 2862.07943 |
| *MBR1* | 0.078112977 | 0.67331977 | 0.99979869 | 3625.98742 | 3827.72371 |
| *YDR524C-B* | 0.065965242 | 0.67353909 | 0.99979869 | 517202.183 | 541399.432 |
| *SSP2* | 0.115676989 | 0.67371713 | 0.99979869 | 690.66427 | 748.32306 |
| *YDL183C* | 0.089451684 | 0.67391884 | 0.99979869 | 1716.90553 | 1826.72835 |
| *RTS1* | 0.067458271 | 0.67443235 | 0.99979869 | 26653.9829 | 27929.8768 |
| *YHR131C* | 0.069864605 | 0.67465298 | 0.99979869 | 10994.0485 | 11539.5516 |
| *SSA3* | 0.07348858 | 0.67476961 | 0.99979869 | 5743.82941 | 6043.99008 |
| *GUP2* | 0.110960011 | 0.67494317 | 0.99979869 | 774.558518 | 836.481667 |
| *RAD59* | 0.08148483 | 0.67507204 | 0.99979869 | 2676.8118 | 2832.35153 |
| *MAG2* | 0.069191209 | 0.67520385 | 0.99979869 | 12648.5211 | 13269.9206 |
| *CWC15* | 0.073540452 | 0.67533969 | 0.99979869 | 5595.55121 | 5888.17487 |
| *EFT2* | 0.0657171 | 0.67536792 | 0.99979869 | 182442.674 | 190945.392 |
| *GRX7* | 0.070449109 | 0.67569659 | 0.99979869 | 9120.08512 | 9576.48496 |
| *YNK1* | 0.067811818 | 0.67592362 | 0.99979869 | 18951.3203 | 19863.3643 |
| *MRPL9* | 0.070188442 | 0.67606887 | 0.99979869 | 9447.85799 | 9918.86839 |
| *THP1* | 0.075353975 | 0.67612305 | 0.99979869 | 4410.30108 | 4646.77867 |
| *SBA1* | 0.065795548 | 0.67626525 | 0.99979869 | 87548.5249 | 91633.6963 |
| *KEI1* | 0.071947836 | 0.67651586 | 0.99979869 | 6731.05009 | 7075.24076 |
| *GSF2* | 0.066390039 | 0.67661774 | 0.99979869 | 38952.2942 | 40786.6821 |
| *YIL077C* | 0.071338878 | 0.6767249 | 0.99979869 | 7374.88967 | 7748.73152 |
| *NBL1* | 0.082817112 | 0.67701945 | 0.99979869 | 2359.76959 | 2499.194 |
| *tG(GCC)G2* | 0.458554979 | 0.67723404 | 0.99979869 | 25.3633772 | 34.8534028 |
| *TRM112* | 0.068221233 | 0.67735044 | 0.99979869 | 14479.5618 | 15180.7071 |
| *ULI1* | 0.118983826 | 0.67737212 | 0.99979869 | 612.62311 | 665.289953 |
| *GPI16* | 0.066489223 | 0.67751124 | 0.99979869 | 31046.7247 | 32511.0491 |
| *PEX32* | 0.077075525 | 0.67810972 | 0.99979869 | 3545.99523 | 3740.5902 |
| *PGA1* | 0.074745011 | 0.67876388 | 0.99979869 | 4372.25602 | 4604.74957 |
| *HCS1* | 0.073134506 | 0.6788205 | 0.99979869 | 5258.9987 | 5532.46514 |
| *YFR054C* | 0.173551135 | 0.67924081 | 0.99979869 | 226.319365 | 255.24992 |
| *HKR1* | 0.068372966 | 0.67934372 | 0.99979869 | 12050.5307 | 12635.3836 |
| *CTS1* | 0.064720857 | 0.67979148 | 0.99979869 | 171962.722 | 179852.784 |
| *ENV11* | 0.072273575 | 0.67990696 | 0.99979869 | 5674.56788 | 5966.08248 |
| *DSF2* | 0.072011186 | 0.68008719 | 0.99979869 | 5845.28292 | 6144.44989 |
| *EDC3* | 0.067029351 | 0.68025962 | 0.99979869 | 17226.6107 | 18045.8618 |
| *YPT35* | 0.081295196 | 0.6805058 | 0.99979869 | 2441.71281 | 2583.25221 |
| *HFI1* | 0.071529726 | 0.68117573 | 0.99979869 | 6017.94899 | 6323.8424 |

| | | | | | |
|---|---|---|---|---|---|
| *RFC1* | 0.066347067 | 0.68119182 | 0.99979869 | 20947.223 | 21933.0413 |
| *YRA2* | 0.069849457 | 0.68218573 | 0.99979869 | 7542.67816 | 7916.84793 |
| *ECM15* | 0.067186228 | 0.68218953 | 0.99979869 | 13953.7595 | 14618.9523 |
| *YPL068C* | 0.084533616 | 0.68248739 | 0.99979869 | 1939.32284 | 2056.35076 |
| *SMP3* | 0.071529726 | 0.68259384 | 0.99979869 | 5724.31912 | 6015.28728 |
| *CNA1* | 0.068469612 | 0.68269363 | 0.99979869 | 9668.32427 | 10138.2398 |
| *YGL041W-A* | 0.072064554 | 0.68299737 | 0.99979869 | 5261.92525 | 5531.44004 |
| *NUR1* | 0.071286664 | 0.68322655 | 0.99979869 | 5790.65411 | 6083.96899 |
| *YOL159C-A* | 0.095736673 | 0.68326905 | 0.99979869 | 1153.05815 | 1232.1703 |
| *SGM1* | 0.066883356 | 0.68346286 | 0.99979869 | 13955.7105 | 14617.9272 |
| *PUF4* | 0.066011825 | 0.68401803 | 0.99979869 | 18143.5943 | 18993.0543 |
| *GPA1* | 0.065677269 | 0.6840473 | 0.99979869 | 20963.8067 | 21940.2171 |
| *DAL81* | 0.067384393 | 0.68411906 | 0.99979869 | 11559.8469 | 12112.5826 |
| *YKL023C-A* | 0.068505369 | 0.68422952 | 0.99979869 | 8850.84311 | 9281.25614 |
| *RAX2* | 0.068045299 | 0.68434969 | 0.99979869 | 9705.39382 | 10174.1183 |
| *CYK3* | 0.06642283 | 0.68478489 | 0.99979869 | 14631.7421 | 15321.1458 |
| *PAU12* | 0.216960921 | 0.6848045 | 0.99979869 | 128.767915 | 149.664612 |
| *THI22* | 0.0780093 | 0.68499303 | 0.99979869 | 2817.28589 | 2973.81534 |
| *RTG1* | 0.070371399 | 0.68552022 | 0.99979869 | 6077.45537 | 6381.24801 |
| *TFA2* | 0.067672818 | 0.68559235 | 0.99979869 | 9865.3782 | 10339.1594 |
| *FMP27* | 0.065340803 | 0.68660545 | 0.99979869 | 18686.9559 | 19552.759 |
| *DGA1* | 0.068357025 | 0.6869675 | 0.99979869 | 7993.36586 | 8381.21827 |
| *SUP5* | 0.371091757 | 0.68697309 | 0.99979869 | 38.0450657 | 49.2048039 |
| *AIP1* | 0.064301275 | 0.68729221 | 0.99979869 | 29473.2198 | 30816.5587 |
| *PIR5* | 0.11883564 | 0.6875327 | 0.99979869 | 556.043268 | 603.783948 |
| *VPS70* | 0.064691252 | 0.68765777 | 0.99979869 | 22485.6094 | 23516.821 |
| *YGL185C* | 0.071249294 | 0.6876774 | 0.99979869 | 5019.02213 | 5273.11482 |
| *RML2* | 0.066409019 | 0.68817255 | 0.99979869 | 11563.749 | 12108.4822 |
| *ASM4* | 0.06656522 | 0.68821087 | 0.99979869 | 11075.0162 | 11597.9823 |
| *YNL193W* | 0.069160589 | 0.68844295 | 0.99979869 | 6539.84925 | 6860.99485 |
| *BIT61* | 0.077929886 | 0.68859813 | 0.99979869 | 2632.91365 | 2779.04632 |
| *RMD8* | 0.068958074 | 0.68913174 | 0.99979869 | 6573.01674 | 6894.82315 |
| *MGM101* | 0.065923839 | 0.68954711 | 0.99979869 | 12073.943 | 12638.4589 |
| *APL6* | 0.064480592 | 0.68981284 | 0.99979869 | 19613.6947 | 20510.2024 |
| *PAH1* | 0.066330333 | 0.68996336 | 0.99979869 | 10575.5528 | 11073.1311 |
| *CHS3* | 0.064313109 | 0.69019023 | 0.99979869 | 20332.6489 | 21259.5506 |
| *UBC7* | 0.066160937 | 0.69036981 | 0.99979869 | 10767.7291 | 11273.0256 |
| *tL(CAA)L* | 0.282097924 | 0.69053601 | 0.99979869 | 68.2860154 | 83.0331066 |
| *MST1* | 0.071223977 | 0.69059436 | 0.99979869 | 4603.45295 | 4836.42219 |
| *FCY22* | 0.073462349 | 0.69060836 | 0.99979869 | 3638.66911 | 3828.74881 |
| *NOT3* | 0.064390882 | 0.69084778 | 0.99979869 | 18379.6688 | 19218.5763 |
| *SFT1* | 0.070929232 | 0.69113821 | 0.99979869 | 4688.32272 | 4924.58079 |
| *ARC35* | 0.063250616 | 0.69114147 | 0.99979869 | 33072.8683 | 34555.0987 |
| *tE(UUC)J* | 0.357835616 | 0.69117593 | 0.99979869 | 39.9960948 | 51.2550041 |
| *YCF1* | 0.062874078 | 0.69159396 | 0.99979869 | 41423.2725 | 43268.4494 |

| | | | | | |
|---|---|---|---|---|---|
| *DAL4* | 0.096816271 | 0.69214821 | 0.99979869 | 993.073767 | 1062.00368 |
| *RVS161* | 0.063092747 | 0.69259302 | 0.99979869 | 29277.1414 | 30585.9111 |
| *PFS1* | 0.221106903 | 0.69259444 | 0.99979869 | 116.086226 | 135.313211 |
| *YML039W* | 0.21875649 | 0.69355478 | 0.99979869 | 118.037255 | 137.363411 |
| *LEU3* | 0.065477792 | 0.69406586 | 0.99979869 | 10253.633 | 10729.7226 |
| *TUL1* | 0.066070434 | 0.69445377 | 0.99979869 | 8781.58158 | 9193.09753 |
| *tP(UGG)O3* | 0.216478862 | 0.69448494 | 0.99979869 | 119.988284 | 139.413611 |
| *YAP1* | 0.06284058 | 0.69533979 | 0.99979869 | 23393.8134 | 24435.3106 |
| *PCF11* | 0.067400594 | 0.69578672 | 0.99979869 | 6485.22044 | 6795.38844 |
| *DFM1* | 0.064762785 | 0.69583032 | 0.99979869 | 11048.6773 | 11555.9532 |
| *TFB1* | 0.065158233 | 0.69672801 | 0.99979869 | 9519.07055 | 9958.84729 |
| *EAR1* | 0.063805985 | 0.6971056 | 0.99979869 | 13337.2343 | 13940.336 |
| *FOX2* | 0.069187691 | 0.69714818 | 0.99979869 | 4811.23754 | 5047.5928 |
| *MUM2* | 0.066717597 | 0.69762009 | 0.99979869 | 6737.87869 | 7056.78896 |
| *PCK1* | 0.090830367 | 0.69766818 | 0.99979869 | 1158.91123 | 1234.2205 |
| *EMC10* | 0.062951236 | 0.69796086 | 0.99979869 | 16783.7271 | 17532.2867 |
| *VPS5* | 0.064187744 | 0.69840045 | 0.99979869 | 10954.0524 | 11452.4181 |
| *TMC1* | 0.061423829 | 0.69907361 | 0.99979869 | 33540.1397 | 34998.967 |
| *YTA7* | 0.061390415 | 0.69946299 | 0.99979869 | 32176.3705 | 33575.103 |
| *SPO77* | 0.107625691 | 0.69946603 | 0.99979869 | 654.570234 | 705.268856 |
| *ABP1* | 0.060883507 | 0.69948525 | 0.99979869 | 51082.8171 | 53284.7023 |
| *CHD1* | 0.062028298 | 0.69976299 | 0.99979869 | 20954.0516 | 21874.6106 |
| *PAP1* | 0.064295975 | 0.69991321 | 0.99979869 | 9752.21852 | 10196.6705 |
| *TOF1* | 0.067611841 | 0.69993 | 0.99979869 | 5395.57073 | 5654.45205 |
| *ARP3* | 0.060934454 | 0.69993877 | 0.99979869 | 43328.4523 | 45197.6877 |
| *YPQ1* | 0.061933932 | 0.70007518 | 0.99979869 | 21190.1261 | 22119.6096 |
| *BUD6* | 0.063970209 | 0.70039425 | 0.99979869 | 10266.3147 | 10731.7728 |
| *SMX2* | 0.070764583 | 0.70045792 | 0.99979869 | 3679.64072 | 3864.62731 |
| *YPR098C* | 0.066262805 | 0.70066013 | 0.99979869 | 6424.73854 | 6726.70674 |
| *ATP25* | 0.06612434 | 0.70085208 | 0.99979869 | 6521.31447 | 6827.16654 |
| *MPS2* | 0.076432717 | 0.70106789 | 0.99979869 | 2292.45909 | 2417.18599 |
| *SDD3* | 0.061247045 | 0.70109018 | 0.99979869 | 27333.9165 | 28519.3094 |
| *YER085C* | 0.287259739 | 0.70109417 | 0.99979869 | 60.4818994 | 73.8072059 |
| *SDH6* | 0.08342648 | 0.70117469 | 0.99979869 | 1531.55777 | 1622.73343 |
| *SEH1* | 0.062203928 | 0.70127767 | 0.99979869 | 16653.9836 | 17387.7476 |
| *YML020W* | 0.066310373 | 0.70132206 | 0.99979869 | 6213.05189 | 6505.28512 |
| *RPN10* | 0.060766972 | 0.70135122 | 0.99979869 | 37275.3848 | 38878.9708 |
| *MSB3* | 0.065331452 | 0.70221753 | 0.99979869 | 7053.94539 | 7380.72059 |
| *SCP1* | 0.068892262 | 0.7022554 | 0.99979869 | 4272.75354 | 4481.73756 |
| *YRF1-4* | 0.084037392 | 0.70254852 | 0.99979869 | 1456.44316 | 1543.80072 |
| *YER134C* | 0.064389335 | 0.70259172 | 0.99979869 | 8298.7019 | 8677.47219 |
| *ACB1* | 0.060130715 | 0.70260191 | 0.99979869 | 54415.1747 | 56731.0887 |
| *tK(CUU)F* | 0.61210085 | 0.70282849 | 0.99979869 | 10.7306596 | 16.4016013 |
| *EMT2* | 0.507631408 | 0.70298939 | 0.99979869 | 16.5837466 | 23.5773019 |
| *CSC1* | 0.061425762 | 0.70301828 | 0.99979869 | 19568.821 | 20419.9936 |

| | | | | | |
|---|---|---|---|---|---|
| *YER010C* | 0.06553652 | 0.7039112 | 0.99979869 | 6353.52598 | 6648.79913 |
| *WHI4* | 0.06296223 | 0.70396645 | 0.99979869 | 10709.1982 | 11186.9172 |
| *TNA1* | 0.059876723 | 0.70475855 | 0.99979869 | 41713.9758 | 43481.6702 |
| *UBP13* | 0.061998898 | 0.70547705 | 0.99979869 | 12741.195 | 13300.6736 |
| *MRM2* | 0.064768398 | 0.70584222 | 0.99979869 | 6676.42128 | 6982.98176 |
| *YME2* | 0.061326226 | 0.70589192 | 0.99979869 | 15502.8765 | 16176.0793 |
| *MMS2* | 0.064214199 | 0.70596863 | 0.99979869 | 7329.04048 | 7662.62311 |
| *YFL065C* | 0.187007606 | 0.70636915 | 0.99979869 | 152.180263 | 173.241914 |
| *ECM19* | 0.069541183 | 0.70664623 | 0.99979869 | 3541.11766 | 3715.9878 |
| *NAB3* | 0.059598756 | 0.70684072 | 0.99979869 | 35179.0041 | 36662.7044 |
| *YRF1-5* | 0.140133905 | 0.70693714 | 0.99979869 | 300.458468 | 331.107326 |
| *BRN1* | 0.067390913 | 0.70696642 | 0.99979869 | 4426.88483 | 4638.57787 |
| *TUS1* | 0.063947317 | 0.70729593 | 0.99979869 | 7257.82793 | 7586.76571 |
| *ROM2* | 0.061345082 | 0.70756335 | 0.99979869 | 13451.3695 | 14035.6703 |
| *EMP24* | 0.059057404 | 0.70771451 | 0.99979869 | 49297.6256 | 51357.5141 |
| *VHR1* | 0.062204282 | 0.70784032 | 0.99979869 | 10295.5801 | 10749.1995 |
| *RPL26A* | 0.058463557 | 0.70835087 | 0.99979869 | 103330.399 | 107603.73 |
| *YOL038C-A* | 0.087074954 | 0.70840102 | 0.99979869 | 1170.61741 | 1243.4464 |
| *IRA1* | 0.059782912 | 0.70851248 | 0.99979869 | 23939.126 | 24951.9611 |
| *GRX4* | 0.064246156 | 0.70905635 | 0.99979869 | 6392.54656 | 6683.65253 |
| *SIP4* | 0.132071591 | 0.7091677 | 0.99979869 | 341.430077 | 374.16153 |
| *VHS2* | 0.061457858 | 0.70947031 | 0.99979869 | 11357.9154 | 11852.2071 |
| *TOR1* | 0.060648801 | 0.70970475 | 0.99979869 | 14411.2758 | 15030.0174 |
| *NUP42* | 0.064707306 | 0.70971991 | 0.99979869 | 5789.67859 | 6055.26618 |
| *HDA2* | 0.062562664 | 0.70977712 | 0.99979869 | 8501.60892 | 8878.39181 |
| *TAF4* | 0.060715954 | 0.71015626 | 0.99979869 | 13585.9905 | 14169.9584 |
| *GLC8* | 0.059312133 | 0.7107817 | 0.99979869 | 23367.4745 | 24348.1771 |
| *IRC13* | 0.574032849 | 0.71097468 | 0.99979869 | 11.7061741 | 17.4267014 |
| *YLR053C* | 0.116477372 | 0.71164539 | 0.99979869 | 462.393876 | 501.27394 |
| *CDH1* | 0.062558049 | 0.71177953 | 0.99979869 | 7710.46666 | 8052.16114 |
| *TOK1* | 0.060810557 | 0.71194121 | 0.99979869 | 11596.9164 | 12096.181 |
| *MAP1* | 0.058296456 | 0.7119456 | 0.99979869 | 40598.9627 | 42273.0772 |
| *DUG3* | 0.063556564 | 0.71197943 | 0.99979869 | 6372.06075 | 6659.05013 |
| *RIB7* | 0.065384987 | 0.71224215 | 0.99979869 | 4820.01717 | 5043.4924 |
| *KAP123* | 0.057851499 | 0.71239753 | 0.99979869 | 60994.0445 | 63489.5736 |
| *RIM8* | 0.06369398 | 0.71241071 | 0.99979869 | 6123.30455 | 6399.69981 |
| *POT1* | 0.090182537 | 0.71302212 | 0.99979869 | 974.538992 | 1037.40128 |
| *GAL83* | 0.059418969 | 0.71310751 | 0.99979869 | 17154.4226 | 17875.6952 |
| *FAS1* | 0.057320118 | 0.7132069 | 0.99979869 | 125012.184 | 130079.05 |
| *YKL050C* | 0.073430124 | 0.71346184 | 0.99979869 | 2220.27102 | 2336.20309 |
| *ACF2* | 0.062042393 | 0.7134717 | 0.99979869 | 7888.0103 | 8234.62896 |
| *YER145C-A* | 0.064929619 | 0.71347614 | 0.99979869 | 4915.6176 | 5141.90201 |
| *YJL077W-B* | 0.09719475 | 0.71354552 | 0.99979869 | 760.901315 | 813.929465 |
| *YGR240C-A* | 0.090716854 | 0.71370315 | 0.99979869 | 947.224585 | 1008.69848 |
| *MAL13* | 0.069204689 | 0.71431459 | 0.99979869 | 3028.97254 | 3177.81025 |

| | | | | | |
|---|---|---|---|---|---|
| *GUT1* | 0.059689592 | 0.71518004 | 0.99979869 | 13009.4615 | 13558.9988 |
| *CDC21* | 0.06318422 | 0.71537879 | 0.99979869 | 5919.42202 | 6184.42879 |
| *ERG25* | 0.057044278 | 0.71542083 | 0.99979869 | 76554.4764 | 79642.0756 |
| *VID30* | 0.059750702 | 0.71547943 | 0.99979869 | 12476.8305 | 13004.4196 |
| *ECI1* | 0.076249606 | 0.71562461 | 0.99979869 | 1786.16706 | 1883.10885 |
| *ECO1* | 0.078784078 | 0.71578129 | 0.99979869 | 1548.14152 | 1635.03463 |
| *RAD18* | 0.067120885 | 0.71624978 | 0.99979869 | 3516.72979 | 3684.20969 |
| *YHR130C* | 0.102894073 | 0.7164072 | 0.99979869 | 621.40274 | 667.340153 |
| *RPS26B* | 0.056472867 | 0.71665652 | 0.99979869 | 170911.117 | 177733.902 |
| *GTO1* | 0.074397946 | 0.716721 | 0.99979869 | 1960.78416 | 2064.55156 |
| *PRE7* | 0.057452458 | 0.71675848 | 0.99979869 | 33152.8605 | 34499.7433 |
| *PDR10* | 0.063627514 | 0.71721237 | 0.99979869 | 5179.00651 | 5412.52843 |
| *PDE2* | 0.061004175 | 0.7174361 | 0.99979869 | 8051.89673 | 8399.67007 |
| *RAD26* | 0.06190614 | 0.7174535 | 0.99979869 | 6749.58487 | 7045.51286 |
| *YNL165W* | 0.066236822 | 0.71749168 | 0.99979869 | 3729.39196 | 3904.60621 |
| *UBA2* | 0.059113272 | 0.71772181 | 0.99979869 | 12977.2695 | 13520.045 |
| *BUD5* | 0.068237527 | 0.71844467 | 0.99979869 | 2995.80505 | 3140.90665 |
| *BOI2* | 0.057970456 | 0.71860645 | 0.99979869 | 18770.8501 | 19540.4578 |
| *NNR2* | 0.059078069 | 0.71877424 | 0.99979869 | 12146.1311 | 12653.8354 |
| *SOR2* | 0.237540771 | 0.71895707 | 0.99979869 | 79.9921895 | 94.3092075 |
| *RIM101* | 0.057610702 | 0.71948473 | 0.99979869 | 20319.9672 | 21147.8147 |
| *GDI1* | 0.05682 | 0.71961486 | 0.99979869 | 33080.6724 | 34409.5344 |
| *SEG2* | 0.058571766 | 0.71965187 | 0.99979869 | 13419.1775 | 13975.1894 |
| *THI7* | 0.061050538 | 0.72017614 | 0.99979869 | 7009.07173 | 7312.03888 |
| *TAF11* | 0.061047811 | 0.72063013 | 0.99979869 | 6872.4997 | 7169.54997 |
| *ELO1* | 0.056838359 | 0.72074921 | 0.99979869 | 27055.8948 | 28143.0976 |
| *VAM6* | 0.059871846 | 0.72095853 | 0.99979869 | 8605.98897 | 8970.65082 |
| *DBF4* | 0.058958274 | 0.72102674 | 0.99979869 | 10794.068 | 11244.3228 |
| *SAN1* | 0.058413208 | 0.72104375 | 0.99979869 | 12718.7581 | 13244.2931 |
| *MLC1* | 0.056227766 | 0.7218207 | 0.99979869 | 36152.5676 | 37589.3949 |
| *YLR358C* | 0.082039164 | 0.72221883 | 0.99979869 | 1200.85836 | 1271.1241 |
| *YBR063C* | 0.064068668 | 0.72222352 | 0.99979869 | 4160.56937 | 4349.49965 |
| *EAP1* | 0.05743586 | 0.72241278 | 0.99979869 | 16255.9737 | 16916.2015 |
| *NAM9* | 0.058034732 | 0.72307883 | 0.99979869 | 12363.6709 | 12871.1566 |
| *SMD3* | 0.064130376 | 0.7232705 | 0.99979869 | 4004.48705 | 4186.50873 |
| *ARP5* | 0.056662447 | 0.72357292 | 0.99979869 | 20647.74 | 21474.8216 |
| *YET1* | 0.056395126 | 0.72389923 | 0.99979869 | 22995.8035 | 23912.5096 |
| *UGO1* | 0.061172239 | 0.72393265 | 0.99979869 | 5863.8177 | 6117.79729 |
| *DAL1* | 0.080704366 | 0.72513586 | 0.99979869 | 1223.29519 | 1293.6763 |
| *ASI3* | 0.058394316 | 0.7252052 | 0.99979869 | 9686.85905 | 10086.9848 |
| *RPH1* | 0.058072532 | 0.72544277 | 0.99979869 | 10401.9112 | 10829.1573 |
| *YAP1801* | 0.056575189 | 0.72547607 | 0.99979869 | 17500.7302 | 18200.652 |
| *SUP35* | 0.054762903 | 0.72573362 | 0.99979869 | 79698.5596 | 82781.9571 |
| *YOR29-23* | 0.115924118 | 0.72585123 | 0.99979869 | 405.814035 | 439.767935 |
| *RRT1* | 0.117000522 | 0.72587851 | 0.99979869 | 396.058889 | 429.516934 |

| | | | | | |
|---|---|---|---|---|---|
| *UTR5* | 0.187007704 | 0.72607158 | 0.99979869 | 128.767915 | 146.589312 |
| *MST28* | 0.063742387 | 0.72657659 | 0.99979869 | 3805.48209 | 3977.38832 |
| *GTT3* | 0.057841542 | 0.7268753 | 0.99979869 | 10123.8895 | 10538.0288 |
| *AIM21* | 0.056582394 | 0.72739999 | 0.99979869 | 14575.1622 | 15158.1549 |
| *PRE10* | 0.054710894 | 0.72744234 | 0.99979869 | 48480.1444 | 50353.9411 |
| *URN1* | 0.0602558 | 0.72762628 | 0.99979869 | 5890.15659 | 6141.37459 |
| *PZF1* | 0.066508176 | 0.72777439 | 0.99979869 | 2807.53075 | 2939.98703 |
| *tS(AGA)G* | 0.405951136 | 0.72869525 | 0.99979869 | 22.4368336 | 29.7279024 |
| *GIP1* | 0.084278321 | 0.72939295 | 0.99979869 | 989.171709 | 1048.67738 |
| *ARO80* | 0.059443532 | 0.72976861 | 0.99979869 | 6197.44366 | 6458.13052 |
| *YBR032W* | 0.145530787 | 0.72979876 | 0.99979869 | 222.417307 | 246.02402 |
| *PRE8* | 0.054536652 | 0.73034211 | 0.99979869 | 30824.3074 | 32011.8254 |
| *CLC1* | 0.054365248 | 0.73036113 | 0.99979869 | 35549.6996 | 36914.879 |
| *BPL1* | 0.056382871 | 0.73046458 | 0.99979869 | 12235.8785 | 12723.5422 |
| *SBH1* | 0.055175106 | 0.7305117 | 0.99979869 | 19991.2188 | 20770.5779 |
| *DAD4* | 0.057548605 | 0.73051915 | 0.99979869 | 8800.11636 | 9158.24413 |
| *SOK1* | 0.055895421 | 0.73066468 | 0.99979869 | 14300.0671 | 14864.9763 |
| *YBT1* | 0.053878551 | 0.73098293 | 0.99979869 | 52304.1613 | 54294.4258 |
| *YJR085C* | 0.055197 | 0.73130001 | 0.99979869 | 18111.4023 | 18817.7622 |
| *RGD1* | 0.057397762 | 0.73178495 | 0.99979869 | 8506.48649 | 8851.73921 |
| *HEM4* | 0.061830904 | 0.73184526 | 0.99979869 | 4076.67512 | 4255.19044 |
| *RPL12A* | 0.053090049 | 0.73215643 | 0.99979869 | 182888.484 | 189743.975 |
| *ISR1* | 0.060545063 | 0.73223945 | 0.99979869 | 4758.55976 | 4962.5095 |
| *DAL80* | 0.235029631 | 0.73238877 | 0.99979869 | 73.163588 | 86.1084069 |
| *DMC1* | 0.085514069 | 0.73275952 | 0.99979869 | 901.375404 | 956.418376 |
| *FBP26* | 0.056695833 | 0.73299415 | 0.99979869 | 9441.02939 | 9819.43368 |
| *STS1* | 0.056160611 | 0.73337457 | 0.99979869 | 10678.9573 | 11102.859 |
| *HMO1* | 0.053674507 | 0.73350641 | 0.99979869 | 35610.1815 | 36959.9834 |
| *ECM34* | 0.075602327 | 0.73357346 | 0.99979869 | 1380.35303 | 1454.61702 |
| *SNU56* | 0.061422634 | 0.73368099 | 0.99979869 | 4052.28726 | 4228.53784 |
| *NTC20* | 0.063832933 | 0.7336845 | 0.99979869 | 3116.76885 | 3257.76806 |
| *IRC18* | 0.120578233 | 0.73384331 | 0.99979869 | 338.503534 | 368.010929 |
| *SUT2* | 0.059557084 | 0.73445738 | 0.99979869 | 5075.60198 | 5289.51642 |
| *SHO1* | 0.05560577 | 0.73467233 | 0.99979869 | 11553.0183 | 12006.9973 |
| *SPS100* | 0.086310006 | 0.73473933 | 0.99979869 | 852.599678 | 905.163372 |
| *SEC27* | 0.052929611 | 0.73475251 | 0.99979869 | 63435.7573 | 65806.2998 |
| *LRO1* | 0.054443979 | 0.73483391 | 0.99979869 | 17814.8459 | 18499.9812 |
| *FRT1* | 0.058728441 | 0.73563776 | 0.99979869 | 5521.4121 | 5750.81146 |
| *ACC1* | 0.052442661 | 0.73587497 | 0.99979869 | 106138.905 | 110068.071 |
| *LGE1* | 0.055447729 | 0.73618302 | 0.99979869 | 10912.1053 | 11339.6571 |
| *snR85* | 0.091764365 | 0.73646871 | 0.99979869 | 690.66427 | 736.021859 |
| *KRE6* | 0.053126941 | 0.73669341 | 0.99979869 | 31247.6807 | 32419.8152 |
| *LMO1* | 0.062088249 | 0.73673466 | 0.99979869 | 3439.66415 | 3590.92559 |
| *IMD3* | 0.052337599 | 0.73693794 | 0.99979869 | 79060.5731 | 81981.3539 |
| *AFB1* | 0.062020607 | 0.73726211 | 0.99979869 | 3415.27629 | 3565.29808 |

| | | | | | |
|---|---|---|---|---|---|
| *PRE6* | 0.052708481 | 0.73727402 | 0.99979869 | 41169.6387 | 42701.569 |
| *YMR252C* | 0.062211063 | 0.73741479 | 0.99979869 | 3333.33307 | 3480.21478 |
| *RFA1* | 0.055748465 | 0.73816205 | 0.99979869 | 8877.18201 | 9226.92584 |
| *AIM26* | 0.075996351 | 0.73825766 | 0.99979869 | 1258.41371 | 1326.47951 |
| *SPE2* | 0.053961745 | 0.73835289 | 0.99979869 | 15555.5543 | 16148.4016 |
| *COQ5* | 0.052902157 | 0.73843008 | 0.99979869 | 27679.2486 | 28713.0533 |
| *YGR066C* | 0.093489149 | 0.73877555 | 0.99979869 | 636.035458 | 678.616254 |
| *PKH2* | 0.054954642 | 0.73903832 | 0.99979869 | 10418.4949 | 10823.0067 |
| *PKH3* | 0.054884844 | 0.7390446 | 0.99979869 | 10630.1816 | 11042.3781 |
| *MMM1* | 0.057980745 | 0.73920353 | 0.99979869 | 5426.7872 | 5649.32655 |
| *NSP1* | 0.052488008 | 0.73924253 | 0.99979869 | 33553.797 | 34797.0223 |
| *ADK2* | 0.061937895 | 0.73956822 | 0.99979869 | 3238.70816 | 3380.78007 |
| *SWP1* | 0.052875092 | 0.73965587 | 0.99979869 | 23463.0749 | 24338.9512 |
| *YSF3* | 0.061687793 | 0.73984059 | 0.99979869 | 3300.16557 | 3444.33627 |
| *ECM25* | 0.058111756 | 0.73997251 | 0.99979869 | 5163.39828 | 5375.62483 |
| *ICP55* | 0.056219343 | 0.74007409 | 0.99979869 | 7208.07669 | 7494.5067 |
| *PDR12* | 0.05245409 | 0.74047426 | 0.99979869 | 27701.6854 | 28727.4047 |
| *MXR2* | 0.055902332 | 0.74059215 | 0.99979869 | 7515.36376 | 7812.28772 |
| *YGR164W* | 0.137254373 | 0.74087274 | 0.99979869 | 230.221423 | 253.19972 |
| *tT(AGU)C* | 0.55695955 | 0.74122315 | 0.99979869 | 9.75514506 | 14.3514011 |
| *ATG32* | 0.061724544 | 0.74123644 | 0.99979869 | 3168.47112 | 3306.97286 |
| *SLY41* | 0.054566617 | 0.74134868 | 0.99979869 | 10014.6319 | 10400.6654 |
| *VPS13* | 0.052339298 | 0.7414029 | 0.99979869 | 25986.7309 | 26946.8059 |
| *TOM40* | 0.051904102 | 0.74153961 | 0.99979869 | 36316.454 | 37646.8005 |
| *ALG8* | 0.053333493 | 0.7416011 | 0.99979869 | 14943.9067 | 15506.6889 |
| *COA6* | 0.057916618 | 0.74173377 | 0.99979869 | 4985.85464 | 5190.08171 |
| *YIL161W* | 0.057895176 | 0.74186572 | 0.99979869 | 4978.05052 | 5181.88091 |
| *PBA1* | 0.05566001 | 0.74194014 | 0.99979869 | 7401.22856 | 7692.35101 |
| *BUD13* | 0.062874057 | 0.74213106 | 0.99979869 | 2772.41223 | 2895.90773 |
| *YJL055W* | 0.051966584 | 0.7427317 | 0.99979869 | 27595.3543 | 28607.468 |
| *SEC13* | 0.051558712 | 0.74273525 | 0.99979869 | 39666.3708 | 41109.5886 |
| *ATG36* | 0.068379779 | 0.74287574 | 0.99979869 | 1789.0936 | 1875.93315 |
| *RGR1* | 0.05293286 | 0.74298775 | 0.99979869 | 15537.9951 | 16118.6737 |
| *ISC1* | 0.053113143 | 0.74313338 | 0.99979869 | 14227.8791 | 14761.4412 |
| *REG1* | 0.051727491 | 0.74313491 | 0.99979869 | 31128.6679 | 32265.0251 |
| *PEA2* | 0.055766998 | 0.74344665 | 0.99979869 | 6733.00112 | 6998.35826 |
| *CWC23* | 0.060531857 | 0.74539618 | 0.99979869 | 3211.39375 | 3349.00197 |
| *SUS1* | 0.056252365 | 0.74543523 | 0.99979869 | 5649.2045 | 5873.82347 |
| *TRR1* | 0.050336006 | 0.74586164 | 0.99979869 | 96661.7814 | 100093.847 |
| *PAA1* | 0.051383147 | 0.74612089 | 0.99979869 | 24691.2477 | 25586.498 |
| *FLC3* | 0.052391147 | 0.74616033 | 0.99979869 | 14508.8272 | 15045.3939 |
| *KAR5* | 0.063677509 | 0.74658181 | 0.99979869 | 2336.35724 | 2441.78839 |
| *NTA1* | 0.054608128 | 0.7468092 | 0.99979869 | 7278.31373 | 7559.088 |
| *UNG1* | 0.054997996 | 0.74691369 | 0.99979869 | 6678.37231 | 6937.87735 |
| *ATG21* | 0.054378988 | 0.74696556 | 0.99979869 | 7594.38043 | 7886.09493 |

| | | | | | |
|---|---|---|---|---|---|
| *COS111* | 0.058148523 | 0.74719256 | 0.99979869 | 4015.21771 | 4180.35813 |
| *CLB3* | 0.053994576 | 0.74785547 | 0.99979869 | 7913.37367 | 8215.15206 |
| *ARC40* | 0.050737919 | 0.74824784 | 0.99979869 | 28225.5367 | 29235.8543 |
| *WWM1* | 0.050606014 | 0.7485062 | 0.99979869 | 30013.6548 | 31085.1349 |
| *ORC5* | 0.055711393 | 0.7485741 | 0.99979869 | 5457.02815 | 5671.87875 |
| *YKU80* | 0.058639723 | 0.74862988 | 0.99979869 | 3619.15882 | 3769.293 |
| *MND1* | 0.106719434 | 0.74892022 | 0.99979869 | 395.083375 | 425.416534 |
| *BRE5* | 0.050206874 | 0.74943469 | 0.99979869 | 35972.0974 | 37245.9864 |
| *SWR1* | 0.051074543 | 0.74997927 | 0.99979869 | 18015.8019 | 18665.0223 |
| *RLF2* | 0.054211258 | 0.7502102 | 0.99979869 | 6703.73569 | 6960.42956 |
| *SPC25* | 0.054285289 | 0.75032789 | 0.99979869 | 6568.13917 | 6819.99084 |
| *RPT5* | 0.049784872 | 0.7505375 | 0.99979869 | 43691.3437 | 45225.3654 |
| *YHL015W-A* | 0.131902813 | 0.75138115 | 0.99979869 | 228.270394 | 250.12442 |
| *MSH3* | 0.053942121 | 0.75143559 | 0.99979869 | 6682.27437 | 6936.85225 |
| *PKC1* | 0.050213951 | 0.75150178 | 0.99979869 | 24676.6149 | 25550.6195 |
| *MRPL25* | 0.056830897 | 0.75161874 | 0.99979869 | 4138.13253 | 4304.39524 |
| *SIZ1* | 0.052993018 | 0.7517977 | 0.99979869 | 8099.69694 | 8402.74537 |
| *ECM12* | 0.122514056 | 0.75186543 | 0.99979869 | 271.193033 | 295.228824 |
| *LSM2* | 0.051616842 | 0.75217261 | 0.99979869 | 11742.2681 | 12169.9882 |
| *SCS7* | 0.049917663 | 0.75221228 | 0.99979869 | 27227.5854 | 28186.1518 |
| *YSC83* | 0.052612631 | 0.7523916 | 0.99979869 | 8611.84206 | 8931.69701 |
| *PLB2* | 0.049955274 | 0.75241153 | 0.99979869 | 25631.6436 | 26534.7156 |
| *IRC5* | 0.0518373 | 0.75342823 | 0.99979869 | 10001.9502 | 10367.8622 |
| *YHR213W* | 0.257086924 | 0.75371123 | 0.99979869 | 49.7512398 | 59.4558047 |
| *STE5* | 0.051864145 | 0.75419479 | 0.99979869 | 9439.07836 | 9784.58028 |
| *YLR412C-A* | 0.167390511 | 0.75452347 | 0.99979869 | 127.7924 | 143.514011 |
| *YMR105W-A* | 0.100352815 | 0.75472811 | 0.99979869 | 435.07947 | 466.420537 |
| *MET2* | 0.05446903 | 0.75488981 | 0.99979869 | 5227.78224 | 5428.93003 |
| *CLD1* | 0.064027434 | 0.75573421 | 0.99979869 | 1880.79197 | 1966.14196 |
| *SWE1* | 0.053327306 | 0.75588811 | 0.99979869 | 6146.7169 | 6378.17271 |
| *tE(UUC)P* | 0.486570269 | 0.75592141 | 0.99979869 | 11.7061741 | 16.4016013 |
| *HOS4* | 0.049522858 | 0.75615302 | 0.99979869 | 20040.97 | 20740.85 |
| *ZRT2* | 0.049732587 | 0.75620027 | 0.99979869 | 17760.2171 | 18383.1198 |
| *ATP4* | 0.049140784 | 0.75662108 | 0.99979869 | 23945.9546 | 24775.6439 |
| *RAM1* | 0.052637764 | 0.75685758 | 0.99979869 | 6748.60935 | 6999.38336 |
| *YKL070W* | 0.060754385 | 0.75707479 | 0.99979869 | 2359.76959 | 2461.2653 |
| *MCM6* | 0.049485572 | 0.75711672 | 0.99979869 | 18206.0272 | 18841.3395 |
| *ERP5* | 0.052256023 | 0.75761178 | 0.99979869 | 7056.87194 | 7317.16438 |
| *PIP2* | 0.052623618 | 0.7583909 | 0.99979869 | 6294.01959 | 6527.83732 |
| *TAH11* | 0.049236814 | 0.75901092 | 0.99979869 | 16732.0248 | 17312.9153 |
| *SIP1* | 0.050191607 | 0.75911761 | 0.99979869 | 11229.1475 | 11626.6851 |
| *THI2* | 0.080783874 | 0.75946329 | 0.99979869 | 757.974771 | 801.628264 |
| *QRI1* | 0.049763639 | 0.75987781 | 0.99979869 | 12312.9441 | 12745.0693 |
| *REX2* | 0.049526578 | 0.76023314 | 0.99979869 | 13083.6006 | 13540.547 |
| *snR52* | 0.235917966 | 0.76025575 | 0.99979869 | 56.5798414 | 66.6315053 |

| | | | | | |
|---|---|---|---|---|---|
| *YMR114C* | 0.052272104 | 0.76030199 | 0.99979869 | 6179.8844 | 6407.90061 |
| *YLL056C* | 0.061767649 | 0.76030736 | 0.99979869 | 2025.16811 | 2113.75637 |
| *EMC1* | 0.04770285 | 0.76061703 | 0.99979869 | 40140.4709 | 41489.9007 |
| *RPN12* | 0.048307698 | 0.76129036 | 0.99979869 | 21321.8206 | 22047.8526 |
| *TEM1* | 0.053260291 | 0.76132187 | 0.99979869 | 4961.46678 | 5148.05261 |
| *BIT2* | 0.058705707 | 0.761789 | 0.99979869 | 2535.3622 | 2640.65781 |
| *PAU7* | 0.096952574 | 0.76189182 | 0.99979869 | 438.981528 | 469.495837 |
| *RER1* | 0.049030001 | 0.76285917 | 0.99979869 | 12670.9579 | 13108.9798 |
| *VTS1* | 0.053989147 | 0.76287553 | 0.99979869 | 4197.63892 | 4357.70045 |
| *SBE2* | 0.049418507 | 0.7632284 | 0.99979869 | 10711.1493 | 11084.4072 |
| *KRE28* | 0.055317172 | 0.76403886 | 0.99979869 | 3404.54563 | 3537.62038 |
| *RFC3* | 0.048479488 | 0.76404027 | 0.99979869 | 14278.6058 | 14766.5667 |
| *ECM5* | 0.049505622 | 0.76411151 | 0.99979869 | 9787.33704 | 10129.0139 |
| *YDL109C* | 0.054601916 | 0.76438212 | 0.99979869 | 3679.64072 | 3821.5731 |
| *MAD2* | 0.052537009 | 0.7649667 | 0.99979869 | 4848.3071 | 5028.1159 |
| *YCR100C* | 0.065531047 | 0.76587213 | 0.99979869 | 1410.59398 | 1476.14412 |
| *SOL1* | 0.050812829 | 0.76594608 | 0.99979869 | 6363.28112 | 6591.39353 |
| *RAD33* | 0.057690841 | 0.76704301 | 0.99979869 | 2452.44347 | 2552.4992 |
| *SIP3* | 0.04862715 | 0.76718702 | 0.99979869 | 10405.8132 | 10762.5258 |
| *PMT7* | 0.04928211 | 0.76746057 | 0.99979869 | 8414.78813 | 8707.20009 |
| *SIL1* | 0.049940844 | 0.76775976 | 0.99979869 | 7027.6065 | 7275.13528 |
| *SPT20* | 0.049090799 | 0.76784464 | 0.99979869 | 8659.64227 | 8959.37471 |
| *ERG12* | 0.046913832 | 0.7688937 | 0.99979869 | 18394.3015 | 19002.2802 |
| *HEL2* | 0.047051837 | 0.76920764 | 0.99979869 | 16467.6604 | 17013.5861 |
| *NUP159* | 0.046121377 | 0.77000575 | 0.99979869 | 26766.167 | 27635.6731 |
| *APM2* | 0.04870045 | 0.77009909 | 0.99979869 | 8388.44924 | 8676.44709 |
| *NUP2* | 0.045649358 | 0.77066046 | 0.99979869 | 36762.2642 | 37944.0795 |
| *KAE1* | 0.046904758 | 0.77073621 | 0.99979869 | 15044.3847 | 15541.5423 |
| *HAL9* | 0.050104397 | 0.77142598 | 0.99979869 | 5691.15163 | 5892.27527 |
| *YIL012W* | 0.146008629 | 0.77146985 | 0.99979869 | 147.30269 | 162.990913 |
| *MKS1* | 0.04811097 | 0.77232033 | 0.99979869 | 8603.06243 | 8894.79341 |
| *SRN2* | 0.054519496 | 0.77268142 | 0.99979869 | 2912.88632 | 3025.07034 |
| *YNL143C* | 0.073172071 | 0.77278893 | 0.99979869 | 856.501736 | 901.062972 |
| *YOR029W* | 0.135660667 | 0.77289328 | 0.99979869 | 171.690553 | 188.618415 |
| *REB1* | 0.046308239 | 0.77307896 | 0.99979869 | 15704.808 | 16217.0833 |
| *KCS1* | 0.04617255 | 0.77372261 | 0.99979869 | 15677.4936 | 16187.3554 |
| *TDA11* | 0.051871204 | 0.7738449 | 0.99979869 | 3892.30288 | 4034.79392 |
| *PUT2* | 0.046184156 | 0.77445409 | 0.99979869 | 14452.2474 | 14922.3819 |
| *STL1* | 0.06762536 | 0.77462145 | 0.99979869 | 1082.8211 | 1134.78579 |
| *PUT1* | 0.058779062 | 0.77462282 | 0.99979869 | 1884.69403 | 1963.06666 |
| *AYT1* | 0.059968329 | 0.77499352 | 0.99979869 | 1711.05244 | 1783.67414 |
| *CAB3* | 0.046411946 | 0.77544464 | 0.99979869 | 11982.2447 | 12373.9831 |
| *AGE1* | 0.047808571 | 0.77560212 | 0.99979869 | 7656.81336 | 7914.79773 |
| *MIP1* | 0.045861208 | 0.77630839 | 0.99979869 | 13940.1023 | 14390.3549 |
| *UTP22* | 0.04496817 | 0.77633269 | 0.99979869 | 23047.5057 | 23777.1964 |

| | | | | | |
|---|---|---|---|---|---|
| *PRE2* | 0.044170897 | 0.77638507 | 0.99979869 | 53280.6513 | 54937.1636 |
| *THP3* | 0.045977375 | 0.77641749 | 0.99979869 | 13113.8415 | 13538.4968 |
| *SRS2* | 0.046479655 | 0.77709218 | 0.99979869 | 10257.535 | 10593.3842 |
| *YGL007C-A* | 0.178445962 | 0.77715599 | 0.99979869 | 88.7718201 | 100.459808 |
| *ARR3* | 0.062408398 | 0.77716938 | 0.99979869 | 1393.03471 | 1454.61702 |
| *RPS10B* | 0.043616295 | 0.77759657 | 0.99979869 | 106347.665 | 109611.902 |
| *PBS2* | 0.045192028 | 0.77825718 | 0.99979869 | 15692.1263 | 16191.4558 |
| *MAD3* | 0.048899818 | 0.77866282 | 0.99979869 | 5202.41886 | 5381.77543 |
| *UBX3* | 0.048085996 | 0.77884946 | 0.99979869 | 6052.092 | 6257.2109 |
| *YCR101C* | 0.073341154 | 0.77899362 | 0.99979869 | 776.509547 | 817.004765 |
| *MCD4* | 0.045923435 | 0.77919844 | 0.99979869 | 10591.161 | 10933.7175 |
| *ERI1* | 0.056401264 | 0.77940178 | 0.99979869 | 2057.36009 | 2139.38387 |
| *NDE2* | 0.068570455 | 0.77971845 | 0.99979869 | 952.102158 | 998.44748 |
| *MRPL50* | 0.046968433 | 0.7803544 | 0.99979869 | 7223.68492 | 7462.7286 |
| *tL(UAA)L* | 0.376387175 | 0.78048112 | 0.99979869 | 16.5837466 | 21.5271017 |
| *GPX2* | 0.044054074 | 0.78116845 | 0.99979869 | 21253.5345 | 21912.5393 |
| *SED4* | 0.044071266 | 0.78128844 | 0.99979869 | 20645.789 | 21286.2032 |
| *VPH2* | 0.047866072 | 0.78131642 | 0.99979869 | 5636.52282 | 5826.66886 |
| *USB1* | 0.053799675 | 0.78161175 | 0.99979869 | 2475.85582 | 2569.92591 |
| *RAD61* | 0.050986711 | 0.78241341 | 0.99979869 | 3311.87175 | 3431.00997 |
| *YOR365C* | 0.05623217 | 0.78258045 | 0.99979869 | 1942.24938 | 2019.44716 |
| *ERG9* | 0.042835199 | 0.78306313 | 0.99979869 | 48495.7526 | 49957.2274 |
| *CMR2* | 0.045748447 | 0.78333873 | 0.99979869 | 8371.86549 | 8641.59369 |
| *YIL163C* | 0.08729729 | 0.78382826 | 0.99979869 | 443.8591 | 471.546038 |
| *GSH2* | 0.043250537 | 0.78384047 | 0.99979869 | 25429.7121 | 26203.6083 |
| *SPO73* | 0.0631594 | 0.78390921 | 0.99979869 | 1180.37255 | 1233.1954 |
| *MCM2* | 0.044968658 | 0.78395328 | 0.99979869 | 10267.2902 | 10592.3591 |
| *ATG23* | 0.050592878 | 0.78395747 | 0.99979869 | 3317.72484 | 3436.13547 |
| *RPL33A* | 0.04211761 | 0.78414464 | 0.99979869 | 205877.459 | 211976.346 |
| *COM2* | 0.043015535 | 0.7844006 | 0.99979869 | 28163.1038 | 29015.4578 |
| *KAP122* | 0.044692998 | 0.78456998 | 0.99979869 | 10797.9701 | 11137.7124 |
| *YGL015C* | 0.140477801 | 0.78499018 | 0.99979869 | 139.498574 | 153.765012 |
| *YMR082C* | 0.26947114 | 0.78515215 | 0.99979869 | 33.1674932 | 39.9789032 |
| *PFK2* | 0.041963506 | 0.78521917 | 0.99979869 | 142779.23 | 146993.201 |
| *IMP1* | 0.046450527 | 0.78525364 | 0.99979869 | 6283.28893 | 6488.88352 |
| *SAC3* | 0.044101647 | 0.78528634 | 0.99979869 | 12846.5505 | 13245.3182 |
| *GWT1* | 0.046068506 | 0.78542917 | 0.99979869 | 6803.23817 | 7023.98576 |
| *LTP1* | 0.043925489 | 0.78551861 | 0.99979869 | 13588.9171 | 14009.0177 |
| *WIP1* | 0.051900937 | 0.78615643 | 0.99979869 | 2670.95872 | 2768.79532 |
| *PEF1* | 0.047079007 | 0.78665965 | 0.99979869 | 5166.32482 | 5337.69613 |
| *PFY1* | 0.041828566 | 0.78676092 | 0.99979869 | 77459.7538 | 79738.435 |
| *HDA3* | 0.045753307 | 0.78720193 | 0.99979869 | 6665.69062 | 6880.47175 |
| *YMR141C* | 0.064636855 | 0.78751618 | 0.99979869 | 1023.31472 | 1070.20449 |
| *ALG7* | 0.043951696 | 0.78760101 | 0.99979869 | 11128.6695 | 11472.9201 |
| *YJL133C-A* | 0.049705962 | 0.78760568 | 0.99979869 | 3313.82278 | 3429.98487 |

| | | | | | |
|---|---|---|---|---|---|
| *RAD5* | 0.046307137 | 0.7881243 | 0.99979869 | 5628.7187 | 5812.31746 |
| *YHR007C-A* | 0.083502602 | 0.78883909 | 0.99979869 | 468.246963 | 496.14844 |
| *MBA1* | 0.046852603 | 0.78919011 | 0.99979869 | 4831.72335 | 4991.2123 |
| *PCL9* | 0.044805206 | 0.78972111 | 0.99979869 | 7334.89357 | 7566.2637 |
| *RRM3* | 0.04542724 | 0.79001066 | 0.99979869 | 6202.32123 | 6400.72491 |
| *SNU66* | 0.045219786 | 0.79025413 | 0.99979869 | 6423.76302 | 6628.29713 |
| *YCR041W* | 0.101483339 | 0.79069577 | 0.99979869 | 278.997149 | 299.329224 |
| *SRB7* | 0.045924545 | 0.790762 | 0.99979869 | 5378.98699 | 5552.96714 |
| *TCM62* | 0.044442461 | 0.79144756 | 0.99979869 | 7290.0199 | 7518.084 |
| *SKS1* | 0.044632287 | 0.79164135 | 0.99979869 | 6865.67109 | 7081.39137 |
| *CLN1* | 0.045127343 | 0.79207726 | 0.99979869 | 5971.12429 | 6160.85149 |
| *PKP2* | 0.045984583 | 0.79211712 | 0.99979869 | 5002.43839 | 5164.45421 |
| *STE14* | 0.045460097 | 0.79229959 | 0.99979869 | 5501.90181 | 5678.02935 |
| *SEI1* | 0.050452349 | 0.79247638 | 0.99979869 | 2623.15851 | 2716.51522 |
| *RBL2* | 0.04441565 | 0.79261117 | 0.99979869 | 6863.72006 | 7078.31606 |
| *ELP4* | 0.042325228 | 0.79314344 | 0.99979869 | 13338.2098 | 13735.316 |
| *YPR027C* | 0.053617127 | 0.79326639 | 0.99979869 | 1897.37571 | 1969.21726 |
| *ALF1* | 0.04526475 | 0.79352001 | 0.99979869 | 5407.27691 | 5579.61975 |
| *MDM12* | 0.049115591 | 0.79492518 | 0.99979869 | 2847.52684 | 2946.13764 |
| *BRL1* | 0.046964585 | 0.79520733 | 0.99979869 | 3755.73085 | 3880.00381 |
| *CSG2* | 0.040407204 | 0.79544445 | 0.99979869 | 39356.1572 | 40474.0265 |
| *STP1* | 0.041523277 | 0.79569152 | 0.99979869 | 15165.3485 | 15608.1738 |
| *PCT1* | 0.041665621 | 0.79662539 | 0.99979869 | 12808.5055 | 13183.8122 |
| *SUR2* | 0.040337375 | 0.79740237 | 0.99979869 | 27000.2905 | 27765.8608 |
| *RPN3* | 0.039926769 | 0.79769383 | 0.99979869 | 40159.9812 | 41286.9309 |
| *SAP155* | 0.040435816 | 0.7977407 | 0.99979869 | 23287.4823 | 23949.4132 |
| *ATG4* | 0.045391434 | 0.79831498 | 0.99979869 | 4292.26383 | 4429.45745 |
| *SYG1* | 0.039969439 | 0.79831629 | 0.99979869 | 32414.396 | 33324.9786 |
| *MPP6* | 0.043484599 | 0.79835358 | 0.99979869 | 6283.28893 | 6475.55722 |
| *PET494* | 0.043394889 | 0.79837009 | 0.99979869 | 6414.98339 | 6610.87043 |
| *YPK1* | 0.040130159 | 0.79837813 | 0.99979869 | 27095.8909 | 27860.17 |
| *GIN4* | 0.04168535 | 0.79864966 | 0.99979869 | 10577.5038 | 10887.588 |
| *ZIP2* | 0.095376842 | 0.79914916 | 0.99979869 | 292.654352 | 312.655525 |
| *UBP3* | 0.040058357 | 0.79934905 | 0.99979869 | 24096.1838 | 24774.6188 |
| *MSO1* | 0.043467543 | 0.80004984 | 0.99979869 | 5773.09485 | 5949.68087 |
| *tK(CUU)E1* | 0.211709015 | 0.80044363 | 0.99979869 | 47.8002108 | 55.3554044 |
| *WTM2* | 0.041498552 | 0.80143172 | 0.99979869 | 9091.7952 | 9357.11355 |
| *SAM35* | 0.04004514 | 0.80204561 | 0.99979869 | 16313.5291 | 16772.6875 |
| *TRA1* | 0.039237056 | 0.80254163 | 0.99979869 | 27808.992 | 28575.6899 |
| *CDC28* | 0.04170598 | 0.80293003 | 0.99979869 | 7676.32365 | 7901.47143 |
| *UIP3* | 0.039813022 | 0.80331431 | 0.99979869 | 15971.1235 | 16418.0029 |
| *DRS2* | 0.039426811 | 0.80334845 | 0.99979869 | 20479.9515 | 21047.3549 |
| *YOL107W* | 0.047136993 | 0.80349061 | 0.99979869 | 2792.89803 | 2885.65673 |
| *PAU17* | 0.071529911 | 0.80354499 | 0.99979869 | 583.357675 | 613.009849 |
| *COA1* | 0.040800379 | 0.80370951 | 0.99979869 | 9766.85124 | 10047.0059 |

| | | | | | |
|---|---|---|---|---|---|
| *COQ11* | 0.043229353 | 0.80441129 | 0.99979869 | 4921.47068 | 5071.1701 |
| *EXG2* | 0.039340148 | 0.80445444 | 0.99979869 | 18480.1468 | 18991.0041 |
| *YER135C* | 0.10127753 | 0.80548448 | 0.99979869 | 234.123481 | 251.14952 |
| *VPS64* | 0.040979277 | 0.80567803 | 0.99979869 | 7961.17388 | 8190.54965 |
| *MVP1* | 0.039930663 | 0.80594446 | 0.99979869 | 11392.0584 | 11711.7684 |
| *GET3* | 0.037990277 | 0.80595006 | 0.99979869 | 60620.4224 | 62237.9264 |
| *PDS5* | 0.041537669 | 0.80617887 | 0.99979869 | 6590.576 | 6783.08724 |
| *GAP1* | 0.046998159 | 0.80638589 | 0.99979869 | 2603.64822 | 2689.86261 |
| *MAM3* | 0.038802968 | 0.80641033 | 0.99979869 | 20224.3667 | 20775.7034 |
| *YML054C-A* | 0.082557037 | 0.8064706 | 0.99979869 | 381.426172 | 403.889432 |
| *RRN11* | 0.045427257 | 0.8071138 | 0.99979869 | 3101.16061 | 3200.36246 |
| *CMR1* | 0.045450614 | 0.80752764 | 0.99979869 | 3049.45835 | 3147.05725 |
| *VMA7* | 0.038014303 | 0.80764118 | 0.99979869 | 33432.8332 | 34325.4762 |
| *MRX1* | 0.041401929 | 0.80818792 | 0.99979869 | 6089.16155 | 6266.4368 |
| *YOL019W* | 0.040402788 | 0.80831926 | 0.99979869 | 7952.39425 | 8178.24845 |
| *DUN1* | 0.044359532 | 0.80884348 | 0.99979869 | 3398.69254 | 3504.81718 |
| *COG6* | 0.039738841 | 0.80891046 | 0.99979869 | 9533.70327 | 9799.95678 |
| *PRM8* | 0.040453313 | 0.8091394 | 0.99979869 | 7415.86128 | 7626.74461 |
| *SLZ1* | 0.063916683 | 0.80924641 | 0.99979869 | 741.391025 | 774.975662 |
| *RRT5* | 0.106979103 | 0.80943886 | 0.99979869 | 196.078416 | 211.170617 |
| *RPL16B* | 0.036891981 | 0.80962137 | 0.99979869 | 261363.749 | 268133.378 |
| *SKM1* | 0.039676712 | 0.80977062 | 0.99979869 | 9113.25652 | 9367.36455 |
| *APC4* | 0.043493282 | 0.80979388 | 0.99979869 | 3750.85328 | 3865.65241 |
| *DAD2* | 0.0441468 | 0.81038044 | 0.99979869 | 3320.65138 | 3423.83427 |
| *SAT4* | 0.038000092 | 0.81049217 | 0.99979869 | 19490.7798 | 20010.9787 |
| *ECM27* | 0.03986751 | 0.81059552 | 0.99979869 | 8044.09262 | 8269.48236 |
| *MCA1* | 0.037313395 | 0.81067903 | 0.99979869 | 36876.3994 | 37842.5946 |
| *LST4* | 0.03814438 | 0.81078407 | 0.99979869 | 16930.0543 | 17383.6472 |
| *DDI1* | 0.038816362 | 0.81094096 | 0.99979869 | 11617.4023 | 11934.2152 |
| *SNF5* | 0.039179961 | 0.81114435 | 0.99979869 | 9810.74939 | 10080.8342 |
| *YPR174C* | 0.044236505 | 0.81129553 | 0.99979869 | 3175.29972 | 3274.16966 |
| *YKR023W* | 0.039954488 | 0.81174082 | 0.99979869 | 7254.90138 | 7458.6282 |
| *MVD1* | 0.0369952 | 0.81250753 | 0.99979869 | 34189.8324 | 35077.8997 |
| *tH(GUG)H* | 0.351640879 | 0.81254669 | 0.99979869 | 13.6572031 | 17.4267014 |
| *MDM30* | 0.040986847 | 0.81263707 | 0.99979869 | 5308.74994 | 5461.73324 |
| *HOG1* | 0.036883449 | 0.81266767 | 0.99979869 | 38027.5065 | 39012.2338 |
| *SGT2* | 0.036420547 | 0.81283487 | 0.99979869 | 96569.1075 | 99037.9942 |
| *YDR541C* | 0.04215296 | 0.81316587 | 0.99979869 | 4113.74467 | 4235.71354 |
| *CAF120* | 0.040418527 | 0.81343261 | 0.99979869 | 5807.23785 | 5972.23308 |
| *HXT3* | 0.036106602 | 0.81346212 | 0.99979869 | 290549.192 | 297912.536 |
| *PSR2* | 0.037902844 | 0.81372271 | 0.99979869 | 13591.8436 | 13953.6623 |
| *DON1* | 0.047914788 | 0.81393413 | 0.99979869 | 1922.73909 | 1987.66906 |
| *YHR138C* | 0.038841756 | 0.81395053 | 0.99979869 | 8883.03509 | 9125.44093 |
| *HXT10* | 0.124382375 | 0.81452028 | 0.99979869 | 130.718944 | 142.488911 |
| *YCL049C* | 0.0390437 | 0.81456528 | 0.99979869 | 7929.95742 | 8147.49545 |

| | | | | | |
|---|---|---|---|---|---|
| *GRS2* | 0.039621832 | 0.81476031 | 0.99979869 | 6600.33115 | 6784.11234 |
| *HRD1* | 0.038652719 | 0.81477019 | 0.99979869 | 8919.12913 | 9161.31943 |
| *VPS38* | 0.039868308 | 0.81549595 | 0.99979869 | 5932.10371 | 6098.32039 |
| *ITR2* | 0.036672713 | 0.81566404 | 0.99979869 | 23987.9017 | 24605.4773 |
| *DFG10* | 0.04599466 | 0.81580461 | 0.99979869 | 2224.17307 | 2296.22418 |
| *RPS26A* | 0.035623981 | 0.81594476 | 0.99979869 | 244941.937 | 251065.462 |
| *YOR296W* | 0.03924243 | 0.81688908 | 0.99979869 | 6435.4692 | 6612.92063 |
| *YIL152W* | 0.041414398 | 0.81808485 | 0.99979869 | 3824.99238 | 3936.38431 |
| *YKL183C-A* | 0.085837936 | 0.81854141 | 0.99979869 | 293.629866 | 311.630425 |
| *ADE17* | 0.035238952 | 0.81905631 | 0.99979869 | 75359.4711 | 77222.8394 |
| *ULS1* | 0.035945574 | 0.82066769 | 0.99979869 | 18578.6738 | 19047.3846 |
| *YDR246W-A* | 0.07152997 | 0.82083605 | 0.99979869 | 450.687702 | 473.596238 |
| *COA3* | 0.037039038 | 0.82102851 | 0.99979869 | 9869.28026 | 10125.9386 |
| *RGL1* | 0.036736112 | 0.82258856 | 0.99979869 | 9743.43889 | 9994.7258 |
| *RTN1* | 0.034451525 | 0.82319514 | 0.99979869 | 65936.001 | 67529.493 |
| *POP2* | 0.035809172 | 0.82328664 | 0.99979869 | 14200.5647 | 14557.4463 |
| *MCM16* | 0.044148366 | 0.8236735 | 0.99979869 | 2179.29941 | 2247.01938 |
| *YOR111W* | 0.040744414 | 0.82375429 | 0.99979869 | 3419.17834 | 3517.11838 |
| *TFC3* | 0.036041575 | 0.82416746 | 0.99979869 | 11401.8135 | 11690.2413 |
| *SNA4* | 0.037848465 | 0.82457141 | 0.99979869 | 5918.44651 | 6075.76818 |
| *NRM1* | 0.041273691 | 0.8245889 | 0.99979869 | 3055.31143 | 3143.98195 |
| *YCR050C* | 0.044471928 | 0.82462616 | 0.99979869 | 2047.60495 | 2111.70617 |
| *SWM1* | 0.038420915 | 0.82483704 | 0.99979869 | 5073.65095 | 5210.58372 |
| *RPD3* | 0.035142979 | 0.82507575 | 0.99979869 | 17116.3775 | 17538.4373 |
| *HSC82* | 0.033750579 | 0.82509462 | 0.99979869 | 383576.206 | 392655.36 |
| *YGR161C-D* | 0.043490626 | 0.82543874 | 0.99979869 | 2230.02616 | 2298.27438 |
| *SPE3* | 0.034099179 | 0.82565486 | 0.99979869 | 47802.1618 | 48945.4536 |
| *DPP1* | 0.035721321 | 0.82597922 | 0.99979869 | 11062.3345 | 11339.6571 |
| *GEP4* | 0.038949731 | 0.82624906 | 0.99979869 | 4237.63501 | 4353.60005 |
| *SPO19* | 0.28303674 | 0.82722725 | 0.99979869 | 18.5347756 | 22.5522018 |
| *MLH1* | 0.036921355 | 0.82761023 | 0.99979869 | 6337.91775 | 6502.20982 |
| *THI12* | 0.109005123 | 0.82789693 | 0.99979869 | 148.278205 | 159.915613 |
| *URA10* | 0.04718403 | 0.82790115 | 0.99979869 | 1457.41867 | 1505.87202 |
| *PEX6* | 0.0351715 | 0.82827032 | 0.99979869 | 11406.6911 | 11688.1911 |
| *MBP1* | 0.035544333 | 0.82862997 | 0.99979869 | 9345.42897 | 9578.53516 |
| *PPT1* | 0.034439046 | 0.82880258 | 0.99979869 | 16295.9698 | 16689.6544 |
| *PSA1* | 0.033065513 | 0.82881394 | 0.99979869 | 203266.982 | 207979.48 |
| *COQ10* | 0.039039529 | 0.82882813 | 0.99979869 | 3723.53887 | 3825.67351 |
| *HIS2* | 0.035023434 | 0.82928265 | 0.99979869 | 11048.6773 | 11320.1802 |
| *ISM1* | 0.036400744 | 0.82931127 | 0.99979869 | 6610.08629 | 6778.98684 |
| *ZDS1* | 0.034567472 | 0.8298454 | 0.99979869 | 13150.9111 | 13469.8151 |
| *RNR1* | 0.033637292 | 0.82989012 | 0.99979869 | 26868.596 | 27502.4101 |
| *NGL2* | 0.035207367 | 0.82993076 | 0.99979869 | 9573.69936 | 9810.20778 |
| *GCD1* | 0.033419449 | 0.83039361 | 0.99979869 | 30833.087 | 31555.6558 |
| *STT3* | 0.032854307 | 0.83085357 | 0.99979869 | 77298.7939 | 79079.2956 |

| | | | | | |
|---|---|---|---|---|---|
| *ADY2* | 0.060088959 | 0.83094654 | 0.99979869 | 617.500682 | 643.762851 |
| *RPS15* | 0.032367076 | 0.83212507 | 0.99979869 | 296511.536 | 303238.956 |
| *HXT8* | 0.042467291 | 0.83219949 | 0.99979869 | 2054.43355 | 2115.80657 |
| *SIA1* | 0.035467852 | 0.83225255 | 0.99979869 | 7231.48903 | 7411.47359 |
| *AVT1* | 0.033256551 | 0.83226416 | 0.99979869 | 24033.7509 | 24594.2012 |
| *PMI40* | 0.03291961 | 0.83229673 | 0.99979869 | 36091.1102 | 36924.1049 |
| *YIL134C-A* | 0.139434922 | 0.8324555 | 0.99979869 | 80.967704 | 89.1837071 |
| *SHR5* | 0.035589636 | 0.83362856 | 0.99979869 | 6343.77083 | 6502.20982 |
| *RPL12B* | 0.032023915 | 0.83402238 | 0.99979869 | 208163.089 | 212835.379 |
| *LHS1* | 0.032759319 | 0.83567099 | 0.99979869 | 19903.4225 | 20360.5378 |
| *FUN26* | 0.032950209 | 0.8357115 | 0.99979869 | 16969.0748 | 17361.095 |
| *ENT1* | 0.03259108 | 0.83642184 | 0.99979869 | 20148.2766 | 20608.612 |
| *YGR122W* | 0.034881603 | 0.83648907 | 0.99979869 | 6494.97558 | 6653.92463 |
| *YRR1* | 0.034316917 | 0.83657714 | 0.99979869 | 7776.80164 | 7964.00254 |
| *SSN3* | 0.035747037 | 0.83659255 | 0.99979869 | 5097.06329 | 5224.93512 |
| *SHE3* | 0.033254818 | 0.83660227 | 0.99979869 | 12407.569 | 12696.8896 |
| *TAF1* | 0.03339349 | 0.83663447 | 0.99979869 | 11479.8547 | 11748.672 |
| *STB5* | 0.035302795 | 0.83784293 | 0.99979869 | 5310.70097 | 5442.25633 |
| *MCM1* | 0.032016537 | 0.83802407 | 0.99979869 | 25744.8033 | 26322.5199 |
| *PPH21* | 0.032231521 | 0.83834714 | 0.99979869 | 19530.7759 | 19972.0249 |
| *POL32* | 0.035340294 | 0.83860177 | 0.99979869 | 5040.48345 | 5165.47931 |
| *AME1* | 0.03593001 | 0.83875837 | 0.99979869 | 4362.50087 | 4472.51166 |
| *UBX2* | 0.03244647 | 0.83902179 | 0.99979869 | 14928.2985 | 15267.8406 |
| *RIB1* | 0.032716572 | 0.83906308 | 0.99979869 | 12568.5289 | 12856.8052 |
| *RPA190* | 0.031096955 | 0.83971987 | 0.99979869 | 73030.918 | 74622.1605 |
| *ORC4* | 0.034560792 | 0.83974443 | 0.99979869 | 5783.82551 | 5924.05337 |
| *YOR015W* | 0.035583748 | 0.8397478 | 0.99979869 | 4479.56261 | 4591.42327 |
| *MGR3* | 0.033696289 | 0.83985111 | 0.99979869 | 7537.80059 | 7715.92832 |
| *SEM1* | 0.032116139 | 0.83985869 | 0.99979869 | 16687.1511 | 17062.7909 |
| *HOP2* | 0.052045647 | 0.84004557 | 0.99979869 | 799.921895 | 829.305966 |
| *NEW1* | 0.031018824 | 0.84033863 | 0.99979869 | 63802.5508 | 65189.1895 |
| *RPN1* | 0.030916362 | 0.84048525 | 0.99979869 | 80090.7165 | 81825.5387 |
| *SPO71* | 0.034608079 | 0.84101827 | 0.99979869 | 5289.23965 | 5417.65393 |
| *MMS4* | 0.036510119 | 0.84187881 | 0.99979869 | 3376.25571 | 3462.78808 |
| *GRX5* | 0.03227281 | 0.84193467 | 0.99979869 | 11519.8508 | 11780.4501 |
| *AIM25* | 0.033763671 | 0.84220323 | 0.99979869 | 6229.63564 | 6377.14761 |
| *BUD32* | 0.032497472 | 0.84261579 | 0.99979869 | 9611.74443 | 9830.70978 |
| *MRP20* | 0.031908293 | 0.84290803 | 0.99979869 | 12603.6474 | 12885.508 |
| *YKL106C-A* | 0.209035556 | 0.84297323 | 0.99979869 | 29.2654352 | 33.8283027 |
| *SWS2* | 0.032344111 | 0.84349864 | 0.99979869 | 9465.41725 | 9680.02007 |
| *CLP1* | 0.033488609 | 0.84365307 | 0.99979869 | 6147.69242 | 6292.0643 |
| *SLD3* | 0.034321391 | 0.84376142 | 0.99979869 | 4865.86636 | 4983.0115 |
| *MCM7* | 0.031464343 | 0.84383178 | 0.99979869 | 14780.9958 | 15106.8999 |
| *CCE1* | 0.035633073 | 0.8440461 | 0.99979869 | 3612.33022 | 3702.6615 |
| *YPR170W-B* | 0.030776622 | 0.84445106 | 0.99979869 | 23886.4482 | 24401.4823 |

| | | | | | |
|---|---|---|---|---|---|
| *AAR2* | 0.038552429 | 0.84457222 | 0.99979869 | 2244.65888 | 2305.45008 |
| *YPR003C* | 0.0332795 | 0.84468527 | 0.99979869 | 6114.52492 | 6257.2109 |
| *UFE1* | 0.034320826 | 0.84480328 | 0.99979869 | 4597.59987 | 4708.28468 |
| *SAM50* | 0.03165169 | 0.84553337 | 0.99979869 | 10734.5616 | 10972.6713 |
| *YDL211C* | 0.04653544 | 0.84556847 | 0.99979869 | 1022.3392 | 1055.85308 |
| *UBC12* | 0.035013585 | 0.84612241 | 0.99979869 | 3707.93064 | 3799.0209 |
| *NUD1* | 0.033081696 | 0.84632311 | 0.99979869 | 5823.8216 | 5958.90678 |
| *PUP1* | 0.030228422 | 0.84638746 | 0.99979869 | 28793.2862 | 29402.9456 |
| *BYE1* | 0.031800705 | 0.84640551 | 0.99979869 | 9158.13018 | 9362.23905 |
| *YBR201C-A* | 0.044562758 | 0.84665773 | 0.99979869 | 1158.91123 | 1195.2667 |
| *RKM1* | 0.032346453 | 0.84680131 | 0.99979869 | 7135.88861 | 7297.68748 |
| *tD(GUC)M* | 0.334567951 | 0.84733395 | 0.99979869 | 9.75514506 | 12.301201 |
| *YGR026W* | 0.030150024 | 0.84755699 | 0.99979869 | 23771.3375 | 24273.3448 |
| *CSR1* | 0.030351061 | 0.84810816 | 0.99979869 | 17610.9634 | 17985.3809 |
| *SNO3* | 0.061591748 | 0.84848143 | 0.99979869 | 426.299839 | 444.893435 |
| *MGL2* | 0.031526986 | 0.84851535 | 0.99979869 | 8489.90275 | 8677.47219 |
| *tH(GUG)M* | 0.175868316 | 0.84915228 | 0.99979869 | 39.0205802 | 44.0793035 |
| *BTT1* | 0.045108642 | 0.85025469 | 0.99979869 | 1021.36369 | 1053.80288 |
| *tL(CAA)N* | 0.166688771 | 0.8505785 | 0.99979869 | 42.9226383 | 48.1797038 |
| *GEA1* | 0.030518592 | 0.8507956 | 0.99979869 | 10894.546 | 11127.4614 |
| *RER2* | 0.03042451 | 0.85107022 | 0.99979869 | 11113.0613 | 11349.9081 |
| *PRS5* | 0.029123878 | 0.85109962 | 0.99979869 | 36406.2014 | 37148.6019 |
| *PRP19* | 0.030386391 | 0.85109986 | 0.99979869 | 11311.0907 | 11551.8528 |
| *MET4* | 0.030149604 | 0.85129535 | 0.99979869 | 12696.3213 | 12964.4407 |
| *SUP4* | 0.312541579 | 0.85135499 | 0.99979869 | 10.7306596 | 13.3263011 |
| *TRR4* | 0.312541579 | 0.85135499 | 0.99979869 | 10.7306596 | 13.3263011 |
| *SNF7* | 0.02939545 | 0.85204844 | 0.99979869 | 20165.8359 | 20580.9343 |
| *SNF6* | 0.029380734 | 0.8522885 | 0.99979869 | 19515.1677 | 19916.6695 |
| *PCI8* | 0.033686379 | 0.85234508 | 0.99979869 | 3617.20779 | 3702.6615 |
| *EST2* | 0.036146871 | 0.85294263 | 0.99979869 | 2335.38173 | 2394.63379 |
| *YML002W* | 0.032055438 | 0.85318787 | 0.99979869 | 5059.99374 | 5173.68011 |
| *RGI2* | 0.133815333 | 0.8532626 | 0.99979869 | 66.3349864 | 72.7821058 |
| *GAL10* | 0.033986988 | 0.85330397 | 0.99979869 | 3266.02257 | 3343.87647 |
| *MNS1* | 0.03056578 | 0.85351825 | 0.99979869 | 8119.20723 | 8293.05966 |
| *TGL1* | 0.029450506 | 0.85408328 | 0.99979869 | 13744.0239 | 14027.4695 |
| *YAL065C* | 0.063603121 | 0.85410764 | 0.99979869 | 356.062795 | 372.11133 |
| *PRM7* | 0.028946691 | 0.85454392 | 0.99979869 | 19050.8228 | 19436.9227 |
| *YML083C* | 0.050156235 | 0.85490057 | 0.99979869 | 663.349864 | 686.817055 |
| *RME2* | 0.039215101 | 0.85541228 | 0.99979869 | 1497.41477 | 1538.67522 |
| *THI72* | 0.038166229 | 0.85561918 | 0.99979869 | 1664.22775 | 1708.84184 |
| *YOR1* | 0.028430357 | 0.85582956 | 0.99979869 | 25057.0656 | 25555.745 |
| *INP54* | 0.03132398 | 0.85586882 | 0.99979869 | 5253.14562 | 5368.44913 |
| *SAP30* | 0.030122008 | 0.85600985 | 0.99979869 | 7826.55288 | 7991.68024 |
| *SNF11* | 0.030344736 | 0.85624045 | 0.99979869 | 7036.38613 | 7185.95157 |
| *ARA2* | 0.033394606 | 0.85668436 | 0.99979869 | 3141.15671 | 3214.71386 |

| | | | | | |
|---|---|---|---|---|---|
| *AIR2* | 0.032020694 | 0.85730917 | 0.99979869 | 4044.48314 | 4135.25373 |
| *tI(UAU)L* | 0.27798419 | 0.85771029 | 0.99979869 | 12.6816886 | 15.3765012 |
| *YFH1* | 0.02927978 | 0.85782051 | 0.99979869 | 9633.20575 | 9830.70978 |
| *CPT1* | 0.028327563 | 0.85880843 | 0.99979869 | 15342.8922 | 15647.1276 |
| *PHO80* | 0.033017548 | 0.85931889 | 0.99979869 | 2999.70711 | 3069.14964 |
| *ACS1* | 0.032022801 | 0.85932055 | 0.99979869 | 3647.44874 | 3729.3141 |
| *YMR090W* | 0.031303279 | 0.85950522 | 0.99979869 | 4257.1453 | 4350.52475 |
| *ERC1* | 0.027692318 | 0.8598125 | 0.99979869 | 23142.1306 | 23590.6282 |
| *UBP1* | 0.027527723 | 0.86047786 | 0.99979869 | 23998.6324 | 24460.9382 |
| *SPS18* | 0.038912841 | 0.86130585 | 0.99979869 | 1309.14047 | 1344.93131 |
| *MKT1* | 0.02710624 | 0.86133883 | 0.99979869 | 34455.1724 | 35108.6527 |
| *TAF14* | 0.027515487 | 0.86147811 | 0.99979869 | 19446.8817 | 19821.3352 |
| *YHR214C-B* | 0.028621122 | 0.86183181 | 0.99979869 | 8788.41019 | 8964.50022 |
| *IRC20* | 0.027372819 | 0.86286064 | 0.99979869 | 17054.9201 | 17381.597 |
| *YDL085C-A* | 0.02793889 | 0.86298079 | 0.99979869 | 11111.1102 | 11328.381 |
| *SPT16* | 0.026643321 | 0.86300829 | 0.99979869 | 44677.5889 | 45510.3432 |
| *IES5* | 0.02953137 | 0.86308318 | 0.99979869 | 5576.04092 | 5691.35565 |
| *STE18* | 0.029253356 | 0.86315528 | 0.99979869 | 6080.38192 | 6204.9308 |
| *YPL109C* | 0.029936197 | 0.86434396 | 0.99979869 | 4565.40789 | 4661.13007 |
| *RBH2* | 0.030488686 | 0.86452835 | 0.99979869 | 3930.34795 | 4014.29192 |
| *YLR255C* | 0.053112331 | 0.86512708 | 0.99979869 | 461.418361 | 478.721738 |
| *RPO41* | 0.027169301 | 0.86550144 | 0.99979869 | 13047.5065 | 13295.5481 |
| *RBD2* | 0.02648875 | 0.8661049 | 0.99979869 | 21422.2986 | 21819.2552 |
| *SRL2* | 0.030085656 | 0.866143 | 0.99979869 | 3961.56441 | 4045.04492 |
| *BEM3* | 0.026919168 | 0.86643058 | 0.99979869 | 13616.2315 | 13872.6794 |
| *OCH1* | 0.026922456 | 0.86718437 | 0.99979869 | 12175.3966 | 12404.7361 |
| *PDS1* | 0.029957043 | 0.86724186 | 0.99979869 | 3846.4537 | 3927.15841 |
| *IML1* | 0.02753596 | 0.86747313 | 0.99979869 | 8347.47763 | 8508.33068 |
| *NUP145* | 0.026047137 | 0.86751419 | 0.99979869 | 26152.5684 | 26629.0248 |
| *CEX1* | 0.027624893 | 0.86760552 | 0.99979869 | 7908.4961 | 8061.38704 |
| *PEX29* | 0.02719947 | 0.86771983 | 0.99979869 | 9639.05883 | 9822.50898 |
| *SPT3* | 0.027394064 | 0.86776418 | 0.99979869 | 8677.20153 | 8843.53841 |
| *UME1* | 0.027576912 | 0.86815069 | 0.99979869 | 7672.42159 | 7820.48852 |
| *CUE4* | 0.028140708 | 0.86845225 | 0.99979869 | 5992.58561 | 6110.62159 |
| *LTE1* | 0.028577575 | 0.86863824 | 0.99979869 | 5121.45116 | 5223.91002 |
| *PAU10* | 0.215922704 | 0.86891637 | 0.99979869 | 18.5347756 | 21.5271017 |
| *tV(AAC)K2* | 0.215922704 | 0.86891637 | 0.99979869 | 18.5347756 | 21.5271017 |
| *RPB8* | 0.025637282 | 0.8689387 | 0.99979869 | 31717.8787 | 32286.5522 |
| *SEC15* | 0.026386708 | 0.86978734 | 0.99979869 | 12159.7883 | 12384.2341 |
| *GTF1* | 0.028343019 | 0.86987145 | 0.99979869 | 5060.96926 | 5161.37891 |
| *MPH3* | 0.058552211 | 0.87008422 | 0.99979869 | 326.79736 | 340.333227 |
| *tA(AGC)K2* | 0.209036241 | 0.8700971 | 0.99979869 | 19.5102901 | 22.5522018 |
| *RPB7* | 0.026192983 | 0.87030545 | 0.99979869 | 12928.4937 | 13165.3604 |
| *KEL1* | 0.025935327 | 0.870403 | 0.99979869 | 15616.0362 | 15899.3023 |
| *CHA1* | 0.024879425 | 0.87127535 | 0.99979869 | 64046.4294 | 65160.4867 |

| | | | | | |
|---|---|---|---|---|---|
| *YNL234W* | 0.027162109 | 0.87171069 | 0.99979869 | 6635.44967 | 6761.56014 |
| *COS6* | 0.02550593 | 0.87184073 | 0.99979869 | 17740.7068 | 18057.1379 |
| *YGR045C* | 0.065653615 | 0.8718479 | 0.99979869 | 239.976569 | 251.14952 |
| *YIH1* | 0.025597257 | 0.87185451 | 0.99979869 | 16218.9042 | 16509.2368 |
| *NPY1* | 0.026492825 | 0.87194876 | 0.99979869 | 8760.12026 | 8922.47111 |
| *NUP57* | 0.025248688 | 0.87196831 | 0.99979869 | 23020.1913 | 23426.6122 |
| *RAD10* | 0.03241816 | 0.87233106 | 0.99979869 | 1969.56379 | 2014.32166 |
| *NDC80* | 0.026698925 | 0.87333593 | 0.99979869 | 6950.54086 | 7080.36626 |
| *RXT3* | 0.026339354 | 0.87346362 | 0.99979869 | 8066.52945 | 8215.15206 |
| *MTC3* | 0.029381099 | 0.87355754 | 0.99979869 | 3150.91185 | 3215.73896 |
| *CLB1* | 0.026089281 | 0.87375012 | 0.99979869 | 8872.30443 | 9034.20702 |
| *CTF3* | 0.028759353 | 0.87379144 | 0.99979869 | 3573.30964 | 3645.25589 |
| *SHE1* | 0.030094561 | 0.87394791 | 0.99979869 | 2687.54246 | 2744.19292 |
| *YER138C* | 0.071530294 | 0.8742211 | 0.99979869 | 188.2743 | 197.844316 |
| *YLR046C* | 0.029928837 | 0.87500633 | 0.99979869 | 2642.6688 | 2698.06342 |
| *AAD4* | 0.031394643 | 0.87510204 | 0.99979869 | 2062.23767 | 2107.60577 |
| *tD(GUC)D* | 0.171067881 | 0.87591004 | 0.99979869 | 27.3144062 | 30.7530025 |
| *GIM4* | 0.024676367 | 0.87599704 | 0.99979869 | 17523.1671 | 17825.4653 |
| *SCT1* | 0.024363608 | 0.87600671 | 0.99979869 | 25537.0187 | 25971.9357 |
| *JIP3* | 0.030887769 | 0.87612546 | 0.99979869 | 2142.22986 | 2188.58867 |
| *YLR236C* | 0.046726547 | 0.87623331 | 0.99979869 | 515.071659 | 532.026942 |
| *ARP4* | 0.024705832 | 0.87626016 | 0.99979869 | 16159.3978 | 16438.5049 |
| *SUP7* | 0.16774747 | 0.87632593 | 0.99979869 | 28.2899207 | 31.7781025 |
| *MLH3* | 0.026361781 | 0.87633715 | 0.99979869 | 6139.8883 | 6253.1105 |
| *ATO3* | 0.024274229 | 0.87649713 | 0.99979869 | 25217.05 | 25644.9287 |
| *YML053C* | 0.0260834 | 0.87656068 | 0.99979869 | 6732.02561 | 6854.84425 |
| *FRE8* | 0.026695079 | 0.87670528 | 0.99979869 | 5292.1662 | 5391.00133 |
| *SPR2* | 0.158994852 | 0.87730354 | 0.99979869 | 31.2164642 | 34.8534028 |
| *LSM3* | 0.024577318 | 0.87749542 | 0.99979869 | 14470.7822 | 14719.4121 |
| *MTG2* | 0.027027224 | 0.87765329 | 0.99979869 | 4463.95438 | 4548.36906 |
| *LEM3* | 0.024662091 | 0.87769415 | 0.99979869 | 13062.1392 | 13287.3473 |
| *FHN1* | 0.034741955 | 0.87774804 | 0.99979869 | 1239.87894 | 1270.099 |
| *VBA1* | 0.024600947 | 0.8777608 | 0.99979869 | 13563.5537 | 13796.822 |
| *LIH1* | 0.026922926 | 0.87783077 | 0.99979869 | 4549.79966 | 4635.50257 |
| *HOT1* | 0.025384899 | 0.87791343 | 0.99979869 | 8119.20723 | 8263.33176 |
| *RRT2* | 0.025110671 | 0.87814336 | 0.99979869 | 9150.32607 | 9310.98404 |
| *OCA4* | 0.02670978 | 0.87814555 | 0.99979869 | 4751.73116 | 4840.52259 |
| *tR(UCU)G1* | 0.147480659 | 0.87824071 | 0.99979869 | 36.0940367 | 39.9789032 |
| *HSP32* | 0.135661912 | 0.87858037 | 0.99979869 | 42.9226383 | 47.1546038 |
| *CBP6* | 0.027846544 | 0.87862994 | 0.99979869 | 3401.61908 | 3467.91358 |
| *NCA2* | 0.026072429 | 0.8790269 | 0.99979869 | 5503.85284 | 5604.22215 |
| *MRP10* | 0.025852144 | 0.879464 | 0.99979869 | 5759.43764 | 5863.57247 |
| *tK(UUU)D* | 0.056833598 | 0.88006143 | 0.99979869 | 288.752294 | 300.354324 |
| *IZH4* | 0.025902704 | 0.8806992 | 0.99979869 | 5139.01042 | 5232.11082 |
| *YHR078W* | 0.024421209 | 0.88073377 | 0.99979869 | 9990.24406 | 10160.792 |

| | | | | | |
|---|---|---|---|---|---|
| *MAD1* | 0.024078116 | 0.88079789 | 0.99979869 | 12542.19 | 12753.2701 |
| *SLH1* | 0.023984128 | 0.88089458 | 0.99979869 | 13300.1648 | 13523.1203 |
| *MDM20* | 0.024323288 | 0.88090384 | 0.99979869 | 10394.1071 | 10570.832 |
| *MNL1* | 0.024045496 | 0.88105374 | 0.99979869 | 12352.9402 | 12560.5513 |
| *YGR125W* | 0.023880235 | 0.88130875 | 0.99979869 | 13511.8514 | 13737.3662 |
| *PFD1* | 0.025420697 | 0.88136216 | 0.99979869 | 5799.43374 | 5902.52627 |
| *RPB9* | 0.024780345 | 0.88141888 | 0.99979869 | 7587.55183 | 7719.00362 |
| *ACT1* | 0.022636935 | 0.88157314 | 0.99979869 | 358505.483 | 364175.005 |
| *GLY1* | 0.022822978 | 0.88166331 | 0.99979869 | 67938.7323 | 69022.0387 |
| *VEL1* | 0.048016394 | 0.88196852 | 0.99979869 | 422.397781 | 436.692635 |
| *SBH2* | 0.023061006 | 0.88239779 | 0.99979869 | 26633.497 | 27062.6422 |
| *CAB5* | 0.024168888 | 0.88245707 | 0.99979869 | 9363.96374 | 9522.15466 |
| *YLR462W* | 0.059848374 | 0.88271749 | 0.99979869 | 241.927598 | 252.17462 |
| *PFK1* | 0.022422562 | 0.88297916 | 0.99979869 | 159915.117 | 162419.932 |
| *BET4* | 0.02452194 | 0.88309399 | 0.99979869 | 7242.21969 | 7366.36919 |
| *PHO90* | 0.023061664 | 0.88316638 | 0.99979869 | 21318.894 | 21662.4149 |
| *TOS1* | 0.022338553 | 0.88361191 | 0.99979869 | 116519.355 | 118337.553 |
| *RNR3* | 0.030453827 | 0.88434111 | 0.99979869 | 1668.12981 | 1703.71634 |
| *SEC21* | 0.022392761 | 0.88434522 | 0.99979869 | 48884.0074 | 49648.6723 |
| *PET111* | 0.026770173 | 0.8847676 | 0.99979869 | 3129.45054 | 3188.06125 |
| *VMA22* | 0.02408654 | 0.88479127 | 0.99979869 | 7508.53515 | 7634.94541 |
| *ZIP1* | 0.02718268 | 0.88513831 | 0.99979869 | 2803.62869 | 2856.95393 |
| *YMR030W-A* | 0.06426228 | 0.88543948 | 0.99979869 | 194.127387 | 202.969816 |
| *BRF1* | 0.023187469 | 0.88566747 | 0.99979869 | 11545.2142 | 11732.2704 |
| *GCN20* | 0.022192087 | 0.88570723 | 0.99979869 | 40709.1959 | 41340.2361 |
| *YPL071C* | 0.02633318 | 0.88572832 | 0.99979869 | 3289.43491 | 3350.02707 |
| *ROT1* | 0.022617075 | 0.88662235 | 0.99979869 | 15977.9521 | 16230.4096 |
| *MOT2* | 0.02299776 | 0.88704026 | 0.99979869 | 10793.0925 | 10966.5207 |
| *CDC7* | 0.025380561 | 0.88738756 | 0.99979869 | 3811.33518 | 3878.97871 |
| *YBR096W* | 0.022837569 | 0.88772798 | 0.99979869 | 10934.5421 | 11109.0096 |
| *SUF5* | 0.086030303 | 0.88788016 | 0.99979869 | 96.5759361 | 102.510008 |
| *APL5* | 0.022074034 | 0.88788018 | 0.99979869 | 22666.0795 | 23015.547 |
| *ESS1* | 0.021864496 | 0.88875184 | 0.99979869 | 23898.1544 | 24263.0938 |
| *GPA2* | 0.021718133 | 0.88897975 | 0.99979869 | 28255.7777 | 28684.3505 |
| *YCR090C* | 0.02262585 | 0.89066411 | 0.99979869 | 8458.68628 | 8592.38889 |
| *PCH2* | 0.02848675 | 0.89186765 | 0.99979869 | 1659.35017 | 1692.44024 |
| *MRS1* | 0.024142835 | 0.89193315 | 0.99979869 | 4045.45866 | 4113.72663 |
| *HLJ1* | 0.021992019 | 0.8921434 | 0.99979869 | 10433.1276 | 10593.3842 |
| *YGL204C* | 0.027071504 | 0.89215831 | 0.99979869 | 2057.36009 | 2096.32967 |
| *TIF4631* | 0.020676432 | 0.89245948 | 0.99979869 | 80525.7959 | 81688.1753 |
| *ASK1* | 0.022134439 | 0.89279059 | 0.99979869 | 8694.76079 | 8829.187 |
| *SLX8* | 0.023042098 | 0.89287857 | 0.99979869 | 5519.46108 | 5608.32255 |
| *MZM1* | 0.024089267 | 0.89297786 | 0.99979869 | 3829.86995 | 3894.35521 |
| *CDC53* | 0.021002156 | 0.89307226 | 0.99979869 | 23979.1221 | 24330.7504 |
| *CUP2* | 0.021692938 | 0.89312901 | 0.99979869 | 11204.7596 | 11374.5105 |

| | | | | | |
|---|---|---|---|---|---|
| *YMR242W-A* | 0.054182931 | 0.89323571 | 0.99979869 | 244.854141 | 254.22482 |
| *SUI2* | 0.020629551 | 0.89382562 | 0.99979869 | 37229.5356 | 37765.7121 |
| *YGR117C* | 0.021859039 | 0.89384931 | 0.99979869 | 8993.26823 | 9130.56643 |
| *HHO1* | 0.020960888 | 0.89441127 | 0.99979869 | 17559.2611 | 17816.2394 |
| *RTA1* | 0.027437183 | 0.89451879 | 0.99979869 | 1750.07302 | 1783.67414 |
| *AVT5* | 0.022691722 | 0.89469045 | 0.99979869 | 5421.90962 | 5507.86274 |
| *MYO4* | 0.020672589 | 0.89476418 | 0.99979869 | 23713.7821 | 24056.0236 |
| *YPR203W* | 0.063018866 | 0.89515032 | 0.99979869 | 165.837466 | 173.241914 |
| *ZRT1* | 0.023047081 | 0.89521851 | 0.99979869 | 4545.8976 | 4619.10097 |
| *GCS1* | 0.020437625 | 0.89550484 | 0.99979869 | 27615.8402 | 28009.8346 |
| *YML082W* | 0.021366652 | 0.89554271 | 0.99979869 | 9877.08437 | 10024.4537 |
| *SRP14* | 0.020973906 | 0.89650173 | 0.99979869 | 11444.7362 | 11612.3337 |
| *tW(CCA)P* | 0.053382848 | 0.89678956 | 0.99979869 | 234.123481 | 242.948719 |
| *RSC30* | 0.021304793 | 0.89715208 | 0.99979869 | 8268.46095 | 8391.46927 |
| *AIM41* | 0.0211128 | 0.89751639 | 0.99979869 | 8890.83921 | 9021.90582 |
| *YJR142W* | 0.020842784 | 0.89753199 | 0.99979869 | 10737.4882 | 10893.7386 |
| *VPS36* | 0.021261495 | 0.89761703 | 0.99979869 | 8005.07204 | 8123.91815 |
| *MAL12* | 0.045766967 | 0.89781114 | 0.99979869 | 330.699418 | 341.358327 |
| *YOR097C* | 0.022144438 | 0.89789277 | 0.99979869 | 5101.94087 | 5180.85581 |
| *GUS1* | 0.019538066 | 0.89799198 | 0.99979869 | 121704.214 | 123363.619 |
| *RRG7* | 0.021797806 | 0.89799401 | 0.99979869 | 5844.30741 | 5933.27927 |
| *OST4* | 0.021384901 | 0.89818058 | 0.99979869 | 6996.39004 | 7100.86827 |
| *MOT1* | 0.020139184 | 0.89845826 | 0.99979869 | 17757.2906 | 18006.908 |
| *MSN5* | 0.019902915 | 0.89861769 | 0.99979869 | 23865.9624 | 24197.4874 |
| *CDC16* | 0.020762983 | 0.89861914 | 0.99979869 | 9648.81398 | 9788.68068 |
| *OMA1* | 0.021537206 | 0.89862674 | 0.99979869 | 6186.713 | 6279.7631 |
| *POP1* | 0.020067345 | 0.89866636 | 0.99979869 | 18456.7345 | 18715.2522 |
| *HIR3* | 0.02073312 | 0.89870795 | 0.99979869 | 9727.83065 | 9868.63849 |
| *RTR2* | 0.025325966 | 0.89919788 | 0.99979869 | 2042.72738 | 2078.90297 |
| *POF1* | 0.022276993 | 0.89956501 | 0.99979869 | 4243.4881 | 4309.52074 |
| *YGR035W-A* | 0.02725677 | 0.89979751 | 0.99979869 | 1452.5411 | 1480.24452 |
| *TRS130* | 0.020801545 | 0.89993633 | 0.99979869 | 7877.27964 | 7991.68024 |
| *ERG8* | 0.019674507 | 0.90052446 | 0.99979869 | 18982.5368 | 19243.1787 |
| *UBA3* | 0.022647984 | 0.90068673 | 0.99979869 | 3484.53782 | 3539.67058 |
| *AIM14* | 0.02035582 | 0.90114354 | 0.99979869 | 8901.56987 | 9028.05642 |
| *GRX8* | 0.026606198 | 0.90145448 | 0.99979869 | 1495.46374 | 1523.29872 |
| *PAU24* | 0.044730017 | 0.90168969 | 0.99979869 | 318.017729 | 328.032026 |
| *SPC72* | 0.022136039 | 0.90201349 | 0.99979869 | 3709.88167 | 3767.2428 |
| *VPS63* | 0.024541281 | 0.90202188 | 0.99979869 | 2070.04178 | 2105.55557 |
| *TIP41* | 0.020442031 | 0.90202609 | 0.99979869 | 7514.38824 | 7621.61911 |
| *RPL21B* | 0.01868949 | 0.90216265 | 0.99979869 | 187787.518 | 190236.023 |
| *YNL277W-A* | 0.096193319 | 0.90221209 | 0.99979869 | 56.5798414 | 60.4809048 |
| *RQC1* | 0.019708996 | 0.90230495 | 0.99979869 | 12248.5601 | 12417.0373 |
| *ARP8* | 0.019461546 | 0.90261092 | 0.99979869 | 14750.7548 | 14951.0847 |
| *CET1* | 0.020097518 | 0.90262567 | 0.99979869 | 8606.96449 | 8727.7021 |

| | | | | | |
|---|---|---|---|---|---|
| *CMG1* | 0.021983408 | 0.90317393 | 0.99979869 | 3583.06478 | 3638.08019 |
| *YFR006W* | 0.019064781 | 0.90324049 | 0.99979869 | 20951.125 | 21229.8227 |
| *CMC2* | 0.020903647 | 0.90334041 | 0.99979869 | 5205.3454 | 5281.31562 |
| *SAP1* | 0.020116287 | 0.90341619 | 0.99979869 | 7662.66645 | 7770.25862 |
| *JAC1* | 0.021809926 | 0.90342569 | 0.99979869 | 3714.75924 | 3771.3432 |
| *UBP5* | 0.019119537 | 0.90345154 | 0.99979869 | 18210.9048 | 18453.8517 |
| *DAL7* | 0.022141706 | 0.90393781 | 0.99979869 | 3246.51228 | 3296.72186 |
| *SFB3* | 0.018552799 | 0.90445627 | 0.99979869 | 36120.3756 | 36587.8721 |
| *ERG7* | 0.018692988 | 0.90484045 | 0.99979869 | 22868.0111 | 23166.2367 |
| *MF(ALPHA)1* | 0.100100598 | 0.90508605 | 0.99979869 | 48.7757253 | 52.2801042 |
| *YCS4* | 0.019626018 | 0.90546691 | 0.99979869 | 7950.44322 | 8059.33684 |
| *DOA1* | 0.018613372 | 0.90549847 | 0.99979869 | 21019.4111 | 21292.3538 |
| *DAK1* | 0.018411176 | 0.90561722 | 0.99979869 | 29306.4068 | 29682.798 |
| *SPO23* | 0.036340524 | 0.90564489 | 0.99979869 | 485.806224 | 498.19864 |
| *CTI6* | 0.018763455 | 0.9058059 | 0.99979869 | 15699.9305 | 15905.4529 |
| *BNA4* | 0.020161327 | 0.90584159 | 0.99979869 | 5689.2006 | 5769.26326 |
| *VPS62* | 0.019021754 | 0.90597858 | 0.99979869 | 11572.5286 | 11726.1198 |
| *tI(AAU)N1* | 0.209037586 | 0.90600204 | 0.99979869 | 9.75514506 | 11.2761009 |
| *YFR018C* | 0.018556971 | 0.90606301 | 0.99979869 | 19292.7504 | 19542.508 |
| *YLR352W* | 0.020383513 | 0.90630903 | 0.99979869 | 4929.2748 | 4999.4131 |
| *RTC2* | 0.027766991 | 0.90669668 | 0.99979869 | 1044.77604 | 1065.07898 |
| *CDC24* | 0.018728296 | 0.90702276 | 0.99979869 | 12486.5857 | 12649.735 |
| *PDC2* | 0.019056696 | 0.90709607 | 0.99979869 | 9313.23699 | 9437.07135 |
| *TEP1* | 0.030414008 | 0.90732763 | 0.99979869 | 763.827858 | 780.101162 |
| *SEC8* | 0.019022147 | 0.90742187 | 0.99979869 | 9088.86865 | 9209.49913 |
| *YLL066W-B* | 0.197064805 | 0.90751932 | 0.99979869 | 10.7306596 | 12.301201 |
| *RCK2* | 0.017959463 | 0.90760625 | 0.99979869 | 33854.2554 | 34278.3216 |
| *IMO32* | 0.019271166 | 0.90761544 | 0.99979869 | 7486.09832 | 7586.76571 |
| *RMI1* | 0.021450813 | 0.90793751 | 0.99979869 | 3030.92357 | 3076.32535 |
| *UBR1* | 0.018050345 | 0.90821357 | 0.99979869 | 21847.6229 | 22122.6849 |
| *YHK8* | 0.020625759 | 0.9083342 | 0.99979869 | 3798.65349 | 3853.35121 |
| *IRC6* | 0.022054492 | 0.90859983 | 0.99979869 | 2488.53751 | 2526.8717 |
| *SUF8* | 0.187011003 | 0.90872501 | 0.99979869 | 11.7061741 | 13.3263011 |
| *tH(GUG)G2* | 0.187011003 | 0.90872501 | 0.99979869 | 11.7061741 | 13.3263011 |
| *KAR9* | 0.020888724 | 0.90881995 | 0.99979869 | 3365.52505 | 3414.60837 |
| *tR(UCU)M1* | 0.109006525 | 0.90915529 | 0.99979869 | 37.0695512 | 39.9789032 |
| *RPS22A* | 0.017322263 | 0.90922177 | 0.99979869 | 204062.026 | 206526.913 |
| *NPP1* | 0.018188861 | 0.90931225 | 0.99979869 | 13491.3656 | 13662.5339 |
| *RIM9* | 0.023617636 | 0.90967286 | 0.99979869 | 1702.27281 | 1730.36894 |
| *LRS4* | 0.020049807 | 0.91004133 | 0.99979869 | 4063.01792 | 4119.87723 |
| *SWP82* | 0.018761093 | 0.91008425 | 0.99979869 | 7442.20017 | 7539.6111 |
| *SBP1* | 0.017183652 | 0.91034598 | 0.99979869 | 91537.4037 | 92634.194 |
| *RNA170* | 0.040503178 | 0.91062255 | 0.99979869 | 320.944273 | 330.082226 |
| *STU2* | 0.018107144 | 0.91066103 | 0.99979869 | 11136.4736 | 11277.126 |
| *tT(AGU)B* | 0.164642805 | 0.91104603 | 0.99979869 | 14.6327176 | 16.4016013 |

| | | | | | |
|---|---|---|---|---|---|
| *MCH4* | 0.01852361 | 0.91130114 | 0.99979869 | 7355.37938 | 7450.42739 |
| *TIM21* | 0.017949381 | 0.91163251 | 0.99979869 | 10729.6841 | 10864.0107 |
| *DID2* | 0.017575511 | 0.9119512 | 0.99979869 | 14802.4571 | 14983.8879 |
| *DBF2* | 0.017910343 | 0.91203757 | 0.99979869 | 10320.9435 | 10449.8702 |
| *AMF1* | 0.020691686 | 0.91211287 | 0.99979869 | 2845.57581 | 2886.68183 |
| *RPL13A* | 0.016768 | 0.91222833 | 0.99979869 | 146222.796 | 147932.193 |
| *ANS1* | 0.141922208 | 0.91242514 | 0.99979869 | 19.5102901 | 21.5271017 |
| *BRR2* | 0.017298297 | 0.91242603 | 0.99979869 | 19011.8022 | 19241.1285 |
| *tF(GAA)H1* | 0.138646989 | 0.912484 | 0.99979869 | 20.4858046 | 22.5522018 |
| *SLY1* | 0.017309986 | 0.91254113 | 0.99979869 | 18037.2632 | 18254.9823 |
| *RPL37B* | 0.016650935 | 0.91286315 | 0.99979869 | 137788.497 | 139387.984 |
| *BIG1* | 0.019975822 | 0.91324376 | 0.99979869 | 3251.38985 | 3296.72186 |
| *LAM6* | 0.01762639 | 0.91392992 | 0.99979869 | 9442.98042 | 9559.05826 |
| *YMR111C* | 0.019116774 | 0.91489982 | 0.99979869 | 3827.91892 | 3878.97871 |
| *1-Oct* | 0.018550628 | 0.9149609 | 0.99979869 | 4788.80071 | 4850.77359 |
| *NAS6* | 0.017617785 | 0.91503907 | 0.99979869 | 7978.73315 | 8076.76354 |
| *PDR1* | 0.016916732 | 0.91515461 | 0.99979869 | 15073.6501 | 15251.439 |
| *VMA9* | 0.016813115 | 0.91533312 | 0.99979869 | 16513.5096 | 16707.0811 |
| *TOM71* | 0.016672029 | 0.91575261 | 0.99979869 | 17962.1486 | 18170.924 |
| *GIM5* | 0.017367628 | 0.91580631 | 0.99979869 | 8525.02127 | 8628.26739 |
| *TSC11* | 0.017029982 | 0.91627729 | 0.99979869 | 10420.446 | 10544.1794 |
| *PMT1* | 0.015841362 | 0.91744456 | 0.99979869 | 74687.3416 | 75511.9473 |
| *PEP5* | 0.016930675 | 0.9178631 | 0.99979869 | 8589.40523 | 8690.79849 |
| *PMT3* | 0.016599995 | 0.91820105 | 0.99979869 | 10784.3129 | 10909.1151 |
| *PYC2* | 0.015717919 | 0.91829878 | 0.99979869 | 58998.1418 | 59644.4232 |
| *CRH1* | 0.015675881 | 0.91848325 | 0.99979869 | 60932.5871 | 61598.2639 |
| *CTF8* | 0.018259619 | 0.91887446 | 0.99979869 | 3767.43702 | 3815.4225 |
| *LSM7* | 0.016640104 | 0.91894278 | 0.99979869 | 9067.40733 | 9172.59553 |
| *SFB2* | 0.016179874 | 0.91914399 | 0.99979869 | 13865.9632 | 14022.344 |
| *CTF4* | 0.016868243 | 0.91917987 | 0.99979869 | 7346.59975 | 7433.00069 |
| *YOL075C* | 0.016633552 | 0.91976064 | 0.99979869 | 7968.978 | 8061.38704 |
| *YDR344C* | 0.024369099 | 0.91983381 | 0.99979869 | 970.636934 | 987.171379 |
| *APE2* | 0.015556365 | 0.91995249 | 0.99979869 | 32376.3509 | 32727.3452 |
| *YPL199C* | 0.015868862 | 0.92002645 | 0.99979869 | 16651.0571 | 16835.2186 |
| *EPS1* | 0.015902264 | 0.92032501 | 0.99979869 | 14597.5991 | 14759.391 |
| *LEU5* | 0.018506337 | 0.92062761 | 0.99979869 | 3000.68262 | 3039.42174 |
| *SSZ1* | 0.015116956 | 0.92080481 | 0.99979869 | 146704.7 | 148249.974 |
| *EHD3* | 0.015924113 | 0.92111347 | 0.99979869 | 11739.3416 | 11869.6338 |
| *CSE1* | 0.015418313 | 0.92117305 | 0.99979869 | 25035.6043 | 25304.5955 |
| *MTR3* | 0.016102243 | 0.92187121 | 0.99979869 | 8566.96839 | 8663.12079 |
| *BPT1* | 0.015372534 | 0.92188476 | 0.99979869 | 20595.0623 | 20815.6823 |
| *UFD4* | 0.015454034 | 0.92196285 | 0.99979869 | 17400.2522 | 17587.6421 |
| *NFI1* | 0.017167435 | 0.92201035 | 0.99979869 | 4431.7624 | 4484.81286 |
| *FMP42* | 0.015607545 | 0.92203122 | 0.99979869 | 13701.1012 | 13850.1272 |
| *SDC25* | 0.016727333 | 0.92205684 | 0.99979869 | 5496.04873 | 5560.14284 |

| | | | | | |
|---|---|---|---|---|---|
| *RSC9* | 0.015376547 | 0.9224991 | 0.99979869 | 16610.0855 | 16788.064 |
| *RIM13* | 0.017437026 | 0.92283643 | 0.99979869 | 3658.1794 | 3702.6615 |
| *NUP188* | 0.015086687 | 0.92287458 | 0.99979869 | 24840.5014 | 25101.6257 |
| *YBL036C* | 0.015435603 | 0.92288904 | 0.99979869 | 13685.493 | 13832.7005 |
| *YML6* | 0.01575419 | 0.92323859 | 0.99979869 | 9054.72565 | 9154.14373 |
| *DFG16* | 0.015768158 | 0.92382661 | 0.99979869 | 8079.21114 | 8167.99745 |
| *YTA12* | 0.014825541 | 0.92413093 | 0.99979869 | 25664.8111 | 25929.9066 |
| *VPS29* | 0.015173484 | 0.92450418 | 0.99979869 | 12629.9863 | 12763.5211 |
| *CYC2* | 0.016606351 | 0.92452819 | 0.99979869 | 4439.56652 | 4490.96346 |
| *MLC2* | 0.015931106 | 0.92458905 | 0.99979869 | 6399.37516 | 6470.43172 |
| *SSP1* | 0.025593597 | 0.92476581 | 0.99979869 | 684.811183 | 697.068056 |
| *SDD4* | 0.015115259 | 0.92525406 | 0.99979869 | 11319.8703 | 11439.0918 |
| *YPL113C* | 0.017849759 | 0.92566176 | 0.99979869 | 2537.31323 | 2568.9008 |
| *RGP1* | 0.015351216 | 0.9257434 | 0.99979869 | 8199.19942 | 8286.90906 |
| *YDL012C* | 0.014698489 | 0.92682346 | 0.99979869 | 12728.5133 | 12858.8554 |
| *IMA5* | 0.022027863 | 0.92722492 | 0.99979869 | 983.318622 | 998.44748 |
| *GAS3* | 0.014125199 | 0.92750768 | 0.99979869 | 28258.7042 | 28536.7361 |
| *RDS2* | 0.015558401 | 0.92771349 | 0.99979869 | 5332.16229 | 5389.97623 |
| *HAP5* | 0.015425056 | 0.92805281 | 0.99979869 | 5524.33865 | 5583.72015 |
| *YJR012C* | 0.017061399 | 0.92844494 | 0.99979869 | 2632.91365 | 2664.23511 |
| *SUB1* | 0.013939604 | 0.92926448 | 0.99979869 | 19469.3185 | 19658.3443 |
| *YNR075C-A* | 0.020455624 | 0.92973947 | 0.99979869 | 1121.84168 | 1137.86109 |
| *MPH2* | 0.0557635 | 0.92985205 | 0.99979869 | 89.7473346 | 93.2841074 |
| *YVC1* | 0.014084894 | 0.93005662 | 0.99979869 | 12095.4044 | 12214.0675 |
| *GIM3* | 0.013759034 | 0.93027418 | 0.99979869 | 18738.6581 | 18918.222 |
| *BCD1* | 0.01430681 | 0.9307179 | 0.99979869 | 8278.2161 | 8360.71627 |
| *RPL40B* | 0.013155354 | 0.93098544 | 0.99979869 | 163592.807 | 165091.343 |
| *MDH3* | 0.013221006 | 0.93180488 | 0.99979869 | 34577.1117 | 34895.4319 |
| *SRF1* | 0.01433861 | 0.93249212 | 0.99979869 | 6023.80208 | 6083.96899 |
| *FUR4* | 0.015158234 | 0.93272038 | 0.99979869 | 3716.71027 | 3755.9667 |
| *GVP36* | 0.01296149 | 0.9327449 | 0.99979869 | 47741.6799 | 48172.5281 |
| *MIP6* | 0.023215953 | 0.93289708 | 0.99979869 | 651.64369 | 662.214653 |
| *SPT5* | 0.012765826 | 0.93355017 | 0.99979869 | 59635.1528 | 60165.174 |
| *YOR376W-A* | 0.017476868 | 0.93360392 | 0.99979869 | 1644.71746 | 1664.76253 |
| *CSE4* | 0.014571512 | 0.93451454 | 0.99979869 | 4057.16483 | 4098.35013 |
| *BIO3* | 0.018032239 | 0.93460128 | 0.99979869 | 1366.69582 | 1383.88511 |
| *YAL037C-A* | 0.017758526 | 0.9347173 | 0.99979869 | 1439.85941 | 1457.69232 |
| *MPD2* | 0.0133177 | 0.93481199 | 0.99979869 | 9546.38496 | 9634.91567 |
| *SNX41* | 0.013776025 | 0.93513805 | 0.99979869 | 6015.99796 | 6073.71798 |
| *RED1* | 0.015273131 | 0.93539389 | 0.99979869 | 2754.85297 | 2784.17182 |
| *RPN2* | 0.012342359 | 0.93546163 | 0.99979869 | 91859.3235 | 92648.5454 |
| *tD(GUC)G2* | 0.071531623 | 0.93565418 | 0.99979869 | 43.8981528 | 46.1295037 |
| *RPN9* | 0.012552485 | 0.93584071 | 0.99979869 | 23866.9379 | 24075.5005 |
| *RPA14* | 0.012882815 | 0.9362939 | 0.99979869 | 11166.7146 | 11266.875 |
| *YPP1* | 0.012644158 | 0.93660736 | 0.99979869 | 14415.1779 | 14542.0698 |

| | | | | | |
|---|---|---|---|---|---|
| *YCR001W* | 0.029164065 | 0.93708779 | 0.99979869 | 303.385011 | 309.580225 |
| *RDR1* | 0.013985052 | 0.93755637 | 0.99979869 | 3866.9395 | 3904.60621 |
| *OST2* | 0.012863462 | 0.93769465 | 0.99979869 | 8249.92618 | 8323.81266 |
| *YLR111W* | 0.030010863 | 0.93785552 | 0.99979869 | 275.095091 | 280.877422 |
| *YHL049C* | 0.036973156 | 0.93867688 | 0.99979869 | 164.861952 | 169.141513 |
| *AIM33* | 0.017082012 | 0.93931236 | 0.99979869 | 1264.2668 | 1279.3249 |
| *PUT3* | 0.012338407 | 0.93945724 | 0.99979869 | 9852.69651 | 9937.32019 |
| *EXO1* | 0.012762821 | 0.93989488 | 0.99979869 | 6012.0959 | 6065.51718 |
| *PRP43* | 0.011521921 | 0.94025157 | 0.99979869 | 44440.5388 | 44796.8736 |
| *MFG1* | 0.012304137 | 0.94090455 | 0.99979869 | 7421.71436 | 7485.2808 |
| *SLM5* | 0.013582384 | 0.94099019 | 0.99979869 | 3221.1489 | 3251.61746 |
| *HEM3* | 0.011715234 | 0.94120136 | 0.99979869 | 14652.2279 | 14771.6922 |
| *VTH2* | 0.0209883 | 0.94123729 | 0.99979869 | 595.063849 | 603.783948 |
| *IRC25* | 0.012614359 | 0.9412446 | 0.99979869 | 5412.15448 | 5459.68304 |
| *SGV1* | 0.011734552 | 0.9413181 | 0.99979869 | 13599.6477 | 13710.7136 |
| *YOR011W-A* | 0.051631488 | 0.94173282 | 0.99979869 | 71.2125589 | 73.8072059 |
| *YIL029C* | 0.014784944 | 0.943016 | 0.99979869 | 1745.19545 | 1763.17214 |
| *MUD1* | 0.011826476 | 0.94378825 | 0.99979869 | 6593.50255 | 6647.77403 |
| *KNH1* | 0.015794006 | 0.94391479 | 0.99979869 | 1261.34026 | 1275.2245 |
| *SND1* | 0.011035003 | 0.94410855 | 0.99979869 | 18030.4346 | 18168.8738 |
| *tC(GCA)P2* | 0.071532105 | 0.94448594 | 0.99979869 | 32.1919787 | 33.8283027 |
| *HRT3* | 0.011746581 | 0.94499803 | 0.99979869 | 5671.64134 | 5718.00826 |
| *OPT2* | 0.012732842 | 0.94522429 | 0.99979869 | 3028.97254 | 3055.82334 |
| *PEX9* | 0.018600432 | 0.94550848 | 0.99979869 | 677.007067 | 685.791955 |
| *MAC1* | 0.011547612 | 0.94577991 | 0.99979869 | 5820.89506 | 5867.67287 |
| *SRB5* | 0.012050883 | 0.94589663 | 0.99979869 | 4009.36462 | 4042.99472 |
| *MTC6* | 0.011832101 | 0.94591883 | 0.99979869 | 4596.62435 | 4634.47747 |
| *JIP5* | 0.010596094 | 0.94608822 | 0.99979869 | 20309.2365 | 20458.9474 |
| *tA(AGC)P* | 0.071532212 | 0.9461156 | 0.99979869 | 30.2409497 | 31.7781025 |
| *ENA2* | 0.031341248 | 0.94629268 | 0.99979869 | 177.54364 | 181.442714 |
| *NNF2* | 0.010675489 | 0.94672335 | 0.99979869 | 12989.9512 | 13086.4276 |
| *tS(GCU)O* | 0.07153227 | 0.94695112 | 0.99979869 | 29.2654352 | 30.7530025 |
| *EFT1* | 0.01008017 | 0.94715561 | 0.99979869 | 117344.64 | 118167.387 |
| *SUL2* | 0.011577564 | 0.94731911 | 0.99979869 | 4433.71343 | 4469.43636 |
| *LDS1* | 0.071532331 | 0.94780132 | 0.99979869 | 28.2899207 | 29.7279024 |
| *DYN2* | 0.010751285 | 0.94798789 | 0.99979869 | 8063.60291 | 8123.91815 |
| *MEP1* | 0.010638011 | 0.9487292 | 0.99979869 | 7694.85842 | 7751.80682 |
| *SOG2* | 0.010554026 | 0.94901295 | 0.99979869 | 7920.20228 | 7978.35394 |
| *RAD50* | 0.010613667 | 0.94938052 | 0.99979869 | 6818.8464 | 6869.19565 |
| *YRF1-1* | 0.071532462 | 0.94954876 | 0.99979869 | 26.3388917 | 27.6777022 |
| *tH(GUG)E2* | 0.035310414 | 0.94966852 | 0.99979869 | 118.037255 | 120.96181 |
| *RPL5* | 0.009482115 | 0.95003101 | 0.99979869 | 531457.376 | 534961.804 |
| *RRP7* | 0.009870828 | 0.95018827 | 0.99979869 | 16344.7455 | 16456.9567 |
| *ERG28* | 0.010262906 | 0.95026214 | 0.99979869 | 8235.29346 | 8294.08476 |
| *ULP1* | 0.010070642 | 0.95036164 | 0.99979869 | 10455.5645 | 10528.8029 |

| | | | | | |
|---|---|---|---|---|---|
| *AUS1* | 0.010399636 | 0.95057626 | 0.99979869 | 6560.33505 | 6607.79513 |
| *YMR230W-A* | 0.029219563 | 0.95105297 | 0.99979869 | 168.76401 | 172.216814 |
| *MKK1* | 0.009956494 | 0.95108965 | 0.99979869 | 9922.93356 | 9991.6505 |
| *MBB1* | 0.012972288 | 0.95123968 | 0.99979869 | 1569.60284 | 1583.77963 |
| *GAL7* | 0.016603673 | 0.95130662 | 0.99979869 | 678.958096 | 686.817055 |
| *RAD14* | 0.010129171 | 0.95186211 | 0.99979869 | 6555.45748 | 6601.64453 |
| *TAE1* | 0.009410567 | 0.95236578 | 0.99979869 | 17545.6039 | 17660.4242 |
| *IRR1* | 0.009975611 | 0.95271563 | 0.99979869 | 6375.96281 | 6420.20181 |
| *COG1* | 0.009836431 | 0.9529041 | 0.99979869 | 7107.59869 | 7156.22367 |
| *YPT11* | 0.010387997 | 0.9529718 | 0.99979869 | 4255.19428 | 4285.94344 |
| *CDC9* | 0.009652282 | 0.95316817 | 0.99979869 | 8341.62454 | 8397.61987 |
| *MON1* | 0.009763527 | 0.95329816 | 0.99979869 | 7029.55753 | 7077.29096 |
| *RKM5* | 0.0099907 | 0.95354185 | 0.99979869 | 5302.89686 | 5339.74633 |
| *HNT3* | 0.011522445 | 0.95368538 | 0.99979869 | 2107.11133 | 2124.00737 |
| *MSK1* | 0.009283663 | 0.9537576 | 0.99979869 | 12451.4672 | 12531.8485 |
| *NSE5* | 0.0097424 | 0.95385169 | 0.99979869 | 6329.13812 | 6372.02211 |
| *SNT309* | 0.009834869 | 0.95416719 | 0.99979869 | 5406.30139 | 5443.28143 |
| *RFC2* | 0.009154455 | 0.95448095 | 0.99979869 | 12057.3593 | 12134.1097 |
| *YPT52* | 0.00865251 | 0.95598565 | 0.99979869 | 19900.4959 | 20020.2046 |
| *RPN14* | 0.00922423 | 0.95598636 | 0.99979869 | 6831.52809 | 6875.34625 |
| *KAP114* | 0.009051001 | 0.95601 | 0.99979869 | 8516.24164 | 8569.83668 |
| *YPR071W* | 0.01028255 | 0.95635848 | 0.99979869 | 2792.89803 | 2812.87462 |
| *RSC2* | 0.008884345 | 0.95641616 | 0.99979869 | 9687.83456 | 9747.67668 |
| *COS4* | 0.04456436 | 0.95642401 | 0.99979869 | 52.6777833 | 54.3303043 |
| *CBS2* | 0.009641907 | 0.95667759 | 0.99979869 | 4019.11977 | 4046.07002 |
| *PGS1* | 0.009818114 | 0.95668267 | 0.99979869 | 3541.11766 | 3565.29808 |
| *CLB4* | 0.009866713 | 0.95701323 | 0.99979869 | 3264.07154 | 3286.47086 |
| *VPS27* | 0.008588533 | 0.95702237 | 0.99979869 | 13550.872 | 13631.7809 |
| *SNO1* | 0.012636125 | 0.95711793 | 0.99979869 | 1073.06596 | 1082.50569 |
| *CDC5* | 0.008619254 | 0.95722375 | 0.99979869 | 11636.9125 | 11706.6429 |
| *MLH2* | 0.009266283 | 0.9572278 | 0.99979869 | 4965.36884 | 4997.3629 |
| *THI5* | 0.03590703 | 0.9572763 | 0.99979869 | 79.9921895 | 82.0080065 |
| *BUD2* | 0.008983977 | 0.95748738 | 0.99979869 | 6254.0235 | 6293.0894 |
| *EXO70* | 0.009088644 | 0.95763381 | 0.99979869 | 5412.15448 | 5446.35673 |
| *YPL245W* | 0.00860753 | 0.95802663 | 0.99979869 | 8891.81472 | 8945.02331 |
| *RPL6B* | 0.007969923 | 0.95818965 | 0.99979869 | 119166.901 | 119827.024 |
| *TGL2* | 0.009738771 | 0.95820247 | 0.99979869 | 2978.24579 | 2998.41774 |
| *RPS18B* | 0.007882259 | 0.95849577 | 0.99979869 | 283739.125 | 285293.554 |
| *CWC22* | 0.009223969 | 0.95879409 | 0.99979869 | 3854.25781 | 3878.97871 |
| *KRE5* | 0.008215901 | 0.95916865 | 0.99979869 | 11905.179 | 11973.169 |
| *MSS18* | 0.008785898 | 0.96072054 | 0.99979869 | 3873.7681 | 3897.43051 |
| *PET112* | 0.008854741 | 0.96082701 | 0.99979869 | 3602.57507 | 3624.75389 |
| *YKR073C* | 0.015063005 | 0.96094242 | 0.99979869 | 482.879681 | 487.947639 |
| *MRPL17* | 0.008008166 | 0.96143765 | 0.99979869 | 7564.13948 | 7606.24261 |
| *MCP2* | 0.007920325 | 0.96147021 | 0.99979869 | 8594.2828 | 8641.59369 |

| | | | | | |
|---|---|---|---|---|---|
| *INP51* | 0.007660912 | 0.96226234 | 0.99979869 | 10272.1677 | 10326.8582 |
| *ADE2* | 0.007298472 | 0.96235894 | 0.99979869 | 31699.3439 | 31860.1105 |
| *SSH1* | 0.00721151 | 0.9623939 | 0.99979869 | 60272.1638 | 60574.1889 |
| *INN1* | 0.007913619 | 0.96305907 | 0.99979869 | 5472.63638 | 5502.73724 |
| *APL2* | 0.007246892 | 0.96358328 | 0.99979869 | 14684.4199 | 14758.3659 |
| *VAC7* | 0.007293382 | 0.96394291 | 0.99979869 | 10864.3051 | 10919.3661 |
| *SNZ2* | 0.009784551 | 0.96412626 | 0.99979869 | 1420.34912 | 1430.01461 |
| *CCL1* | 0.007449185 | 0.96465022 | 0.99979869 | 6421.81199 | 6455.05522 |
| *SHS1* | 0.0068723 | 0.96522943 | 0.99979869 | 17116.3775 | 17198.1041 |
| *NUP157* | 0.006750768 | 0.96533831 | 0.99979869 | 26438.3941 | 26562.3933 |
| *IOC4* | 0.006934595 | 0.96535083 | 0.99979869 | 13077.7475 | 13140.7579 |
| *tL(CAA)M* | 0.071534234 | 0.96564013 | 0.99979869 | 11.7061741 | 12.301201 |
| *GPI14* | 0.007045129 | 0.96599529 | 0.99979869 | 7766.07098 | 7804.08692 |
| *YBL071C-B* | 0.038364836 | 0.96599757 | 0.99979869 | 42.9226383 | 44.0793035 |
| *AIM23* | 0.007336241 | 0.96604742 | 0.99979869 | 5065.84683 | 5091.67211 |
| *tT(AGU)O2* | 0.071534445 | 0.96701204 | 0.99979869 | 10.7306596 | 11.2761009 |
| *RPB11* | 0.006639594 | 0.96703859 | 0.99979869 | 11622.2798 | 11675.8899 |
| *RSC1* | 0.006731345 | 0.9670849 | 0.99979869 | 9195.19973 | 9238.20194 |
| *YAL037W* | 0.009624682 | 0.96717104 | 0.99979869 | 1091.60073 | 1098.90729 |
| *LOA1* | 0.006619595 | 0.9675328 | 0.99979869 | 9601.01377 | 9645.16667 |
| *SEC72* | 0.006502495 | 0.96754821 | 0.99979869 | 12771.4359 | 12829.1275 |
| *RLI1* | 0.006234309 | 0.96774606 | 0.99979869 | 36308.6499 | 36465.8852 |
| *GCN2* | 0.00640835 | 0.96807816 | 0.99979869 | 12333.4299 | 12388.3345 |
| *GIT1* | 0.010389582 | 0.96913093 | 0.99979869 | 705.296988 | 710.394357 |
| *PRE9* | 0.005958873 | 0.96915115 | 0.99979869 | 37410.9813 | 37565.8176 |
| *YFH7* | 0.007129678 | 0.96921313 | 0.99979869 | 3083.60135 | 3098.87755 |
| *YAR023C* | 0.00839544 | 0.96926144 | 0.99979869 | 1412.545 | 1420.78871 |
| *LSM5* | 0.006438076 | 0.96945843 | 0.99979869 | 6390.59553 | 6419.17671 |
| *YBL044W* | 0.01841906 | 0.96974036 | 0.99979869 | 161.935408 | 164.016013 |
| *BCS1* | 0.006257013 | 0.97008471 | 0.99979869 | 6946.6388 | 6976.83116 |
| *LDB17* | 0.006240239 | 0.97031787 | 0.99979869 | 6570.0902 | 6598.56923 |
| *SMC5* | 0.006361167 | 0.97040422 | 0.99979869 | 5300.94583 | 5324.36982 |
| *CTM1* | 0.006860311 | 0.97077193 | 0.99979869 | 2848.50236 | 2862.07943 |
| *AKR2* | 0.005982047 | 0.97082651 | 0.99979869 | 8850.84311 | 8887.61771 |
| *CTK1* | 0.006158342 | 0.97094988 | 0.99979869 | 6033.55722 | 6059.36658 |
| *MRPL8* | 0.005783893 | 0.97114264 | 0.99979869 | 12678.762 | 12729.6928 |
| *SCO1* | 0.006018246 | 0.97127658 | 0.99979869 | 6812.0178 | 6840.49285 |
| *ARP7* | 0.005648694 | 0.97141483 | 0.99979869 | 17111.5 | 17178.6272 |
| *RTT103* | 0.00577686 | 0.97175779 | 0.99979869 | 9152.2771 | 9188.99713 |
| *IRC8* | 0.005808862 | 0.97214979 | 0.99979869 | 7163.20302 | 7192.10217 |
| *PDX1* | 0.005439326 | 0.97239469 | 0.99979869 | 18387.4729 | 18456.927 |
| *PFF1* | 0.005380471 | 0.97267691 | 0.99979869 | 18676.2252 | 18746.0052 |
| *INP2* | 0.006519752 | 0.97277993 | 0.99979869 | 2546.09286 | 2557.6247 |
| *MRPL33* | 0.005679975 | 0.97286285 | 0.99979869 | 6887.13241 | 6914.30005 |
| *BSP1* | 0.005557101 | 0.9731286 | 0.99979869 | 7922.1533 | 7952.72643 |

| | | | | | |
|---|---|---|---|---|---|
| *MRPL1* | 0.005524754 | 0.97323919 | 0.99979869 | 8092.86834 | 8123.91815 |
| *PAF1* | 0.005264111 | 0.9732528 | 0.99979869 | 18948.3938 | 19017.6567 |
| *LAM5* | 0.005246322 | 0.97408823 | 0.99979869 | 10644.8143 | 10683.5931 |
| *MIL1* | 0.005138142 | 0.97442263 | 0.99979869 | 12146.1311 | 12189.4651 |
| *BIL1* | 0.006071946 | 0.97472249 | 0.99979869 | 2507.07228 | 2517.6458 |
| *YIL156W-B* | 0.005397298 | 0.97473695 | 0.99979869 | 5595.55121 | 5616.52335 |
| *YOS9* | 0.00512915 | 0.97525485 | 0.99979869 | 7698.76048 | 7726.17932 |
| *MSD1* | 0.005176322 | 0.97556231 | 0.99979869 | 6076.47986 | 6098.32039 |
| *VPS25* | 0.005611903 | 0.97576956 | 0.99979869 | 3079.6993 | 3091.70185 |
| *DNM1* | 0.004894552 | 0.97591077 | 0.99979869 | 10071.2118 | 10105.4366 |
| *FAR8* | 0.004772467 | 0.97641963 | 0.99979869 | 10700.4186 | 10735.8732 |
| *CTR3* | 0.006324016 | 0.97703507 | 0.99979869 | 1368.64685 | 1374.65921 |
| *ATG27* | 0.004445275 | 0.97735572 | 0.99979869 | 20268.2649 | 20330.8099 |
| *RPL25* | 0.004280758 | 0.97744612 | 0.99979869 | 274466.859 | 275282.426 |
| *SHG1* | 0.005982828 | 0.97746188 | 0.99979869 | 1581.30901 | 1587.88003 |
| *UPS1* | 0.004738542 | 0.97765083 | 0.99979869 | 6015.99796 | 6035.78928 |
| *SUP11* | 0.028463407 | 0.97772666 | 0.99979869 | 33.1674932 | 33.8283027 |
| *YPL152W-A* | 0.028463407 | 0.97772666 | 0.99979869 | 33.1674932 | 33.8283027 |
| *ATF1* | 0.004507235 | 0.97779346 | 0.99979869 | 10229.2451 | 10261.2518 |
| *MED7* | 0.00473526 | 0.97799442 | 0.99979869 | 5239.48841 | 5256.71322 |
| *ATG17* | 0.005362826 | 0.97811537 | 0.99979869 | 2263.19365 | 2271.62178 |
| *YBR071W* | 0.004387672 | 0.97816787 | 0.99979869 | 12034.9225 | 12071.5786 |
| *IGO1* | 0.004115375 | 0.9792377 | 0.99979869 | 15855.0373 | 15900.3274 |
| *GPB2* | 0.004037559 | 0.97972267 | 0.99979869 | 14364.4511 | 14404.7063 |
| *SMF1* | 0.004037284 | 0.97978881 | 0.99979869 | 13467.9533 | 13505.6936 |
| *MDR1* | 0.003967874 | 0.98014692 | 0.99979869 | 13326.5037 | 13363.2047 |
| *PUT4* | 0.006703875 | 0.98070281 | 0.99979869 | 643.839574 | 646.838152 |
| *THI21* | 0.004779006 | 0.98093724 | 0.99979869 | 2028.09466 | 2034.82366 |
| *MPS1* | 0.003924021 | 0.98126636 | 0.99979869 | 6818.8464 | 6837.41755 |
| *MUM3* | 0.005052396 | 0.98130106 | 0.99979869 | 1473.0269 | 1478.19432 |
| *TTI2* | 0.004176147 | 0.98196765 | 0.99979869 | 3079.6993 | 3088.62655 |
| *RSC8* | 0.003435771 | 0.98214527 | 0.99979869 | 45366.3021 | 45474.4647 |
| *HSM3* | 0.003628021 | 0.98260502 | 0.99979869 | 7150.52133 | 7168.52487 |
| *IES1* | 0.003445038 | 0.98276174 | 0.99979869 | 13332.3568 | 13364.2298 |
| *MRPL15* | 0.003561031 | 0.98283447 | 0.99979869 | 7609.98866 | 7628.79481 |
| *snR24* | 0.007399726 | 0.98336247 | 0.99979869 | 336.552505 | 338.283027 |
| *MNN2* | 0.003149119 | 0.98374363 | 0.99979869 | 32136.3744 | 32206.5944 |
| *RPL8B* | 0.003081514 | 0.98374653 | 0.99979869 | 421327.642 | 422228.473 |
| *MCD1* | 0.003456064 | 0.98375964 | 0.99979869 | 5799.43374 | 5813.34256 |
| *YLR050C* | 0.003481237 | 0.98384675 | 0.99979869 | 5166.32482 | 5178.80561 |
| *COF1* | 0.003061393 | 0.98398547 | 0.99979869 | 75410.1979 | 75570.378 |
| *ATG15* | 0.00336906 | 0.98404074 | 0.99979869 | 6278.41136 | 6293.0894 |
| *tD(GUC)O* | 0.006860618 | 0.98421857 | 0.99979869 | 356.062795 | 357.759929 |
| *MOB2* | 0.00308792 | 0.98509572 | 0.99979869 | 7727.0504 | 7743.60602 |
| *SPC29* | 0.004040643 | 0.98515868 | 0.99979869 | 1430.10427 | 1434.11501 |

| | | | | | |
|---|---|---|---|---|---|
| *RIF1* | 0.002947992 | 0.98534702 | 0.99979869 | 11788.1173 | 11812.2282 |
| *HEM13* | 0.002803435 | 0.98566022 | 0.99979869 | 22835.8191 | 22880.2338 |
| *USE1* | 0.003066832 | 0.98575265 | 0.99979869 | 5219.97812 | 5231.08572 |
| *TRM5* | 0.002763435 | 0.98604993 | 0.99979869 | 16032.5809 | 16063.3183 |
| *MGA2* | 0.002659706 | 0.98637799 | 0.99979869 | 23793.7743 | 23837.6773 |
| *tH(GUG)E1* | 0.019065081 | 0.98642245 | 0.99979869 | 27.3144062 | 27.6777022 |
| *RPS14A* | 0.002546053 | 0.98658334 | 0.99979869 | 273553.778 | 274036.93 |
| *JLP1* | 0.003461622 | 0.98662802 | 0.99979869 | 1756.90163 | 1761.12194 |
| *NSG1* | 0.002554765 | 0.98697451 | 0.99979869 | 20686.7606 | 20723.4233 |
| *BZZ1* | 0.002606767 | 0.98697793 | 0.99979869 | 12902.1549 | 12925.4869 |
| *RHO4* | 0.002832557 | 0.98710839 | 0.99979869 | 4405.42351 | 4414.08095 |
| *CIT3* | 0.00353955 | 0.98712575 | 0.99979869 | 1377.42648 | 1380.80981 |
| *SWI6* | 0.002467646 | 0.9877976 | 0.99979869 | 10810.6518 | 10829.1573 |
| *SRC1* | 0.002366055 | 0.98789324 | 0.99979869 | 23069.9426 | 23107.806 |
| *YJR098C* | 0.002841408 | 0.98792041 | 0.99979869 | 2811.43281 | 2816.97502 |
| *YOL085C* | 0.004415757 | 0.9884628 | 0.99979869 | 493.61034 | 495.12334 |
| *FRQ1* | 0.002303379 | 0.98865479 | 0.99979869 | 10160.9591 | 10177.1936 |
| *SUF23* | 0.014949128 | 0.98973056 | 0.99979869 | 25.3633772 | 25.627502 |
| *YCR022C* | 0.005256979 | 0.98976563 | 0.99979869 | 239.001054 | 239.873419 |
| *ATP22* | 0.00173275 | 0.99180489 | 0.99979869 | 6175.98234 | 6183.40369 |
| *tN(GUU)N1* | 0.005642744 | 0.99226633 | 0.99979869 | 109.257625 | 109.685709 |
| *LSM4* | 0.001512327 | 0.99250675 | 0.99979869 | 11160.8615 | 11172.5658 |
| *FLC2* | 0.001428353 | 0.99266216 | 0.99979869 | 26388.6429 | 26414.7789 |
| *YER160C* | 0.001781731 | 0.99346101 | 0.99979869 | 1426.20221 | 1427.96441 |
| *MRPS28* | 0.001332834 | 0.99348256 | 0.99979869 | 9140.57092 | 9149.01823 |
| *TOR2* | 0.001272908 | 0.99356654 | 0.99979869 | 16459.8563 | 16474.3834 |
| *MDV1* | 0.001256143 | 0.99383083 | 0.99979869 | 9725.87963 | 9734.35038 |
| *HEM25* | 0.001227615 | 0.9943469 | 0.99979869 | 4840.50298 | 4844.62299 |
| *MUS81* | 0.001289435 | 0.99436825 | 0.99979869 | 3305.04315 | 3307.99796 |
| *PFA3* | 0.001255352 | 0.99442812 | 0.99979869 | 3672.81212 | 3676.00889 |
| *BRR6* | 0.001239835 | 0.9947229 | 0.99979869 | 2830.9431 | 2833.37663 |
| *RPS20* | 0.000915842 | 0.995168 | 0.99979869 | 459377.585 | 459669.228 |
| *RGD2* | 0.000857002 | 0.99574243 | 0.99979869 | 11672.0311 | 11678.9652 |
| *SUI1* | 0.000818621 | 0.99575852 | 0.99979869 | 38056.7719 | 38078.3676 |
| *YOP1* | 0.000789984 | 0.99589946 | 0.99979869 | 41976.3892 | 41999.3755 |
| *FAD1* | 0.000781165 | 0.99625891 | 0.99979869 | 7052.96988 | 7056.78896 |
| *TFG2* | 0.00065627 | 0.99669556 | 0.99979869 | 15119.4993 | 15126.3768 |
| *YMR001C-A* | 0.001025891 | 0.99709821 | 0.99979869 | 613.598624 | 614.034949 |
| *LOS1* | 0.000575169 | 0.99711638 | 0.99979869 | 13822.065 | 13827.575 |
| *GEF1* | 0.000542333 | 0.99729556 | 0.99979869 | 12474.8795 | 12479.5684 |
| *ADD66* | 0.000536905 | 0.9974558 | 0.99979869 | 6297.92165 | 6300.2651 |
| *VPS33* | 0.000516611 | 0.99755999 | 0.99979869 | 6093.06361 | 6095.24509 |
| *HSL7* | 0.000459914 | 0.99770311 | 0.99979869 | 12826.0647 | 12830.1526 |
| *SAG1* | 0.000507204 | 0.99770432 | 0.99979869 | 4223.97781 | 4225.46254 |
| *MPH1* | 0.000453275 | 0.99773647 | 0.99979869 | 12804.6034 | 12808.6255 |

| | | | | | |
|---|---|---|---|---|---|
| *SRP54* | 0.000299928 | 0.9984428 | 0.99999915 | 42545.1142 | 42553.9546 |
| *SIS2* | 0.000247748 | 0.99873682 | 0.99999915 | 20660.4217 | 20663.9674 |
| *YNL035C* | 0.00024213 | 0.99881503 | 0.99999915 | 9246.902 | 9248.45294 |
| *DPB11* | 0.000244126 | 0.99888182 | 0.99999915 | 4633.6939 | 4634.47747 |
| *APL1* | 0.000177753 | 0.99911238 | 0.99999915 | 12796.7993 | 12798.3745 |
| *CFT1* | 0.000161954 | 0.99919381 | 0.99999915 | 12086.6247 | 12087.9802 |
| *YDL129W* | 0.000138779 | 0.99936081 | 0.99999915 | 4849.28261 | 4849.74849 |
| *FLX1* | 0.000105643 | 0.99952744 | 0.99999915 | 3877.67016 | 3877.95361 |

**Supplementary Table 2: List of strains used in this study**

| Strains | Genotype | Source |
|---|---|---|
| CSY064 | *MATa his3Δ1 leu2Δ0 met15Δ0 ura3Δ0* | This Study |
| CSY032 | *MATα SUC2 gal2 mal2 mel flo1 hap1 ho bio1 bio6* | This Study |
| CSY036 | *MATα SUC2 gal2 mal2 mel flo1 hap1 ho bio1 bio6 ykl075cΔ::kanMX-6* | This Study |
| CSY046 | *MATa his3Δ1 leu2Δ0 met15Δ0 ura3Δ0 TOM70-mCherry hphMX-4 [pmito-roGFP1:URA3]* | This Study |
| CSY065 | *MATα SUC2 gal2 mal2 mel flo1 hap1 ho bio1 bio6 tor1Δ::hphMX-4* | This Study |
| CSY074 | *MATα SUC2 gal2 mal2 mel flo1 hap1 ho bio1 bio6 ykl075cΔ::kanMX-6 tor1Δ::hphMX-4* | This Study |
| CSY093 | *MATα SUC2 gal2 mal2 mel flo1 hap1 ho bio1 bio6 ykl075cΔ::hphMX-4 [pmitoHo-roGFP1:KanMX]* | This Study |
| CSY104 | *MATα SUC2 gal2 mal2 mel flo1 hap1 ho bio1 bio6 [pmitoHo-roGFP1:KanMX]* | This Study |
| CSY107 | *MATα SUC2 gal2 mal2 mel flo1 hap1 ho bio1 bio6 ykl075cΔ::hphMX-4 [pmitoHo-roGFP1:KanMX]* | This Study |
| CSY113 | *MATα SUC2 gal2 mal2 mel flo1 hap1 ho bio1 bio6 ykl075cΔ::hphMX-4 ABP140-GFP (S65T) kanMX-6* | This Study |
| CSY125 | *MATα SUC2 gal2 mal2 mel flo1 hap1 ho bio1 bio6 ABP140-GFP (S65T) kanMX-6* | This Study |
| CSY152 | *MATa his3Δ1 leu2Δ0 met15Δ0 ura3Δ0 ABP140-GFP (S65T) kanMX-6* | This Study |
| CSY154 | *MATa his3Δ1 leu2Δ0 met15Δ0 ura3Δ0 act1-V159N [pML104]* | This Study |
| CSY160 | *MATα SUC2 gal2 mal2 mel flo1 hap1 ho bio1 bio6 bat1Δ::hphMX-4* | This Study |
| CSY164 | *MATα SUC2 gal2 mal2 mel flo1 hap1 ho bio1 bio6 ykl075cΔ::kanMX-6 bat1Δ::hphMX-4* | This Study |
| CSY183 | *MATa his3Δ1 leu2Δ0 met15Δ0 ura3Δ0 act1-V159N [pML104] [pmito-roGFP1:URA3]* | This Study |
| CSY186 | *MATa his3Δ1 leu2Δ0 met15Δ0 ura3Δ0 ykl075cΔ::LEU2* | This Study |
| CSY187 | *MATa his3Δ1 leu2Δ0 met15Δ0 ura3Δ0 act1-V159N ABP140-ENVY his3 [pML104]* | This Study |
| CSY225 | *MATα SUC2 gal2 mal2 mel flo1 hap1 ho bio1 bio6 YKL075C-13Myc-kanMX-6* | This Study |
| CSY258 | MATα SUC2 gal2 mal2 mel flo1 hap1 ho bio1 bio6 ykl075cΔ::hphMX1 Ho-promoter-ykl075c-13xmyc-adh1terminator-Ho::KanMX | This Study |
| CSY265 | MATα SUC2 gal2 mal2 mel flo1 hap1 ho bio1 bio6 ykl075cΔ::hphMX1 Ho-promoter-ykl075c-adh1terminator-Ho::KanMX | This Study |
| EGS164 | *MATa his3Δ1 leu2Δ0 met15Δ0 ura3Δ0 ABP140-ENVY his3* | This Study |
| RHY055 | *MATa his3Δ1 leu2Δ0 met15Δ0 ura3Δ0 ykl075cΔ::LEU2* | This Study |
| CTY056 | *MATα SUC2 gal2 mal2 mel flo1 hap1 ho bio1 bio6 bat1Δ::hphMX-4 [pmitoHo-roGFP1:KanMX]* | This Study |

**Supplementary Table 3: List of primers and plasmids used in this study**

| Primer Name | Plasmid | Primer Sequence 5' - 3' | Source |
|---|---|---|---|
| FW *YKL075C* deletion pFA6 | pFA6a-kanMX6 Addgene #39296 | TAACATTATAAATCAAGCCTAGTATAGCGCGCCAAAGAAGcggatccccgggttaattaa | This Study |
| RV *YKL075C* deletion pFA6 | pFA6a-kanMX6 Addgene #39296 | AGAAAACGCTGGATCTTGCTTCTTTTCCTTTGATTTCTCCgaattcgagctcgtttaaac | This Study |
| FW *YKL075C* deletion PCY | pCY 3090-07 Addgene #36232 | TAACATTATAAATCAAGCCTAGTATAGCGCGCCAAAGAAGgtttagcttgcctcgtcc | This Study |
| RV *YKL075C* deletion PCY | pCY 3090-07 Addgene #36232 | AGAAAACGCTGGATCTTGCTTCTTTTCCTTTGATTTCTCCatcgatgaattcgagctcg | This Study |
| FW *ABP140* Tag pFA6 GFP | pFA6a-GFP(S65T)-KanMX6 Addgene #39292 | GTACCGCTGCTGGGTACAAGCTGTGTTTGACGTTCCTCAAcggatccccgggttaattaa | This Study |
| RV *ABP140* Tag pFA6 GFP | pFA6a-GFP(S65T)-KanMX6 Addgene #39292 | TTTATGATGAGAGAGGAGGTGGTACTTGTCTCAGAACTTCgaattcgagctcgtttaaac | This Study |
| FW *BAT1* deletion PCY | pCY 3090-07 Addgene #36232 | TATAAACGCAAAATCAGCTAGAACCTTAGCATACTAAAACgtttagcttgcctcgtcc | This Study |
| RV *BAT1* deletion PCY | pCY 3090-07 Addgene #36232 | TTTTTTTTGGGGGGGGAGGGGATGTTTACCTTCATTATCAtcgatgaattcgagctcg | This Study |
| FW *YKL075C* deletion pOM | pOM13 Euroscarf #P30388 | TAACATTATAAATCAAGCCTAGTATAGCGCGCCAAAGAAGtgcaggtcgacaaccttaat | This Study |
| RV *YKL075C* deletion pOM | pOM13 Euroscarf #P30388 | AGAAAACGCTGGATCTTGCTTCTTTTCCTTTGATTTCTCCgcagcgtacggatatcaccta | This Study |
| FW *ABP140* Tag pFA6 GFPEnvy-linker | pFA6a-link-GFPEnvy-SpHis5 Addgene #60782 | GTACCGCTGCTGGGTACAAGCTGTGTTTGACGTTCCTCAAggtgacggtgctggtttaattaac | This Study |
| FW *YKL075C* pFA6 13Myc Tag | pFA6a-13Myc-kanMX6 Addgene #41602 | GGAAAAAATATGTGAAGTTGCTGCAAGACAGAAATGGATAcggatccccgggttaattaa | This Study |
| RV *YKL075C* pFA6 13Myc Tag | pFA6a-13Myc-kanMX6 Addgene #41602 | AGAAAACGCTGGATCTTGCTTCTTTTCCTTTGATTTCTCCgattcgagctcgtttaaac | This Study |
| FW Ho mito-roGFP1 | HO-poly-KanMX4-HO Adddgene#51662 | GCCGCCAGCTGAAGCTTAATTAT | This Study |
| RV Ho mito-roGFP1 | HO-poly-KanMX4-HO Adddgene#51662 | CATAGGCCACTGTAAGATTCCGCC | This Study |
| FW *YKL075C* rescue pFA6 | pFA6a-13Myc-kanMX6 Addgene #41602 | CAAGACAGAAATGGATATAAaacggtgaacaaaaattaatcaat | This Study |
| RV *YKL075C* rescue pFA6 | pFA6a-13Myc-kanMX6 Addgene #41602 | CCTTAATTAACCCGGGGATcccggtagaggtgtggtcaat | This Study |
| FW *YKL075C* rescue | N/A | acgctgcaggtcgacggatcACCCGGCGTCGATTACAGAC | This Study |
| RV *YKL075C r*escue | N/A | attaatttttgttcaccgttTTATATCCATTTCTGTCTTGCAGCA | This Study |
| FW *BAT1* RT-qPCR | N/A | ACCCAAATCCATCCAAGCCA | This Study |
| RV *BAT1* RT-qPCR | N/A | CACCCTTCTTTGGCTGACCA | This Study |
| FW *BAT2* RT-qPCR | N/A | TTTCCACGCCTGATAGAGCC | This Study |
| RV *BAT2* RT-qPCR | N/A | CAGTGGCTTCCAGTCTGACC | This Study |
| FW *ACT1* RT-qPCR | N/A | TCGCCTTGGACTTCGAACAA | This Study |
| RV *ACT1* RT-qPCR | N/A | CAAAGCTTCTGGGGCTCTGA | This Study |

**Supplementary Table 2: List of strains used in this study**

| Strains | Genotype | Source |
|---|---|---|
| CSY064 | *MATa his3Δ1 leu2Δ0 met15Δ0 ura3Δ0* | This Study |
| CSY032 | *MATα SUC2 gal2 mal2 mel flo1 hap1 ho bio1 bio6* | This Study |
| CSY036 | *MATα SUC2 gal2 mal2 mel flo1 hap1 ho bio1 bio6 ykl075cΔ::kanMX-6* | This Study |
| CSY046 | *MATa his3Δ1 leu2Δ0 met15Δ0 ura3Δ0 TOM70-mCherry hphMX-4 [pmito-roGFP1:URA3]* | This Study |
| CSY065 | *MATα SUC2 gal2 mal2 mel flo1 hap1 ho bio1 bio6 tor1Δ::hphMX-4* | This Study |
| CSY074 | *MATα SUC2 gal2 mal2 mel flo1 hap1 ho bio1 bio6 ykl075cΔ::kanMX-6 tor1Δ::hphMX-4* | This Study |
| CSY093 | *MATα SUC2 gal2 mal2 mel flo1 hap1 ho bio1 bio6 ykl075cΔ::hphMX-4 [pmitoHo-roGFP1:KanMX]* | This Study |
| CSY104 | *MATα SUC2 gal2 mal2 mel flo1 hap1 ho bio1 bio6 [pmitoHo-roGFP1:KanMX]* | This Study |
| CSY107 | *MATα SUC2 gal2 mal2 mel flo1 hap1 ho bio1 bio6 ykl075cΔ::hphMX-4 [pmitoHo-roGFP1:KanMX]* | This Study |
| CSY113 | *MATα SUC2 gal2 mal2 mel flo1 hap1 ho bio1 bio6 ykl075cΔ::hphMX-4 ABP140-GFP (S65T) kanMX-6* | This Study |
| CSY125 | *MATα SUC2 gal2 mal2 mel flo1 hap1 ho bio1 bio6 ABP140-GFP (S65T) kanMX-6* | This Study |
| CSY152 | *MATa his3Δ1 leu2Δ0 met15Δ0 ura3Δ0 ABP140-GFP (S65T) kanMX-6* | This Study |
| CSY154 | *MATa his3Δ1 leu2Δ0 met15Δ0 ura3Δ0 act1-V159N [pML104]* | This Study |
| CSY160 | *MATα SUC2 gal2 mal2 mel flo1 hap1 ho bio1 bio6 bat1Δ::hphMX-4* | This Study |
| CSY164 | *MATα SUC2 gal2 mal2 mel flo1 hap1 ho bio1 bio6 ykl075cΔ::kanMX-6 bat1Δ::hphMX-4* | This Study |
| CSY183 | *MATa his3Δ1 leu2Δ0 met15Δ0 ura3Δ0 act1-V159N [pML104] [pmito-roGFP1:URA3]* | This Study |
| CSY186 | *MATa his3Δ1 leu2Δ0 met15Δ0 ura3Δ0 ykl075cΔ::LEU2* | This Study |
| CSY187 | *MATa his3Δ1 leu2Δ0 met15Δ0 ura3Δ0 act1-V159N ABP140-ENVY his3 [pML104]* | This Study |
| CSY225 | *MATα SUC2 gal2 mal2 mel flo1 hap1 ho bio1 bio6 YKL075C-13Myc-kanMX-6* | This Study |
| CSY258 | MATα SUC2 gal2 mal2 mel flo1 hap1 ho bio1 bio6 ykl075cΔ::hphMX1 Ho-promoter-ykl075c-13xmyc-adh1terminator-Ho::KanMX | This Study |
| CSY265 | MATα SUC2 gal2 mal2 mel flo1 hap1 ho bio1 bio6 ykl075cΔ::hphMX1 Ho-promoter-ykl075c-adh1terminator-Ho::KanMX | This Study |
| EGS164 | *MATa his3Δ1 leu2Δ0 met15Δ0 ura3Δ0 ABP140-ENVY his3* | This Study |
| RHY055 | *MATa his3Δ1 leu2Δ0 met15Δ0 ura3Δ0 ykl075cΔ::LEU2* | This Study |
| CTY056 | *MATα SUC2 gal2 mal2 mel flo1 hap1 ho bio1 bio6 bat1Δ::hphMX-4 [pmitoHo-roGFP1:KanMX]* | This Study |

**Supplementary Table 3: List of primers and plasmids used in this study**

| Primer Name | Plasmid | Primer Sequence 5' - 3' | Source |
|---|---|---|---|
| FW *YKL075C* deletion pFA6 | pFA6a-kanMX6 Addgene #39296 | TAACATTATAAATCAAGCCTAGTATAGCGCGCCAAAGAAGcggatccccgggttaattaa | This Study |
| RV *YKL075C* deletion pFA6 | pFA6a-kanMX6 Addgene #39296 | AGAAAACGCTGGATCTTGCTTCTTTTCCTTTGATTTCTCCgaattcgagctcgtttaaac | This Study |
| FW *YKL075C* deletion PCY | pCY 3090-07 Addgene #36232 | TAACATTATAAATCAAGCCTAGTATAGCGCGCCAAAGAAGgtttagcttgcctcgtcc | This Study |
| RV *YKL075C* deletion PCY | pCY 3090-07 Addgene #36232 | AGAAAACGCTGGATCTTGCTTCTTTTCCTTTGATTTCTCCatcgatgaattcgagctcg | This Study |
| FW *ABP140* Tag pFA6 GFP | pFA6a-GFP(S65T)-KanMX6 Addgene #39292 | GTACCGCTGCTGGGTACAAGCTGTGTTTGACGTTCCTCAAcggatccccgggttaattaa | This Study |
| RV *ABP140* Tag pFA6 GFP | pFA6a-GFP(S65T)-KanMX6 Addgene #39292 | TTTATGATGAGAGAGGAGGTGGTACTTGTCTCAGAACTTCgaattcgagctcgtttaaac | This Study |
| FW *BAT1* deletion PCY | pCY 3090-07 Addgene #36232 | TATAAACGCAAAATCAGCTAGAACCTTAGCATACTAAAACgtttagcttgcctcgtcc | This Study |
| RV *BAT1* deletion PCY | pCY 3090-07 Addgene #36232 | TTTTTTTTGGGGGGGGAGGGGATGTTTACCTTCATTATCAtcgatgaattcgagctcg | This Study |
| FW *YKL075C* deletion pOM | pOM13 Euroscarf #P30388 | TAACATTATAAATCAAGCCTAGTATAGCGCGCCAAAGAAGtgcaggtcgacaacccttaat | This Study |
| RV *YKL075C* deletion pOM | pOM13 Euroscarf #P30388 | AGAAAACGCTGGATCTTGCTTCTTTTCCTTTGATTTCTCCgcagcgtacggatatcaccta | This Study |
| FW *ABP140* Tag pFA6 GFPEnvy-linker | pFA6a-link-GFPEnvy-SpHis5 Addgene #60782 | GTACCGCTGCTGGGTACAAGCTGTGTTTGACGTTCCTCAAggtgacggtgctggtttaattaac | This Study |
| FW *YKL075C* pFA6 13Myc Tag | pFA6a-13Myc-kanMX6 Addgene #41602 | GGAAAAAATATGTGAAGTTGCTGCAAGACAGAAATGGATAcggatccccgggttaattaa | This Study |
| RV *YKL075C* pFA6 13Myc Tag | pFA6a-13Myc-kanMX6 Addgene #41602 | AGAAAACGCTGGATCTTGCTTCTTTTCCTTTGATTTCTCCgattcgagctcgtttaaac | This Study |
| FW Ho mito-roGFP1 | HO-poly-KanMX4-HO Adddgene#51662 | GCCGCCAGCTGAAGCTTAATTAT | This Study |
| RV Ho mito-roGFP1 | HO-poly-KanMX4-HO Adddgene#51662 | CATAGGCCACTGTAAGATTCCGCC | This Study |
| FW *YKL075C* rescue pFA6 | pFA6a-13Myc-kanMX6 Addgene #41602 | CAAGACAGAAATGGATATAAaacggtgaacaaaaattaatcaat | This Study |
| RV *YKL075C* rescue pFA6 | pFA6a-13Myc-kanMX6 Addgene #41602 | CCTTAATTAACCCGGGGATcccggtagaggtgtggtcaat | This Study |
| FW *YKL075C* rescue | N/A | acgctgcaggtcgacggatcACCCGGCGTCGATTACAGAC | This Study |
| RV *YKL075C r*escue | N/A | attaatttttgttcaccgttTTATATCCATTTCTGTCTTGCAGCA | This Study |
| FW *BAT1* RT-qPCR | N/A | ACCCAAATCCATCCAAGCCA | This Study |
| RV *BAT1* RT-qPCR | N/A | CACCCTTCTTTGGCTGACCA | This Study |
| FW *BAT2* RT-qPCR | N/A | TTTCCACGCCTGATAGAGCC | This Study |
| RV *BAT2* RT-qPCR | N/A | CAGTGGCTTCCAGTCTGACC | This Study |
| FW *ACT1* RT-qPCR | N/A | TCGCCTTGGACTTCGAACAA | This Study |
| RV *ACT1* RT-qPCR | N/A | CAAAGCTTCTGGGGCTCTGA | This Study |
